# Post-transcriptional modifications on tRNA fragments confer functional changes to high-density lipoproteins in atherosclerosis

**DOI:** 10.1101/2024.12.19.628772

**Authors:** Elizabeth M. Semler, Danielle L. Michell, Philip J. Kingsley, Clark Massick, Marisol A. Ramirez, Mark A. Castleberry, Amanda C. Doran, John J. Carr, Lawrence J. Marnett, Quanhu Sheng, MacRae F. Linton, Kasey C. Vickers

**Author notes:** Corresponding Author: Kasey C. Vickers, PhD Vanderbilt University Medical Center 2220 Pierce Ave. 312 Preston Research Building Nashville, TN 37232.

## Abstract

Epitranscriptomic modifications on RNA play critical roles in stability, processing, and function, partly by influencing interactions with RNA-binding proteins and receptors. The role of post-transcriptional RNA modifications on cell-free non-coding small RNA (sRNA) remains poorly understood in disease contexts. High-density lipoproteins (HDL), which transport sRNAs, can lose their beneficial properties in atherosclerosis cardiovascular disease (ASCVD). We hypothesize that changes to regulatory modifications on HDL-sRNAs contribute to this dysfunction. To assess changes in HDL-sRNA modification status, HDL-derived RNA from healthy subjects and those with atherosclerotic lesion development were analyzed using LC-MS/MS and AlkB-facilitated RNA (de)Methylation Sequencing. ASVD-HDL showed an enrichment in modified nucleosides including m^1^A tRNA-derived sRNAs (tDRs), particularly tDR-ArgACG-1. Functional studies revealed that ASCVD-HDL induced cell adhesion genes, including TMEM123, in primary macrophages. Recombinant HDL loaded with m^1^A-tDR-ArgACG-1 induced immune signaling, and similarly upregulated TMEM123. These findings suggest HDL-delivered-m^1^A-tDR-ArgACG-1 act on adhesion genes and immune pathways, promoting macrophage activation.

## INTRODUCTION

Atherosclerotic cardiovascular disease (ASCVD) is a complex, multi-faceted disease, resulting mainly from hypercholesterolemia-associated accumulation of apolipoprotein B containing lipoproteins (e.g., low-density lipoproteins) in the arterial wall. In response to LDL particles and other inflammatory stimuli, the atherosclerotic lesion becomes a site of chronic inflammation. Historically, HDL particles have been reported to antagonize atherosclerosis-associated inflammation by accepting excess cholesterol from lesion immune cells (macrophage foam cells) and dampening inflammatory responses to lesion stimuli^1^. In an active inflammatory setting, however, HDL often lose these beneficial properties and adopt pro-inflammatory, pro-atherogenic features^2^. Disease-associated changes in HDL function are likely mediated by changes to HDL cargo and its ability to interact with receptors, deliver cargo, drive cell signaling, or be taken up by phagocytes. HDL transport many different types of cargo, including cell-free small non-coding RNAs (sRNA) of host and microbial origin^3–5^. Here, we sought to define the direct impact of disease HDL on macrophages and determine if changes to RNA modifications on HDL-sRNA cargo may contribute to these newly discovered functions.

We posit that HDL functionality is predominantly shaped, not by its ability to accept more cholesterol, but by its ability to deliver non-cholesterol cargo to immune cells, including, but not limited to, sRNAs. HDL transport many classes of non-coding sRNAs, including fragments of parent transcripts of rRNAs (rDR), lncRNAs (lncDR), snRNAs (snDR), snoRNAs (snoDR), Y RNAs (yDR), and tRNAs (tDR)^5^. The most widely studied class of functional sRNAs on circulating HDL are microRNAs (miRNA)^3,4^. Nevertheless, miRNAs are only a small fraction of sRNAs transported by HDL, and the many other sRNA classes, including HDL-tDRs, are ripe for investigation. tDRs are generated by the targeted cleavage of full-length mature tRNAs through specific RNases, a process that can occur under normal cellular conditions but is often upregulated during stress responses^6–9^. These tDR fragments are involved in a variety of cellular processes, including the regulation of protein translation, gene silencing, apoptosis, cell proliferation and innate immune responses^10–17^. Moreover, multiple cardiovascular diseases have previously been linked to alterations in cellular tDR abundance^18–21^. tDR formation and function is further regulated by post-transcriptional RNA modifications present on parental tRNAs. Mature tRNAs are highly decorated with extensive modifications that influence their stability and function^22,23^. These modifications provide positional cues that direct RNase cleavage of tRNAs at specific locations, resulting in tDRs with inherited modifications^24–32^. Modified tDRs have been reported to dynamically respond to shifts in cellular metabolism or environmental stresses, and their dysregulation can result in significant biological consequences^26,27,31,33–35^. Upon resolution of the cellular stress and the return of protein synthesis, cells likely respond by secreting cleaved tDR fragments through various efflux routes to extracellular carriers, including HDL. Despite their biological significance, the presence and function of cell-free modified tDRs residing on circulating HDL and other lipoproteins remain to be explored, particularly in disease context. Here, we sought to define the impact of ASCVD on HDL-sRNA modifications and their contribution to HDL functionality in direct immune cell signaling.

A notable challenge in quantifying and identifying post-transcriptionally modified sRNAs arises from the fact that common RNA modifications impede first-strand synthesis by reverse transcriptase, and fail to be captured by PCR or sequencing approaches^36^. To overcome this barrier, several methods have been developed, including AlkB-mediated RNA (de)Methylase sequencing (ARM-seq) which utilizes a pool of mutated bacterial AlkB enzymes to remove specific sequencing-inhibitory methyl groups (*i.e*., m^1^A, m^3^C, m^1^G, and m^2^_2_G) from sRNAs prior to reverse transcription^37–39^. Another analytical technique for surveying the modification landscape of sRNA is liquid chromatography tandem mass spectrometry (LC-MS/MS). A key advantage of LC-MS/MS is its ability to quantify absolute levels of modified RNA nucleosides with high-sensitivity using standards. In the current study, ARM-seq and LC-MS/MS approaches were used to assess the impact of ASCVD on HDL-sRNAs and their regulatory modifications. LC-MS/MS revealed a significant enrichment in the relative levels of several HDL-RNA nucleoside modifications from subjects with advanced lesion development (CAC^+^) compared to healthy controls. ARM-seq revealed a dramatic increase in HDL-RNA reads mapping to host tDRs, which remained undetected by conventional sRNA-seq methods. Additionally, ARM-seq identified many HDL-tDRs, particularly from the 3’ terminal ends of parent tRNAs, that harbor m^1^A methylations and are up-regulated in subjects with advanced lesion development compared to healthy controls. Collectively, we highlight that HDL from ASCVD (CAC^+^) subjects transport multiple modified nucleosides that are increased in ASCVD, and that ASCVD HDL directly alters primary macrophage gene expression, inducing TMEM123 activation, independent of cholesterol flux or prior endotoxin challenge. Most importantly, we present evidence that the observed changes in macrophages responses to disease (ASCVD) HDL are likely conferred through the m^1^A modification on HDL, as recombinant HDL (rHDL) delivery of unmodified tDRs failed to induce candidate gene expression. Overall, rHDL delivery of m^1^A-tDRs to recipient macrophages recapitulated the observed responses with disease (ASCVD) CAC^+^HDL treatments, as compared to relevant controls. Results presented below support a model in which pro-inflammatory m^1^A modifications accumulate on critical tDR fragments circulating on HDL during advanced atherosclerotic disease and drive inflammatory signaling and novel receptor expression in macrophages. These sRNA methylations likely differ from other reported anti-inflammatory RNA modifications and provide a readily accessible drug target in plasma to reduce atherosclerosis-associated inflammation.

## RESULTS

### ASCVD HDL induces changes in macrophage gene expression

To determine if HDL from individuals with ASCVD (CAC^+^HDL) directly elicit inflammatory gene expression changes in macrophages without a pre-stimulus, bone marrow-derived macrophages (BMDM) were treated with physiological levels (1 mg/mL HDL total protein) of disease HDL (CAC^+^HDL) or HDL from healthy controls subjects (CAC^-^; Ctr-HDL). HDL were isolated from human plasma by density-gradient ultracentrifugation (Table 1). Total RNA (bulk) sequencing were used to quantify protein coding gene (mRNA) expression in macrophages 6h after HDL treatments. Remarkably, CAC^+^HDL directly induced changes to macrophage gene expression without a prior endotoxin challenge, as we found 269 genes to be significantly up-regulated and 88 genes down-regulated, as compared to Ctr-HDL treatments (Fig.1A, Table S1). Gene set enrichment analyses indicated that CAC^+^HDL treatments were associated with a significant enrichment in genes associated with immune cell signaling and nuclear factor κ beta (NF-κB) transactivation (Table S2,S3). Gene ontology and KEGG pathway analyses found that CAC^+^HDL induced gene expression changes in macrophages associated with cellular adhesion (GO:0045785), myeloid differentiation (GO:0030099), calcium handling (GO:0019722; KEGG:mmu04020), and efferocytosis (KEGG:mmu04148) compared to Ctr-HDL responses (Figs.1B, Tables S2,S3). Of note, multiple target genes within these altered pathways were significantly increased, including RAS guanyl-releasing protein 1 (*Rasgrp1*), proto-oncogene c-Rel (*Rel*), and OX40 ligand (*Tnfs4*), nuclear receptor 4A3 (*Nr4a3*), and *Tmem123* (Tables S1-S3). Transcription factor (TF) analyses (MetaCore) were used to identify TFs whose gene targets were significantly altered greater than chance, and multiple TFs were identified to be significantly enriched in CAC^+^ HDL-treated macrophages including the transcription factor Early Growth Receptor 1 (EGR1), which showed high activity in response to CAC^+^HDL (p=1.597x10^-10^) (Table S4). *Egr1* was also ranked as one of the most highly expressed gene network objects (mRNA) within the altered pathway maps derived from CAC^+^HDL datasets, highlighting its potential role in the transcriptional responses to CAC^+^HDL (Tables S1-S4). Notably, studies indicate that *Tmem123*, a gene increased by CAC^+^ HDL in macrophages, harbors a putative EGR1 TF response element within its promoter^40^. In orthogonal studies, *Tmem123* expression levels were confirmed to be significantly increased at the mRNA level by RT-PCR upon CAC^+^HDL treatments in BMDMs, as compared to Ctr-HDL treatments (Fig.1C). Additionally, TMEM123 protein levels in both BMDMs and human monocyte derived macrophages were slightly increased upon CAC+HDL treatments, as compared to Ctr-HDL treatments (Figs.1D,E). Although many immune related genes were confirmed by PCR to be significantly increased by disease HDL treatments in macrophages, including *Tmem123* and *Nr4a3*; classic inflammatory cytokines, interleukin 6 (*Il6*), tumor necrosis factor (*Tnf*) and interleukin 1beta (*Il1b*), were not increased upon these treatments (Fig.1C). Moreover, we failed to observe differences in responses to HDL treatments in macrophages loaded with acetylated LDL (AcLDL) or pre-stimulated with inflammatory agonists including endotoxin lipopolysaccharide (LPS) and heat-killed *Listeria monocytogenes* (HKLM), with and without AcLDL, with the exception of *Il1b* mRNA, which was increased in CAC^+^HDL treatments after pre-stimulating macrophages with LPS and AcLDL (Figs.S1A-E). These results suggest that HDL from ASCVD subjects (CAC^+^HDL) directly stimulates gene expression in primary macrophages associated with cell adhesion and immune signaling, independent of prior inflammatory challenge.

**Figure 1.**
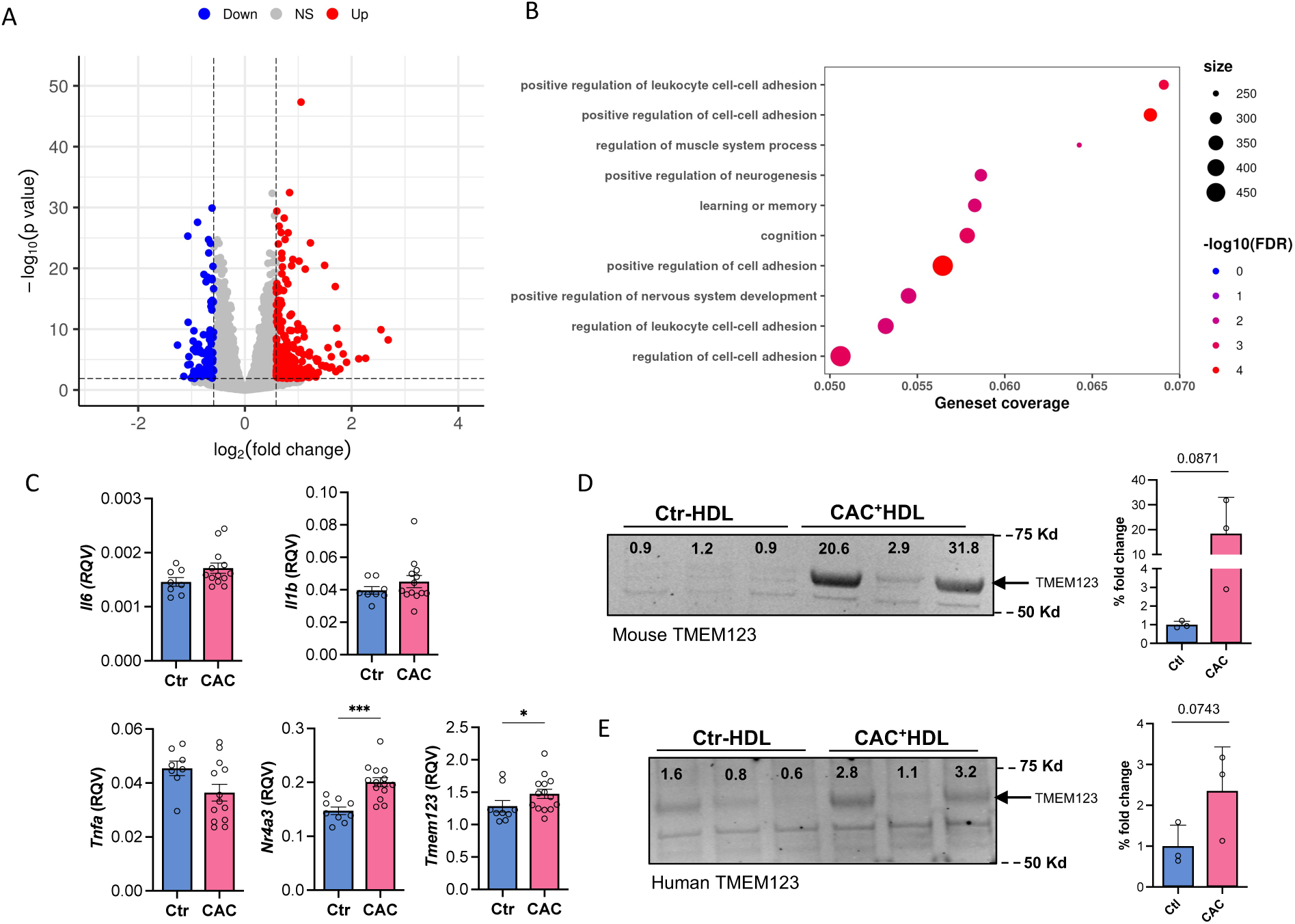
Atherosclerotic HDL induces macrophage TMEM123. (A-B) Total RNA sequencing analysis from murine bone marrow derived macrophages (BMDM) treated with HDL (1 mg/mL) from CAC^+^HDL and Ctr-HDL subjects for 6h, n=4. (A) Volcano plot demonstrating significant (adjusted p > 0.05) differential (>1.5-absolute fold change) abundances. Differential expression analysis revealed 357 genes differentially expressed between CAC^+^HDL treatments and Ctr-HDL (269 upregulated, 88 downregulated). (B) KEGG pathway analysis of differentially expressed gene-sets. Color scale indicates false discovery rate (FDR); dot size represents the percentage of cells that express the gene for each pathway. (C) mRNA expression by qPCR displaying relative quantitative values (RQV) from BMDMs treated with Ctr-HDL (n=9) and CAC^+^HDL (n=14). Data are presented as mean + s.e.m. Mann-Whitney U-test was performed, \**p* < 0.05. (D-E) Western blotting of TMEM123 expression in (D) BMDMs or (E) human monocyte derived macrophages treated with healthy Ctr-HDL or CAC^+^ HDL (1 mg/mL) for 24h, n=3. Densitometric analysis of normalized TMEM123 protein expression was used to calculate significant fold differences in TMEM123 protein expression. Statistical analysis assessed by one-way Students t-test. Data are mean + s.e.m., \**p* < 0.05.

**Table 1.**
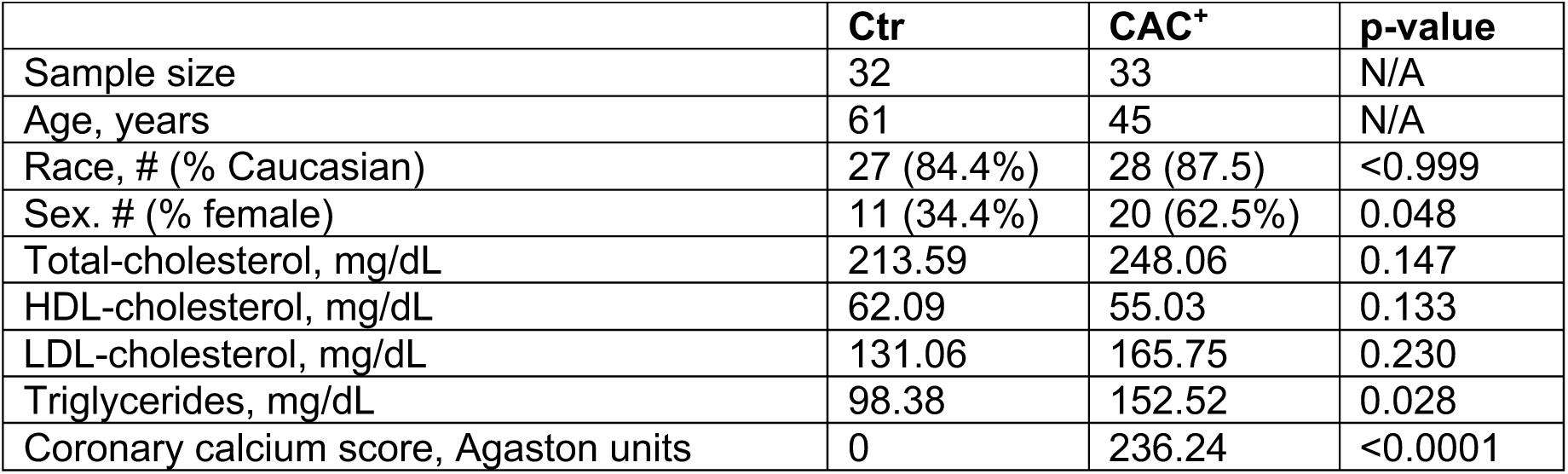
Clinical characteristics for healthy control (Ctr) and ASCVD (CAC+) subjects.

|  | **Ctr** | **CAC<sup>+</sup>** | **p-value** |
| --- | --- | --- | --- |
| Sample size | 32 | 33 | N/A |
| Age, years | 61 | 45 | N/A |
| Race, # (% Caucasian) | 27 (84.4%) | 28 (87.5) | <0.999 |
| Sex, # (% female) | 11 (34.4%) | 20 (62.5%) | 0.048 |
| Total-cholesterol, mg/dL | 213.59 | 248.06 | 0.147 |
| HDL-cholesterol, mg/dL | 62.09 | 55.03 | 0.133 |
| LDL-cholesterol, mg/dL | 131.06 | 165.75 | 0.230 |
| Triglycerides, mg/dL | 98.38 | 152.52 | 0.028 |
| Coronary calcium score, Agaston units | 0 | 236.24 | <0.0001 |

### Post-transcriptional modifications are increased on HDL-sRNAs in ASCVD

To determine if HDL-RNAs harbor modifications that are altered in ASCVD, HDL-RNA nucleosides were quantified by LC-MS/MS. HDL particles were purified from plasma by DGUC from both healthy control subjects (Ctr) and individuals with atherosclerotic lesion development (CAC^+^), stratified by coronary artery calcium scores (Ctr=0, ASCVD=CAC^+^ >21) (Table 1). The purity of isolated native HDL were confirmed by fast-protein liquid chromatography (FPLC)-based size-exclusion chromatography (SEC), and total protein and cholesterol levels were quantified across FPLC-SEC fractions to assess for contaminating LDL or excess free protein (Fig.S2). Total RNA were isolated from approximately 1 mg/mL (total protein) of HDL from the plasma of healthy control (n=27) and ASCVD (CAC^+^) (n=25) subjects, then hydrolyzed to single nucleosides using Nucleoside Digestion Mix prior to LC-MS/MS analysis. The levels of each modified and canonical nucleoside were calculated based on nucleoside standards (Fig.S3, Tables S5,S6). Remarkably, all 4 main and >20 modified nucleosides were consistently detected across HDL samples (Table S7, Figs.2A-J). The levels (nM) of each main nucleoside and inosine were significantly higher in CAC^+^ subjects compared to healthy (Ctr) subjects (Fig.2A). The most abundant modifications detected on HDL-sRNAs for both healthy Ctr and CAC^+^ subjects were pseudouridine (Ψ), inosine (I), and 1-methylguanosine (m^1^G) (Table S7). After normalizing modified nucleosides relative to their respective canonical counterparts, we observed a significant decrease in the relative levels of Ψ, 5-methyluridine (m^5^U), and 7-methylguanosine (m^7^G) in CAC^+^ compared to Ctr subjects (Table S7, Figs.2B-D). Conversely, CAC^+^ subjects showed a significant increase in several modifications, including Inosine, 3-methylcytidine (m^3^C), 5-methylcytidine (m^5^C), 6-methyladenosine (m^6^A), 2,2-dimethylguanosine (m^2^_2_G) and 2,7-dimethylguanosine (m^2^_7_G) (Figs.2E-J). Collectively, LC-MS/MS studies here demonstrated that HDL transports modified ribonucleosides, and that the levels of these modifications are altered in atherosclerosis.

**Figure 2.**
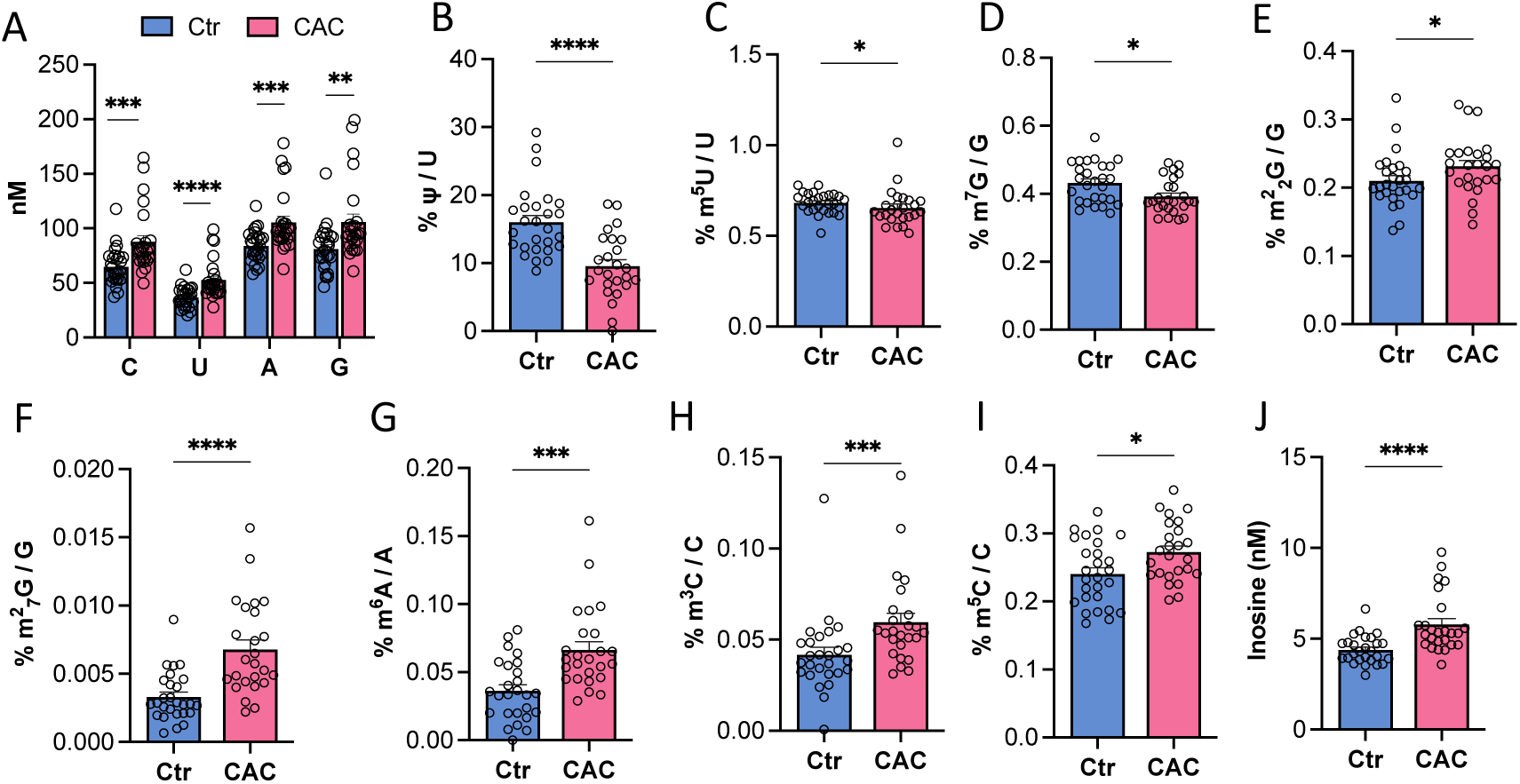
Mass spectrometry reveals altered RNA modifications on atherosclerotic HDL. (A) LC-MS/MS quantification of Ctr-HDL (n=27) and CAC^+^HDL (n=25) total unmodified nucleosides (cytidine [C], uridine [U], guanosine [G], and adenosine [A]) represented in nanomolar (nM). (B-J) Percent ratios of modified nucleosides to their respective unmodified counterparts (A) in Ctr-HDL and CAC^+^HDL samples. Modifications include pseudouridine (Ψ), 5-methyluridine (m^5^U), 7-methylguanosine (m^7^G), 2,2-dimethylguanosine (m^2^_2_G), 2,7-dimethylguanosine (m^2^_7_G), 6-methyladenosine (m^6^A), 3-methylcytidine (m^3^C), 5-methylcytidine (m^5^C), Inosine. (A-J) Mann-Whitney U-test performed. Data are presented as mean + s.e.m., \**p* < 0.05.

### Demethylation sequencing revealed 3’ tRNA-derived fragments on HDL

A major limitation to investigating modified sRNAs is that some modifications impede polymerase activity during first strand synthesis reactions for PCR or sRNA sequencing. To overcome this barrier, *E.coli* AlkB enzymes were used to enzymatically remove inhibitory methyl groups (i.e., m^1^A, m^3^C, m^1^G, and m^2^_2_G) from HDL-sRNAs prior to PCR or sRNA library generation^37–39^. Enzymatic efficiency of AlkB enzymes was tested prior to use by demethylating and assessing reverse transcriptase read-through efficiency of a fluorogenic (reporter) RNA aptamer containing either m^1^G or m^1^A methylations (Fig.S4A,B)^39^. Once validated, these enzymes were used to prepare HDL-sRNAs for sequencing from healthy Ctr (n=24) and CAC^+^ (n=24) subjects (Table 1). Comparative analysis of the AlkB-treated and untreated sRNA reads were carried out using our in-house data analysis pipeline (TIGER), which is specifically designed for lipoprotein sRNA analyses^5^. Using this approach, we detected 1.12x10^7^ and 1.15x10^7^ (mean) read depth for Ctr and CAC^+^ libraries, respectively. The most abundant class of host sRNAs observed on human HDL prior to RNA demethylation were rDRs, tDRs, yDRs and miRNAs (Fig.3A). AlkB treatments prior to sequencing significantly increased the detected levels of tDRs in HDL-sRNA datasets for both healthy and CAC^+^ subjects at both the parent transcript and read levels, suggesting that tDRs represent the most abundant class of host sRNAs on circulating HDL particles (Figs.3A-C, Table S8). At the read level, some rDRs and miRNAs were found to be increased after AlkB treatment; however, many were observed to be decreased (Figs.3D,E, S5A-D).

**Figure 3.**
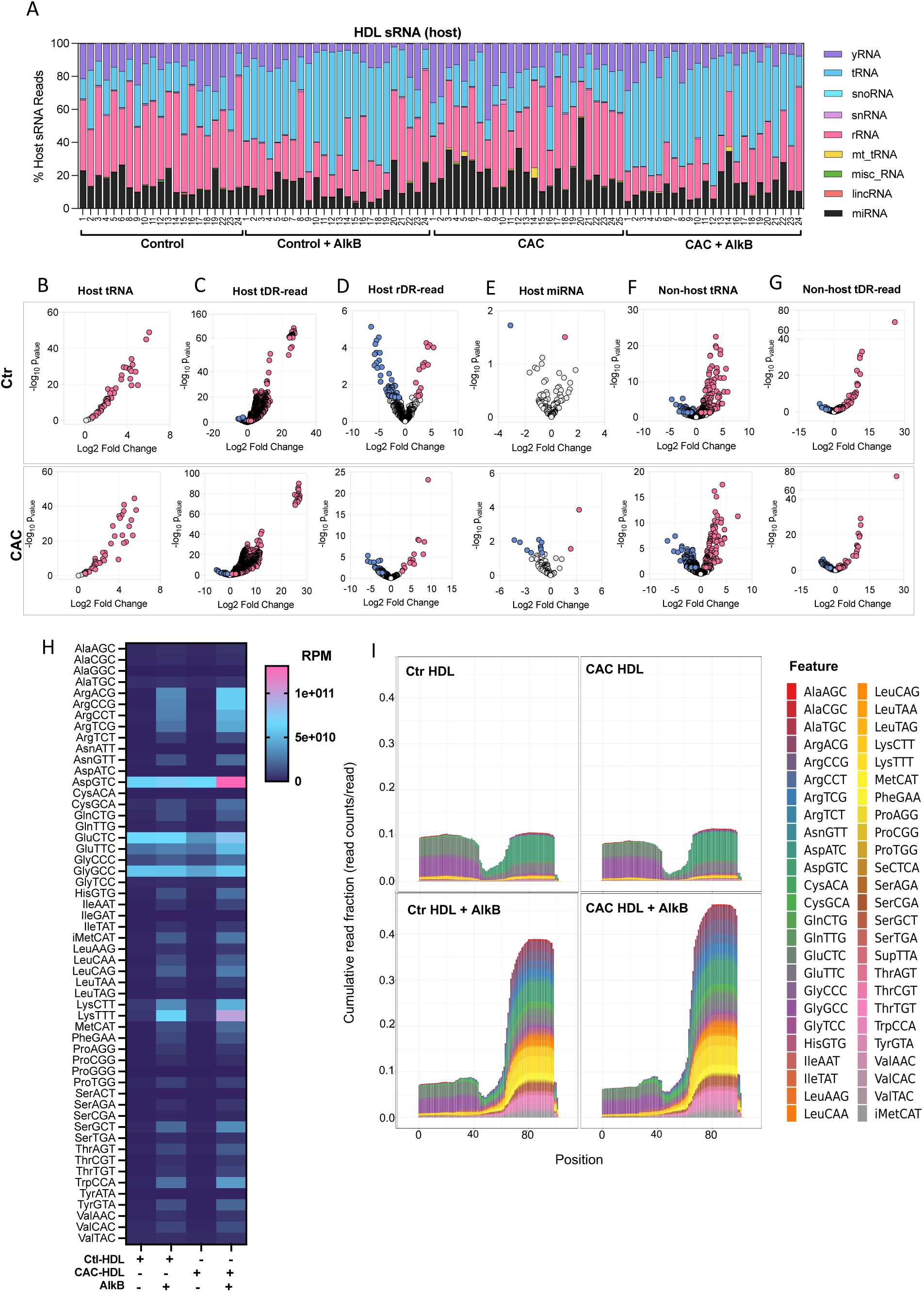
Demethylation increases capture of tRNA-derived sRNAs by sequencing. AlkB-mediated RNA (de)Methylase sequencing. (A) Distribution of host sRNAs abundance as percentage of total host reads for Ctr-HDL and CAC^+^HDL samples with and without AlkB demethylation treatment. (B-G) Differential expression (DEseq2) analysis of AlkB vs. non-AlkB treated samples. Adjusted p>0.05 differential (>1.5-absolute fold change) abundances for AlkB-treated host (B-C) tDRs (D) rDR and (C) miRNA at the parent and individual fragment levels, and non-host (F-G) tDRs at the parent and individual fragment level. (H) Heat-map representing the normalized reads per million (RPM) of total host tRNA reads aligning to multiple tRNA isoacceptors. (I) Positional coverage maps of healthy Ctr-HDL and CAC^+^HDL host tDRs for parent tRNA isoacceptors. Reported as mean cumulative read fraction (read counts/total counts).

We have previously reported that HDL transport a wide variety of non-host microbial sRNAs originating from bacteria, fungi, viruses, protozoa, and non-human eukaryotes from the microbiome, diet, and environment^5^. Strikingly, many microbial sRNAs were significantly increased after AlkB-treatments (at the read level), with 997 and 789 non-host sRNAs significantly upregulated in Ctr-HDL and CAC^+^HDL samples, respectively (Fig.S5E). Specifically, non-host tDRs, predominantly bacterial, were increased in HDL-sRNA datasets at both the parent transcript and read levels following AlkB treatment (Figs.3F-G). Among host (human) tRNAs, AlkB treatments specifically improved the detection of tDRs derived from arginine (ArgCCG, ArgACG, ArgTCG, ArgCCT), leucine (LeuTAA), methionine (MetCAT), asparagine (AsnATT), and alanine (AlaGGC) (Fig.3H). Most of these AlkB-sensitive tDRs corresponded to 3′ terminal fragments and halves of the parent tRNAs (Figs.3I). Interestingly, many AlkB-sensitive tDRs aligned with positions on parent eukaryotic tRNA transcripts known to carry modifications, as cataloged in the MODOMICS RNA modification database^41^. These results suggest that both host and non-host HDL-tDRs likely harbor AlkB-sensitive modifications and would not likely be captured by conventional sRNA sequencing approaches.

To assess the impact of ASCVD on HDL-sRNAs, including modified tDRs, we conducted differential expression analyses of AlkB-treated HDL samples from CAC^+^ and Ctr (CAC^-^) subjects, and reported changes at both the parent transcript and read levels (Figs.4A,B, Tables S9,S10). Many non-tDRs were also significantly altered on CAC^+^HDL compared to Ctr-HDL (Table S11). Nevertheless, among the many candidate tDRs identified by ARM-seq, tRNA-ArgACG-1 was observed to be enriched in CAC^+^HDL samples following demethylation and was therefore selected for down-stream functional investigation (Figs.4C,D, S5A-D, Table S12,). Strikingly, results from ARM-seq analyses of the 3’ terminal ends of ASCVD-associated tDRs showed unexpected variability in 3’ terminal non-templated “CCA” of ArgACG-1 tDRs on HDL (Fig.4E). Approximately 40% of ArgACG-1 tDRs on Ctr-HDL retained the full-length non-templated addition “CCA”, while the remaining tDRs exhibited truncated forms, e.g., CC, C, or other variations (Fig.4E). In CAC^+^HDL samples, only 49% of ArgACG-1 tDRs retained the full-length non-templated addition “CCA” (Fig.4E). Nevertheless, to confirm the presence of m^1^A-containing tDR-ArgACG-1 on HDL, RT-PCR were performed. In untreated samples, reverse transcription failed, indicating an obstructive modification; however, when samples underwent AlkB treatment to remove methyl groups, reverse transcription proceeded, enabling robust amplification of tDR-ArgACG-1 (Fig.4F). Quantification of AlkB-treated samples revealed approximately 8.6x10¹⁰ copies of tDR-ArgACG-1 per mg of HDL total protein in Ctr-HDL samples, compared to 2.5 x 10¹¹ copies per mg in CAC^+^HDL samples, a significant increase compared to Ctr-HDL (Fig.4F). These results confirm that HDL transport m^1^A-modified tDR-ArgACG-1, and that their levels are markedly elevated in ASCVD.

**Figure 4.**
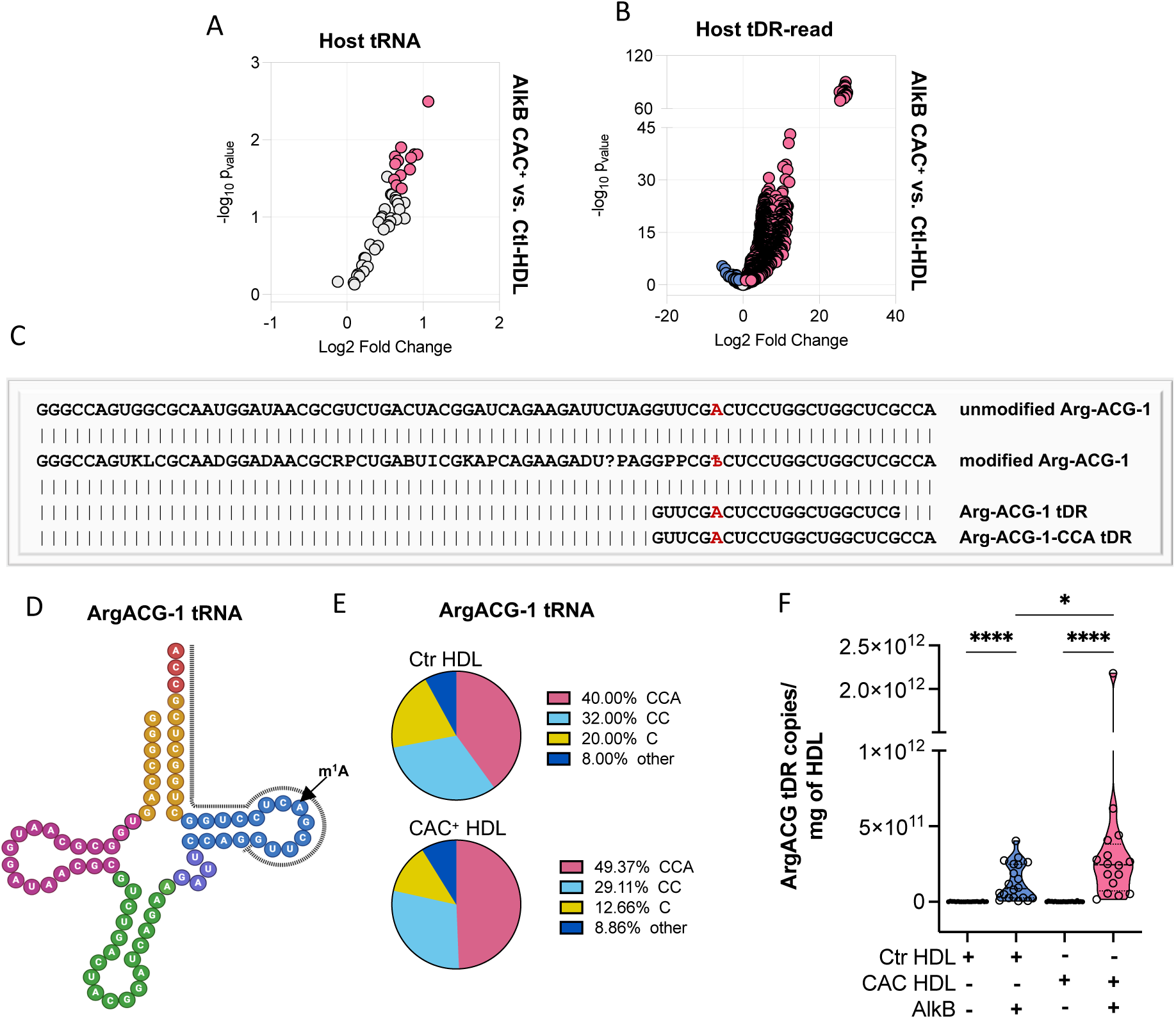
CAC^+^HDL contain increased methylated ArgACG-1 tDRs. (A-B) Differential expression analysis (DEseq2) of AlkB-treated healthy Ctr-HDL and CAC^+^HDL tDRs, n=24. Volcano plots demonstrating significant (adjusted p > 0.05) differential (>1.5-absolute fold change) abundances for reads mapping to host (A) parental tRNAs and (B) tDRs at the read level. (C) Alignment of HDL tDR-ArgACG-1 to both unmodified and modified forms of parental eukaryotic ArgACG-1 tRNA^41^. The modified tRNA sequence contains symbols representing RNA modifications according to the MODOMICS modification code (Ѣ: 1-methyladenosine, P: pseudouridine, ?: 5-methylcytidine, D: dihydrouridine, K: 1-methylguanosine, I: Inosine, B: 2’-O-methylcytidine, R: N2,N2-dimethylguanosine, L: N2-methylguanosine). (D) Schematic representation of unmodified ArgACG-1 tRNA highlighting tDR-ArgACG-1. (E) Chart depicting the percentage of healthy Ctr-HDL AlkB treated and CAC^+^HDL tDRs mapping to host ArgACG-1 tRNAs containing either a 3’ terminal CCA, CC, C or some other terminal end. (F) Quantification of Ctr-HDL (n=24) and CAC^+^HDL (n=24) tDR-ArgACG-1 copy numbers per 1 mg of HDL, as measured by qPCR before and after AlkB treatment. Statistical analysis assessed by Mann-Whitney U-test. Data are presented as mean + s.e.m., \**p* < 0.05.

### HDL delivery of m^1^A-tDR-ArgACG-1 drives pro-inflammatory macrophage activation

Based on our observations that the levels of HDL-tDR-ArgACG-1 harboring m^1^A were increased in the setting of ASCVD, we hypothesized that delivery of modified sRNAs by HDL could recapitulate the observed primary macrophage gene changes associated with disease CAC^+^HDL treatments. To evaluate the effects of HDL mediated delivery of m^1^A-containing tDRs on macrophage gene expression, primary human monocyte-derived macrophages (hMDM) were treated with rHDL (1 mg total protein) loaded with either unmodified tDR-ArgACG-1 or m^1^A-containing tDR-ArgACG-1. rHDL particles were composed of human apolipoprotein A-I (apoA-I) derived from endotoxin-free *Escherichia coli* (Clear Coli) and egg PC (1:100 molar ratio). For rigor, rHDL particles were demonstrated to be endotoxin free and were confirmed to be the appropriate size of HDL based on size-exclusion chromatography (Figs.S6A-B). To assess the biological activity of these candidate sRNAs and evaluate the impact of 3’ terminal non-terminal CCA additions, multiple forms of tDR-ArgACG-1 were synthesized with or without internal m^1^A modifications at position 6. First, candidates were synthesized without a 3’ terminal CCA and complexed to rHDL prior to macrophage delivery. Bulk mRNA sequencing revealed substantial changes in macrophage gene expression following treatments. We specifically tested gene expression changes between rHDL-m^1^A-tDR-ArgACG-21 compared to the unmodified rHDL-tDR-ArgACG-21 treatments. Remarkably, a single methylation (m^1^A) on tDR-ArgACG-21 was sufficient to cause robust changes to macrophage gene expression, with >950 genes significantly up-regulated and >550 down-regulated (Fig.5A, Table S13). rHDL delivery of m^1^A-tDR-ArgACG-1 lacking CCA were found to be highly inflammatory and significantly increased the expression of classic cytokines in hMDMs, including *CXCL5* (C-X-C motif chemokine 5, 34.53-fold), *SRPINB2* (plasminogen activator inhibitor-2, 32.98-fold), *IL6* (interleukin 6, 7.18-fold), *CSF2* (Colony Stimulating Factor 2, 5.85-fold), *CXCL8* (interleukin 8, 4.89-fold), *IL23A* (interleukin-23, 4.40-fold), *IL6* (Interleukin 6, 7.18-fold), and *IL1B* (4.38-fold) (Fig.5A, Table S13). Pathway analyses revealed that many of the most significantly altered pathways in response to rHDL-m1A-tDR-ArgACG-1 treatments were associated with immune cell activation, cytokine responses, cytoskeleton remodeling, and protein folding (Fig.5B, Table S14). These observations were confirmed in PCR assays, as rHDL-delivery were found to increase cytokine mRNA expression in treated macrophages compared to Ctr-HDL treatments (Fig.5C). In addition to classic cytokines, rHDL-m1A-tDR-ArgACG-1-21 deliveries were found to significantly increase *TMEM123* expression in treated macrophages similar to CAC^+^HDL treatments, as quantified by both RT-PCR and bulk sequencing (Figs.5A,C, Table S13). To assess the impact of rHDL-delivered m^1^A-tDR-ArgACG-1-21 on cytokine protein secretion, proximity extension assays (PEA, Olink) were completed on hMDM cell culture media. Results from PEAs demonstrated that rHDL-m^1^A-tDR-ArgACG-1-21 stimulated the secretion of 22 out of 45 measured cytokines from hMDMs, including TNF, IL6, IL1B, CSF1, CCL2, and CCL7, compared to delivery of unmodified tDR-ArgACG-1-21 (Fig.5D). To confirm PEA observations, IL-6 ELISAs were performed on treated-hMDM media samples, and results showed that rHDL-m^1^A-tDR-ArgACG-1 delivery significantly increased IL-6 protein secretion from primary macrophages compared to delivery of unmodified tDRs (Fig.5E).

**Figure 5.**
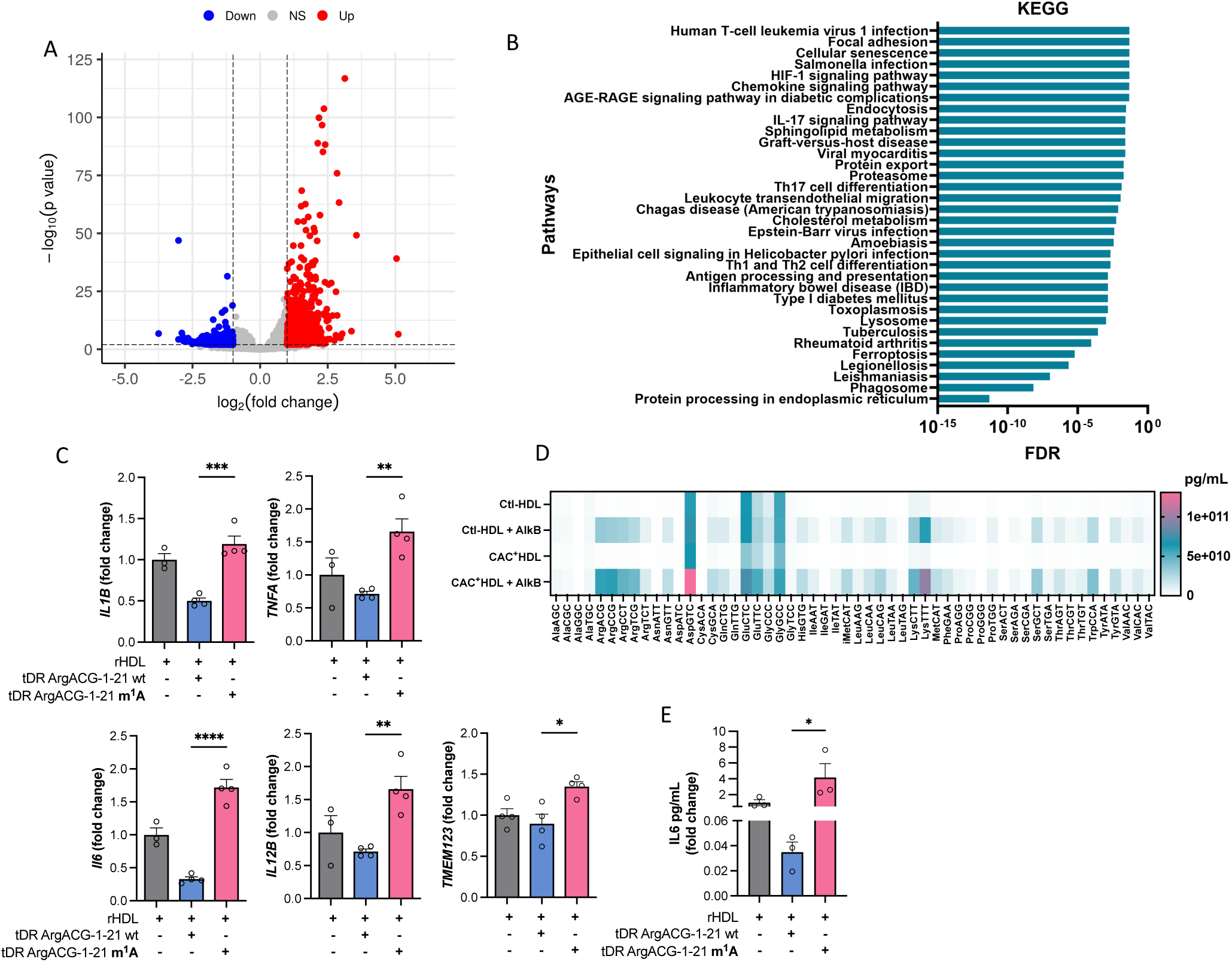
HDL delivery of m^1^A tDR-ArgACG-1-21 drives macrophage activation. (A-B) Total RNA sequencing of human monocyte derive macrophages (hMDM) treated with reconstituted HDL (rHDL, 1 mg/mL) complexed with tDR-ArgACG-1-21-m^1^A or its unmodified RNA counterpart (wt), n=4. (A) Volcano plots demonstrating significant (adjusted p > 0.05) differentially expressed genes (>1.5-absolute fold change) in hMDMs . (B) KEGG pathway analysis of differentially expressed gene-sets in rHDL m^1^A-tDR-ArgACG-1-21 induced macrophages compared to tDR-ArgACG-1-21 (wt) treatment, sorted by adjusted p-value with false discovery rate (FDR). (C) mRNA expression by qPCR displaying fold changes for *IL1B*, *TNFA*, *IL6, IL12B*, *TMEM123* in hMDM treated with rHDL, or rHDL complexed with tDR-ArgACG-1-21 (wt) or m^1^A-tDR-ArgACG-1-21. Statistical analysis assessed by two-way Students t-test. Data are presented as mean + s.e.m., \**p* < 0.05. (D) Heatmap analysis from Olink Target 48 Cytokine multiplex proximity extension assay representing human primary macrophage inflammatory cytokine secretion (pg/mL) in cell culture media after treatment with rHDL, or rHDL complexed with m^1^A-tDR-ArgACG-1-21 or tDR-ArgACG-1-21 (wt). (E) Secretion of IL-6 in hMDM supernatants after stimulation with rHDL m^1^A-tDR-ArgACG-1-21 or tDR-ArgACG-1-21 (wt) for 6 h by ELISA. Statistical analysis assessed by one-way Students t-test. Data are mean + s.e.m., \**p* < 0.05.

Non-templated CCAs are added to the 3’ terminal ends of parent tRNAs during maturation and these extra terminal nucleotides likely provide stability and resistance to RNase digestion^42^. Based on sequencing results, nearly half of tDR-ArgACG-1 molecules on circulating HDL do not contain the full 3’ terminal CCA addition and likely harbor some shortened form or lack non-templated additions. Nonetheless, many tDR-ArgACG-1 molecules are modified with terminal CCAs. To assess the impact of terminal CCAs and further investigate the role of HDL-m^1^A-tDR delivery to macrophages, candidate forms of modified and unmodified tDR-ArgACG-1-24 harboring 3’ non-templated CCA additions were synthesized. To determine if rHDL delivery of m^1^A-modified tDR-ArgACG-1-24, with a 3’ CCA terminus, also induced pro-inflammatory gene expression in recipient macrophages, RT-PCR were used to quantify target gene expression in treated hMDMs. Results revealed that rHDL-delivery of modified tDRs harboring CCAs significantly increased *IL1B* and *TNF* mRNA levels in treated hMDMs compared to unmodified HDL-tDR deliveries (Fig.6A). Of note, unmodified tDR-ArgACG-1-24 deliveries likely decreased inflammatory gene expression compared to rHDL alone; however, these beneficial features were lost with disease (CAC^+^) HDL treatments (Fig.6A). Strikingly, *TMEM123* mRNA levels were also significantly increased with rHDL delivery of m^1^A-modified tDR-ArgACG-1-24 compared to unmodified tDR (Fig.6A). These results are in agreement with macrophage gene responses observed with both CAC^+^HDL and rHDL-m^1^A-tDR-ArgACG-1-21 deliveries (Figs.1,5). To determine if m^1^A-tDR mediated effects on macrophage gene expression are dependent on the HDL receptor SR-BI (scavenger receptor class B type I), primary macrophages were pre-treated with block lipid transport-1 (BLT-1) prior to rHDL-tDR delivery. BLT-1 is an inhibitor of SR-BI mediated bidirectional cholesterol flux with HDL; however, BLT-1 also binds to HDL and increases HDL particle residency on SR-BI receptors^43,44^. BLT-1 likely increases HDL binding to SR-BI and promotes increased sRNA delivery to cells independent of cholesterol flux. Based on RT-PCR assays, BLT-1 treatments were found to enhance the m^1^A-modified tDR effects on *TNF* and *TMEM123* mRNA induction in macrophages by rHDL deliveries (Fig.6B). These results suggest that macrophage responses to rHDL-m^1^A-tDR-1 and CAC^+^HDL may potentially be mediated through SR-BI, independent of cholesterol flux. To further determine if candidate gene expression changes were associated with cholesterol efflux, macrophages were treated with methyl-beta cyclodextrin (MβCD), a strong cholesterol acceptor that mimics HDL and apolipoprotein A-I-mediated effects on inflammation and cholesterol efflux^45,46^. Results showed that cholesterol efflux alone is not sufficient to drive the expression of candidate genes in primary macrophages, including *TMEM123* (Fig.6C).

**Figure 6.**
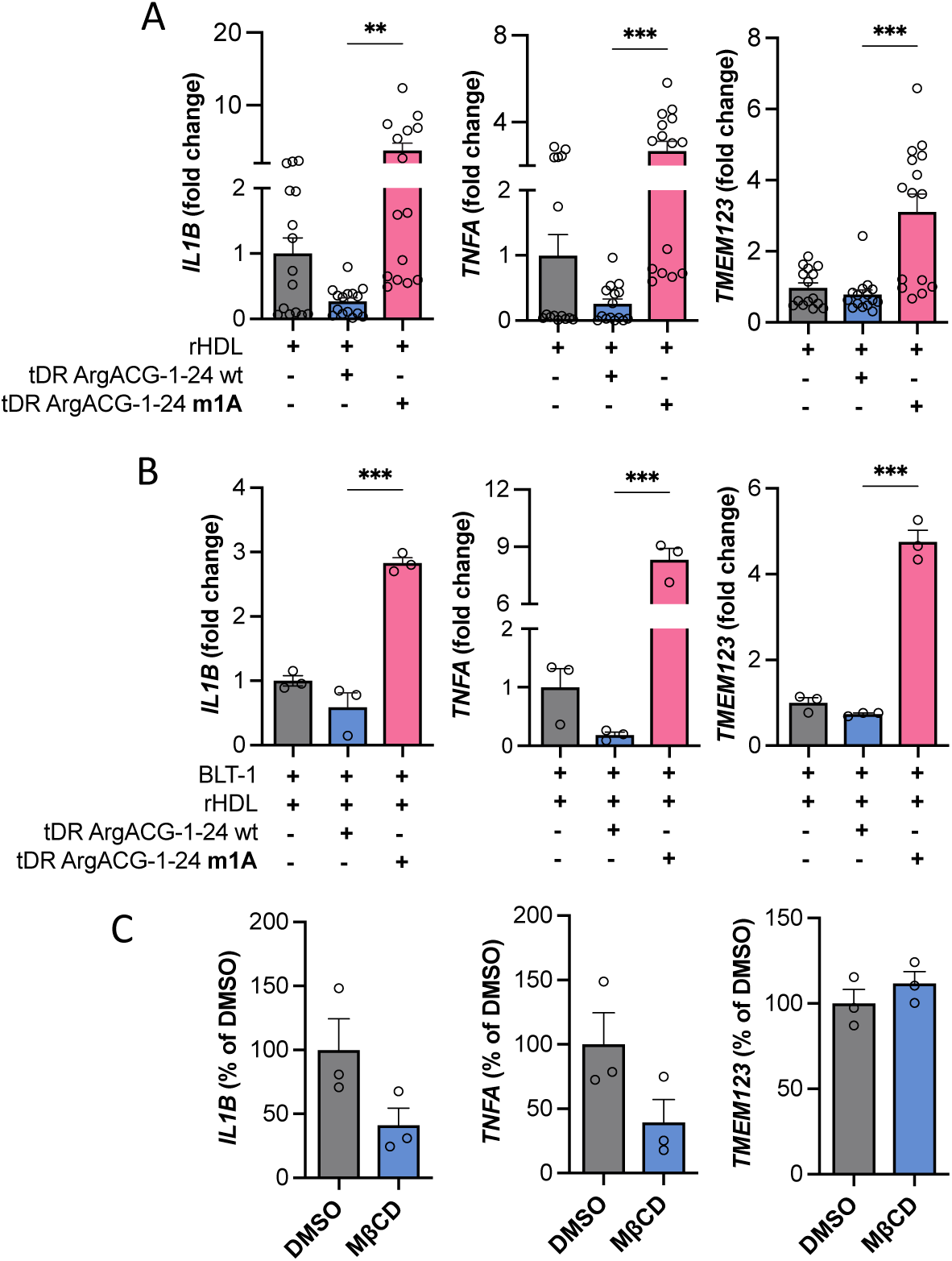
HDL delivery of m^1^A tDR-ArgACG-1-24 drives pro-inflammatory macrophage activation and *TMEM123* expression. (A-C) mRNA expression by qPCR displaying fold changes of *IL1B*, *TNFA* and *TMEM123* from human primary macrophages treated with (A) rHDL, rHDL complexed with m^1^A-tDR-ArgACG-1-24 or tDR-ArgACG-1-24 (wt), with and without (B) pre-treatment (1h) and co-treatment (6h) with block lipid transport-1 (BLT-1, 10 μM). (C) Methyl-β-cyclodextrin (MβCD, 10 μM) or dimethyl-sulfoxide (DMSO) treatment of primary human macrophages for 6h. Data are percent (%) of MβCD to DMSO. (A) Mann-Whitney U-test, or (B-C) two-way Students t-test. Data are presented as mean + s.e.m., \**p* < 0.05.

Collectively, results strongly suggest that m^1^A methylation on HDL-tDR-ArgACG-1 drives pro-inflammatory responses and likely influences macrophage phenotypes. The consistent upregulation of *TMEM123* following both m^1^A-tDR delivery and CAC^+^HDL treatment points to a macrophage activation signaling pathway potentially linked to SR-BI uptake of HDL-sRNAs and EGR1 transactivation of *TMEM123* expression. Together, these findings highlight how modifications on HDL cargo can reshape particle function and influence macrophage gene expression, potentially shedding light on how HDL may become dysregulated in the setting of ASCVD.

## DISCUSSION

ASCVD remains as the leading cause of morbidity and mortality worldwide, driven by complex interactions between lipid metabolism, immune responses, and inflammatory processes^47^. Therapeutically targeting atherosclerotic lesion inflammation has been historically challenging, and many patients on cholesterol-lowering drugs still carry residual inflammatory risk for cardiovascular events^48^. It is therefore critically important that new inflammatory drug targets and pathways are discovered. HDL have traditionally been viewed as a protective molecule, primarily due to its role in cholesterol efflux; however, many of HDL’s beneficial properties are lost during ASCVD^49,50^. While the underlying reasons for this switch are unknown, changes to its composition are likely a contributing factor. Our prior studies have revealed that HDL transports a diverse array of non-coding sRNAs, including tRNA-derived fragments (tDR), many of which are likely to carry modifications^3,5,51^. This study investigates whether these RNA modifications are altered on HDL-sRNAs in individuals with ASCVD and whether they contribute to pro-inflammatory gene expression changes and other phenotypic changes in macrophages. By using coronary artery calcium (CAC^+^) scores to stratify participants, we compared the modification profiles of HDL-sRNAs from individuals with advanced atherosclerosis (CAC^+^>21) to those from healthy controls (Ctr; CAC=0). Our findings suggest that disease-associated HDL particles acquire modified RNA cargo that confer changes to macrophage responses upon uptake. Our study demonstrates that HDL-associated sRNAs are decorated with key regulatory modifications, which introduce an additional layer of HDL-mediated gene regulation. Notably, we found that this novel regulatory mechanism is heightened in the context of ASCVD, potentially contributing to altered HDL function in this disease. This study provides a novel perspective on HDL function in atherosclerotic cardiovascular disease, highlighting HDL-sRNA modifications as potential targets and biomarkers.

HDL particles are dynamic transporters in extracellular biofluids, carrying a wide range of cargo that can undergo changes in composition, quality, and function under disease conditions. HDL play critical roles in systemic processes by delivering proteins, small molecules, bioactive lipids, and sRNAs, all of which likely contribute to both local and distal intercellular communication networks^52–55^. Our studies demonstrate that HDL from ASCVD subjects directly regulates an array of genes and pathways in primary macrophages, as compared to HDL from healthy subjects. For example, diseased HDL were found to increase the expression of multiple genes curated in cell adhesion, inflammation, and myeloid differentiation pathways. Although HDL have been reported to exhibit pro-inflammatory properties under certain conditions, our data show that physiological levels of CAC^+^HDL (1 mg/mL) did not significantly increase classic cytokine mRNA expression (*IL6*, *IL1B*, or *TNF*) at 6h post-treatment, but did induce the expression of other less-known inflammatory genes, including *TMEM123*. These direct gene changes did not require prior stimulus (endotoxin challenge) and were likely not associated with cholesterol flux. Studies of HDL dysfunction in atherosclerosis are often tested through HDL’s ability to dampen prior inflammatory stimuli, such as endotoxin (LPS). We did not observe a significant difference in HDL-mediated gene regulation between CAC^+^ and Ctr-HDL treatments following endotoxin challenges. Therefore, CAC^+^HDL induction of *TMEM123* in macrophages does not likely require prior stimulation or manipulation of the cells and is likely a direct effect of the transfer of modified RNA cargo to recipient macrophages. HDL’s ability to facilitate cholesterol efflux has previously been linked to increased pro-inflammatory gene expression in macrophages^56,57^. For example, passive cholesterol efflux to accepting HDL was reported to trigger Protein Kinase C signaling and NF-kB transcriptional activation in macrophages^57^. The ability of CAC^+^HDL to regulate *TMEM123* and other candidate genes is likely not linked to HDL-mediated cholesterol efflux in macrophages, as MβCD treatments failed to alter macrophage candidate gene expression. Overall, results from our analyses of macrophage gene responses to disease HDL showed that HDL likely do not turn on classic pro-inflammatory cytokine activation in primary macrophages during ASCVD. Nonetheless, results do support that disease HDL likely induces *TMEM123* expression and promotes cell adhesion, differentiation, and specific macrophage sub-phenotypes. We posit that the observed enrichment of modified sRNA cargo on HDL during ASCVD may play a pivotal role in driving these specific gene and phenotypic changes.

Of the >170 chemical modifications reported to date, eukaryotic tRNAs are predicted to harbor >100 of these distinct modifications, averaging approximately 13 modifications per tRNA molecule^41^. While m^6^A has become the most studied RNA modification, the role of m^1^A modifications, particularly on cell-free sRNAs circulating on HDL and other lipoproteins remains unexplored. Evidence suggests that m^1^A introduces a unique positive charge that influences RNA structure, protein interactions, and cellular pathways^58^. Gaining a strong understanding for how modifications regulate sRNA function in disease states is essential for evaluating their potential as disease biomarkers or therapeutic targets. To this end, it is necessary to accurately quantify and detect changes in nucleoside modification status in disease states. In this present study, we aimed to profile the modification status of HDL-sRNAs from subjects with advanced atherosclerosis using CAC scores >21 as indication of ASCVD disease status. To globally characterize the modification status on HDL-sRNAs, we first performed LC-MS/MS. Pseudouridine (Ψ) emerged as the most abundant modification on HDL-sRNAs for both healthy control (Ctr) and ASCVD (CAC^+^) subjects. This is not surprising given that Ψ represents the most abundant RNA modification^59^. Moreover, Ψ was observed to be significantly decreased on HDL-sRNA in ASCVD (CAC^+^) subjects compared to healthy individuals. Approximately 8.9% of all uridines on HDL-sRNA from diseased subjects were modified with Ψ, compared to approximately twice the amount (15%) of uridines on HDL-sRNAs from healthy control subjects. At positions along the sRNAs, Ψ modifications may antagonize sRNA recognition by TLR7/8 in recipient macrophages. Toll-like receptor 7 and 8 (TLR7, TLR8) are known single-stranded RNA (ssRNA) receptors for detecting viral and bacterial ssRNA^60–62^. Most, if not all, of the RNA cargo on HDL are likely in ssRNA form. As such, HDL bearing decreased Ψ levels would be predicted have reduced anti-inflammatory properties towards macrophage TLR7/8 activation. Similarly, levels of m^5^U, another methylation with reported anti-inflammatory effects, were significantly decreased on diseased CAC^+^HDL^60^. We observed multiple modifications to be significantly increased on HDL from ASCVD (CAC^+^) subjects including m^3^C, m^5^C, m^6^A, m^2^_2_G, and m^2^_7_G. While some HDL function(s) may be attributed to a few individual modifications, the collective impact of the overall modification profile likely governs HDL functionality. Interestingly, studies have shown that in monocyte-derived dendritic cells, m^6^A and m^5^C-modified RNAs reduced inflammatory gene expression, whereas the same modifications increased inflammatory gene expression in primary blood dendritic cells (DC1 and DC2)^60^. These studies suggest the effects of modified RNAs are likely cell specific, and that primary blood DCs may possess an additional RNA-sensing receptor that specifically recognizes RNA modified with m^5^C and m^6^A. At this time, the key receptor for m^1^A-tDRs, specifically responsible for the HDL m^1^A-tDR-ArgACG-1 effects in macrophages, is unknown.

Conventional sRNA-seq methods are limited in their ability to detect modified sRNAs due to the inhibitory effects of certain post-transcriptional modifications, which block first-strand synthesis by reverse transcriptase, an essential step in sRNA-seq library preparation. To overcome this limitation, we employed ARM-seq, which enzymatically removes specific methyl groups (i.e., m^1^A, m^3^C, m^1^G, and m^2^_2_G) from HDL-sRNAs prior to reverse transcription^37,63^. This ARM-seq approach was markedly more effective than conventional sRNA-seq, allowing us to capture many more (modified) sRNAs, particularly tDRs, that remained undetected with conventional sequencing methods. While rDRs appeared as the predominant sRNA class on HDL in conventional sequencing datasets, ARM-seq revealed tDRs to be the most abundant class on HDL, comprising approximately half of the total HDL-sRNAs (CAC^+^HDL: 51.74%; Ctr-HDL: 45.71%). These observed increases in tDRs mainly correspond to fragments that aligned to the 3’ terminal ends of parent tRNAs, indicating that a large portion of 3’ terminal tDRs likely contain AlkB-sensitive modifications, e.g., m^1^A at position 58. Indeed, others have reported that sRNAs, e.g., 3’ fragments from tRNAs are underrepresented by conventional sRNA-seq studies^37^. Although we did not observe an increase in overall m^1^A content within the CAC^+^HDL samples using LC-MS/MS, it is possible that m^1^A rearranged to m^6^A under alkaline conditions or heat prior to LC-MS/MS analysis^64^. Another reason could be that LC-MS/MS provides a broad, global profile of modifications across all HDL-sRNAs, whereas our ARM-seq approach allowed for focused analyses of specific classes of sRNAs, including tDR fragments.

We observed notable differences in host tRNA isodecoder representation between CAC^+^ and Ctr-HDL samples following AlkB treatment at the parent tRNA level. Specifically, 65% of the HDL-tDRs aligning to host parent tRNA isodecoders in both CAC^+^ and Ctr-HDL samples overlapped with tRNA transcript sequences listed in the MODOMICS database as containing AlkB-sensitive modifications. Strikingly, only after AlkB treatments did certain isodecoders, i.e., AlaGGC and ProGGG, appear, suggesting that these fragments likely harbor AlkB-sensitive methylations. Additionally, we detected an increase in HDL-tDR reads mapping to isodecoders harboring AlkB-sensitive methylations (e.g., m^1^G, m^2^_2_G, m^3^C, and m^1^A), not cataloged in the MODOMICS database, including ArgACG^41^. Although MODOMICS does not report methylations such as m^1^A within ArgACG tRNAs, other studies utilizing AlkB have demonstrated the presence of m^1^A at position 58 within ArgACG tRNAs^37,65,66^. In addition to tDRs, ARM-seq also increased the detection of rDRs, indicating that HDL-rDRs may also contain AlkB-sensitive modifications. Although miRNAs have been reported to contain modifications such as m^1^A, m^6^A, and m^5^C, they are less frequently modified compared to tDRs and rDRs, and our results indicate that miRNAs on HDL are not readily modified on HDL particles^67^. Notably, bacterial sRNAs were significantly enriched in HDL libraries following AlkB treatment, particularly bacterial tDRs, at both the transcript and read levels. Since bacterial tRNAs harbor methylations, e.g., m^1^A and m^1^G, it is likely that these non-host tDRs on HDL exhibit similar methylations.

Differential expression analysis of AlkB-treated host tDRs revealed an increase in many tDRs within CAC^+^HDL samples. Within the upregulated CAC^+^HDL-tDRs that were treated with AlkB, 33 sequences aligned to the MODOMICS reference database for eukaryotic tRNA sequences reported to harbor either m^1^A, m^1^G or m^2^_2_G methylations. (m^1^A,78.8%, m^1^G,18.2%, m^2^_2_G,3%)^41^. These results indicate a significant enrichment for methylated 3′ terminal tDRs, predominantly m^1^A-modified tDRs, in CAC^+^HDL compared to HDL from healthy controls. ARM-seq analysis revealed substantial diversity in the 3′ termini of ArgACG-1 tDRs circulating on HDL, with approximately half of the sequences containing the non-templated terminal CCA, while the rest displayed truncated forms such as CC, C, or other variations. This aligns with recent findings on EV-associated tRNAs, where tRNAs frequently lacked functional 3′ CCA ends^6968^. The observed variation between cellular tRNAs, which predominantly retain the 3′ CCA, and HDL-associated tDRs with variable 3′ termini, suggest a selective packaging mechanism within HDL, possibly tailored for specific roles in disease-associated signaling pathways. Notably, the tDR sequence, 5′-GUUCGACUCCUGGCUGGCUCGCCA-3′, derived from tRNA-ArgACG-1, emerged as particularly interesting among the tDRs uniquely enriched in CAC^+^HDL after demethylation. To confirm the presence and enrichment of m^1^A-containing tDR-ArgACG-1 on HDL, RT-PCR were performed. Due to m^1^A causing premature termination of cDNA synthesis and preventing PCR amplification, we designed reverse-complementary oligomers to probe for m^1^A by comparing amplification in untreated versus AlkB-treated samples. In untreated samples, PCR amplification of m^1^A-containing tDR-ArgACG-1 were undetectable, reflecting the inhibitory effect of m^1^A on reverse transcription. Following AlkB treatment and subsequent demethylation, reverse transcription was successful, enabling amplification of tDR-ArgACG-1. These RT-PCR results not only confirm the presence of m^1^A at position 6 in tDR-ArgACG-1, but also suggest that HDL-tDRs with m^1^A methylations are enriched in individuals with ASCVD. Our findings underscore the significant role of m^1^A-modified tDR-ArgACG-1 in enhancing pro-inflammatory responses in macrophages. We demonstrated that rHDL delivery of m^1^A-modified tDR-ArgACG-1 induced similar pro-inflammatory responses in macrophages, regardless of whether they contain a 3’ CCA terminus or not. Using synthetic rHDL particles loaded with tDR-ArgACG-1, we observed substantial upregulation of pro-inflammatory genes and cytokine secretion in treated macrophages. Bulk RNA sequencing showed marked changes in gene expression, with significant upregulation of *TMEM123* and pro-inflammatory markers such as *IL6, IL1B*, and *TNF*, while cytokine protein profiling of macrophage culture media confirmed elevated secretion of key inflammatory cytokines in response the HDL treatments, indicating that m^1^A modifications directly and strongly drive macrophage activation and cytokine secretion. Evidence may support that 3’ terminal CCA additions dampen some of the pro-inflammatory effects of m^1^A-methylations on tDRs, as PCR analysis of macrophage responses to rHDL delivery of CCA-containing m^1^A-tDRs were not as strong as responses to modified tDRs lacking the 3’ terminal CCA, particularly for classic cytokines *IL-6*, *IL-1B*, and *TNF*. The observed upregulation of *TMEM123* in response to HDL delivery of m^1^A-tDRs to macrophages suggests a pathway through which disease (ASCVD)-HDL could impact macrophage behavior. While TMEM123 itself is not an inflammatory marker, its involvement in immune cell adhesion, migration, and cytoskeleton organization implies a role in modulating immune cell activity^69,70^. Nevertheless, HDL-m^1^A-tDR induced macrophage gene responses may be linked to HDL binding to its receptor SR-BI, as BLT-1, an SR-BI cholesterol flux inhibitor and binding agent for HDL-SR-BI, increased the gene responses upon rHDL delivery of modified tDRs. Experiments using the cholesterol depletion agent, MβCD, failed to replicate the gene-expression changes seen after treatment of macrophages with CAC^+^HDL or HDL-m^1^A-tDR. These results suggest that the observed gene changes in macrophages are likely independent of HDL’s ability to accept cholesterol from macrophages.

One limitation of the study is related to the applied methods. The LC-MS/MS method employed in this study is sensitive, highly quantitative, and enabled detection of HDL-sRNA nucleoside modifications associated with disease states; however, it did not provide site-specific mapping of these modifications along sRNAs. Similarly, ARM-seq, while effective, only captures specific inhibitory RNA modifications, e.g., m^1^A, m^3^C, m^1^G, and m^2^_2_G. In addition to base methylations which interfere with sRNA-seq library construction, sRNAs can also harbor unique termini, e.g., 5’-phosphate and 2′-3′ cyclic phosphates, that prevent adapter ligation. While this study focused on AlkB-sensitive modifications within HDL-sRNA, future studies could use a combination of AlkB and T4 PNK treatments to test ligation efficiencies and assess ASCVD effects on sRNA termini. Another limitation of the study is that we currently do not know the RNA binding protein or receptor complex in macrophages that is responsible for mediating HDL-m^1^A-modified tDR signaling. Although the precise cell signaling mechanisms through which these modifications exert their effects remain unclear, it is possible that modifications alter HDL-sRNA interactions with cellular RNA-binding proteins or receptors involved in immune regulation. Further research is needed to elucidate these pathways and understand how modified HDL-sRNA cargo might contribute to altered gene expression and macrophage function in disease contexts.

Taken together, these results support a model whereby HDL delivery of m^1^A-tDRs in the setting of ASCVD influences macrophage gene expression, immune cell signaling, and sub-phenotypes within the atherosclerotic lesion. The distinct macrophage response to m^1^A-modified HDL-tDRs, compared to unmodified tDRs, points to m^1^A as a powerful signaling factor on circulating HDL that governs disease-associated changes to HDL’s biological functions, particularly related to atherosclerosis-associated inflammation.

## EXPERIMENTAL MODEL AND STUDY PARTICIPANT DETAILS

### Human plasma samples

Following informed written consent and institutional review board approval (IRB #101615 & 170046) plasma was collected from healthy adult donors (Ctr, n=21, females, n=11 males) with coronary artery calcium (CAC) = 0 and ASCVD subjects (CAC^+^, n=13 females, n=19 males) with CAC>21 and mean total cholesterol >200 mg/dL (Table 1). CAC was determined using non-contrast coronary CT examinations as previously described^55^. CAC was evaluated on 3.0 mm slices and reported as Agatston scores, based on a minimum lesion volume of 0.5 mm^3^. Plasma lipoprotein cholesterol and plasma triglyceride levels were quantified using Ace Axel clinical chemistry system (Alfa Wassermann).

### Animals

Adult (12-week-old) wild-type (RRID: IMSR_JAX:002052) mice on a C57BL/6 background purchased were housed with a maximum of 5 mice per cage in a 12 h light-dark cycle with unrestricted access to water and food (chow diet, NIH-31).

### Cell Culture

Peripheral blood mononuclear cells (PBMCs) were isolated from whole blood obtained from healthy donors using the standard Ficoll density gradient procedure. Primary human CD14^+^ monocytes were isolated from PBMCs following CD14^+^ selection using magnetic-activated cell sorting, according to manufacturer’s instructions, then seeded directly to tissue culture plates with hGM-CSF (20 ng/mL) and maintained in DMEM supplemented with sodium pyruvate (110 mg/L), L-glutamine, 10% heat-inactivated FBS and 1% penicillin-streptomycin for 6 days. On the sixth day, cells were washed with PBS and given fresh FBS-free medium containing hGM-CSF. Murine bone marrow cells were isolated from the femurs and tibiae of mice and cultured for seven days in treated cell culture plates. Cells were maintained in DMEM supplemented with 10% heat-inactivated fetal bovine serum, 1% penicillin/streptomycin, L-glutamine (300 mg/L), sodium pyruvate (110 mg/L), and mGM-CSF (40 ng/ml) for bone marrow-derived macrophage differentiation.

## METHOD DETAILS

### Density-Gradient Ultracentrifugation

Blood was collected in EDTA-containing collection tubes from fasted healthy and CAC subjects. Cell-free plasma was isolated by centrifugation (1,500 x *g;* 15 min; 4°C) and stored at -80°C until further use. Plasma was subjected to potassium bromide density gradient ultracentrifugation to separate very low-density lipoproteins (VLDL) (d = 0.94 -1.006 g/mL), LDL (d = 1.006 - 1.063 g/mL) and HDL (d = 1.063-1.21 g/mL). Briefly, plasma density was adjusted from d=1.006 g/mL to 1.025 g/mL with KBr, overlaid with saline buffer d=1.019 g/mL and centrifuged. After removing the VLDL fraction the density of the remaining sample was adjusted to d=1.080 g/mL with KBr, overlaid with saline buffer d=1.063 g/mL and centrifuged. The LDL fraction was then removed, and stored for later use, and the density of the remaining sample was adjusted to d=1.225 g/mL with KBr, overlaid with saline buffer d=1.210 g/mL, centrifuged, and the resultant HDL fraction was collected. The samples were centrifuged at 4°C for 24 h at 274,400 x *g* in a SW-40Ti rotor (Beckman Coulter Inc.). To remove KBr, the fractions of interest were extensively dialyzed in PBS utilizing 10k MWCO SnakeSkin dialysis tubing, then concentrated using Amicon Ultra 3K centrifugal filter device.

### Size-Exclusion Chromatography

Isolated human plasma HDL samples were injected into an AKTA pure fast-protein liquid chromatography (FPLC) system (Cytiva) with three tandem Superdex-200 Increase size-exclusion chromatography (SEC) columns and collected in 1.5 mL fractions containing buffer (10 mM Tris-HCl, 0.15 M NaCl, 0.2% NaN_3_). Total protein and total cholesterol were quantified in individual FPLC-SEC fractions, according to the manufacturer’s instructions.

### LC-MS/MS

Total RNA was isolated from equivalent amounts of HDL (1 mg), as determined by protein concentration, and isolated using RNeasy Mini Kit (Qiagen), according to the manufacturer’s instructions. RNA samples and ribonucleoside standards were enzymatically digested to single nucleosides under neutral conditions using the Nucleoside Digestion Mix prior to LC-MS/MS analysis. Synthetic ribonucleoside standards were purchased from Biosynth, Sigma-Aldrich and Cambridge Isotope Laboratories. LC-MS/MS analysis was performed on a Shimadzu Nexera system in-line with a QTRAP 6500 (Applied Biosystems). All separations were run on a Hypersil GOLD aQ C18 column (100-mm length × 2.1-mm inner diameter, pore size 175 Å, particle size 1.9 µm) at a flow rate of 0.4 mL/min using 0.1% (v/v) formic acid in water (solvent A) and acetonitrile with 0.1% (v/v) formic acid (solvent B). The gradient profile applied to each sample was as follows: 0–6 min, 0% B; 6–7.65 min, 1% B; 7.65-9.35 min, 6% B; 9.35-10 min, 6% B; 10-12 min, 50% B; 12-14 min, 75% B; 14-17 min, 75% B; 17-17.5 min, 0% B. The column was equilibrated prior to injections (20 µL) and maintained at 40°C. Data acquisition and sample processing were performed using AB SCIEX Analyst Software 1.6.2 (Applied Biosystems). Quantitative determination of RNA nucleosides was performed in ESI positive-ion mode using MRM. Table S6 summarizes the retention times (s), *m/z* values for precursor ion (Q1) and product ions (Q3), along with operational parameters such as collision energy (CE), declustering potential (DP), and dwell time for nucleosides. To quantify the amount of nucleoside in each sample, synthetic RNA nucleoside standards were used to prepare 16 calibration solutions with concentrations ranging from 30 pM to 1 μM. A linear regression curve was produced by extracting the peak areas of each nucleoside standard, and the lower and upper limits of quantification (LLOQ and ULOQ) were established from the resulting data (Table S5). The amount of HDL RNA nucleoside was calculated from the peak areas and linear fit equations of the calibration curve.

### ARM-seq

Purification of *E. coli* AlkB was adapted from previously described methods with some minor changes. AlkB wild-type protein (pET30a-AlkB ΔN11), D135S mutant protein (pET30a-AlkB-D135S) and D135T mutant protein (pET28a(+)-AlkB-D135T) were over-expressed in BL21(DE3) competent *E. coli* cells^37–39^. Transformed cells were grown in LB (Luria-Bertani) media supplemented with 50 µM kanamycin at 37°C until the OD_600_ reached 0.6. Protein expression was induced by addition of 1 mM isopropyl β-d-1-thiogalactopyranoside (IPTG) supplemented with freshly prepared iron sulfate (5 μM) for an additional 4 h at 30°C. Cells were harvested by centrifugation and resuspended in buffer A (10 mM Tris-HCL pH 7.4, 300 mM NaCl, 2 mM CaCl_2_, 10 mM MgCl_2_, 2 mM 2-mercaptoethanol, 5% glycerol) containing 1× protease inhibitor and lysed by sonication. Following centrifugation, protein lysate was loaded onto a Ni-NTA column pre-equilibrated with buffer A containing 20 mM imidazole. The protein was eluted off the column with buffer A containing either 70 mM or 250 mM of imidazole, then dialyzed against the storage buffer (20 mM Tris pH 8, 2 mM 2-mercaptoethanol). Purified AlkB protein was then aliquoted, supplemented with 30% glycerol, snap-frozen, and stored at −80°C until further use. Total RNA was isolated from equivalent amounts of HDL (2 mg), as determined by protein concentration, using miRNeasy Mini Kit, according to the manufacturer’s instructions. Demethylation reaction was performed as previously described with some edits. Briefly, RNA was demethylated in a 50 μL demethylation mixture containing 1 μM purified AlkB-wt, D135S and D135T enzymes, 25 mM MES buffer pH 6.0, 2 mM MgCl_2_, 270 mM KCl, 2 mM L-ascorbic acid, 300 μM α-ketoglutarate, 283 μM (NH_4_)_2_Fe(SO_4_)_2_, and 1 U/mL SUPERase•In RNase Inhibitor. The demethylation reaction was conducted at 25°C for 2h then stopped by addition of 5 mM EDTA and heated at 65°C for 10 min to denature the AlkB enzymes. RNA was purified from the reaction using RNeasy Mini Kit (Qiagen) and subsequently used for sRNA-seq library preparation. Untreated control reactions were performed similarly without the addition of AlkB enzymes. AlkB-treated and untreated HDL-sRNA libraries were generated using NEXTFlex Small RNA-Seq Kit v3 with a 1:8 adapter dilution, 22 PCR cycles, and the no size selection bead clean up. Pippin Prep 3% agarose gel cassettes with Marker P (Sage Sciences) were used to size select (156-216 bp) NEXTFlex sRNA-libraries. Libraries were quantified on a Qubit Fluorometer and qualitatively analyzed using Bioanalyzer High-sensitivity DNA Chips (Agilent). Paired-end sequencing (PE-150) of equimolar multiplexed libraries was performed on the NovaSeq6000 (S4) (Illumina) by the Vanderbilt Technologies for Advanced Genomics core. (GEO: GSE284076)

### Modified-RNA fluorescence assay

Broccoli RNA-based fluorescence assay was carried out, as previously reported with slight changes^39^. RNA demethylation was conducted in a 50 μl mixture containing 25 mM MES buffer (pH 6.0), 2 mM MgCl_2_, 50 μM (NH_4_)_2_Fe(SO_4_)_2_6H_2_O, 2 mM l-ascorbic acid, 283 μM α-ketoglutarate, 50 μg/ml BSA, 270 mM KCl, 0.8 U/μl SUPERaseIn RNase Inhibitor (Thermo), 1-2.5 μM of RNA and 10 μM of purified AlkB-WT, D135S, and D135T enzymes. The reactions were incubated at 25°C for 2h then quenched by addition of EDTA at 2.5 mM and heated to 65°C for 5 min to denature the protein. Reverse Transcriptase was conducted in a 20 μL total volume reaction containing 2.5 μL of the demethylation reaction, 0.5 μM primer, 5 μM dNTP, 5 μM DTT, 2U/μL SUPERaseIn RNase Inhibitor and 10 U/μL M-MLV RT in 1× RT buffer. A mixture containing the RNA template, primer, and dNTPs were first heated at 70°C for 5 min followed by quick chill on ice. After addition of RT buffer, DTT, and SUPERaseIn RNase Inhibitor, the reactions were incubated 37°C for 2 min. Lastly, M-MLV RT was added to the mixture, and the reaction was incubated at 42°C for 60 min, followed by heating at 70°C for 10 min to denature the M-MLV RT. PCR amplification was performed in a 20 μL total volume reaction using the Q5 high fidelity DNA polymerase (New England Biolabs Inc.) following the manufacturer’s protocol. *In vitro* transcription was conducted in a 20 μL reaction containing 7 μL of the PCR product, 6 mM rNTP, 25 mM MgCl_2_, 10 mM DTT, 50 μM DFHBI-IT and 25 U T7 polymerase in 1× RNAPol buffer. Fluorescence was recorded on a microplate fluorometer (BioTek Inc.) with an excitation wavelength of 472 nm and emission wavelength of 507 nm. The *in vitro* transcription reactions were conducted at 37°C for up to 3h and fluorescence was recorded at 8 intervals.

### Synthetic rHDL production

Recombinant human apo-A1 (pET20b apo-A1) was over-expressed in ClearColi BL21 (DE3) (Lucigen). Upon reaching OD_600_ 0.7-0.8, cells were induced with 0.5 mM IPTG at 37°C for 3-4 h. After harvesting by centrifugation, cells were resuspended in lysis buffer (20 mM Tris-HCL pH 8, 500 mM NaCl, 5 mM imidazole) containing 1× protease inhibitor (Thermo Scientific) and lysed by sonication. Following centrifugation, protein lysate was loaded onto a Ni-NTA column (Qiagen) and eluted with a buffer containing 20 mM Tris–HCl pH 8, 500 mM NaCl and a gradient of imidazole (100 mM – 1M). Elution’s containing purified apo-AI were pooled and dialyzed against storage buffer (20 mM Tris–HCl pH 8, 250 mM NaCl) utilizing 10k MWCO SnakeSkin dialysis tubing (Thermo Scientific). Purified protein was then aliquoted, supplemented with 30% glycerol, snap-frozen, and stored at −80°C. Prior to sHDL production, purified apo-AI (1 mg) was incubated with RNA oligonucleotides (5 μg) at 37°C for 1 h. rHDL particles were prepared using the previously described sodium cholate dialysis method with some slight changes. Briefly, Egg-PC (L-α-Phosphatidylcholine (Egg, Chicken); Avanti) dissolved in chloroform was dried under N_2_ gas and resuspended in an appropriate amount of PBS. The mixture was then incubated in the presence of 10% sodium deoxycholate (Sigma) at 37°C until the solution was clear. Apo-AI complexed with RNA oligonucleotides (See Table S15) was then added to Egg-PC to a final molar ratio of 100:1 (Egg-PC:apo-AI) and incubated at 37°C for 12h. The mixture was dialyzed extensively against PBS utilizing 10k MWCO SnakeSkin dialysis tubing, then sterile filtered (0.20 μm) prior to use.

### Chemical Modification of LDL

Acetylated LDL (AcLDL) was prepared by addition of 2 μL of acetic anhydride per 1 mg of LDL in 50% saturated sodium acetate over a 2 h period at 4°C. The AcLDL was then dialyzed for 24 h against PBS with 0.15M NaCl.

### Tissue Culture Treatments

For western blotting experiments, HDL and rHDL treatments (1 mg/mL) were carried out for 24 h in serum free DMEM. All other experiments involving native HDL or rHDL (1 mg/mL) were carried out for 6 h in serum free DMEM. For experiments involving the small molecule inhibitor Block Lipid Transport-1 (BLT-1, Sigma-Aldrich), the inhibitor was added to the media at a final concentration of 10 μM, 1 h before and during HDL treatments. For lipopolysaccharide (LPS) treatments (*Escherichia coli* O111:B4, Millipore Sigma), cells were exposed to LPS (10 ng/mL) 4 h prior and throughout the HDL experiments. Similarly, treatments with the TLR2 agonist, heat-killed Listeria monocytogenes (HKLM, InvivoGen) (10^7^-10^8^ cells/ml) and LPS (10 μg/ml) were administered 6 h prior to and during HDL treatments. For experiments involving the cholesterol-depleting agent, methyl-β-cyclodextrin (MβCD) was added to the media at a final concentration of 10 μM for 6 h.

### Endotoxin Assay

Endotoxin limulus amebocyte lysate (LAL) activity (EU/mL) of reconstituted HDL particles (1 mg) made with BL21 (DE3) competent *E. coli* or LPS free ClearColi BL21 (DE3) electrocompetent cells was measured using the Pierce Chromogenic Endotoxin Quantitation Kit (Thermo Scientific; #A39552S).

### Western Blotting

Whole cell lysates were prepared using RIPA with the addition of 1x Halt protease and phosphatase inhibitor (Thermo Scientific). Protein concentrations were determined by BCA protein assay according to manufacturer’s instructions (Pierce BCA). Cell lysates were resolved by SDS-PAGE and transferred onto PVDF membranes using iBlot Transfer Stack (Thermo Fisher) and probed with polyclonal rabbit TMEM123 antibody (Novus Biological, #NB100-56371, 1 μg/ml), human TMEM123 antibody (R&D Systems, #MAB3010, 1 μg/ml) or Rabbit Anti-GAPDH Monoclonal Antibody (Cell Signaling Technology, Cat# 2118; 0.5 μg/ml) (Figure S7). Band intensities were quantified using densitometry and normalized with corresponding GAPDH and compared to control (Ctl) samples. Proteins were visualized by ChemiDoc Imaging System (Bio-Rad) and quantified with Image Studio software.

### Transcriptomics

Total cellular RNA was isolated using Total RNA Purification kits (Norgen) and cDNA was synthesized using High-Capacity cDNA Reverse Transcription Kit (Thermo), according to manufacturer’s instructions. Total HDL-RNA was isolated using miRNeasy Mini kits (Qiagen), and cDNA was generated using the miRCURY LNA RT kit (Qiagen), as per the manufacturer’s instructions. Real-time PCR was carried out using Power SYBR 2× Master Mix (Thermo) on QuantStudio12 instruments (Life Technologies). *Rplp01 and PPARA* served as a housekeeper for human cells and an arbitrary Ct of 32 was used to generate relative quantitative results for HDL-sRNAs. The delta-Ct method was used to generate relative quantitative results, and data is reported as fold change.

### Total RNA sequencing

For hMDM treated with rHDL complexed with synthetic RNA, total RNA-seq libraries were generated using the Universal Plus Total RNA sequencing kit with NuQuant (Tecan) as per manufacturer’s instructions (GEO: GSE284268). For mouse BMDM, total RNA-seq libraries were generated using the NEBNext Ultra II Directional RNA Library prep kit for Illumina with multiplex oligos set 1 (NEB) (GEO: GSE284071). Briefly, approximately 100ng of total RNA from hMDMs or mouse BMDMs were subjected to chemical fragmentation followed by cDNA synthesis. Synthesized cDNA then underwent end repair, adapter ligation, and rRNA depletion. Libraries were then PCR amplified (13 cycles) and assessed for quality using the Agilent 2100 Bioanalyzer, and quantity using the Qubit fluorometer 3.0. Paired-end (150) sequencing of equimolar pooled libraries were performed on the NovaSeq6000 (Illumina) platform by the Vanderbilt Technologies for Advanced Genomics core.

### Informatics

Paired-end sequencing (PE-150) of equimolar multiplexed libraries was conducted on the NovaSeq6000 (S4) platform (Illumina) by the Vanderbilt Technologies for Advanced Genomics core, using the in-house data analysis pipeline, TIGER, as previously described^5^. Briefly, Cutadapt (v1.16) was used to trim 3′ adaptors from the raw reads and reads shorter than 16 nucleotides (nts) were excluded as ‘too short.’ The trimmed reads were aligned to the hg19 genome, supplemented with rRNA and tRNA reference sequences, using Bowtie1 (v1.1.2) with a maximum of one mismatch allowed. Reads shorter than 20 nts that did not perfectly align to the human genome or annotate as sRNA were removed from further analysis. The remaining unmapped reads were aligned against non-host structural RNA databases and curated microbial genome databases, allowing only perfect matches (no mismatches). Reads that failed to align to any reference were labeled as ‘unknown.’

### Statistics

Comparisons between two variables were performed using one-way Student’s t-test, two-way Student’s t-tests, or Mann-Whitney non-parametric test when appropriate. For comparisons involving more than two variables, a one-way analysis of variance (ANOVA) was used. Correlation coefficients (R) were calculated using Spearman’s rank method. A one-sided or two-sided p-value of <0.05 was considered statistically significant. Unless otherwise stated, the standard error of the mean (S.E.M.) was reported.

## ACKNOWLEDGMENTS

The authors would like to thank Anca Ifrim for assistance with sample procurement, Angela Jones of VANTAGE at Vanderbilt University Medical Center for expertise in high-throughput sequencing technologies, Jason Shrand for technical assistance and Wade Calcutt for his expertise in mass spectrometry.

## AUTHOR CONTRIBUTIONS

E.M.S.: conceptualization, data curation, formal analysis, investigation, methodology, visualization, writing – original draft. D.L.M.: project administration, methodology, supervision, writing – review & editing. P.J.K.: investigation, methodology, project administration. C.M.: investigation. M.A.R.: data curation, resources, software. Q.S.: data curation, resources, software. M.A.C.: investigation. A.C.D.: resources. J.J.C.: resources. L.J.M.: resources. M.F.L.: resources. K.C.V.: conceptualization, funding acquisition, methodology, project administration, supervision, validation, writing – original draft.

