## Supplemental Figures and Legends for "Post-transcriptional modifications on tRNA fragments confer functional changes to high-density lipoproteins in atherosclerosis"

### SUPPLEMENTAL INFORMATION:

**Figure S1. CAC+HDL directly stimulates macrophage gene expression, independent of cholesterol loading, prior inflammation and cholesterol efflux.** (A-E) mRNA expression by qPCR displaying relative quantitative values (RQV) of *Il6*, *Il1b*, and *Tnfa* from BMDMs treated with Ctr-HDL and $CAC^+$HDL pre-treated with (A) Acetylated LDL (AcLDL, 10 µg/ml), (B) heat-killed Listeria monocytogenes (HKLM, $10^7$-$10^8$ cells/ml), (C) AcLDL with HKLM, (D) lipopolysaccharide (LPS, 10 ng/mL), (E) LPS with AcLDL. (A) Mann-Whitney U-test performed. Data are presented as mean + s.e.m., $^*p < 0.05$. (B-E) Two-way Students t-test. Data are presented as mean + s.e.m., $^*p < 0.05$; ns: not significant.

**Figure S2. Characterization of isolated native HDL.** (A-B) The purity of isolated plasma HDL from healthy Ctl (n=32) and $CAC^+$ (n=33) subjects was assessed by size exclusion chromatography, and (A) total protein and (B) total cholesterol levels were quantified across fractions.

**Figure S3.** (A) Total ion chromatogram from LC-MS/MS analysis of commercial nucleoside standards as described in Materials and Methods. (B) A linear regression curve of Cytidine.

**Figure S4. Enzymatic efficiency of AlkB enzymes.** (A-B) Demethylation activity of AlkB-wt, AlkB-D135T and AlkB D135S mutants determined using an *in vitro* Broccoli RNA-based fluorescence assay[39]. The demethylation activity was assessed by comparing with the positive unmodified RNA control (pos cntl), and the negative $m^1$G-RNA control (neg cntl) which did not receive demethylase treatment. (A) Demethylation activity of D135S and D135T or the combination of D135S/D135T (1:1 Molar ratio) on $m^1$G containing RNA. (B) Demethylation activity of AlkB-wt on $m^1$A-containing RNA. Data are presented as fluorescent arbitrary units (AU), mean $+ \mathrm{s.e.m.}$

**Figure S5. ARM-seq alters healthy Ctr-HDL and $CAC^+$HDL small RNA profiles.** (A-E) Differential expression analysis of AlkB vs. non-AlkB treated Ctr-HDL and $CAC^+$HDL RNAs. (A) Volcano plots demonstrating significant (adjusted p > 0.05) differential (>1.5-absolute fold change) abundances for reads mapping to host (B) parental yRNAs (C) yDRs at the read level (D) parental rRNAs and (E) miRNAs at the read level and (F) non-host sRNAs - pink, increased; blue, decreased.

**Figure S6. Characterization of synthetic reconstituted rHDL**. (A) Endotoxin limulus amebocyte lysate (LAL) activity (EU/mL) of reconstituted HDL (rHDL) particles (1 mg) made with BL21 (DE3) competent *E. coli* or LPS free ClearColi BL21 (DE3) electrocompetent cells. (B) Purity of ClearColi BL21 (DE3) rHDL particles assessed by fast protein liquid chromatography and phospholipid concentration of each fraction.

**Figure S7. Atherosclerotic $CAC^+$HDL induces macrophage TMEM123.** (A-B) Western blot detection of TMEM123 and GAPDH protein expression in bone marrow derived macrophages treated with healthy Ctr-HDL or $CAC^+$ HDL (1 mg/mL) for 24h, n=3 (C-D) Western blot detection of TMEM123 and GAPDH protein expression in human monocyte derived macrophages treated with healthy Ctr-HDL or $CAC^+$ HDL (1 mg/mL) for 24h, n=3.

### Figure S1.

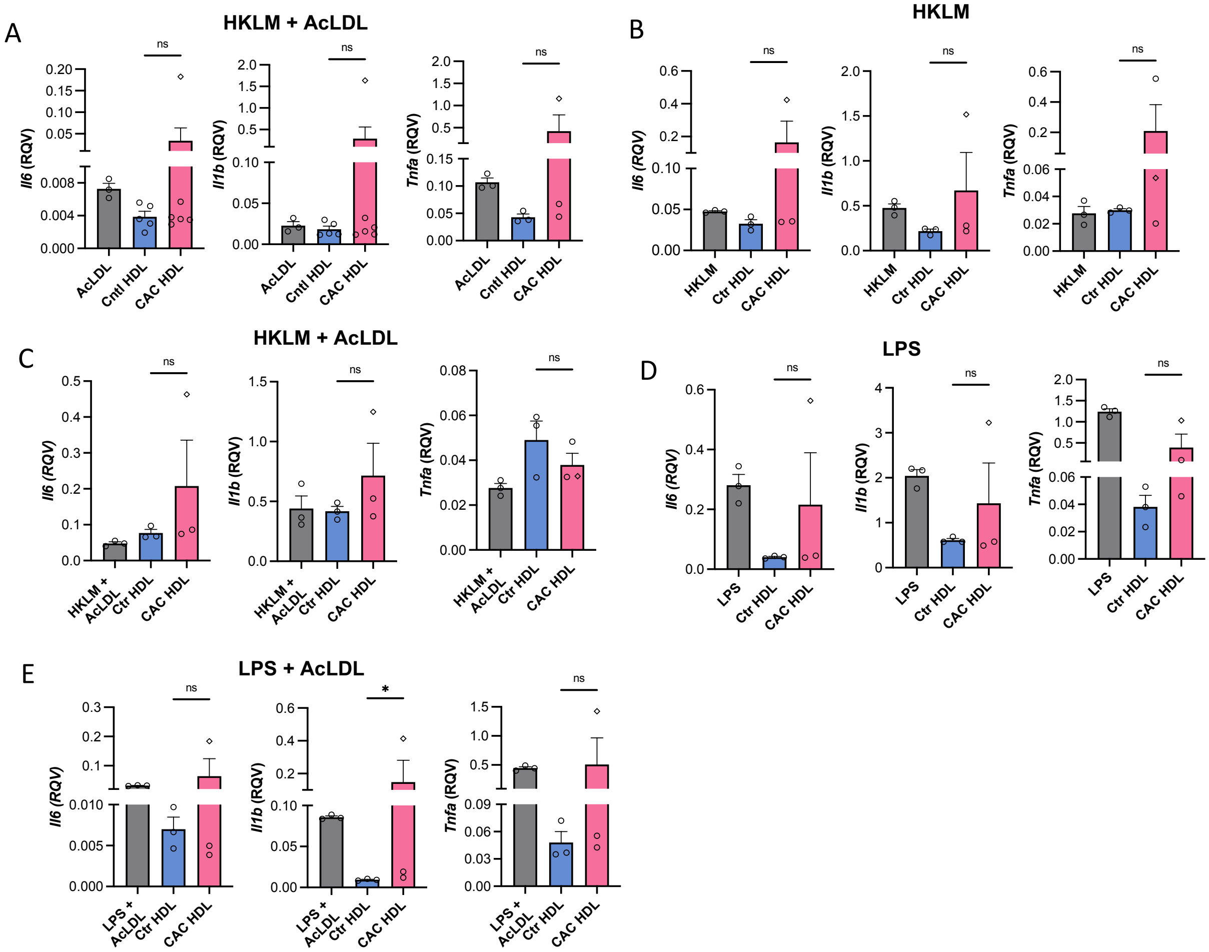


### Figure S2.

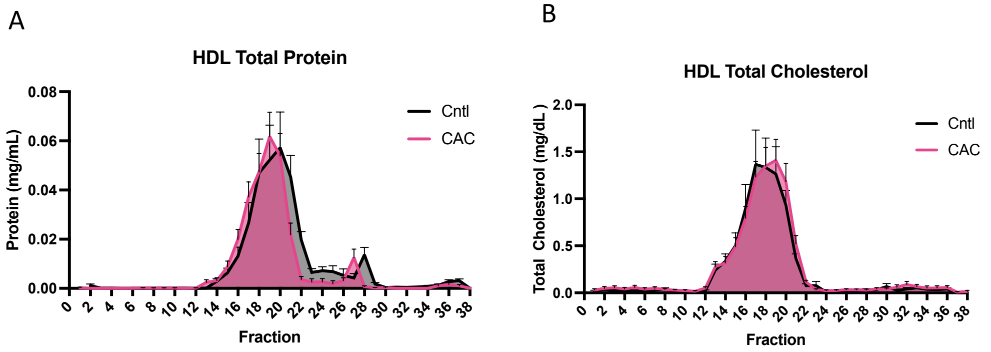

**Figure S3.**

A

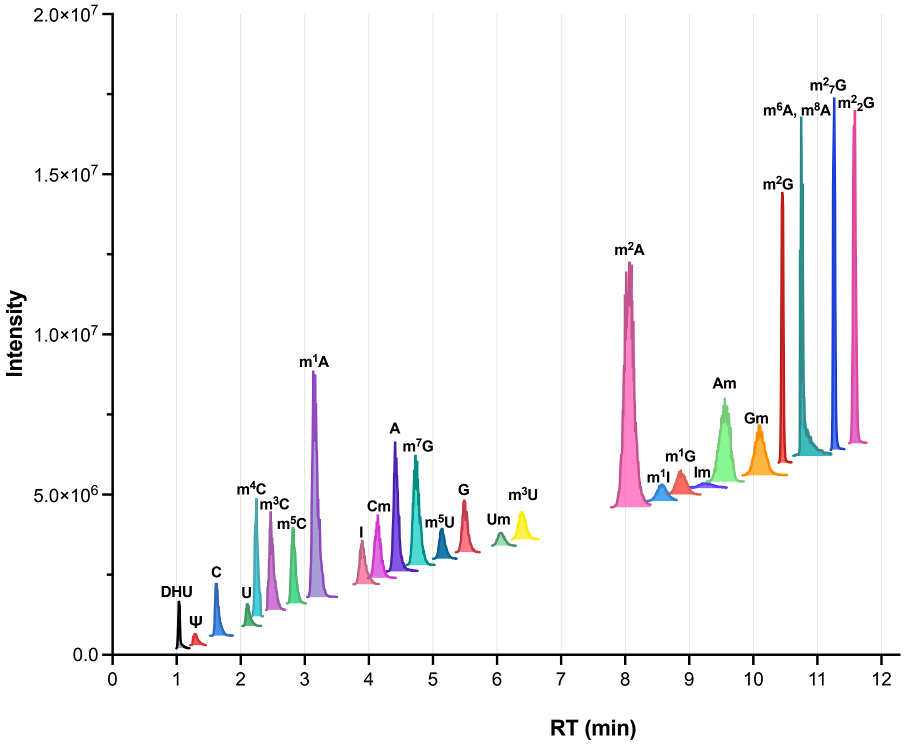

2.0×10^7
1.5×10^7
1.0×10^7
5.0×10^6
0.0
Intensity
RT (min)
DHU
Ψ
C
U
m4C
m3C
m5C
m1A
I
Cm
A
m7G
m5U
G
Um
m3U
m2A
m1I
m1G
Im
Am
Gm
m2G
m6A, m6A
m27G
m22G


B

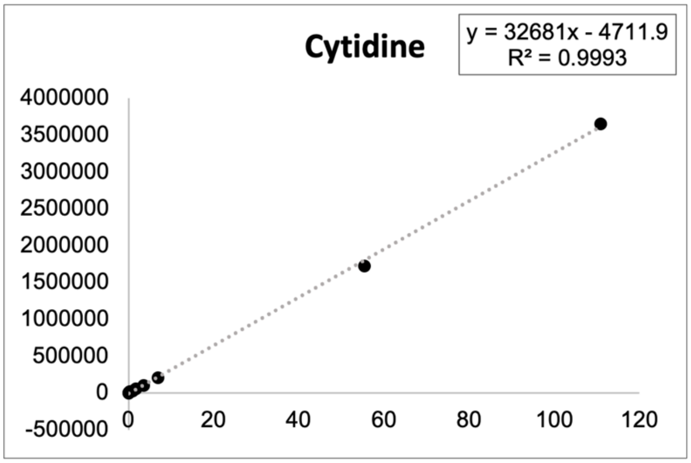

Cytidine
y = 32681x - 4711.9
R² = 0.9993
4000000
3500000
3000000
2500000
2000000
1500000
1000000
500000
0
-500000
0
20
40
60
80
100
120

### Figure S4.

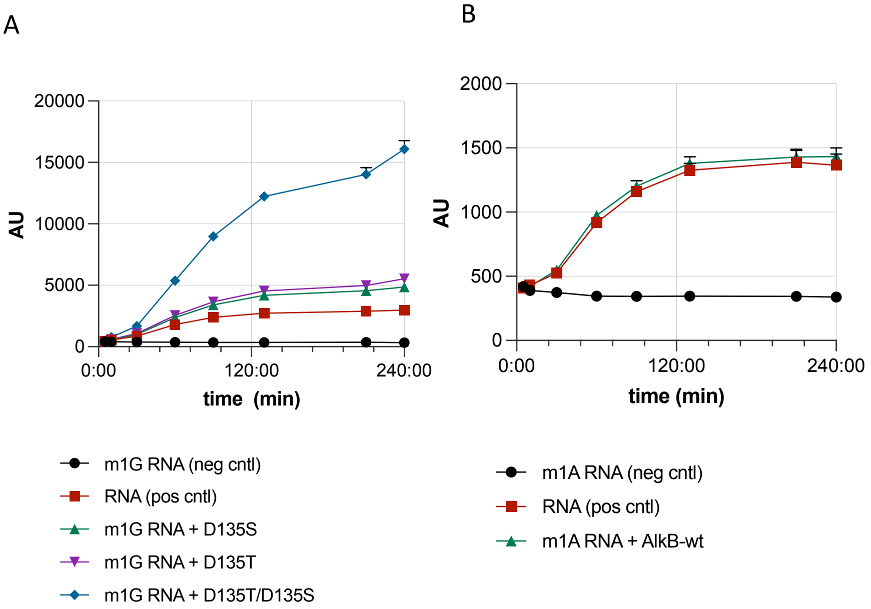


### Figure S5.

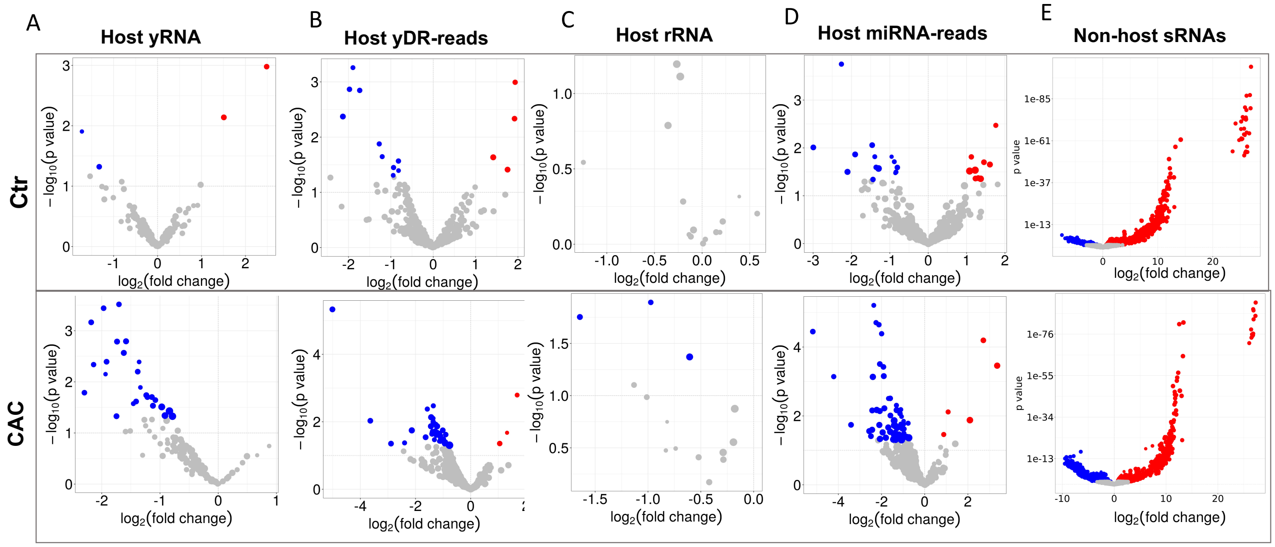

**Figure S6.**

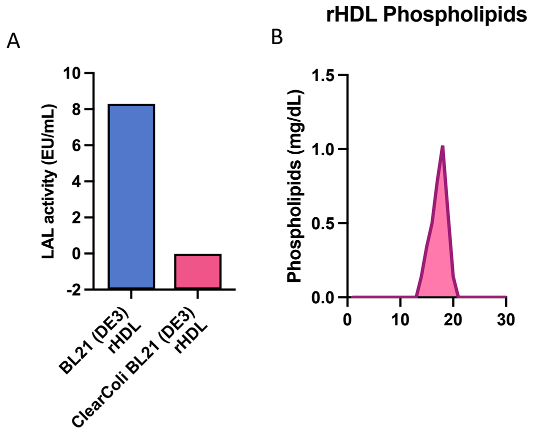


**Figure S7.**

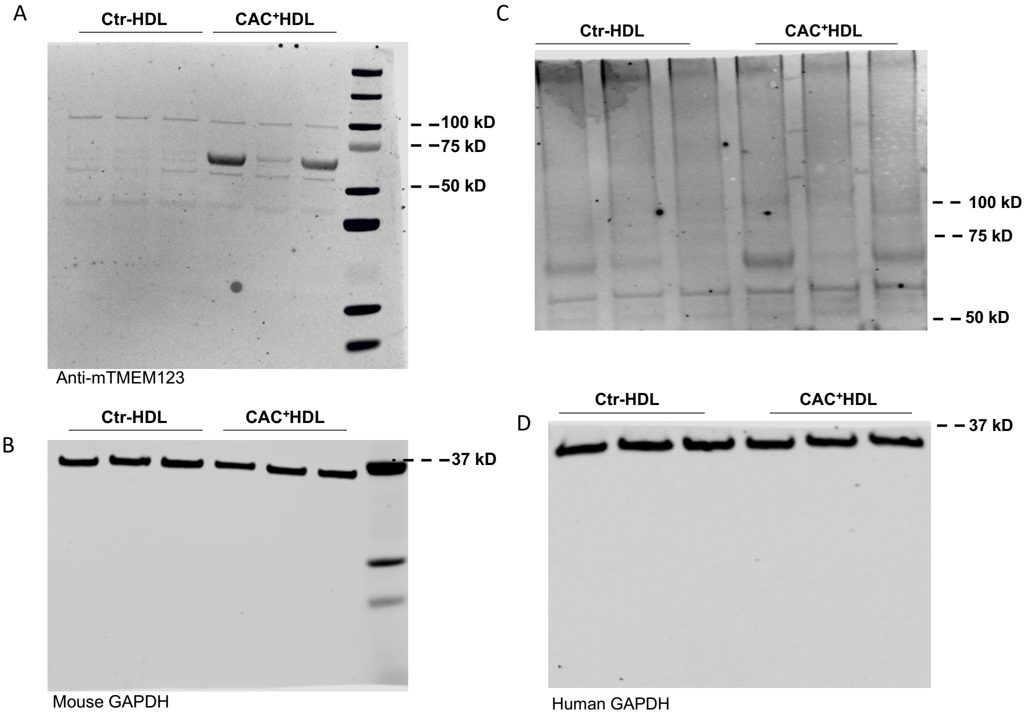
