## Supplemental Tables for "Post-transcriptional modifications on tRNA fragments confer functional changes to high-density lipoproteins in atherosclerosis"

**Table S1. Significant differentially altered protein coding genes in response to CAC-HDL compared to Ctr-HDL**

| Gene Symbol | Fold Change | padj |
|---|---|---|
| Aatk | -1.53 | 3.07E-08 |
| Abca13 | 1.67 | 7.49E-03 |
| Adgrg6 | 1.62 | 1.02E-19 |
| Adra2c | -1.59 | 2.57E-02 |
| Adrb1 | -1.81 | 1.52E-03 |
| Adrb2 | -1.58 | 1.53E-07 |
| Afmid | 1.56 | 2.52E-02 |
| Ager | 2.04 | 4.89E-03 |
| Ak8 | -1.63 | 2.12E-05 |
| Akap5 | 1.77 | 9.71E-05 |
| Alms1 | 1.62 | 1.15E-03 |
| Alox15 | -1.70 | 1.69E-05 |
| Angptl2 | -1.60 | 1.64E-22 |
| Ank2 | 1.69 | 1.82E-03 |
| Aoc1 | 2.34 | 6.55E-03 |
| Apol8 | 6.43 | 1.68E-07 |
| Ar | 1.92 | 2.24E-04 |
| Arfgef3 | 1.58 | 9.99E-10 |
| Arhgap28 | 1.55 | 6.22E-06 |
| Arid5a | 1.53 | 1.23E-09 |
| Arntl2 | 2.30 | 3.66E-02 |
| Asb4 | -1.55 | 3.88E-02 |
| Atp11b | 1.54 | 6.18E-22 |
| Atp23 | -1.75 | 2.68E-02 |
| Atp2b4 | 1.79 | 6.72E-09 |
| Bach2 | 2.14 | 2.29E-05 |
| Bbs10 | 1.51 | 3.38E-02 |
| Bcl2l14 | 1.90 | 8.89E-04 |
| Bclaf3 | 1.71 | 3.43E-05 |
| Bicd1 | 1.77 | 1.17E-04 |
| Brca2 | 1.72 | 1.99E-03 |
| Btla | 1.63 | 4.32E-04 |
| Cacna2d2 | 2.29 | 1.62E-03 |
| Cacna2d4 | 1.88 | 2.32E-02 |
| Cacnb2 | 2.26 | 3.25E-02 |
| Calcb | 1.83 | 2.51E-11 |
| Cald1 | -1.67 | 4.33E-03 |
| Caprin2 | 1.57 | 1.64E-02 |
| Car4 | -2.10 | 5.04E-23 |
| Cbr3 | -1.54 | 5.75E-12 |
| Ccin | 3.74 | 3.02E-04 |
| Ccr6 | 1.80 | 1.23E-02 |

| | | |
|---|---|---|
| Cd276 | -1.56 | 3.24E-13 |
| Cd46 | 2.08 | 6.55E-03 |
| Cd5 | 1.96 | 2.31E-02 |
| Cd5l | 1.73 | 9.43E-03 |
| Cd72 | -1.55 | 1.93E-05 |
| Cd80 | 1.55 | 3.74E-11 |
| Cdc14a | 1.59 | 8.43E-09 |
| Cdh23 | 1.83 | 1.18E-18 |
| Cdkl5 | 1.67 | 1.30E-04 |
| Cdon | 1.54 | 2.95E-04 |
| Cecr2 | 2.45 | 1.03E-02 |
| Cenpn | 1.87 | 8.99E-07 |
| Cep290 | 1.60 | 1.02E-03 |
| Ch25h | -1.95 | 4.38E-06 |
| Chrna2 | 1.92 | 9.95E-03 |
| Chst10 | 1.66 | 3.73E-02 |
| Chst13 | 2.08 | 4.31E-02 |
| Cldn15 | 2.19 | 2.55E-02 |
| Clec2i | 1.66 | 3.75E-08 |
| Cmc2 | 1.52 | 2.88E-06 |
| Cmklr1 | -1.53 | 3.70E-27 |
| Coch | 2.94 | 2.44E-06 |
| Col17a1 | 2.15 | 1.40E-02 |
| Col1a2 | -1.51 | 3.51E-03 |
| Crem | 1.75 | 1.55E-23 |
| Csf3 | -1.68 | 2.34E-04 |
| Csrp2 | 1.72 | 2.06E-03 |
| Cyb5rl | -1.60 | 2.60E-02 |
| Cytip | 1.53 | 6.31E-15 |
| Ddb2 | 1.61 | 4.28E-02 |
| Diablo | 1.54 | 1.58E-02 |
| Dixdc1 | 1.61 | 3.57E-02 |
| Dmrta2 | 1.78 | 2.54E-04 |
| Dnah2 | 1.52 | 1.98E-04 |
| Dock9 | 1.83 | 7.20E-09 |
| Dok2 | -1.51 | 3.46E-13 |
| Dpm1 | 1.50 | 7.46E-03 |
| Dpp4 | 1.51 | 1.91E-07 |
| Dpy19l3 | 1.54 | 1.42E-05 |
| Efna2 | 1.53 | 1.16E-03 |
| Egr1 | 3.38 | 7.72E-07 |
| Emilin1 | -1.50 | 3.58E-15 |
| Eml5 | 1.85 | 6.26E-05 |
| Emp2 | -1.60 | 1.68E-20 |
| En2 | 1.98 | 5.78E-09 |
| Engase | -1.52 | 1.13E-04 |

| | | |
|---|---|---|
| Ets1 | 1.57 | 4.21E-03 |
| Exph5 | 2.85 | 1.27E-03 |
| F5 | 2.19 | 3.75E-18 |
| Fam20c | -1.57 | 5.16E-22 |
| Fam71a | 3.59 | 1.90E-05 |
| Fam71f2 | 3.07 | 4.87E-05 |
| Fam83f | -1.94 | 5.23E-03 |
| Farp1 | 1.92 | 3.00E-03 |
| Fgf1 | 2.52 | 9.50E-05 |
| Flvcr2 | -1.56 | 9.48E-05 |
| Fndc9 | 2.37 | 7.62E-03 |
| Fos | 1.53 | 5.36E-04 |
| Fosb | 2.58 | 1.18E-04 |
| Frmd5 | 1.66 | 3.46E-03 |
| Frzb | 2.02 | 1.12E-05 |
| Fsd1l | 1.74 | 6.14E-05 |
| Fut7 | -1.75 | 1.53E-05 |
| Fyn | 1.55 | 8.05E-15 |
| Galnt9 | -1.70 | 2.40E-17 |
| Gas6 | -1.66 | 1.05E-05 |
| Gas7 | -1.53 | 2.09E-13 |
| Gem | 1.63 | 4.90E-05 |
| Ggt5 | -2.09 | 4.13E-10 |
| Ghrl | 1.97 | 6.50E-03 |
| Gls2 | 2.82 | 1.05E-18 |
| Gm10033 | 1.57 | 3.79E-02 |
| Gm36079 | 1.68 | 1.88E-02 |
| Gnal | 2.08 | 2.41E-02 |
| Gng4 | 1.77 | 5.88E-04 |
| Gp1ba | 1.59 | 2.15E-02 |
| Gpr65 | 1.63 | 2.81E-03 |
| Gprc5a | 1.64 | 1.04E-04 |
| Grcc10 | 1.64 | 4.89E-05 |
| Grip2 | 1.81 | 4.61E-02 |
| Gsta3 | -1.52 | 2.54E-04 |
| H1f3 | 1.58 | 2.97E-09 |
| H1f5 | 1.70 | 2.92E-06 |
| H2bc15 | 1.51 | 1.28E-03 |
| H2bc3 | 1.51 | 3.72E-03 |
| Hapln3 | 1.72 | 3.11E-05 |
| Has3 | 1.68 | 3.67E-02 |
| Hdgfl3 | 1.55 | 1.62E-02 |
| Hepacam2 | 1.57 | 1.68E-09 |
| Hhip | 1.81 | 2.32E-02 |
| Hid1 | 1.73 | 4.68E-06 |
| Hmox1 | -1.63 | 6.38E-17 |

| | | |
|---|---|---|
| Hsf2 | 1.61 | 7.26E-13 |
| Ica1l | 2.52 | 3.23E-02 |
| Icos | 1.59 | 2.79E-03 |
| Idi1 | 1.55 | 1.55E-06 |
| Igf1 | -1.51 | 1.44E-18 |
| Ikzf4 | 1.69 | 1.64E-22 |
| Il12rb1 | -1.57 | 4.11E-04 |
| Il12rb2 | 1.70 | 1.62E-05 |
| Il1f9 | -1.55 | 1.71E-12 |
| Il1r2 | 1.55 | 1.05E-02 |
| Il2ra | 1.56 | 1.58E-24 |
| Il33 | 2.16 | 1.33E-02 |
| Insyn2b | 2.01 | 2.75E-03 |
| Iqcb1 | 1.77 | 5.10E-03 |
| Itga11 | 2.01 | 4.40E-02 |
| Itga3 | 1.62 | 6.61E-03 |
| Kcnj2 | -1.71 | 2.91E-08 |
| Kcnk7 | 1.66 | 2.00E-02 |
| Kcp | 1.51 | 6.25E-16 |
| Kif18a | 1.84 | 2.47E-02 |
| Kif26b | -2.40 | 1.00E-06 |
| Klf2 | -1.94 | 7.62E-09 |
| Knl1 | 1.57 | 8.15E-04 |
| Krba1 | 1.78 | 3.51E-05 |
| Krit1 | 1.51 | 9.78E-13 |
| Lama5 | 1.58 | 3.63E-06 |
| Lepr | 2.14 | 8.33E-09 |
| Lipt2 | -1.59 | 2.33E-02 |
| Lpar5 | -1.60 | 1.33E-03 |
| Lrrc4 | 2.74 | 8.20E-04 |
| Lrrtm2 | 2.11 | 4.09E-02 |
| Lyzl4 | -2.21 | 2.65E-02 |
| Mag | -1.65 | 2.33E-05 |
| Mall | 2.07 | 2.30E-02 |
| Map1b | 1.75 | 7.93E-04 |
| Map3k15 | 1.58 | 6.07E-04 |
| Matn2 | -1.96 | 1.65E-02 |
| Mdk | 1.74 | 1.32E-02 |
| Megf11 | 2.10 | 2.25E-04 |
| Mfsd2b | 3.08 | 1.39E-03 |
| Mid1 | 2.05 | 4.40E-06 |
| Morn3 | 2.38 | 4.58E-03 |
| Ms4a4a | 1.51 | 8.78E-05 |
| Msi2 | 1.67 | 8.47E-06 |
| Msrb2 | -1.88 | 1.44E-06 |
| Mtcp1 | 1.67 | 2.95E-02 |

| | | |
|---|---|---|
| Mthfs | 1.54 | 5.58E-04 |
| Mthfsl | 1.66 | 1.65E-06 |
| mt-Nd2 | 1.57 | 4.75E-03 |
| Muc5b | 2.59 | 7.89E-03 |
| Mycbp | 1.52 | 6.49E-03 |
| Mylip | -1.52 | 1.30E-05 |
| Mylk | 1.51 | 3.68E-10 |
| Ncald | 2.26 | 1.31E-02 |
| Ndnf | 1.90 | 3.19E-06 |
| Neurl1a | 1.68 | 8.00E-08 |
| Neurl2 | -1.79 | 7.19E-03 |
| Nfat5 | 1.67 | 9.38E-26 |
| Nid2 | 1.80 | 1.45E-06 |
| Nod1 | -1.51 | 7.30E-08 |
| Nphp4 | 1.76 | 1.80E-02 |
| Nptx2 | 2.31 | 2.21E-02 |
| Nr4a3 | 1.79 | 1.87E-29 |
| Ntrk2 | 3.26 | 6.02E-03 |
| Nxpe5 | -1.90 | 8.97E-06 |
| Oas1c | -1.73 | 1.92E-02 |
| Ocln | 1.69 | 5.82E-03 |
| Ogfrl1 | 1.52 | 2.34E-10 |
| Oit3 | -1.83 | 7.61E-07 |
| Olfr920 | 2.87 | 1.04E-03 |
| Olr1 | 1.56 | 1.62E-09 |
| Orm2 | 1.91 | 1.14E-04 |
| Orm3 | 2.71 | 8.75E-04 |
| Osm | 1.75 | 6.71E-16 |
| Otud1 | -1.53 | 4.48E-02 |
| P2ry10 | 1.57 | 2.04E-06 |
| Papolg | 1.60 | 2.63E-09 |
| Paqr7 | -1.50 | 1.23E-08 |
| Pcgf5 | 1.55 | 3.77E-12 |
| Pde1b | 1.56 | 3.08E-15 |
| Pde3b | 1.52 | 6.71E-06 |
| Pde4b | 1.51 | 1.17E-14 |
| Pdk4 | 2.21 | 2.70E-03 |
| Pdzd2 | 1.96 | 4.48E-02 |
| Peg10 | 5.86 | 5.38E-09 |
| Per2 | 1.54 | 1.31E-05 |
| Perm1 | 1.53 | 1.34E-06 |
| Phka1 | 1.66 | 4.81E-08 |
| Pi16 | -2.10 | 6.85E-04 |
| Pkib | 1.62 | 1.98E-17 |
| Plat | -1.90 | 3.24E-02 |
| Plcb1 | 1.66 | 1.16E-02 |

| | | |
|---|---|---|
| Plekhb1 | 2.86 | 9.50E-04 |
| Plekhg5 | -1.52 | 1.12E-05 |
| Plod2 | 1.89 | 4.25E-03 |
| Pmel | 1.81 | 3.73E-02 |
| Pou4f1 | 2.02 | 2.61E-19 |
| Pparg | -1.54 | 1.98E-05 |
| Ppl | 3.30 | 3.20E-09 |
| Ppp2r3a | 1.69 | 5.15E-08 |
| Prdm9 | 1.72 | 7.27E-05 |
| Prkce | 1.62 | 6.84E-04 |
| Prkcg | 1.91 | 1.27E-03 |
| Prokr1 | -1.70 | 5.69E-03 |
| Prpf39 | 1.63 | 3.28E-10 |
| Prrg4 | 1.72 | 2.89E-07 |
| Prrt1 | -2.01 | 4.27E-02 |
| Prss46 | -1.60 | 1.90E-02 |
| Psip1 | 1.51 | 2.31E-05 |
| Ptbp2 | 1.59 | 6.91E-04 |
| Ptger3 | -1.64 | 1.49E-02 |
| Ptp4a1 | 1.80 | 6.33E-03 |
| Ptpn18 | -1.68 | 2.27E-03 |
| Ptpn4 | 1.57 | 4.39E-13 |
| Ptprf | 1.99 | 7.12E-10 |
| Ptprh | 1.73 | 2.85E-02 |
| Rabgap1l | 1.61 | 1.73E-20 |
| Radx | -1.50 | 4.64E-03 |
| Rasgrp1 | 1.52 | 1.34E-06 |
| Rbm4 | 2.55 | 2.08E-04 |
| Rbm4b | 1.67 | 3.34E-04 |
| Rbpms2 | 1.56 | 1.66E-03 |
| Reck | 1.64 | 4.35E-02 |
| Rel | 1.60 | 1.55E-23 |
| Rflna | 2.18 | 1.25E-02 |
| Rgs1 | 1.70 | 1.60E-16 |
| Ripk4 | 1.54 | 4.14E-02 |
| Rnase6 | -1.64 | 1.37E-03 |
| Sacs | 1.57 | 1.81E-10 |
| Samd11 | 1.68 | 7.56E-05 |
| Samsn1 | 1.54 | 2.08E-13 |
| Satb1 | 1.67 | 4.11E-08 |
| Scin | 1.65 | 6.62E-15 |
| Scube1 | 2.16 | 6.02E-08 |
| Sdcbp2 | 2.11 | 3.75E-09 |
| Septin2 | 1.81 | 9.11E-04 |
| Septin3 | -1.82 | 9.60E-06 |
| Shroom3 | 1.50 | 1.80E-02 |

| | | |
|---|---|---|
| Siglecg | 1.59 | 2.49E-05 |
| Sinhcaf | 1.51 | 1.28E-07 |
| Sirpb1b | -1.80 | 5.07E-06 |
| Sirpb1c | -1.62 | 2.89E-08 |
| Slc15a2 | 1.97 | 1.66E-07 |
| Slc18a2 | 1.84 | 2.12E-02 |
| Slc23a1 | 2.09 | 3.15E-02 |
| Slc24a1 | 3.00 | 1.86E-03 |
| Slc25a34 | 2.40 | 2.32E-02 |
| Slc40a1 | -1.85 | 4.00E-25 |
| Slc51a | 1.83 | 2.10E-03 |
| Slc6a1 | 4.80 | 8.03E-05 |
| Slc6a4 | -1.66 | 3.36E-16 |
| Slco5a1 | 1.70 | 2.72E-06 |
| Slf1 | 1.51 | 2.87E-02 |
| Slpi | -1.52 | 3.04E-03 |
| Smagp | -1.79 | 1.32E-02 |
| Sned1 | 1.51 | 8.62E-06 |
| Sox7 | -1.93 | 4.94E-02 |
| Spata1 | 4.39 | 9.48E-05 |
| Spib | 1.58 | 3.12E-05 |
| Spon1 | 2.31 | 3.63E-05 |
| Spry1 | -2.04 | 5.80E-04 |
| Sptb | 1.63 | 6.93E-03 |
| Srpk3 | 2.15 | 8.87E-03 |
| Stat4 | 1.80 | 4.21E-11 |
| Stk38l | 1.59 | 4.35E-13 |
| Stk39 | 1.53 | 1.04E-11 |
| Strip2 | 1.59 | 2.34E-13 |
| Stxbp6 | 1.68 | 2.09E-04 |
| Susd2 | -1.60 | 3.96E-06 |
| Suv39h2 | 1.54 | 5.74E-03 |
| Syn3 | 1.52 | 2.04E-03 |
| Syne2 | 1.57 | 1.89E-12 |
| Tbc1d10c | 1.54 | 3.74E-02 |
| Tbc1d16 | -1.57 | 5.84E-09 |
| Tead1 | 3.24 | 1.75E-15 |
| Tenm4 | 1.87 | 3.95E-04 |
| Tesc | 1.74 | 8.43E-03 |
| Tex14 | 2.34 | 4.93E-22 |
| Tex15 | 1.98 | 4.82E-02 |
| Tgfbr3 | 1.66 | 3.75E-04 |
| Thbs1 | 1.68 | 4.87E-04 |
| Tifab | -1.53 | 1.03E-16 |
| Timp3 | 1.69 | 8.74E-03 |
| Tk1 | 1.63 | 1.76E-18 |

| | | |
|---|---|---|
| Tlr5 | 2.16 | 2.37E-02 |
| Tmem123 | 1.50 | 3.00E-15 |
| Tmem178 | -2.07 | 4.67E-05 |
| Tmem38a | -1.53 | 3.66E-02 |
| Tmem41a | -1.78 | 8.97E-06 |
| Tmtc2 | 1.86 | 1.46E-19 |
| Tnfrsf11b | 1.73 | 4.89E-02 |
| Tnfrsf8 | -1.53 | 3.05E-02 |
| Tnfsf4 | 1.51 | 4.45E-11 |
| Tnni3 | 1.88 | 1.56E-04 |
| Tnnt3 | 1.65 | 1.17E-02 |
| Tox3 | 2.04 | 1.94E-03 |
| Tppp3 | -1.55 | 3.92E-03 |
| Trim47 | -1.55 | 6.28E-07 |
| Tspan13 | 1.50 | 1.13E-07 |
| Tspoap1 | 2.07 | 5.65E-44 |
| Unc5b | -1.96 | 2.50E-07 |
| Vat1l | 1.58 | 2.08E-02 |
| Vegfa | 1.56 | 3.87E-02 |
| Vsir | -1.53 | 1.62E-16 |
| Wnt11 | 1.79 | 3.24E-02 |
| Xylb | -1.75 | 5.31E-03 |
| Yes1 | 2.31 | 1.09E-05 |
| Zbtb10 | 1.51 | 1.25E-04 |
| Zc3h11a | 1.50 | 5.69E-03 |
| Zdhhc2 | 1.92 | 2.10E-02 |
| Zfc3h1 | 1.51 | 9.39E-27 |
| Zfp280d | 1.53 | 1.41E-04 |
| Zfp386 | 1.51 | 5.33E-05 |
| Zfyve28 | -1.90 | 3.15E-02 |
| Znrf3 | -1.57 | 2.83E-03 |

**Table S2. Gene set enrichment analyses of significantly altered genes upon CAC+HDL compared to Ctr-HDL**

| GeneSet | In Data | In Pathway | NES | FDR.q.val | Description |
|---|---|---|---|---|---|
| DACOSTA UV RESPONSE VIA ERCC3 COMMON DN | 174 | 435 | 2.70 | 0.00E+00 | Common down-regulated transcripts in fibroblasts expressing either XP/CS or TDD mutant forms of ERCC3 [GeneID=2071], after UVC irradiation. |
| SHEN SMARCA2 TARGETS UP | 174 | 415 | 2.00 | 4.98E-03 | Genes whose expression positively correlated with that of |

| | | | | | |
|---|---|---|---|---|---|
| | | | | | SMARCA2 [GeneID=6595] in prostate cancer samples. |
| ZHANG BREAST CANCER PROGENITORS UP | 171 | 407 | 1.85 | 2.12E-02 | Genes up-regulated in cancer stem cells isolated from mammary tumors compared to the non-tumorigenic cells. |
| ZHENG BOUND BY FOXP3 | 145 | 439 | 2.09 | 1.59E-03 | Genes whose promoters are bound by FOXP3 [GeneID=50943] based an a ChIP-chip analysis. |
| DUTERTRE ESTRADIOL RESPONSE 24HR DN | 129 | 415 | 1.74 | 3.78E-02 | Genes down-regulated in MCF7 cells (breast cancer) at 24 h of estradiol [PubChem=5757] treatment. |
| SENESE HDAC3 TARGETS UP | 121 | 403 | 1.79 | 3.05E-02 | Genes up-regulated in U2OS cells (osteosarcoma) upon knockdown of HDAC3 [GeneID=8841] by RNAi. |
| BLANCO MELO COVID19 SARS COV 2 INFECTION A594 ACE2 EXPRESSING CELLS RUXOLITINIB UP | 120 | 300 | 2.50 | 0.00E+00 | Genes up-regulated in SARS-CoV-2 infection with Ruxolitinib (ACE2 expressing A549 cells, MOI: 2, 24hpi) |
| BLANCO MELO COVID19 SARS COV 2 INFECTION A594 ACE2 EXPRESSING CELLS UP | 117 | 344 | 2.19 | 2.88E-04 | Genes up-regulated in SARS-CoV-2 infection (ACE2 expressing A549 cells, MOI: 2, 24hpi) |
| GABRIELY MIR21 TARGETS | 116 | 257 | 2.60 | 0.00E+00 | Genes significantly de-regulated ($p < 0.05$) by MIR21 [GeneID=406991] in A172 cells (glioma). |
| DAZARD RESPONSE TO UV NHEK DN | 113 | 275 | 2.56 | 0.00E+00 | Genes down-regulated in NHEK cells (normal keratinocytes) by UV-B irradiation. |
| FOSTER TOLERANT MACROPHAGE DN | 113 | 404 | 2.02 | 4.03E-03 | Class NT (non-tolerizeable) genes: induced during the first LPS stimulation and induced at equal or greater degree in tolerant macrophages. |
| BASAKI YBX1 TARGETS DN | 107 | 315 | 1.74 | 3.80E-02 | Genes down-regulated in SKOC-3 cells (ovarian cancer) after YB-1 (YBX1) [GeneID=4904] knockdown by RNAi. |
| HORIUCHI WTAP TARGETS DN | 97 | 293 | 1.74 | 3.82E-02 | Genes down-regulated in primary endothelial cells (HUVEC) after knockdown of WTAP [GeneID=9589] by RNAi. |
| SENGUPTA NASOPHARYNGEAL CARCINOMA WITH LMP1 UP | 96 | 291 | 1.93 | 1.02E-02 | Genes up-regulated in nasopharyngeal carcinoma (NPC) positive for LMP1 [GeneID=9260], a latent gene of Epstein-Barr virus (EBV). |
| CHEN HOXA5 TARGETS 9HR UP | 93 | 190 | 2.54 | 0.00E+00 | Genes up-regulated 9 h after induction of HoxA5 |

| | | | | | |
|---|---|---|---|---|---|
| | | | | | [GeneID=3205] expression in a breast cancer cell line. |
| GRESHOCK CANCER COPY NUMBER UP | 90 | 266 | 2.02 | 3.98E-03 | Genes from common genomic gains observed in a meta analyis of copy number alterations across a panel of different cancer cell lines and tumor samples. |
| FISCHER G2 M CELL CYCLE | 90 | 226 | 2.00 | 5.03E-03 | Cell cycle genes with peak expression in G2/M check point. |
| DEBIASI APOPTOSIS BY REOVIRUS INFECTION UP | 90 | 256 | 1.81 | 2.69E-02 | Genes up-regulated in HEK293 cells (embryonic kidney) at 6 h, 12 h or 24 h after infection with reovirus strain T3A (known as a strong inducer of apoptosis). |
| ZHENG FOXP3 TARGETS IN THYMUS UP | 88 | 192 | 2.32 | 0.00E+00 | Genes with promoters bound by FOXP3 [GeneID=50943] and which are up-regulated only in developing (located in the thymus) regulatory CD4+ [GeneID=920] T lymphocytes. |
| DEURIG T CELL PROLYMPHOCYTIC LEUKEMIA DN | 88 | 266 | 1.98 | 6.11E-03 | Genes down-regulated in T-PLL cells (T-cell prolymphocytic leukemia) bearing the inv(14)/t(14:14) chromosomal aberration. |
| FLORIO NEOCORTEX BASAL RADIAL GLIA DN | 85 | 163 | 2.03 | 4.12E-03 | Genes down-regulated in basal radial glia (bRG) relative to apical radial glia (aRG), and up-regulated in both aRG and bRG relative to neurons. |
| BILD HRAS ONCOGENIC SIGNATURE | 81 | 194 | 1.77 | 3.24E-02 | Genes selected in supervised analyses to discriminate cells expressing activated HRAS [GeneID=3265] oncogene from control cells expressing GFP. |
| ZHOU INFLAMMATORY RESPONSE LIVE UP | 79 | 305 | 1.75 | 3.55E-02 | Genes up-regulated in macrophage by live P.gingivalis. |
| REACTOME CHROMATIN MODIFYING ENZYMES | 75 | 242 | 2.17 | 4.13E-04 | Chromatin modifying enzymes |
| LEE RECENT THYMIC EMIGRANT | 72 | 199 | 1.87 | 1.90E-02 | Candidate genes specific for recent thymic emigrants (RTEs). |
| RUTELLA RESPONSE TO HGF VS CSF2RB AND IL4 DN | 71 | 221 | 2.18 | 2.72E-04 | Genes down-regulated in peripheral blood mononucleocytes by HGF [GeneID=3082] compared to those regulated by CSF2RB (GM-CSF) and IL4 [GeneID=1437;3565]. |
| STARK PREFRONTAL CORTEX 22Q11 DELETION UP | 68 | 159 | 1.74 | 3.77E-02 | Genes up-regulated in prefrontal cortex (PFC) of mice carrying a hemizygotic microdeletion in the 22q11.2 region. |

| | | | | | |
|---|---|---|---|---|---|
| DURCHDEWALD SKIN CARCINOGENESIS DN | 66 | 235 | 1.85 | 2.09E-02 | Genes down-regulated upon skin specific knockout of FOS [GeneID=2353] by cre-lox in the K5-SOS-F mice (express a constitutively active form of SOS1 [GeneID=6654] in the skin). |
| PYEON CANCER HEAD AND NECK VS CERVICAL UP | 64 | 161 | 2.12 | 1.26E-03 | Up-regulated genes in head and neck cancer compared to cervical carcinoma samples. |
| ZWANG CLASS 3 TRANSIENTLY INDUCED BY EGF | 64 | 188 | 2.07 | 2.58E-03 | Class III of genes transiently induced by EGF [GeneID =1950] in 184A1 cells (mammary epithelium). |
| BIDUS METASTASIS UP | 64 | 199 | 1.93 | 1.04E-02 | Genes up-regulated in endometroid endometrial tumors from patients with lymph node metastases compared to those without the metastases. |
| VILLANUEVA LIVER CANCER KRT19 UP | 63 | 154 | 1.92 | 1.18E-02 | Genes over-expressed in KRT19-positive [GeneID=3880] hepatocellular carcinoma (HCC). |
| JISON SICKLE CELL DISEASE DN | 62 | 164 | 2.12 | 1.24E-03 | Genes down-regulated in peripheral blood mononuclear cells (PBMC) from sickle cell disease patients compared to those from healthy subjects. |
| THUM SYSTOLIC HEART FAILURE DN | 62 | 162 | 1.78 | 3.15E-02 | Genes down-regulated in samples with systolic heart failure compared to normal hearts. |
| DAZARD UV RESPONSE CLUSTER G6 | 57 | 132 | 2.45 | 0.00E+00 | Cluster G6: genes increasingly down-regulated in NHEK cells (normal keratinocyte) after UV-B irradiation. |
| CHANDRAN METASTASIS UP | 55 | 161 | 2.02 | 4.04E-03 | Genes up-regulated in metastatic tumors from the whole panel of patients with prostate cancer. |
| PICCALUGA ANGIOIMMUNOBLASTIC LYMPHOMA DN | 54 | 126 | 2.32 | 0.00E+00 | Down-regulated genes in angioimmunoblastic lymphoma (AILT) compared to normal T lymphocytes. |
| WINNEPENNINCKX MELANOMA METASTASIS UP | 54 | 154 | 1.81 | 2.64E-02 | Genes from the 254-gene classifier which were up-regulated in melanoma patients with a reported distant metastasis within 4 years. |
| SHEPARD CRASH AND BURN MUTANT DN | 51 | 135 | 1.92 | 1.12E-02 | Human orthologs of genes down-regulated in the crb ('crash and burn') zebrafish mutant that represents a loss-of-function mutation in BMYB [GeneID=4605]. |
| GROSS HYPOXIA VIA ELK3 DN | 51 | 145 | 1.75 | 3.69E-02 | Genes down-regulated in SEND cells (skin endothelium) at hypoxia with ELK3 |

| | | | | | |
|---|---|---|---|---|---|
| | | | | | [GeneID=2004] knockdown by RNAi. |
| REACTOME MITOTIC SPINDLE CHECKPOINT | 49 | 109 | 1.85 | 2.22E-02 | Mitotic Spindle Checkpoint |
| ROSTY CERVICAL CANCER PROLIFERATION CLUSTER | 49 | 139 | 1.81 | 2.67E-02 | The 'Cervical Cancer Proliferation Cluster' (CCPC): genes whose expression in cervical carcinoma positively correlates with that of the HPV E6 and E7 oncogenes; they are also differentially expressed according to disease outcome. |
| PUJANA XPRSS INT NETWORK | 48 | 164 | 1.82 | 2.59E-02 | Genes constituting the XPRSS-Int network: intersection of genes whose expression correlates with BRCA1, BRCA2, ATM, and CHEK2 [GeneID=672;675;472;11200] in a compendium of normal tissues. |
| WP CIRCADIAN RHYTHM GENES | 47 | 143 | 1.84 | 2.26E-02 | Circadian rhythm genes |
| CASORELLI ACUTE PROMYELOCYTIC LEUKEMIA UP | 47 | 134 | 1.81 | 2.66E-02 | Genes up-regulated in APL (acute promyeolocytic leukemia) blasts expressing PML-RARA fusion [GeneID=5371;5914] compared to normal promyeloblasts. |
| BENPORATH NOS TARGETS | 47 | 135 | 1.74 | 3.77E-02 | Set 'NOS targets': genes upregulated and identified by ChIP on chip as targets of the transcription factors NANOG , OCT4, and Sox2 [GeneID=79923;5460;6657] (NOS) in human embryonic stem cells. |
| KEGG RIBOSOME | 46 | 82 | 1.90 | 1.36E-02 | Ribosome |
| REACTOME EUKARYOTIC TRANSLATION ELONGATION | 46 | 87 | 1.80 | 2.71E-02 | Eukaryotic Translation Elongation |
| NAGASHIMA NRG1 SIGNALING UP | 46 | 150 | 1.76 | 3.48E-02 | Genes up-regulated in MCF7 cells (breast cancer) after stimulation with NRG1 [GeneID=3084]. |
| IKEDA MIR30 TARGETS UP | 45 | 108 | 2.09 | 1.64E-03 | Genes up-regulated in hypertrophic hearts (due to expression of constitutively active form of PPP3CA [GeneID=5530]) and predicted to be targets of miR-30 microRNA. |
| IVANOVA HEMATOPOIESIS STEM CELL LONG TERM | 45 | 146 | 1.95 | 8.36E-03 | Genes in the expression cluster 'LT-HSC Shared': up-regulated in long term hematopoietic stem cells (LT-HSC) from adult bone marrow and fetal liver. |

| | | | | | |
|---|---|---|---|---|---|
| KEGG UBIQUITIN MEDIATED PROTEOLYSIS | 43 | 127 | 1.82 | 2.59E-02 | Ubiquitin mediated proteolysis |
| WP ENDODERM DIFFERENTIATION | 41 | 96 | 1.99 | 5.34E-03 | Endoderm differentiation |
| KONG E2F3 TARGETS | 40 | 97 | 2.04 | 3.80E-03 | Genes up-regulated in MEF cells (embryonic fibroblasts) at 16 hr after serum stimulation and knockdown of E2F3 [GeneID=1871] by RNAi. |
| VANTVEER BREAST CANCER METASTASIS DN | 40 | 99 | 1.82 | 2.57E-02 | Genes whose expression is significantly and negatively correlated with poor breast cancer clinical outcome (defined as developing distant metastases in less than 5 years). |
| IVANOVA HEMATOPOIESIS STEM CELL | 39 | 138 | 1.78 | 3.05E-02 | Genes in the expression cluster 'HSC Shared': up-regulated in hematopoietic stem cells (HSC) from adult bone marrow and fetal liver. |
| ODONNELL TFRC TARGETS DN | 39 | 103 | 1.76 | 3.41E-02 | Genes down-regulated in P493-6 cells (B lymphocyte, Burkitt's lymphoma model) upon knockdown of TFRC [GeneID=7037] by RNAi. |
| VERHAAK GLIOBLASTOMA PRONEURAL | 39 | 106 | 1.75 | 3.70E-02 | Genes correlated with proneural type of glioblastoma multiforme tumors. |
| DACOSTA UV RESPONSE VIA ERCC3 TTD DN | 38 | 77 | 2.12 | 1.23E-03 | Genes exclusively down-regulated in fibroblasts expressing the TTD mutant form of ERCC3 [GeneID=2071], after UVC irradiation. |
| PYEON HPV POSITIVE TUMORS UP | 38 | 80 | 1.97 | 7.06E-03 | Up-regulated genes in cervical carcinoma and head and neck tumors positive for human papilloma virus (HPV) compared to those negative for HPV. |
| REACTOME ESTROGEN DEPENDENT GENE EXPRESSION | 38 | 120 | 1.79 | 3.01E-02 | Estrogen-dependent gene expression |
| RODRIGUES DCC TARGETS DN | 38 | 105 | 1.72 | 4.24E-02 | Genes down-regulated in HCT8/S1 cells (colon cancer) which normally lack DCC [GeneID=9423] compared to those stably expressing wild type DCC off a plasmid vector. |
| REACTOME CIRCADIAN CLOCK | 37 | 66 | 1.99 | 5.38E-03 | Circadian Clock |
| BILD CTNNB1 ONCOGENIC SIGNATURE | 36 | 68 | 2.40 | 0.00E+00 | Genes selected in supervised analyses to discriminate cells expressing activated beta-catenin (CTNNB1) [GeneID=1499] oncogene from control cells expressing GFP. |

| | | | | | |
|---|---|---|---|---|---|
| DACOSTA UV RESPONSE VIA ERCC3 XPCS DN | 36 | 77 | 2.29 | 6.48E-05 | Genes exclusively down-regulated in fibroblasts expressing the XP/CS mutant form of ERCC3 [GeneID=2071] after high dose UVC irradiation. |
| DE YY1 TARGETS DN | 36 | 89 | 1.90 | 1.38E-02 | Genes down-regulated in SaOS-2 cells (osteosarcoma) upon knockdown of YY1 [GeneID=7528] by RNAi. |
| REACTOME DNA DAMAGE TELOMERE STRESS INDUCED SENESCENCE | 36 | 63 | 1.88 | 1.60E-02 | DNA Damage/Telomere Stress Induced Senescence |
| HOEBEKE LYMPHOID STEM CELL UP | 36 | 81 | 1.76 | 3.49E-02 | Genes up-regulated in the common lymphoid progenitor (CLP, defined as CD34+CD38-CD7+ [GeneID=947;952;924]) compared to a multipotent cord blood cell (defined as CD34+CD38+CD7-). |
| REACTOME DEPOSITION OF NEW CENPA CONTAINING NUCLEOSOMES AT THE CENTROMERE | 35 | 59 | 1.82 | 2.49E-02 | Deposition of new CENPA-containing nucleosomes at the centromere |
| BROWNE HCMV INFECTION 18HR UP | 35 | 146 | 1.76 | 3.38E-02 | Genes up-regulated in primary fibroblast cell culture after infection with HCMV (AD169 strain) at 18 h time point that were not up-regulated at the previous time point, 16 h. |
| BLANCO MELO BRONCHIAL EPITHELIAL CELLS INFLUENZA A INFECTION UP | 35 | 102 | 1.74 | 3.89E-02 | Genes up-regulated oninfection of normal human bronchial epithelial cells by Influenza A (MOI: 3, 12hpi) |
| EPPERT HSC R | 34 | 99 | 1.81 | 2.70E-02 | Genes up-regulated in human hematopoietic stem cell (HSC) enriched populations compared to committed progenitors and mature cells. |
| PLASARI TGFB1 SIGNALING VIA NFIC 1HR DN | 33 | 77 | 2.50 | 0.00E+00 | Genes down-regulated after 1 h of TGFB1 [GeneID=7040] stimulation in MEF cells (embryonic fibroblast) with NFIC [GeneID=4782] knockout vs wild type MEFs. |
| SENESE HDAC2 TARGETS UP | 32 | 98 | 1.76 | 3.42E-02 | Genes up-regulated in U2OS cells (osteosarcoma) upon knockdown of HDAC2 [GeneID=3066] by RNAi. |
| KIM MYCN AMPLIFICATION TARGETS DN | 31 | 82 | 2.03 | 4.09E-03 | Genes negatively correlated with amplifications of MYCN [GeneID=4613] in the SCLC (small cell lung cancer) cell lines. |

| | | | | | |
|---|---|---|---|---|---|
| SEIDEN ONCOGENESIS BY MET | 31 | 85 | 1.89 | 1.53E-02 | Genes changed in xenograft tumors formed by DLD-1 or DKO-4 cells (colon cancer) overexpressing MET [GeneID=4233]. |
| BOSCO ALLERGEN INDUCED TH2 ASSOCIATED MODULE | 31 | 128 | 1.72 | 4.42E-02 | Genes representing a co-expression network in atopic CD4 [GeneID=920] T lymphocyte responses. |
| CHIANG LIVER CANCER SUBCLASS UNANNOTATED UP | 30 | 65 | 2.16 | 4.91E-04 | Marker genes up-regulated in the 'unannotated' subclass of hepatocellular carcinoma (HCC) samples. |
| HOSHIDA LIVER CANCER SUBCLASS S2 | 30 | 101 | 1.84 | 2.26E-02 | Genes from 'subtype S2' signature of hepatocellular carcinoma (HCC): proliferation, MYC and AKT1 [GeneID=4609;207] activation. |
| WP MESODERMAL COMMITMENT PATHWAY | 28 | 102 | 2.12 | 1.29E-03 | Mesodermal commitment pathway |
| REACTOME ACTIVATION OF GENE EXPRESSION BY SREBF SREBP | 28 | 41 | 1.81 | 2.66E-02 | Activation of gene expression by SREBF (SREBP) |
| REACTOME RUNX1 REGULATES GENES INVOLVED IN MEGAKARYOCYTE DIFFERENTIATION AND PLATELET FUNCTION | 26 | 75 | 2.09 | 1.57E-03 | RUNX1 regulates genes involved in megakaryocyte differentiation and platelet function |
| MOLENAAR TARGETS OF CCND1 AND CDK4 DN | 26 | 54 | 1.85 | 2.12E-02 | Genes commonly down-regulated in SK-N-BE cells (neuroblastoma) after RNAi knockdown of CCND1 and CDK4 [GeneID=595;1019]. |
| REACTOME HDMS DEMETHYLATE HISTONES | 26 | 42 | 1.82 | 2.61E-02 | HDMs demethylate histones |
| DING LUNG CANCER EXPRESSION BY COPY NUMBER | 26 | 99 | 1.78 | 3.14E-02 | The lung adenocarcinoma TSP (tumor sequencing project) genes showing strong correlation between DNA copy number variation and gene expression. |
| NAKAYAMA SOFT TISSUE TUMORS PCA2 UP | 25 | 65 | 1.71 | 4.52E-02 | Top 100 probe sets contrubuting to the positive side of the 2nd principal component; associated with adipocytic differentiation. |
| REACTOME PKMTS METHYLATE HISTONE LYSINES | 24 | 63 | 2.37 | 0.00E+00 | PKMTs methylate histone lysines |
| AMIT EGF RESPONSE 60 HELA | 24 | 42 | 1.90 | 1.36E-02 | Genes whose expression peaked at 60 min after stimulation of HeLa cells with EGF [GeneID=1950]. |

| | | | | | |
|---|---|---|---|---|---|
| NAGASHIMA EGF SIGNALING UP | 24 | 52 | 1.80 | 2.76E-02 | Genes up-regulated in MCF7 cells (breast cancer) after stimulation with EGF [GeneID=1950]. |
| WP MECP2 AND ASSOCIATED RETT SYNDROME | 24 | 52 | 1.79 | 2.96E-02 | MECP2 and associated Rett syndrome |
| NIKOLSKY MUTATED AND AMPLIFIED IN BREAST CANCER | 24 | 71 | 1.76 | 3.38E-02 | Genes both mutated and amplified in a panel of 191 breast tumor samples. |
| WP PATHWAYS AFFECTED IN ADENOID CYSTIC CARCINOMA | 23 | 55 | 2.15 | 6.08E-04 | Pathways affected in adenoid cystic carcinoma |
| TANG SENESCENCE TP53 TARGETS DN | 23 | 49 | 1.89 | 1.50E-02 | Genes down-regulated in WI-38 cells (senescent primary fibroblasts) after inactivation of TP53 [GeneID=7157] by GSE56 polypeptide. |
| RHEIN ALL GLUCOCORTICOID THERAPY UP | 23 | 62 | 1.81 | 2.65E-02 | Genes up-regulated in ALL (acute lymphoblastic leukemia) blasts after 1 week of treatment with glucocorticoids. |
| PHONG TNF TARGETS UP | 23 | 54 | 1.79 | 3.03E-02 | Genes up-regulated in Calu-6 cells (lung cancer) at 1 h time point after TNF [GeneID=7124] treatment. |
| REACTOME DDX58 IFIH1 MEDIATED INDUCTION OF INTERFERON ALPHA BETA | 23 | 63 | 1.76 | 3.39E-02 | DDX58/IFIH1-mediated induction of interferon-alpha/beta |
| KANG DOXORUBICIN RESISTANCE UP | 23 | 53 | 1.74 | 3.78E-02 | Genes up-regulated in gastric cancer cell lines: doxorubicin [PubChem=31703] resistant vs sensitive. |
| GEORGES CELL CYCLE MIR192 TARGETS | 22 | 59 | 1.89 | 1.51E-02 | Experimentally validated direct targets of MIR192 [GeneID=406967] microRNA; MIR192 caused cell cycle arrest in HCT116 cells (colon cancer). |
| SHEPARD BMYB TARGETS | 22 | 53 | 1.86 | 2.08E-02 | Human orthologs of BMYB [GeneID=4605] target genes in zebra fish, identified as commonly changed in the BMYB loss of function mutant crb ('crush and burn') and after knockdown of BMYB by morpholino. |
| WP ONCOSTATIN M SIGNALING PATHWAY | 22 | 61 | 1.80 | 2.70E-02 | Oncostatin M signaling pathway |
| REACTOME SUMOYLATION OF TRANSCRIPTION COFACTORS | 22 | 42 | 1.79 | 3.07E-02 | SUMOylation of transcription cofactors |

| | | | | | |
|---|---|---|---|---|---|
| GAVIN FOXP3 TARGETS CLUSTER P7 | 22 | 76 | 1.73 | 4.13E-02 | Cluster P7 of genes with similar expression profiles in peripheral T lymphocytes after FOXP3 [GeneID=50943] loss of function (LOF). |
| HADDAD T LYMPHOCYTE AND NK PROGENITOR UP | 22 | 61 | 1.72 | 4.43E-02 | Genes up-regulated in hematopoietic progenitor cells (HPC) of T lymphocyte and NK (natural killer) lineage. |
| LINDSTEDT DENDRITIC CELL MATURATION B | 21 | 47 | 2.52 | 0.00E+00 | Maturation of monocyte-derived dendritic cells (DC) in response to inflammatory stimuli: genes up-regulated both at 8 hr and 48 hr after the stimulation (cluster B). |
| NAGASHIMA NRG1 SIGNALING DN | 21 | 39 | 2.04 | 3.65E-03 | Genes down-regulated in MCF7 cells (breast cancer) after stimulation with NRG1 [GeneID=3084]. |
| IKEDA MIR133 TARGETS UP | 21 | 41 | 1.95 | 8.19E-03 | Genes up-regulated in hypertrophic hearts (due to expression of constitutively active form of PPP3CA [GeneID=5530]) and predicted to be targets of miR-133 microRNA. |
| REACTOME HEME SIGNALING | 21 | 43 | 1.87 | 1.82E-02 | Heme signaling |
| HOOI ST7 TARGETS UP | 21 | 67 | 1.83 | 2.37E-02 | Genes up-regulated in PC-3 cells (prostate cancer) stably expressing ST7 [GeneID=7982] off a plasmid vector. |
| LU IL4 SIGNALING | 21 | 79 | 1.83 | 2.41E-02 | Genes up-regulated in peripheral B lymphocytes after incubation with IL4 [GeneID=3565] for 4 h. |
| SCHAEFFER PROSTATE DEVELOPMENT AND CANCER BOX4 DN | 21 | 33 | 1.80 | 2.75E-02 | Early prostate development genes (down-regulated at 6 hr dihydrotestosterone [PubChem=10635]) which are also down-regulated in localized vs metastatic prostate cancers. |
| GENTILE UV RESPONSE CLUSTER D2 | 20 | 33 | 2.00 | 4.93E-03 | Cluster d2: genes down-regulated consistently in WS1 cells (fibroblast) between 6 h and 24 h after irradiation with high dose UV-C. |
| REACTOME CA2 PATHWAY | 20 | 53 | 1.97 | 6.95E-03 | Ca2+ pathway |
| PID HES HEY PATHWAY | 20 | 37 | 1.84 | 2.26E-02 | Notch-mediated HES/HEY network |
| NATSUME RESPONSE TO INTERFERON BETA UP | 20 | 51 | 1.83 | 2.39E-02 | Genes up-regulated in T98 cells (glioma) 48 h after treatment with interferon beta. |
| WP HEAD AND NECK SQUAMOUS CELL CARCINOMA | 20 | 70 | 1.75 | 3.66E-02 | Head and neck squamous cell carcinoma |

| | | | | | |
|---|---|---|---|---|---|
| PID IL12 2PATHWAY | 20 | 50 | 1.71 | 4.68E-02 | IL12-mediated signaling events |
| BURTON ADIPOGENESIS 12 | 19 | 31 | 2.36 | 0.00E+00 | Strongly down-regulated at 2 h during differentiation of 3T3-L1 cells (fibroblast) into adipocytes. |
| WP HISTONE MODIFICATIONS | 19 | 58 | 2.10 | 1.60E-03 | Histone modifications |
| YAO TEMPORAL RESPONSE TO PROGESTERONE CLUSTER 2 | 19 | 37 | 1.79 | 3.03E-02 | Genes co-regulated in uterus during a time course response to progesterone [PubChem=5994]: SOM cluster 2. |
| ZHAN MULTIPLE MYELOMA PR DN | 18 | 41 | 1.92 | 1.10E-02 | Top 50 down-regulated genes in cluster PR of multiple myeloma samples characterized by increased expression of proliferation and cell cycle genes. |
| BILD SRC ONCOGENIC SIGNATURE | 18 | 52 | 1.86 | 2.01E-02 | Genes selected in supervised analyses to discriminate cells expressing c-Src (CSK) [GeneID=1445] from control cells expressing GFP. |
| ZHAN MULTIPLE MYELOMA PR UP | 18 | 42 | 1.85 | 2.12E-02 | Top 50 up-regulated genes in cluster PR of multiple myeloma samples characterized by increased expression of proliferation and cell cycle genes. |
| REACTOME TRANSCRIPTIONAL REGULATION BY E2F6 | 18 | 34 | 1.83 | 2.38E-02 | Transcriptional Regulation by E2F6 |
| REACTOME TRANSCRIPTIONAL REGULATION BY MECP2 | 17 | 43 | 2.03 | 4.20E-03 | Transcriptional Regulation by MECP2 |
| GREENBAUM E2A TARGETS UP | 17 | 32 | 1.85 | 2.11E-02 | Genes up-regulated in pre-B lymphocytes upon Cre-Lox knockout of E2A [GeneID=6929]. |
| WP LEPTIN SIGNALING PATHWAY | 17 | 73 | 1.81 | 2.66E-02 | Leptin signaling pathway |
| PEDERSEN METASTASIS BY ERBB2 ISOFORM 1 | 17 | 40 | 1.79 | 2.85E-02 | Genes regulated in MCF7 cells (breast cancer) by expression of full-length and truncated (611-CTF) forms of ERBB2 [GeneID=2064] at both 15 h and 60 h time points. |
| BILBAN B CLL LPL DN | 17 | 36 | 1.71 | 4.60E-02 | Genes down-regulated in B-CLL (B-cell chronic leukemia) samples expressing high levels of LPL [GeneID=4023] compared with those expressing low levels of the gene. |
| TURASHVILI BREAST LOBULAR CARCINOMA VS | 16 | 51 | 1.99 | 5.25E-03 | Genes down-regulated in lobular carcinoma vs normal ductal breast cells. |

| | | | | | |
|---|---|---|---|---|---|
| DUCTAL NORMAL DN | | | | | |
| REACTOME SIGNALING BY FGFR1 IN DISEASE | 16 | 31 | 1.97 | 6.47E-03 | Signaling by FGFR1 in disease |
| GRAHAM NORMAL QUIESCENT VS NORMAL DIVIDING UP | 16 | 48 | 1.81 | 2.63E-02 | Genes up-regulated in quiescent vs dividing CD34+ [GeneID=8842] cells isolated from peripheral blood of normal donors. |
| SCHMIDT POR TARGETS IN LIMB BUD UP | 16 | 24 | 1.78 | 3.17E-02 | Genes up-regulated in E12.5 forelimb buds with POR [GeneID=5447] knockout. |
| REACTOME RUNX1 INTERACTS WITH CO FACTORS WHOSE PRECISE EFFECT ON RUNX1 TARGETS IS NOT KNOWN | 16 | 36 | 1.72 | 4.46E-02 | RUNX1 interacts with co-factors whose precise effect on RUNX1 targets is not known |
| MARTIN INTERACT WITH HDAC | 16 | 38 | 1.71 | 4.66E-02 | Interaction partners of class IIa histone deacetylases (HDAC). |
| VILIMAS NOTCH1 TARGETS UP | 15 | 38 | 2.24 | 1.84E-04 | Genes up-regulated in bone marrow progenitors by constitutively active NOTCH1 [GeneID=4851]. |
| WP RETT SYNDROME CAUSING GENES | 15 | 36 | 1.97 | 6.53E-03 | Rett syndrome causing genes |
| MA MYELOID DIFFERENTIATION DN | 15 | 33 | 1.75 | 3.70E-02 | Genes down-regulated during myeloid differentiation induced by tretinoin (ATRA) [PubChem=444795] and IL3 [GeneID=3652] in the EML cell line (myeloid progenitor). |
| WP MODULATORS OF TCR SIGNALING AND T CELL ACTIVATION | 15 | 55 | 1.72 | 4.24E-02 | Modulators of TCR signaling and T cell activation |
| WP MELATONIN METABOLISM AND EFFECTS | 15 | 28 | 1.71 | 4.64E-02 | Melatonin metabolism and effects |
| PID GMCSF PATHWAY | 15 | 33 | 1.70 | 4.82E-02 | GMCSF-mediated signaling events |
| REACTOME REGULATION OF MECP2 EXPRESSION AND ACTIVITY | 14 | 29 | 2.31 | 0.00E+00 | Regulation of MECP2 expression and activity |
| REACTOME FGFR1 MUTANT RECEPTOR ACTIVATION | 14 | 24 | 2.11 | 1.32E-03 | FGFR1 mutant receptor activation |
| WP THYMIC STROMAL LYMPHOPOIETIN TSLP SIGNALING PATHWAY | 14 | 44 | 2.00 | 4.83E-03 | Thymic stromal lymphopoietin (TSLP) signaling pathway |

| | | | | | |
|---|---|---|---|---|---|
| REICHERT MITOSIS LIN9 TARGETS | 14 | 27 | 1.83 | 2.43E-02 | Genes with known mitosis function that were down-regulated in MEF cells (embryonic fibroblast) upon knockout of LIN9 [GeneID=286826]. |
| WP MBDNF AND PROBDNF REGULATION OF GABA NEUROTRANSMISSION | 13 | 21 | 1.88 | 1.62E-02 | mBDNF and proBDNF regulation of GABA neurotransmission |
| MCLACHLAN DENTAL CARIES DN | 13 | 45 | 1.73 | 4.20E-02 | Genes down-regulated in pulpal tissue extracted from carious teeth. |
| ZHENG FOXP3 TARGETS IN T LYMPHOCYTE DN | 12 | 34 | 2.08 | 2.27E-03 | Genes with promoters bound by FOXP3 [GeneID=50943] and which are down-regulated only in mature (peripheral blood) regulatory CD4+ [GeneID=920] T lymphocytes. |
| WIERENGA STAT5A TARGETS GROUP2 | 12 | 45 | 1.86 | 2.02E-02 | Genes up-regulated in a linear fashion in CD34+ [GeneID=947] cells upon increasing activity levels of STAT5A [GeneID=6776]; predominant long-term growth and self-renewal phenotype. |
| YANAGIHARA ESX1 TARGETS | 12 | 23 | 1.83 | 2.44E-02 | Genes down-regulated in U2-OS Tet-On cells (osteosarcoma) after induction of ESX1 [GeneID=80712] expression. |
| AMIT EGF RESPONSE 60 MCF10A | 12 | 34 | 1.78 | 3.14E-02 | Genes whose expression peaked at 60 min after stimulation of MCF10A cells with EGF [GeneID=1950]. |
| REACTOME FLT3 SIGNALING IN DISEASE | 12 | 27 | 1.77 | 3.30E-02 | FLT3 signaling in disease |
| REACTOME SIGNALING BY FLT3 FUSION PROTEINS | 11 | 18 | 1.98 | 5.80E-03 | Signaling by FLT3 fusion proteins |
| REACTOME SIGNALING BY CYTOSOLIC FGFR1 FUSION MUTANTS | 11 | 18 | 1.96 | 7.51E-03 | Signaling by cytosolic FGFR1 fusion mutants |
| PID IL23 PATHWAY | 11 | 26 | 1.90 | 1.34E-02 | IL23-mediated signaling events |
| LI WILMS TUMOR ANAPLASTIC UP | 11 | 16 | 1.90 | 1.36E-02 | Selected up-regulated genes distinguishing between Wilms tumors of different histological types: anaplastic vs favorable histology. |
| PID IL12 STAT4 PATHWAY | 11 | 24 | 1.86 | 2.08E-02 | IL12 signaling mediated by STAT4 |
| REACTOME LAMININ INTERACTIONS | 11 | 23 | 1.78 | 3.09E-02 | Laminin interactions |

| | | | | | |
|---|---|---|---|---|---|
| REACTOME DEADENYLATION DEPENDENT MRNA DECAY | 11 | 55 | 1.76 | 3.43E-02 | Deadenylation-dependent mRNA decay |
| REACTOME RHOV GTPASE CYCLE | 11 | 37 | 1.76 | 3.50E-02 | RHOV GTPase cycle |
| GAZDA DIAMOND BLACKFAN ANEMIA MYELOID UP | 11 | 29 | 1.74 | 3.81E-02 | Genes up-regulated in myeloid progenitor cells isolated from bone marrow of patients with Diamond-Blackfan anemia (DBA) and mutated RPS19 [GeneID=6223]. |
| PID HNF3A PATHWAY | 11 | 24 | 1.74 | 3.80E-02 | FOXA1 transcription factor network |
| REACTOME IKK COMPLEX RECRUITMENT MEDIATED BY RIP1 | 11 | 22 | 1.74 | 3.81E-02 | IKK complex recruitment mediated by RIP1 |
| WP PHOSPHODIESTERASES IN NEURONAL FUNCTION | 11 | 27 | 1.72 | 4.33E-02 | Phosphodiesterases in neuronal function |
| YUAN ZNF143 PARTNERS | 11 | 21 | 1.71 | 4.50E-02 | Proteins associated with ZNF143 [GeneID=7702] in HeLa cells, based on MudPIT analysis. |
| REACTOME STRIATED MUSCLE CONTRACTION | 10 | 20 | 1.95 | 8.20E-03 | Striated Muscle Contraction |
| MARSON FOXP3 TARGETS STIMULATED UP | 10 | 22 | 1.93 | 1.01E-02 | Genes with promoters bound by FOXP3 [GeneID=50943], dependent on it, and up-regulated in hybridoma cells stimulated by PMA [PubChem=4792] and ionomycin [PubChem=3733]. |
| REACTOME SIGNALING BY KIT IN DISEASE | 10 | 20 | 1.91 | 1.33E-02 | Signaling by KIT in disease |
| BIOCARTA CTCF PATHWAY | 10 | 23 | 1.76 | 3.39E-02 | CTCF: First Multivalent Nuclear Factor |
| REACTOME INTERACTION BETWEEN L1 AND ANKYRINS | 10 | 19 | 1.75 | 3.65E-02 | Interaction between L1 and Ankyrins |
| REACTOME RORA ACTIVATES GENE EXPRESSION | 10 | 17 | 1.74 | 3.88E-02 | RORA activates gene expression |
| CAFFAREL RESPONSE TO THC UP | 10 | 28 | 1.72 | 4.44E-02 | Genes up-regulated in EVSA-T cells (breast cancer) treated THC (delta-9-tetrahydrocannabinol) [PubChem=6610319]. |
| TIAN TNF SIGNALING VIA NFKB | 9 | 22 | 2.02 | 3.89E-03 | Genes modulated in HeLa cells (cervical carcinoma) by TNF [GeneID=7124] via NFKB pathway. |

| | | | | | |
|---|---|---|---|---|---|
| REACTOME FORMATION OF SENESCENCE ASSOCIATED HETEROCHROMATIN FOCI SAHF | 9 | 16 | 1.89 | 1.53E-02 | Formation of Senescence-Associated Heterochromatin Foci (SAHF) |
| REACTOME DEADENYLATION OF MRNA | 9 | 25 | 1.84 | 2.27E-02 | Deadenylation of mRNA |
| WP INITIATION OF TRANSCRIPTION AND TRANSLATION ELONGATION AT THE HIV1 LTR | 9 | 31 | 1.82 | 2.57E-02 | Initiation of transcription and translation elongation at the HIV-1 LTR |
| WP NEOVASCULARISATION PROCESSES | 9 | 32 | 1.78 | 3.07E-02 | Neovascularisation processes |
| WP LDL INFLUENCE ON CD14 AND TLR4 | 9 | 22 | 1.77 | 3.20E-02 | LDL- influence on CD14 and TLR4 |
| REACTOME CD28 CO STIMULATION | 9 | 32 | 1.75 | 3.71E-02 | CD28 co-stimulation |
| XIE LT HSC S1PR3 OE UP | 9 | 23 | 1.70 | 4.76E-02 | Genes upregulated in long-term hematopoietic stem cells (CD34+,CD38_,CD45RA_,CD90+,CD49f+) upon overexpression of Sphingosine-1-Phosphate Receptor 3 (S1PR3) |
| WP INTERACTOME OF POLYCOMB REPRESSIVE COMPLEX 2 PRC2 | 8 | 16 | 2.07 | 2.54E-03 | Interactome of polycomb repressive complex 2 (PRC2) |
| WP INTERACTIONS BETWEEN IMMUNE CELLS AND MICRORNAS IN TUMOR MICROENVIRONMENT | 8 | 26 | 2.00 | 5.25E-03 | Interactions between immune cells and microRNAs in tumor microenvironment |
| REACTOME NOTCH3 INTRACELLULAR DOMAIN REGULATES TRANSCRIPTION | 8 | 20 | 1.81 | 2.68E-02 | NOTCH3 Intracellular Domain Regulates Transcription |
| REACTOME EPHA MEDIATED GROWTH CONE COLLAPSE | 8 | 17 | 1.80 | 2.74E-02 | EPHA-mediated growth cone collapse |
| WP AMPLIFICATION AND EXPANSION OF ONCOGENIC PATHWAYS AS METASTATIC TRAITS | 8 | 15 | 1.80 | 2.75E-02 | Amplification and expansion of oncogenic pathways as metastatic traits |
| WP GASTRIC CANCER NETWORK 1 | 8 | 23 | 1.80 | 2.77E-02 | Gastric cancer network 1 |
| REACTOME NOTCH HLH | 8 | 27 | 1.78 | 3.13E-02 | Notch-HLH transcription pathway |

| | | | | | |
|---|---|---|---|---|---|
| TRANSCRIPTION PATHWAY | | | | | |
| COLLIS PRKDC SUBSTRATES | 8 | 16 | 1.77 | 3.19E-02 | Substrates of PRKDC [GeneID=5591]. |
| WEST ADRENOCORTICAL TUMOR MARKERS UP | 8 | 17 | 1.73 | 4.04E-02 | Top up-regulated genes in pediatric adrenocortical tumors (ACT) compared to the normal tissue. |
| PID CD40 PATHWAY | 8 | 27 | 1.72 | 4.35E-02 | CD40/CD40L signaling |
| BIOCARTA TNFR2 PATHWAY | 8 | 17 | 1.72 | 4.33E-02 | TNFR2 Signaling Pathway |
| REACTOME TRAF6 MEDIATED NF KB ACTIVATION | 8 | 22 | 1.71 | 4.58E-02 | TRAF6 mediated NF-kB activation |
| BIOCARTA IL7 PATHWAY | 8 | 16 | 1.71 | 4.56E-02 | IL-7 Signal Transduction |
| REACTOME CD28 DEPENDENT PI3K AKT SIGNALING | 7 | 22 | 2.04 | 3.77E-03 | CD28 dependent PI3K/Akt signaling |
| IVANOVA HEMATOPOIESIS STEM CELL SHORT TERM | 7 | 15 | 2.02 | 3.98E-03 | Genes in the expression cluster 'ST-HSC Shared': up-regulated in short term hematopoietic stem cells (ST-HSC) from adult bone marrow and fetal liver. |
| PETROVA PROX1 TARGETS UP | 7 | 23 | 1.85 | 2.10E-02 | Genes specific to LEC (lymphatic endothelium cells) induced in BEC (blood endothelium cells) by expression of PROX1 [GeneID=5629] off adenovirus vector. |
| REACTOME NOTCH4 INTRACELLULAR DOMAIN REGULATES TRANSCRIPTION | 7 | 17 | 1.83 | 2.36E-02 | NOTCH4 Intracellular Domain Regulates Transcription |
| WANG IMMORTALIZED BY HOXA9 AND MEIS1 DN | 7 | 21 | 1.82 | 2.59E-02 | Down-regulated genes in myeloid progenitors immortalized by HOXA9 [GeneID=3205] vs those immortalized by HOXA9 and MEIS1 [GeneID=4211]. |
| KEGG DORSO VENTRAL AXIS FORMATION | 7 | 19 | 1.72 | 4.35E-02 | Dorso-ventral axis formation |
| GAUSSMANN MLL AF4 FUSION TARGETS B UP | 6 | 17 | 1.91 | 1.32E-02 | Up-regulated genes from the set B (Fig. 5a): specific signature shared by cells expressing either AF4-MLL or MLL-AF4 [GeneID=4299;4297] fusion proteins. |
| REACTOME INTERLEUKIN 20 FAMILY SIGNALING | 6 | 16 | 1.83 | 2.45E-02 | Interleukin-20 family signaling |
| REACTOME NEPHRIN FAMILY INTERACTIONS | 6 | 18 | 1.80 | 2.75E-02 | Nephrin family interactions |

| | | | | | |
|---|---|---|---|---|---|
| WP OVERVIEW OF INTERFERONSMEDIATED SIGNALING PATHWAY | 6 | 16 | 1.77 | 3.20E-02 | Overview of interferons-mediated signaling pathway |
| WP EBSTEINBARR VIRUS LMP1 SIGNALING | 6 | 20 | 1.72 | 4.33E-02 | Ebstein-Barr virus LMP1 signaling |
| WP HIPPOYAP SIGNALING PATHWAY | 5 | 21 | 1.78 | 3.09E-02 | Hippo-Yap signaling pathway |
| NIKOLSKY OVERCONNECTED IN BREAST CANCER | 5 | 15 | 1.78 | 3.07E-02 | Overconnected mutated transcription factors regulating genes within the breast cancer amplicome. |
| REACTOME INTERLEUKIN 6 FAMILY SIGNALING | 5 | 17 | 1.76 | 3.36E-02 | Interleukin-6 family signaling |
| PID NFKAPPAB ATYPICAL PATHWAY | 4 | 17 | 1.78 | 3.15E-02 | Atypical NF-kappaB pathway |

**Table S3. Top gene ontology and KEGG pathways enriched with significantly altered genes upon CAC+HDL compared to Ctr-HDL.**

| GO term | In Data | In Pathway | Expected | FDR | Pathway Hits | Description |
|---|---|---|---|---|---|---|
| GO:2000181 | 8 | 107 | 1.76 | 3.98E-02 | Emilin1;Krit1;Atp2b4;Klf2;Adrb2;Pde3b;Pparg;Hhip | negative regulation of blood vessel morphogenesis |
| GO:1905664 | 4 | 16 | 0.26 | 2.04E-02 | Fyn;Adrb2;Akap5;Adrb1 | regulation of calcium ion import across plasma membrane |
| GO:1904996 | 5 | 27 | 0.44 | 1.74E-02 | Nfat5;Fut7;Ets1;Mdk;Gp1ba | positive regulation of leukocyte adhesion to vascular endothelial cell |
| GO:1904994 | 5 | 35 | 0.58 | 2.95E-02 | Nfat5;Fut7;Ets1;Mdk;Gp1ba | regulation of leukocyte adhesion to vascular endothelial cell |
| GO:1904062 | 13 | 248 | 4.09 | 2.95E-02 | Fyn;Stk39;Kcnj2;Tspan13;Adrb2;Akap5;Prkce;Adrb1;Ank2;Tesc;Plcb1;Cacnb2;Tmem38a | regulation of monoatomic cation transmembrane transport |
| GO:1904036 | 7 | 73 | 1.20 | 2.56E-02 | Igf1;Hmox1;Krit1;Ndnf;Gas6;Alms1;Mdk | negative regulation of epithelial cell apoptotic process |
| GO:1904035 | 9 | 124 | 2.04 | 2.67E-02 | Igf1;Hmox1;Krit1;Ndnf;Gas6;Alms1;Ager;Mdk;Wnt11 | regulation of epithelial cell apoptotic process |
| GO:1903708 | 11 | 203 | 3.34 | 4.75E-02 | Il2ra;Pou4f1;Vsir;Tnfsf4;Rasgrp1;Gas6;Fos;Ager;Cd46;Tesc;Mdk | positive regulation of hemopoiesis |

| | | | | | | |
|---|---|---|---|---|---|---|
| GO:1903706 | 18 | 431 | 7.10 | 3.42E-02 | Il2ra;Pou4f1;Vsir;Scin;Tnfsf4;Clec2i;Rasgrp1;Gas6;Tmem178;Csf3;Fos;Ets1;Ager;Cd46;Tesc;Ccr6;Mdk;Tnfrsf11b | regulation of hemopoiesis |
| GO:1903555 | 12 | 209 | 3.44 | 2.50E-02 | Igf1;Vsir;Arid5a;Nod1;Rasgrp1;Gas6;Ager;Ghrl;Il33;Oas1c;Tlr5;Tnfrsf8 | regulation of tumor necrosis factor superfamily cytokine production |
| GO:1903054 | 4 | 12 | 0.20 | 1.18E-02 | Emilin1;Dpp4;Pparg;Tgfbr3 | negative regulation of extracellular matrix organization |
| GO:1903039 | 19 | 275 | 4.53 | 4.81E-04 | Nr4a3;Nfat5;Il2ra;Igf1;Vsir;Cd276;Cd80;Tnfsf4;Dpp4;Rasgrp1;Fut7;Il12rb1;Icos;Ets1;Ager;Cd46;Mdk;Gp1ba;Cd5 | positive regulation of leukocyte cell-cell adhesion |
| GO:1903037 | 20 | 376 | 6.19 | 3.69E-03 | Nr4a3;Nfat5;Il2ra;Igf1;Vsir;Cd276;Cd80;Tnfsf4;Dpp4;Rasgrp1;Fut7;Il12rb1;Btla;Icos;Ets1;Ager;Cd46;Mdk;Gp1ba;Cd5 | regulation of leukocyte cell-cell adhesion |
| GO:1902107 | 11 | 203 | 3.34 | 4.75E-02 | Il2ra;Pou4f1;Vsir;Tnfsf4;Rasgrp1;Gas6;Fos;Ager;Cd46;Tesc;Mdk | positive regulation of leukocyte differentiation |
| GO:1901343 | 8 | 108 | 1.78 | 4.05E-02 | Emilin1;Krit1;Atp2b4;Klf2;Adrb2;Pde3b;Pparg;Hhip | negative regulation of vasculature development |
| GO:1901342 | 16 | 298 | 4.91 | 1.23E-02 | Emp2;Hmox1;Emilin1;Krit1;Atp2b4;Klf2;Adrb2;Pde3b;Pparg;Fgf1;Ets1;Ghrl;Mdk;Hhip;Vegfa;Reck | regulation of vasculature development |
| GO:0120162 | 8 | 98 | 1.61 | 2.67E-02 | Cmklr1;Lepr;Adrb2;Per2;Alms1;Adrb1;Ghrl;Vegfa | positive regulation of cold-induced thermogenesis |
| GO:0106056 | 5 | 39 | 0.64 | 4.06E-02 | Nfat5;Igf1;Atp2b4;Akap5;Tbc1d10c | regulation of calcineurin-mediated signaling |
| GO:0106015 | 3 | 9 | 0.15 | 3.63E-02 | Siglecg;Mdk;Il33 | negative regulation of inflammatory response to wounding |
| GO:0099173 | 13 | 252 | 4.15 | 3.33E-02 | Fyn;Stk38l;Ptprf;Lama5;Cdkl5;Lrrc4;Farp1;Ghrl;Itga3;Caprin2;Zdhhc2;Lrrtm2;Grip2 | postsynapse organization |
| GO:0090594 | 5 | 28 | 0.46 | 1.77E-02 | Hmox1;Pparg;Siglecg;Mdk;Il33 | inflammatory response to wounding |

| | | | | | | |
|---|---|---|---|---|---|---|
| GO:0090287 | 16 | 322 | 5.30 | 1.84E-02 | Fam20c;Vsir;Kcp;Emilin1;Ptprf;Peg10;Atp2b4;Sinhcaf;Pparg;Fgf1;Tgfbr3;Spry1;Rbpms2;Itga3;Hhip;Vegfa | regulation of cellular response to growth factor stimulus |
| GO:0090257 | 16 | 249 | 4.10 | 3.69E-03 | Nr4a3;Igf1;Tifab;Atp2b4;Kcnj2;Adrb2;Pparg;Grcc10;Tnni3;Pi16;Adrb1;Ank2;Ghrl;Tnnt3;Adra2c;Tmem38a | regulation of muscle system process |
| GO:0090132 | 16 | 316 | 5.21 | 1.74E-02 | Emp2;Igf1;Hmox1;Krit1;Atp2b4;Dpp4;Plekhg5;Pparg;Fgf1;Tgfbr3;Prkce;Ets1;Ager;Itga3;Ccr6;Vegfa | epithelium migration |
| GO:0090130 | 16 | 322 | 5.30 | 1.84E-02 | Emp2;Igf1;Hmox1;Krit1;Atp2b4;Dpp4;Plekhg5;Pparg;Fgf1;Tgfbr3;Prkce;Ets1;Ager;Itga3;Ccr6;Vegfa | tissue migration |
| GO:0085029 | 6 | 47 | 0.77 | 2.04E-02 | Emilin1;Gas6;Pparg;Tgfbr3;Col1a2;Has3 | extracellular matrix assembly |
| GO:0071880 | 4 | 21 | 0.35 | 3.63E-02 | Atp2b4;Adrb2;Adrb1;Adra2c | adenylate cyclase-activating adrenergic receptor signaling pathway |
| GO:0071706 | 12 | 209 | 3.44 | 2.50E-02 | Igf1;Vsir;Arid5a;Nod1;Rasgrp1;Gas6;Ager;Ghrl;Il33;Oas1c;Tlr5;Tnfrsf8 | tumor necrosis factor superfamily cytokine production |
| GO:0071349 | 4 | 10 | 0.16 | 8.22E-03 | Stat4;Il12rb2;Il12rb1;Plcb1 | cellular response to interleukin-12 |
| GO:0070884 | 5 | 38 | 0.63 | 3.86E-02 | Nfat5;Igf1;Atp2b4;Akap5;Tbc1d10c | regulation of calcineurin-NFAT signaling cascade |
| GO:0070671 | 4 | 12 | 0.20 | 1.18E-02 | Stat4;Il12rb2;Il12rb1;Plcb1 | response to interleukin-12 |
| GO:0070661 | 18 | 397 | 6.54 | 2.04E-02 | Il2ra;Emp2;Igf1;Vsir;Fyn;Cd276;Cd80;Stat4;Tnfsf4;Clec2i;Satb1;Rasgrp1;Siglecg;Il12rb1;Btla;Ager;Cd46;Il33 | leukocyte proliferation |
| GO:0070588 | 16 | 314 | 5.17 | 1.74E-02 | Fyn;Atp2b4;Tspan13;Adrb2;Gas6;Akap5;Prkce;Adrb1;Cacna2d2;Ank2;Slc24a1;Plcb1;Gp1ba;Cacna2d4;Cacnb2;Tmem38a | calcium ion transmembrane transport |
| GO:0060291 | 9 | 114 | 1.88 | 2.04E-02 | Akap5;Prkcg;Adrb1;Slc24a1;Ager;Ntrk | long-term synaptic potentiation |

| | | | | | | |
|---|---|---|---|---|---|---|
| | | | | | 2;Zdhhc2;Lrrtm2;Prrt1 | |
| GO:0051962 | 20 | 367 | 6.05 | 3.69E-03 | Igf1;Ptprf;Neurl1a;Pparg;Mag;Hapln3;Akap5;Cdkl5;Dmrta2;Cdon;Tenm4;Map1b;Ntrk2;Ghrl;Mdk;Il33;Caprin2;Dixdc1;Vegfa;Lrrtm2 | positive regulation of nervous system development |
| GO:0051402 | 18 | 356 | 5.86 | 1.18E-02 | Nr4a3;Pou4f1;Hmox1;Fyn;Ptprf;En2;Aatk;Unc5b;Egr1;Ndnf;Mag;Prkcg;Tox3;Ager;Ntrk2;Mdk;Diablo;Vegfa | neuron apoptotic process |
| GO:0051251 | 15 | 330 | 5.44 | 3.98E-02 | Il2ra;Igf1;Vsir;Cd276;Cd80;Tnfsf4;Dpp4;Rasgrp1;Gas6;Il12rb1;Icos;Ager;Cd46;Mdk;Cd5 | positive regulation of lymphocyte activation |
| GO:0050900 | 17 | 372 | 6.13 | 2.38E-02 | Cmklr1;Emp2;Emilin1;Stk39;Tnfsf4;Dpp4;Ch25h;Gas6;Fut7;Ager;Itga3;Plcb1;Ccr6;Mdk;Il33;Gp1ba;Vegfa | leukocyte migration |
| GO:0050890 | 21 | 363 | 5.98 | 1.30E-03 | Igf1;Tifab;Slc6a4;Pde1b;Cbr3;Adrb2;Egr1;Grcc10;Slc6a1;Fos;Prkcg;Adrb1;Ager;Ntrk2;Ghrl;Itga3;Plcb1;Mdk;Nptx2;Chst10;Prrt1 | cognition |
| GO:0050878 | 16 | 348 | 5.73 | 2.76E-02 | Nfat5;Emp2;F5;Tifab;Slc6a4;Emilin1;Stk39;Neurl1a;Adrb2;Gas6;Prkce;Adrb1;Gp1ba;Adra2c;Plat;Vegfa | regulation of body fluid levels |
| GO:0050870 | 14 | 249 | 4.10 | 1.74E-02 | Il2ra;Igf1;Vsir;Cd276;Cd80;Tnfsf4;Dpp4;Rasgrp1;Il12rb1;Icos;Ager;Cd46;Mdk;Cd5 | positive regulation of T cell activation |
| GO:0050867 | 18 | 387 | 6.38 | 1.74E-02 | Nr4a3;Il2ra;Igf1;Vsir;Cd276;Cd80;Tnfsf4;Dpp4;Rasgrp1;Gas6;Il12rb1;Icos;Ager;Cd46;Mdk;Il33;Gp1ba;Cd5 | positive regulation of cell activation |
| GO:0050863 | 17 | 382 | 6.29 | 2.67E-02 | Il2ra;Igf1;Vsir;Cd276;Cd80;Tnfsf4;Clec2i;Dpp4;Rasgrp1;Il12rb1;Btla;Icos;Ager;Cd46;Ccr6;Mdk;Cd5 | regulation of T cell activation |

| | | | | | | |
|---|---|---|---|---|---|---|
| GO:0050798 | 7 | 50 | 0.82 | 8.91E-03 | Il2ra;Igf1;Fyn;Tnfsf4;Satb1;Il12rb1;Ager | activated T cell proliferation |
| GO:0050769 | 18 | 307 | 5.06 | 3.69E-03 | Igf1;Ptprf;Neurl1a;Pparg;Mag;Hapln3;Akap5;Cdkl5;Dmrta2;Cdon;Tenm4;Map1b;Ntrk2;Mdk;Il33;Caprin2;Dixdc1;Vegfa | positive regulation of neurogenesis |
| GO:0050767 | 20 | 459 | 7.56 | 1.74E-02 | Pou4f1;Igf1;Ptprf;Neurl1a;Per2;Pparg;Mag;Hapln3;Akap5;Cdkl5;Dmrta2;Cdon;Tenm4;Map1b;Ntrk2;Mdk;Il33;Caprin2;Dixdc1;Vegfa | regulation of neurogenesis |
| GO:0048511 | 16 | 336 | 5.54 | 2.37E-02 | Crem;Igf1;Slc6a4;Lepr;Egr1;Per2;Pparg;Rbm4;Rbm4b;Prkcg;Adrb1;Prokr1;Suv39h2;Ntrk2;Ghrl;Mdk | rhythmic process |
| GO:0048167 | 13 | 240 | 3.95 | 2.50E-02 | Neurl1a;Egr1;Akap5;Map1b;Prkcg;Adrb1;Slc24a1;Ager;Ntrk2;Zdhhc2;Gnal;Lrrtm2;Prrt1 | regulation of synaptic plasticity |
| GO:0046651 | 16 | 355 | 5.85 | 3.27E-02 | Il2ra;Emp2;Igf1;Vsir;Fyn;Cd276;Cd80;Tnfsf4;Clec2i;Satb1;Rasgrp1;Siglecg;Il12rb1;Btla;Ager;Cd46 | lymphocyte proliferation |
| GO:0046530 | 7 | 79 | 1.30 | 3.55E-02 | Ppp2r3a;Samd11;Cep290;Alms1;Ntrk2;Nphp4;Vegfa | photoreceptor cell differentiation |
| GO:0045785 | 28 | 496 | 8.17 | 9.12E-05 | Nr4a3;Nfat5;Il2ra;Emp2;Igf1;Vsir;Emilin1;Cd276;Cd80;Tnfsf4;Dpp4;Kif26b;Rasgrp1;Ndnf;Fut7;Alox15;Il12rb1;Prkce;Icos;Frmd5;Ets1;Ager;Cd46;Itga3;Mdk;Gp1ba;Cd5;Vegfa | positive regulation of cell adhesion |
| GO:0045765 | 16 | 291 | 4.79 | 1.18E-02 | Emp2;Hmox1;Emilin1;Krit1;Atp2b4;Klf2;Adrb2;Pde3b;Pparg;Fgf1;Ets1;Ghrl;Mdk;Hhip;Vegfa;Reck | regulation of angiogenesis |
| GO:0045066 | 5 | 41 | 0.68 | 4.64E-02 | Vsir;Tnfsf4;Fut7;Cd46;Mdk | regulatory T cell differentiation |
| GO:0044092 | 13 | 260 | 4.28 | 3.98E-02 | Cmklr1;Stk39;Tnfsf4;Ptprf;Adrb2;Gas6;Per2;Akap5;Bicd1; | negative regulation of molecular function |

| | | | | | | |
|---|---|---|---|---|---|---|
| | | | | | Timp3;Ptprh;Zfyve28;Reck | |
| GO:0043542 | 13 | 221 | 3.64 | 1.74E-02 | Emp2;Igf1;Hmox1;Krit1;Atp2b4;Dpp4;Plekhg5;Pparg;Fgf1;Tgfbr3;Ets1;Ager;Vegfa | endothelial cell migration |
| GO:0043524 | 12 | 201 | 3.31 | 2.14E-02 | Nr4a3;Pou4f1;Hmox1;Fyn;En2;Unc5b;Ndnf;Mag;Prkcg;Tox3;Ntrk2;Mdk | negative regulation of neuron apoptotic process |
| GO:0043523 | 15 | 298 | 4.91 | 2.16E-02 | Nr4a3;Pou4f1;Hmox1;Fyn;Ptprf;En2;Unc5b;Egr1;Ndnf;Mag;Prkcg;Tox3;Ager;Ntrk2;Mdk | regulation of neuron apoptotic process |
| GO:0043410 | 22 | 484 | 7.97 | 9.20E-03 | Igf1;Osm;Dok2;Stk39;Lepr;Nod1;Adrb2;Rasgrp1;Mid1;Gas6;Alox15;Fgf1;Ar;Cdon;Prkce;Ager;Ntrk2;Ghrl;Plcb1;Adra2c;Dixdc1;Vegfa | positive regulation of MAPK cascade |
| GO:0042461 | 6 | 61 | 1.00 | 4.39E-02 | Samd11;Cep290;Alms1;Ntrk2;Nphp4;Vegfa | photoreceptor cell development |
| GO:0042104 | 5 | 27 | 0.44 | 1.74E-02 | Il2ra;Igf1;Tnfsf4;Il12rb1;Ager | positive regulation of activated T cell proliferation |
| GO:0042098 | 14 | 252 | 4.15 | 1.74E-02 | Il2ra;Igf1;Vsir;Fyn;Cd276;Cd80;Tnfsf4;Clec2i;Satb1;Rasgrp1;Il12rb1;Btla;Ager;Cd46 | T cell proliferation |
| GO:0042088 | 6 | 49 | 0.81 | 2.30E-02 | Stat4;Tnfsf4;Arid5a;Il12rb1;Il33;Vegfa | T-helper 1 type immune response |
| GO:0042060 | 16 | 373 | 6.14 | 4.39E-02 | Igf1;F5;Hmox1;Slc6a4;Emilin1;Ppl;Ndnf;Gas6;Alox15;Pparg;Fgf1;Prkce;Gp1ba;Adra2c;Plat;Vegfa | wound healing |
| GO:0035811 | 3 | 10 | 0.16 | 4.39E-02 | Nfat5;Adrb2;Adrb1 | negative regulation of urine volume |
| GO:0035722 | 4 | 8 | 0.13 | 3.69E-03 | Stat4;Il12rb2;Il12rb1;Plcb1 | interleukin-12-mediated signaling pathway |
| GO:0034764 | 11 | 202 | 3.33 | 4.64E-02 | Nr4a3;Igf1;Kcnj2;Adrb2;Akap5;Adrb1;Ank2;Ocln;Tesc;Plcb1;Cacnb2 | positive regulation of transmembrane transport |
| GO:0034762 | 20 | 411 | 6.77 | 8.91E-03 | Nr4a3;Igf1;Fyn;Stk39;Kcnj2;Tspan13;Adrb2;Yes1;Per2;Akap5;Prkce;Septin2;Adrb1;Ank2;Ocln;T | regulation of transmembrane transport |

| | | | | | | |
|---|---|---|---|---|---|---|
| | | | | | esc;Plcb1;Cacnb2;Tmem38a;Prrt1 | |
| GO:0032943 | 16 | 361 | 5.95 | 3.63E-02 | Il2ra;Emp2;Igf1;Vsir;Fyn;Cd276;Cd80;Tnfsf4;Clec2i;Satb1;Rasgrp1;Siglecg;Il12rb1;Btla;Ager;Cd46 | mononuclear cell proliferation |
| GO:0032689 | 5 | 38 | 0.63 | 3.86E-02 | Vsir;Cd276;Tnfsf4;Gas6;Il33 | negative regulation of type II interferon production |
| GO:0032680 | 12 | 206 | 3.39 | 2.38E-02 | Igf1;Vsir;Arid5a;Nod1;Rasgrp1;Gas6;Ager;Ghrl;Il33;Oas1c;Tlr5;Tnfrsf8 | regulation of tumor necrosis factor production |
| GO:0032649 | 9 | 122 | 2.01 | 2.50E-02 | Vsir;Cd276;Tnfsf4;Arid5a;Rasgrp1;Gas6;Il12rb2;Il12rb1;Il33 | regulation of type II interferon production |
| GO:0032640 | 12 | 206 | 3.39 | 2.38E-02 | Igf1;Vsir;Arid5a;Nod1;Rasgrp1;Gas6;Ager;Ghrl;Il33;Oas1c;Tlr5;Tnfrsf8 | tumor necrosis factor production |
| GO:0032609 | 9 | 122 | 2.01 | 2.50E-02 | Vsir;Cd276;Tnfsf4;Arid5a;Rasgrp1;Gas6;Il12rb2;Il12rb1;Il33 | type II interferon production |
| GO:0031295 | 5 | 32 | 0.53 | 2.38E-02 | Cd80;Tnfsf4;Dpp4;Icos;Cd5 | T cell costimulation |
| GO:0031294 | 5 | 34 | 0.56 | 2.70E-02 | Cd80;Tnfsf4;Dpp4;Icos;Cd5 | lymphocyte costimulation |
| GO:0030099 | 22 | 467 | 7.69 | 6.81E-03 | Fam20c;Rabgap1l;Pou4f1;Tifab;Pde1b;Scin;Klf2;Clec2i;Pparg;Spib;Tmem178;Csf3;Tgfbr3;Fos;Efna2;Ets1;Tesc;Plcb1;Il33;Gp1ba;Vegfa;Tnfrsf11b | myeloid cell differentiation |
| GO:0022409 | 22 | 322 | 5.30 | 9.12E-05 | Nr4a3;Nfat5;Il2ra;Igf1;Vsir;Emilin1;Cd276;Cd80;Tnfsf4;Dpp4;Kif26b;Rasgrp1;Fut7;Alox15;Il12rb1;Icos;Ets1;Ager;Cd46;Mdk;Gp1ba;Cd5 | positive regulation of cell-cell adhesion |
| GO:0022407 | 25 | 494 | 8.14 | 1.30E-03 | Nr4a3;Nfat5;Il2ra;Igf1;Vsir;Emilin1;Cd276;Cd80;Tnfsf4;Dpp4;Kif26b;Rasgrp1;Fut7;Alox15;Il12rb1;Btla;Icos;Ets1;Ager;Cd46;Mdk;Zdhhc2;Gp1ba;Cd5;Vegfa | regulation of cell-cell adhesion |
| GO:0019932 | 18 | 331 | 5.45 | 6.81E-03 | Cmklr1;Nfat5;Adgrg6;Igf1;Atp2b4;Ndnf;Pde3b;Siglecg;Aka | second-messenger-mediated signaling |

| | | | | | | |
|---|---|---|---|---|---|---|
| | | | | | p5;Tnni3;Alms1;Ank2;Tox3;Ccr6;Ncald;Tmem38a;Tbc1d10c;Vegfa | |
| GO:0019722 | 14 | 227 | 3.74 | 1.18E-02 | Cmklr1;Nfat5;Igf1;Atp2b4;Siglecg;Akap5;Tnni3;Alms1;Ank2;Tox3;Ccr6;Ncald;Tmem38a;Tbc1d10c | calcium-mediated signaling |
| GO:0016525 | 8 | 105 | 1.73 | 3.63E-02 | Emilin1;Krit1;Atp2b4;Klf2;Adrb2;Pde3b;Pparg;Hhip | negative regulation of angiogenesis |
| GO:0010632 | 14 | 233 | 3.84 | 1.20E-02 | Emp2;Igf1;Hmox1;Krit1;Atp2b4;Pparg;Fgf1;Tgfbr3;Prkce;Ets1;Ager;Itga3;Ccr6;Vegfa | regulation of epithelial cell migration |
| GO:0010631 | 16 | 313 | 5.16 | 1.74E-02 | Emp2;Igf1;Hmox1;Krit1;Atp2b4;Dpp4;Plekhg5;Pparg;Fgf1;Tgfbr3;Prkce;Ets1;Ager;Itga3;Ccr6;Vegfa | epithelial cell migration |
| GO:0010594 | 11 | 166 | 2.73 | 1.90E-02 | Emp2;Igf1;Hmox1;Krit1;Atp2b4;Pparg;Fgf1;Tgfbr3;Ets1;Ager;Vegfa | regulation of endothelial cell migration |
| GO:0008217 | 13 | 216 | 3.56 | 1.74E-02 | Emp2;Hmox1;Emilin1;Stk39;Adrb2;Gas6;Pparg;Tnni3;Ar;Adrb1;Col1a2;Ncald;Grip2 | regulation of blood pressure |
| GO:0007623 | 14 | 235 | 3.87 | 1.23E-02 | Igf1;Slc6a4;Lepr;Egr1;Per2;Pparg;Rbm4;Rbm4b;Prkcg;Adrb1;Prokr1;Suv39h2;Ntrk2;Ghrl | circadian rhythm |
| GO:0007613 | 10 | 166 | 2.73 | 4.07E-02 | Igf1;Slc6a4;Egr1;Slc6a1;Adrb1;Ntrk2;Itga3;Plcb1;Mdk;Chst10 | memory |
| GO:0007611 | 19 | 326 | 5.37 | 3.17E-03 | Igf1;Tifab;Slc6a4;Pde1b;Adrb2;Egr1;Slc6a1;Fos;Prkcg;Adrb1;Ager;Ntrk2;Ghrl;Itga3;Plcb1;Mdk;Nptx2;Chst10;Prrt1 | learning or memory |
| GO:0007159 | 21 | 411 | 6.77 | 3.69E-03 | Nr4a3;Nfat5;Il2ra;Igf1;Vsir;Cd276;Cd80;Tnfsf4;Olr1;Dpp4;Rasgrp1;Fut7;Il12rb1;Btla;Icos;Ets1;Ager;Cd46;Mdk;Gp1ba;Cd5 | leukocyte cell-cell adhesion |

| | | | | | | |
|---|---|---|---|---|---|---|
| GO:0006942 | 7 | 83 | 1.37 | 4.07E-02 | Kcnj2;Grcc10;Tnni3;Adrb1;Ank2;Tnnt3;Tmem38a | regulation of striated muscle contraction |
| GO:0006939 | 8 | 112 | 1.85 | 4.64E-02 | Tifab;Atp2b4;Adrb2;Tnni3;Adrb1;Ghrl;Adra2c;Grip2 | smooth muscle contraction |
| GO:0006937 | 10 | 155 | 2.55 | 2.95E-02 | Kcnj2;Adrb2;Grcc10;Tnni3;Adrb1;Ank2;Ghrl;Tnnt3;Adra2c;Tmem38a | regulation of muscle contraction |
| GO:0006936 | 14 | 299 | 4.93 | 4.27E-02 | Tifab;Atp2b4;Kcnj2;Adrb2;Grcc10;Tnni3;Adrb1;Ank2;Ghrl;Tnnt3;Adra2c;Cacnb2;Tmem38a;Grip2 | muscle contraction |
| GO:0006816 | 18 | 414 | 6.82 | 2.50E-02 | Cdh23;Igf1;Fyn;Atp2b4;Tspan13;Adrb2;Gas6;Akap5;Prkce;Adrb1;Cacna2d2;Ank2;Slc24a1;Plcb1;Gp1ba;Cacna2d4;Cacnb2;Tmem38a | calcium ion transport |
| GO:0003012 | 19 | 419 | 6.90 | 1.74E-02 | Nr4a3;Igf1;Tifab;Atp2b4;Kcnj2;Adrb2;Pparg;Grcc10;Tnni3;Ar;Pi16;Adrb1;Ank2;Ghrl;Tnnt3;Adra2c;Cacnb2;Tmem38a;Grip2 | muscle system process |
| GO:0002820 | 7 | 57 | 0.94 | 1.26E-02 | Vsir;Samsn1;Cd80;Tnfsf4;Alox15;Cd46;Il33 | negative regulation of adaptive immune response |
| GO:0002819 | 13 | 231 | 3.81 | 2.09E-02 | Vsir;Samsn1;Cd80;Tnfsf4;Arid5a;Dpp4;Fut7;Alox15;Il12rb1;Ager;Cd46;Ccr6;Il33 | regulation of adaptive immune response |
| GO:0002696 | 17 | 365 | 6.01 | 2.09E-02 | Nr4a3;Il2ra;Igf1;Vsir;Cd276;Cd80;Tnfsf4;Dpp4;Rasgrp1;Gas6;Il12rb1;Icos;Ager;Cd46;Mdk;Il33;Cd5 | positive regulation of leukocyte activation |
| GO:0002685 | 14 | 231 | 3.81 | 1.18E-02 | Cmklr1;Emilin1;Stk39;Tnfsf4;Dpp4;Gas6;Fut7;Ager;Plcb1;Ccr6;Mdk;Il33;Gp1ba;Vegfa | regulation of leukocyte migration |
| GO:0002573 | 15 | 272 | 4.48 | 1.47E-02 | Fam20c;Pou4f1;Pde1b;Clec2i;Pparg;Spib;Tmem178;Csf3;Fos;Efna2;Tesc;Plcb1;Il33;Vegfa;Tnfrsf11b | myeloid leukocyte differentiation |
| GO:0002460 | 15 | 332 | 5.47 | 4.06E-02 | Emp2;Vsir;Cd80;Stat4;Tnfsf4;Arid5a;Dpp4;Fut7;Bach2;Il1 | adaptive immune response based on somatic |

| | | | | | | |
|---|---|---|---|---|---|---|
| | | | | | 2rb1;Ager;Cd46;Ccr6;Il33;Vegfa | recombination of immune receptors built from immunoglobulin superfamily domains |
| GO:0002250 | 20 | 464 | 7.64 | 1.84E-02 | Emp2;Vsir;Fyn;Samsn1;Cd80;Stat4;Tnfsf4;Arid5a;Dpp4;Fut7;Alox15;Bach2;Siglecg;Il12rb1;Btla;Ager;Cd46;Ccr6;Il33;Vegfa | adaptive immune response |
| GO:0001818 | 15 | 295 | 4.86 | 2.08E-02 | Cmklr1;Igf1;Hmox1;Vsir;Cd276;Tnfsf4;Klf2;Gas6;Pparg;Ager;Ghrl;Il1r2;Il33;Oas1c;Wnt11 | negative regulation of cytokine production |
| GO:0001754 | 6 | 59 | 0.97 | 4.02E-02 | Ppp2r3a;Samd11;Cep290;Alms1;Ntrk2;Vegfa | eye photoreceptor cell differentiation |
| GO:0001667 | 18 | 449 | 7.40 | 4.39E-02 | Emp2;Igf1;Hmox1;Krit1;Atp2b4;Dpp4;Lama5;Plekhg5;Pparg;Fgf1;Tgfbr3;Prkce;Ets1;Ager;Itga3;Ccr6;Wnt11;Vegfa | ameboidal-type cell migration |
| **KEGG GeneSet** | **In Data** | **In Pathway** | **Expected** | **FDR** | **Pathway Hits** | **Description** |
| mmu04148 | 11 | 161 | 2.8612323 | 0.03898 | Atp11b;Sirpb1c;Ch25h;Sirpb1b;Gas6;Alox15;Pparg;Siglecg;Megf11;Thbs1;Ager | Efferocytosis |
| mmu04151 | 17 | 364 | 6.468873 | 0.03898 | Il2ra;Igf1;Osm;Ppp2r3a;Lama5;Fgf1;Csf3;Thbs1;Gng4;Efna2;Lpar5;Col1a2;Ntrk2;Itga3;Mtcp1;Vegfa;Itga11 | PI3K-Akt signaling pathway |
| mmu05414 | 8 | 101 | 1.7949346 | 0.03898 | Igf1;Tnni3;Adrb1;Cacna2d2;Itga3;Cacna2d4;Cacnb2;Itga11 | Dilated cardiomyopathy |
| mmu04261 | 10 | 156 | 2.7723742 | 0.03898 | Crem;Atp2b4;Ppp2r3a;Adrb2;Tnni3;Adrb1;Cacna2d2;Plcb1;Cacna2d4;Cacnb2 | Adrenergic signaling in cardiomyocytes |
| mmu04020 | 13 | 253 | 4.4962222 | 0.03898 | Pde1b;Mylk;Atp2b4;Phka1;Adrb2;Fgf1;Prkcg;Adrb1;Ntrk2;Plcb1;Ptger3;Gnal;Vegfa | Calcium signaling pathway |

**Table S4. Significantly active transcription factors (TF) in CAC+HDL induced gene expression in macrophages compared to Ctr-HDL.**

| TF | In Data | In Pathway | Expected | p-value | z-score | TF gene | TF gene Fold Change | TF gene p-value |
|---|---|---|---|---|---|---|---|---|
| CUX1 (p110) | 24 | 1325 | 15.00 | 1.69E-02 | 2.38 | | | |
| REV-ERBalpha | 20 | 1051 | 11.89 | 1.75E-02 | 2.40 | | | |
| GATA-6 | 37 | 2224 | 25.17 | 1.27E-02 | 2.45 | | | |
| SOX4 | 7 | 252 | 2.85 | 2.53E-02 | 2.48 | | | |
| c-Rel (NF-kB subunit) | 7 | 246 | 2.78 | 2.26E-02 | 2.55 | *Rel* | 1.60 | 0.00E+00 |
| Ikaros | 9 | 349 | 3.95 | 1.87E-02 | 2.57 | | | |
| SPI-B | 21 | 1079 | 12.21 | 1.20E-02 | 2.57 | *Spib* | 1.58 | 0.00E+00 |
| HNF1-beta | 5 | 146 | 1.65 | 2.56E-02 | 2.63 | | | |
| PAX5 | 48 | 2957 | 33.47 | 7.44E-03 | 2.64 | | | |
| PAX8 | 6 | 188 | 2.13 | 2.06E-02 | 2.68 | | | |
| PPAR-beta(delta) | 7 | 235 | 2.66 | 1.81E-02 | 2.69 | | | |
| DMRT1 | 5 | 142 | 1.61 | 2.31E-02 | 2.70 | | | |
| C/EBPdelta | 7 | 228 | 2.58 | 1.56E-02 | 2.78 | | | |
| NFYA | 7 | 226 | 2.56 | 1.50E-02 | 2.80 | | | |
| EBF4 | 4 | 96 | 1.09 | 2.38E-02 | 2.82 | | | |
| SNAIL1 | 40 | 2287 | 25.88 | 4.33E-03 | 2.89 | | | |
| IRF5 | 4 | 93 | 1.05 | 2.15E-02 | 2.89 | | | |
| E2F6 | 5 | 129 | 1.46 | 1.59E-02 | 2.95 | | | |
| FLI1 | 7 | 215 | 2.43 | 1.16E-02 | 2.95 | | | |
| CSX (Nkx2.5) | 5 | 127 | 1.44 | 1.50E-02 | 2.99 | | | |
| CLOCK | 7 | 212 | 2.40 | 1.08E-02 | 3.00 | | | |
| MEF2A | 6 | 166 | 1.88 | 1.18E-02 | 3.03 | | | |
| KLF3 | 4 | 88 | 1.00 | 1.79E-02 | 3.03 | | | |
| AP-2A | 13 | 512 | 5.80 | 6.03E-03 | 3.03 | | | |
| ROR-gamma | 6 | 164 | 1.86 | 1.12E-02 | 3.07 | | | |
| KLF15 | 3 | 54 | 0.61 | 2.33E-02 | 3.08 | | | |
| p73 | 8 | 251 | 2.84 | 8.16E-03 | 3.09 | | | |
| Islet-1 | 7 | 205 | 2.32 | 9.12E-03 | 3.10 | | | |
| XBP1 | 8 | 249 | 2.82 | 7.80E-03 | 3.12 | | | |
| NF-AT5 | 3 | 53 | 0.60 | 2.22E-02 | 3.12 | *Nfat5* | 1.67 | 0.00E+00 |
| PROX1 | 5 | 121 | 1.37 | 1.24E-02 | 3.13 | | | |
| NOTCH1 precursor | 4 | 83 | 0.94 | 1.47E-02 | 3.18 | | | |
| E2A | 22 | 1008 | 11.41 | 2.82E-03 | 3.20 | | | |
| RFX2 | 3 | 50 | 0.57 | 1.91E-02 | 3.26 | | | |
| MafA | 3 | 50 | 0.57 | 1.91E-02 | 3.26 | | | |

| | | | | | | | | |
|---|---|---|---|---|---|---|---|---|
| ELF3 | 3 | 50 | 0.57 | 1.91E-02 | 3.26 | | | |
| VDR | 18 | 754 | 8.53 | 2.64E-03 | 3.30 | | | |
| HOXA9 | 3 | 49 | 0.55 | 1.81E-02 | 3.31 | | | |
| EPAS1 | 9 | 279 | 3.16 | 4.79E-03 | 3.32 | | | |
| b-Myb | 6 | 149 | 1.69 | 7.17E-03 | 3.35 | | | |
| FOXF1 | 3 | 48 | 0.54 | 1.71E-02 | 3.35 | | | |
| PEA3 | 5 | 109 | 1.23 | 8.10E-03 | 3.42 | | | |
| E2F7 | 18 | 730 | 8.26 | 1.87E-03 | 3.44 | | | |
| NFIA | 4 | 75 | 0.85 | 1.05E-02 | 3.44 | | | |
| FosB | 4 | 75 | 0.85 | 1.05E-02 | 3.44 | *Fosb* | 2.58 | 0.00E+00 |
| IRF3 | 5 | 105 | 1.19 | 6.95E-03 | 3.52 | | | |
| NXF | 5 | 105 | 1.19 | 6.95E-03 | 3.52 | | | |
| MITF | 8 | 218 | 2.47 | 3.57E-03 | 3.55 | | | |
| MYOD | 24 | 1053 | 11.92 | 1.04E-03 | 3.57 | | | |
| ELF5 | 9 | 259 | 2.93 | 2.95E-03 | 3.58 | | | |
| TFIIB | 5 | 101 | 1.14 | 5.91E-03 | 3.63 | | | |
| FOXM1 | 22 | 922 | 10.43 | 9.35E-04 | 3.65 | | | |
| Lef-1 | 20 | 802 | 9.08 | 9.32E-04 | 3.69 | | | |
| OLIG2 | 8 | 208 | 2.35 | 2.68E-03 | 3.71 | | | |
| SALL4 | 15 | 531 | 6.01 | 1.20E-03 | 3.72 | | | |
| TR-alpha | 6 | 131 | 1.48 | 3.87E-03 | 3.74 | | | |
| HOXA10 | 3 | 41 | 0.46 | 1.12E-02 | 3.75 | | | |
| RFX1 | 3 | 41 | 0.46 | 1.12E-02 | 3.75 | | | |
| PIT1 | 3 | 41 | 0.46 | 1.12E-02 | 3.75 | | | |
| BTEB1 | 5 | 96 | 1.09 | 4.77E-03 | 3.78 | | | |
| CDX2 | 19 | 732 | 8.28 | 7.79E-04 | 3.78 | | | |
| CREM (activators) | 5 | 95 | 1.08 | 4.57E-03 | 3.81 | *Crem* | 1.75 | 0.00E+00 |
| AP-4 | 7 | 162 | 1.83 | 2.57E-03 | 3.85 | | | |
| HHEX (PRH) | 2 | 19 | 0.22 | 1.92E-02 | 3.87 | | | |
| NFAT-90 | 2 | 19 | 0.22 | 1.92E-02 | 3.87 | | | |
| SATB2 | 2 | 19 | 0.22 | 1.92E-02 | 3.87 | | | |
| KLF13 | 2 | 19 | 0.22 | 1.92E-02 | 3.87 | | | |
| MEF2D | 6 | 125 | 1.42 | 3.07E-03 | 3.88 | | | |
| NRSF | 17 | 611 | 6.92 | 6.85E-04 | 3.89 | | | |
| CIC | 8 | 197 | 2.23 | 1.92E-03 | 3.90 | | | |
| HOXB3 | 2 | 18 | 0.20 | 1.73E-02 | 4.00 | | | |
| Oct-2 | 4 | 61 | 0.69 | 5.08E-03 | 4.01 | | | |
| GATA-4 | 12 | 356 | 4.03 | 8.30E-04 | 4.01 | | | |
| CREM (repressors) | 3 | 37 | 0.42 | 8.41E-03 | 4.01 | *Crem* | 1.75 | 0.00E+00 |
| SREBP1 (nuclear) | 8 | 188 | 2.13 | 1.43E-03 | 4.06 | | | |
| NUR77 | 6 | 118 | 1.34 | 2.30E-03 | 4.07 | | | |

| | | | | | | | | |
|---|---|---|---|---|---|---|---|---|
| KLF11 (TIEG2) | 3 | 36 | 0.41 | 7.79E-03 | 4.09 | | | |
| FOXD3 | 4 | 59 | 0.67 | 4.51E-03 | 4.11 | | | |
| RFX6 | 4 | 59 | 0.67 | 4.51E-03 | 4.11 | | | |
| IRF8 | 14 | 438 | 4.96 | 5.33E-04 | 4.11 | | | |
| TBX2 | 70 | 3909 | 44.24 | 7.26E-05 | 4.13 | | | |
| LXR-beta | 5 | 85 | 0.96 | 2.83E-03 | 4.15 | | | |
| IRF6 | 2 | 17 | 0.19 | 1.55E-02 | 4.15 | | | |
| MafF | 2 | 17 | 0.19 | 1.55E-02 | 4.15 | | | |
| HOXA2 | 16 | 527 | 5.96 | 3.86E-04 | 4.16 | | | |
| GLI-3 | 5 | 84 | 0.95 | 2.69E-03 | 4.18 | | | |
| Elk-1 | 11 | 298 | 3.37 | 6.48E-04 | 4.20 | | | |
| NURR1 | 13 | 383 | 4.34 | 4.79E-04 | 4.21 | | | |
| ARNT | 8 | 179 | 2.03 | 1.05E-03 | 4.23 | | | |
| COUP-TFII | 8 | 179 | 2.03 | 1.05E-03 | 4.23 | | | |
| KLF5 | 11 | 294 | 3.33 | 5.80E-04 | 4.25 | | | |
| DLX4 (BP1) | 2 | 16 | 0.18 | 1.38E-02 | 4.30 | | | |
| GLIS3 | 53 | 2682 | 30.35 | 5.45E-05 | 4.30 | | | |
| LYL1 | 33 | 1416 | 16.03 | 8.30E-05 | 4.35 | | | |
| TFII-I | 7 | 139 | 1.57 | 1.07E-03 | 4.36 | | | |
| STAT4 | 6 | 107 | 1.21 | 1.40E-03 | 4.38 | *Stat4* | 1.80 | 0.00E+00 |
| LXR-alpha | 9 | 206 | 2.33 | 6.08E-04 | 4.41 | | | |
| SP7 | 3 | 32 | 0.36 | 5.60E-03 | 4.41 | | | |
| ATF-5 | 2 | 15 | 0.17 | 1.22E-02 | 4.47 | | | |
| OASIS | 2 | 15 | 0.17 | 1.22E-02 | 4.47 | | | |
| KLF8 | 2 | 15 | 0.17 | 1.22E-02 | 4.47 | | | |
| NF-AT1(NFATC2) | 46 | 2159 | 24.43 | 3.03E-05 | 4.53 | | | |
| DSIPI (GILZ) | 4 | 51 | 0.58 | 2.66E-03 | 4.53 | | | |
| hASH1 | 23 | 824 | 9.33 | 7.87E-05 | 4.56 | | | |
| STAT1 | 19 | 623 | 7.05 | 1.07E-04 | 4.57 | | | |
| ATF-4 | 10 | 232 | 2.63 | 3.42E-04 | 4.59 | | | |
| FOXK2 | 7 | 130 | 1.47 | 7.24E-04 | 4.59 | | | |
| TCF7L2 (TCF4) | 108 | 6426 | 72.73 | 7.84E-06 | 4.60 | | | |
| c-Jun | 38 | 1645 | 18.62 | 2.88E-05 | 4.63 | | | |
| TBX20 | 6 | 99 | 1.12 | 9.36E-04 | 4.64 | | | |
| FKHR | 84 | 4655 | 52.68 | 8.59E-06 | 4.66 | | | |
| AF-4 | 8 | 158 | 1.79 | 4.62E-04 | 4.68 | | | |
| MAZ | 5 | 71 | 0.80 | 1.28E-03 | 4.71 | | | |
| EGR3 | 4 | 48 | 0.54 | 2.12E-03 | 4.72 | | | |
| HAND2 | 27 | 1002 | 11.34 | 3.49E-05 | 4.75 | | | |
| ETS1 | 127 | 7753 | 87.74 | 3.11E-06 | 4.77 | *Ets1* | 1.57 | 0.00E+00 |

| | | | | | | | | |
|---|---|---|---|---|---|---|---|---|
| ETS2 | 8 | 154 | 1.74 | 3.90E-04 | 4.78 | | | |
| STAT6 | 20 | 643 | 7.28 | 5.39E-05 | 4.79 | | | |
| OC-2 | 3 | 28 | 0.32 | 3.82E-03 | 4.80 | | | |
| TWIST1 | 14 | 374 | 4.23 | 1.08E-04 | 4.80 | | | |
| SP3 | 29 | 1095 | 12.39 | 2.45E-05 | 4.82 | | | |
| Fra-1 | 11 | 252 | 2.85 | 1.57E-04 | 4.87 | | | |
| FOXP1 | 68 | 3450 | 39.05 | 4.44E-06 | 4.91 | | | |
| E2F1 | 98 | 5517 | 62.44 | 2.34E-06 | 4.93 | | | |
| GRHL2 | 9 | 179 | 2.03 | 2.18E-04 | 4.94 | | | |
| NRF1 | 69 | 3496 | 39.57 | 3.56E-06 | 4.96 | | | |
| Fra-2 | 10 | 211 | 2.39 | 1.60E-04 | 4.97 | | | |
| FXR | 16 | 442 | 5.00 | 5.25E-05 | 4.98 | | | |
| MKL1 | 4 | 44 | 0.50 | 1.53E-03 | 4.99 | | | |
| BACH1 | 7 | 116 | 1.31 | 3.66E-04 | 5.00 | | | |
| MEF2C | 7 | 115 | 1.30 | 3.47E-04 | 5.03 | | | |
| Oct-6 | 42 | 1775 | 20.09 | 6.01E-06 | 5.05 | | | |
| BACH2 | 5 | 64 | 0.72 | 7.98E-04 | 5.06 | *Bach2* | 2.14 | 0.00E+00 |
| HOXB9 | 2 | 12 | 0.14 | 7.82E-03 | 5.09 | | | |
| TITF1 | 13 | 311 | 3.52 | 6.29E-05 | 5.10 | | | |
| Menin | 6 | 86 | 0.97 | 4.44E-04 | 5.13 | | | |
| NPAS2 | 3 | 25 | 0.28 | 2.75E-03 | 5.14 | | | |
| SF1 | 8 | 139 | 1.57 | 1.95E-04 | 5.16 | | | |
| ERG | 18 | 509 | 5.76 | 2.46E-05 | 5.17 | | | |
| PPAR-alpha | 17 | 465 | 5.26 | 2.72E-05 | 5.18 | | | |
| PPAR-gamma | 17 | 464 | 5.25 | 2.65E-05 | 5.19 | *Pparg* | -1.54 | 0.00E+00 |
| ATF-3 | 13 | 304 | 3.44 | 4.99E-05 | 5.21 | | | |
| MLL1 (HRX) | 21 | 630 | 7.13 | 1.28E-05 | 5.27 | *Gas7* | -1.53 | 0.00E+00 |
| ITF2 | 5 | 60 | 0.68 | 5.93E-04 | 5.28 | | | |
| USF1 | 13 | 296 | 3.35 | 3.80E-05 | 5.33 | | | |
| RUNX2 | 17 | 450 | 5.09 | 1.80E-05 | 5.34 | | | |
| CITED2 | 2 | 11 | 0.12 | 6.57E-03 | 5.35 | | | |
| ZNF384 | 2 | 11 | 0.12 | 6.57E-03 | 5.35 | | | |
| ATF-2 | 10 | 191 | 2.16 | 7.04E-05 | 5.38 | | | |
| ESR1 (nuclear) | 106 | 5849 | 66.20 | 2.83E-07 | 5.39 | | | |
| E4BP4 | 6 | 80 | 0.91 | 3.00E-04 | 5.39 | | | |
| c-Myb | 11 | 222 | 2.51 | 5.09E-05 | 5.40 | | | |
| ZNF161 | 3 | 23 | 0.26 | 2.15E-03 | 5.40 | | | |
| SLUG | 7 | 104 | 1.18 | 1.87E-04 | 5.41 | | | |
| FEZF2 | 6 | 79 | 0.89 | 2.81E-04 | 5.44 | | | |
| STAT5A | 14 | 325 | 3.68 | 2.39E-05 | 5.44 | | | |
| GFI-1 | 7 | 103 | 1.17 | 1.76E-04 | 5.44 | | | |
| SOX5 | 5 | 57 | 0.65 | 4.68E-04 | 5.46 | | | |

| | | | | | | | | |
|---|---|---|---|---|---|---|---|---|
| TBP | 11 | 219 | 2.48 | 4.50E-05 | 5.46 | | | |
| CSDA | 3 | 22 | 0.25 | 1.89E-03 | 5.55 | | | |
| NRF2 | 52 | 2222 | 25.15 | 5.89E-07 | 5.56 | | | |
| AHR | 17 | 429 | 4.86 | 9.76E-06 | 5.58 | | | |
| ESR2 (nuclear) | 27 | 864 | 9.78 | 2.50E-06 | 5.61 | | | |
| Nkx6.1 | 6 | 75 | 0.85 | 2.11E-04 | 5.63 | | | |
| ZF5 | 2 | 10 | 0.11 | 5.41E-03 | 5.64 | | | |
| TEF-3 | 29 | 950 | 10.75 | 1.68E-06 | 5.67 | | | |
| JunB | 78 | 3788 | 42.87 | 1.29E-07 | 5.71 | | | |
| EBF | 46 | 1836 | 20.78 | 4.42E-07 | 5.72 | | | |
| TCF7 (TCF1) | 63 | 2830 | 32.03 | 1.88E-07 | 5.74 | | | |
| ZNF143 | 99 | 5180 | 58.62 | 6.55E-08 | 5.74 | | | |
| MEIS1 | 35 | 1229 | 13.91 | 6.88E-07 | 5.79 | | | |
| USF2 | 10 | 173 | 1.96 | 3.05E-05 | 5.79 | | | |
| PR (nuclear) | 18 | 449 | 5.08 | 4.57E-06 | 5.80 | | | |
| JunD | 16 | 366 | 4.14 | 5.26E-06 | 5.89 | | | |
| JDP2 | 4 | 33 | 0.37 | 5.10E-04 | 5.97 | | | |
| ID2 | 2 | 9 | 0.10 | 4.36E-03 | 5.98 | | | |
| CTIP1 | 3 | 19 | 0.22 | 1.22E-03 | 6.04 | | | |
| HOXB13 | 10 | 163 | 1.85 | 1.83E-05 | 6.05 | | | |
| C/EBPbeta | 29 | 893 | 10.11 | 4.88E-07 | 6.05 | | | |
| C/EBPalpha | 23 | 627 | 7.10 | 1.01E-06 | 6.06 | | | |
| c-Fos | 20 | 504 | 5.70 | 1.58E-06 | 6.06 | *Fos* | 1.53 | 0.00E+00 |
| Eomesodermin | 142 | 8057 | 91.19 | 4.73E-09 | 6.09 | | | |
| NF-kB1 (p50) | 20 | 498 | 5.64 | 1.32E-06 | 6.13 | | | |
| DEC1 (Stra13) | 8 | 109 | 1.23 | 3.52E-05 | 6.14 | | | |
| HIF1A | 37 | 1265 | 14.32 | 1.70E-07 | 6.14 | | | |
| HNF3-beta | 38 | 1299 | 14.70 | 1.14E-07 | 6.23 | | | |
| EBF2 | 3 | 18 | 0.20 | 1.04E-03 | 6.23 | | | |
| Elk-3 | 3 | 18 | 0.20 | 1.04E-03 | 6.23 | | | |
| RUNX3 | 20 | 483 | 5.47 | 8.25E-07 | 6.30 | | | |
| ATF-1 | 7 | 83 | 0.94 | 4.47E-05 | 6.30 | | | |
| Bcl-6 | 30 | 900 | 10.19 | 1.78E-07 | 6.33 | | | |
| BMAL1 | 60 | 2455 | 27.78 | 1.52E-08 | 6.37 | | | |
| HB9 | 2 | 8 | 0.09 | 3.42E-03 | 6.38 | | | |
| HLF | 2 | 8 | 0.09 | 3.42E-03 | 6.38 | | | |
| MYF6 | 3 | 17 | 0.19 | 8.70E-04 | 6.44 | | | |
| ATH1 | 119 | 6209 | 70.27 | 1.32E-09 | 6.44 | | | |
| MTB-Zf | 1 | 2 | 0.02 | 2.25E-02 | 6.53 | | | |
| IRX2 | 1 | 2 | 0.02 | 2.25E-02 | 6.53 | | | |

| | | | | | | | | |
|---|---|---|---|---|---|---|---|---|
| CREB5 | 1 | 2 | 0.02 | 2.25E-02 | 6.53 | | | |
| YY2 | 1 | 2 | 0.02 | 2.25E-02 | 6.53 | | | |
| Zipro1 | 1 | 2 | 0.02 | 2.25E-02 | 6.53 | | | |
| ZBTB9 | 1 | 2 | 0.02 | 2.25E-02 | 6.53 | | | |
| FOXI2 | 1 | 2 | 0.02 | 2.25E-02 | 6.53 | | | |
| MEIS2 | 19 | 423 | 4.79 | 4.63E-07 | 6.57 | | | |
| SMAD4 | 20 | 459 | 5.20 | 3.73E-07 | 6.58 | | | |
| HMGB1 | 57 | 2224 | 25.17 | 7.06E-09 | 6.59 | | | |
| STAT5B | 11 | 170 | 1.92 | 4.21E-06 | 6.60 | | | |
| CXXC1 | 48 | 1728 | 19.56 | 1.09E-08 | 6.63 | | | |
| HOX11 | 3 | 16 | 0.18 | 7.22E-04 | 6.66 | | | |
| SATB1 | 8 | 96 | 1.09 | 1.40E-05 | 6.68 | *Satb1* | 1.67 | 0.00E+00 |
| p63 | 28 | 759 | 8.59 | 5.92E-08 | 6.73 | | | |
| SOX9 | 49 | 1752 | 19.83 | 6.07E-09 | 6.76 | | | |
| CHX10 | 37 | 1147 | 12.98 | 1.44E-08 | 6.82 | | | |
| NRIF | 2 | 7 | 0.08 | 2.58E-03 | 6.86 | | | |
| Androgen receptor | 65 | 2587 | 29.28 | 1.15E-09 | 6.90 | *Ar* | 1.92 | 0.00E+00 |
| SRF | 57 | 2133 | 24.14 | 1.59E-09 | 6.94 | | | |
| RARgamma | 7 | 71 | 0.80 | 1.61E-05 | 6.96 | | | |
| EGR2 (Krox20) | 13 | 209 | 2.37 | 8.89E-07 | 6.98 | | | |
| NFIB | 34 | 984 | 11.14 | 1.09E-08 | 6.99 | | | |
| WT1 | 16 | 288 | 3.26 | 2.30E-07 | 7.13 | | | |
| p53 | 65 | 2516 | 28.47 | 3.74E-10 | 7.14 | | | |
| RXRA | 218 | 13172 | 149.10 | 1.41E-12 | 7.17 | | | |
| NF-AT3(NFATC4) | 6 | 51 | 0.58 | 2.37E-05 | 7.18 | | | |
| LHX2 | 102 | 4689 | 53.07 | 3.23E-11 | 7.26 | | | |
| IRF4 | 44 | 1393 | 15.77 | 1.07E-09 | 7.30 | | | |
| SNFT | 4 | 23 | 0.26 | 1.21E-04 | 7.37 | | | |
| BLIMP1 (PRDI-BF1) | 12 | 167 | 1.89 | 5.08E-07 | 7.41 | | | |
| DRIL2 | 2 | 6 | 0.07 | 1.86E-03 | 7.46 | | | |
| YY1 | 114 | 5326 | 60.28 | 3.80E-12 | 7.55 | | | |
| ZNF145 | 9 | 96 | 1.09 | 1.52E-06 | 7.65 | | | |
| NF-AT2(NFATC1) | 17 | 289 | 3.27 | 4.24E-08 | 7.67 | | | |
| TCF7L1 (TCF3) | 79 | 3143 | 35.57 | 1.20E-11 | 7.67 | | | |
| HOXB8 | 3 | 12 | 0.14 | 2.93E-04 | 7.82 | | | |
| TFE3 | 61 | 2095 | 23.71 | 1.26E-11 | 7.94 | | | |
| GATA-3 | 49 | 1476 | 16.70 | 1.97E-11 | 8.12 | | | |
| STAT3 | 50 | 1503 | 17.01 | 1.12E-11 | 8.22 | | | |

| | | | | | | | | |
|---|---|---|---|---|---|---|---|---|
| EGR1 | 30 | 660 | 7.47 | 1.60E-10 | 8.37 | *Egr1* | 3.38 | 0.00E+00 |
| Oct-3/4 | 175 | 9019 | 102.10 | 1.33E-15 | 8.42 | | | |
| HNF4-alpha | 175 | 8988 | 101.70 | 9.39E-16 | 8.47 | | | |
| IRF1 | 56 | 1718 | 19.44 | 1.25E-12 | 8.55 | | | |
| ZNF263 | 165 | 8231 | 93.15 | 8.20E-16 | 8.55 | | | |
| SOX17 | 192 | 10133 | 114.70 | 1.67E-16 | 8.60 | | | |
| ZBTB2 | 72 | 2481 | 28.08 | 1.46E-13 | 8.65 | | | |
| AP-2C | 92 | 3552 | 40.20 | 2.50E-14 | 8.67 | | | |
| SMAD3 | 57 | 1735 | 19.64 | 5.59E-13 | 8.70 | | | |
| LMO2 | 113 | 4728 | 53.51 | 2.85E-15 | 8.79 | | | |
| ETV2 | 6 | 36 | 0.41 | 2.96E-06 | 8.82 | | | |
| Oct-1 | 136 | 6144 | 69.53 | 5.89E-16 | 8.82 | | | |
| DREAM | 5 | 25 | 0.28 | 7.98E-06 | 8.92 | | | |
| C11orf9 | 53 | 1502 | 17.00 | 2.61E-13 | 8.98 | | | |
| BATF | 12 | 120 | 1.36 | 1.32E-08 | 9.20 | | | |
| ZNF362 | 1 | 1 | 0.01 | 1.13E-02 | 9.35 | | | |
| BBX | 1 | 1 | 0.01 | 1.13E-02 | 9.35 | | | |
| ZBED4 | 1 | 1 | 0.01 | 1.13E-02 | 9.35 | | | |
| DMBX1 | 1 | 1 | 0.01 | 1.13E-02 | 9.35 | | | |
| ELL | 1 | 1 | 0.01 | 1.13E-02 | 9.35 | | | |
| c-Maf | 98 | 3636 | 41.15 | 2.08E-16 | 9.41 | | | |
| CTCF | 175 | 8313 | 94.08 | 2.67E-19 | 9.60 | | | |
| DMTF1 | 3 | 8 | 0.09 | 7.72E-05 | 9.73 | | | |
| TCF8 | 84 | 2800 | 31.69 | 1.20E-16 | 9.74 | | | |
| T-bet | 16 | 181 | 2.05 | 3.15E-10 | 9.83 | | | |
| RBP-J kappa (CBF1) | 164 | 7394 | 83.68 | 4.49E-20 | 9.93 | | | |
| SP1 | 183 | 8627 | 97.64 | 7.50E-21 | 10.00 | | | |
| GCR | 204 | 9920 | 112.30 | 1.92E-22 | 10.27 | | | |
| RelA (p65 NF-kB subunit) | 74 | 2187 | 24.75 | 2.07E-17 | 10.28 | | | |
| TEF-1 | 94 | 3076 | 34.81 | 3.16E-19 | 10.56 | *Tead1* | 3.24 | 0.00E+00 |
| PTF1-p48 | 103 | 3506 | 39.68 | 6.22E-20 | 10.65 | | | |
| CREB1 | 175 | 7649 | 86.57 | 2.40E-23 | 10.80 | | | |
| PU.1 | 60 | 1481 | 16.76 | 1.05E-17 | 10.85 | | | |
| KLF17 | 206 | 9674 | 109.50 | 8.68E-25 | 10.89 | | | |
| SOX2 | 155 | 6341 | 71.76 | 5.87E-23 | 10.91 | | | |
| ZFPM1 (FOG) | 3 | 6 | 0.07 | 2.81E-05 | 11.32 | | | |
| c-Myc | 366 | 22619 | 256.00 | 6.32E-38 | 11.55 | | | |
| ASH2 | 95 | 2851 | 32.27 | 5.00E-22 | 11.59 | | | |
| MeCP2 | 5 | 15 | 0.17 | 4.95E-07 | 11.79 | | | |

| | | | | | |
|---|---|---|---|---|---|
| AML1 (RUNX1) | 182 | 7467 | 84.51 | 5.57E-28 | 12.01 |
| FBI-1 (Pokemon) | 275 | 13781 | 156.00 | 8.26E-34 | 12.28 |
| TEF (thyrotroph embryonic factor) | 3 | 5 | 0.06 | 1.42E-05 | 12.44 |
| Esrrg | 166 | 6204 | 70.21 | 1.52E-29 | 12.66 |
| Esrra | 201 | 8291 | 93.83 | 1.15E-31 | 12.72 |
| HNF3-alpha | 181 | 7051 | 79.80 | 1.21E-30 | 12.74 |
| LBP9 | 296 | 15062 | 170.50 | 1.26E-37 | 12.78 |
| KLF4 | 306 | 15403 | 174.30 | 1.59E-41 | 13.37 |
| CRY1 | 273 | 12825 | 145.10 | 5.72E-39 | 13.38 |
| SMAD1 | 178 | 6478 | 73.32 | 8.64E-34 | 13.61 |
| NANOG | 232 | 9793 | 110.80 | 2.68E-37 | 13.62 |
| FOXP3 | 205 | 7961 | 90.10 | 1.86E-36 | 13.84 |
| GATA-1 | 264 | 11679 | 132.20 | 7.76E-42 | 14.10 |
| TAL1 | 235 | 9192 | 104.00 | 6.56E-44 | 15.02 |
| GATA-2 | 245 | 9335 | 105.60 | 5.05E-49 | 15.90 |

**Table 5. Linear equations for HDL-RNA nucleosides**

| **Nucleoside** | **Linear equation** | **LLOQ (nM)** | **ULOQ (nM)** | **$R^2$** |
|---|---|---|---|---|
| **A** | y = 88922x + 53816 | 1.297 | 443.04 | $R^2 = 1$ |
| **G** | y = 43049x + 1497.7 | 0.947 | > 4000 | $R^2 = 1$ |
| **C** | y = 32681x - 4711.9 | 0.027 | > 4000 | $R^2 = 0.9993$ |
| **U** | y = 14414x + 41.862 | 0.685 | 221.52 | $R^2 = 0.9999$ |
| **Ψ** | y = 12609x + 58184 | 0.488 | 15.62 | $R^2 = 0.9915$ |
| **DHU** | y = 11807x + 948.4 | 0.108 | 110.76 | $R^2 = 0.9999$ |
| **Cm** | y = 82796x + 9920.8 | 0.304 | 110.76 | $R^2 = 0.9992$ |
| **$m^3C$** | y = 110738x - 55.699 | 0.013 | 55.37 | $R^2 = 0.9996$ |
| **$m^4C$** | y = 82738x + 1610.1 | 0.039 | 443.04 | $R^2 = 1$ |
| **$m^5C$** | y = 110502x + 5463.7 | 0.027 | 110.76 | $R^2 = 1$ |
| **Um** | y = 15132x - 21.657 | 0.108 | > 4000 | $R^2 = 1$ |
| **$m^3U$** | y = 31757x + 12.183 | 0.108 | > 4000 | $R^2 = 1$ |
| **$m^5U$** | y = 27643x - 74.067 | 0.224 | > 4000 | $R^2 = 0.9999$ |
| **I** | y = 49202x + 4820 | 0.108 | 443.04 | $R^2 = 0.9999$ |
| **Am** | y = 175686x - 134.76 | 0.054 | 110.76 | $R^2 = 1$ |
| **$m^1A$** | y = 343006x + 14915 | 0.021 | 221.52 | $R^2 = 1$ |
| **$m^2A$** | y = 503749x - 348.67 | 0.023 | 110.76 | $R^2 = 0.9993$ |
| **$m^6A$** | y = 179033x + 17545 | 0.022 | 221.52 | $R^2 = 1$ |

| | | | | |
|---|---|---|---|---|
| **Im** | y = 32802x + 4706.9 | 0.432 | 221.52 | $R^2$ = 0.9997 |
| **$m^1I$** | y = 123542x + 2969.5 | 0.216 | 110.76 | $R^2$ = 1 |
| **Gm** | y = 93067x - 1384.3 | 0.136 | > 4000 | $R^2$ = 1 |
| **$m^1G$** | y = 14029x + 975.7 | 0.643 | 110.76 | $R^2$ = 1 |
| **$m^2G$** | y = 124874x + 3562.4 | 0.015 | 443.03 | $R^2$ = 1 |
| **$m^7G$** | y = 136419x + 5723.2 | 0.030 | 110.76 | $R^2$ = 1 |
| **$m^2{}_2G$** | y = 221702x + 10627 | 0.054 | 110.76 | $R^2$ = 1 |
| **$m^2{}_7G$** | y = 280061x + 90.109 | 0.0015 | 55.38 | $R^2$ = 0.9969 |

**Table S6. Nucleoside standards for LC-MS/MS**

| **Nucleoside** | **Code** | **Q1 Mass (Da)** | **Q2 Mass (Da)** | **Time (msec)** | **DP (volts)** | **CE (volts)** |
|---|---|---|---|---|---|---|
| Cytidine | C | 244.2 | 112.2 | 10 | 34 | 15 |
| Uridine | U | 245.1 | 113.1 | 15 | 40 | 14 |
| Guanosine | G | 284.1 | 152 | 10 | 20 | 24 |
| Adenosine | A | 268.1 | 136.1 | 10 | 23 | 30 |
| Inosine | I | 269.2 | 137.1 | 10 | 25 | 12 |
| 3-Methylcytidine | $m^3C$ | 258.1 | 126 | 10 | 35 | 20 |
| 4-Methylcytidine | $m^4C$ | 258.1 | 126 | 10 | 35 | 20 |
| 5-Methylcytidine | $m^5C$ | 258.1 | 126 | 10 | 35 | 20 |
| 3-Methyluridine | $m^3U$ | 259.1 | 127.1 | 15 | 28 | 14 |
| 5-Methyluridine | $m^5U$ | 259.1 | 127.1 | 15 | 28 | 14 |
| 5,6-Dihydrouridine | DHU | 247.2 | 115.2 | 14 | 30 | 14 |
| Pseudouridine | Ψ | 245.1 | 167 | 20 | 30 | 20 |
| 1-Methylguanosine | $m^1G$ | 298.1 | 166 | 10 | 25 | 18 |
| 2-Methylguanosine | $m^2G$ | 298.1 | 166 | 10 | 25 | 18 |
| 7-Methylguanosine | $m^7G$ | 298.1 | 166 | 10 | 25 | 18 |
| 2,2-Dimethylguanosine | $m^2{}_2G$ | 312.1 | 180.1 | 10 | 37 | 19 |
| 2,7 -Dimethylguanosine | $m^2{}_7G$ | 312.1 | 180.1 | 10 | 37 | 19 |
| 1-Methyladenosine | $m^1A$ | 282.1 | 150 | 10 | 43 | 23 |
| 2-Methyladenosine | $m^2A$ | 282.1 | 150 | 10 | 43 | 23 |
| 6-Methyladenosine | $m^6A$ | 282.1 | 150 | 10 | 43 | 23 |
| 8-Methyladenosine | $m^8A$ | 282.1 | 150 | 10 | 43 | 23 |
| 1-Methylinosine | $m^1I$ | 283.1 | 151.1 | 10 | 29 | 12 |
| 2'-O-Methylcytidine | Cm | 258 | 112 | 10 | 30 | 14 |
| 2'-O-Methyladenosine | Am | 282.1 | 136 | 10 | 38 | 22 |
| 2'-O-Methyluridine | Um | 259 | 113 | 20 | 28 | 12 |
| 2-O-Methylguanosine | Gm | 298.1 | 152 | 10 | 20 | 15 |
| 2′-O-Methylinosine | Im | 283.1 | 137.1 | 10 | 25 | 17 |
| [15N]5-2-deoxyadenosine | N15-dA | 257.1 | 141.1 | 10 | 35 | 21 |

**Table S7. Concentrations (nM) of main and modified nucleosides on HDL from Ctr-HDL and CAC$^{+}$HDL subjects by LC-MS/MS (*p<0.05).**

| Nucleoside | Code | Ctr (nM) | CAC (nM) | p-value |
|---|---|---|---|---|
| Cytidine | **C** | 64.81 | 87.77 | 3.35E-04 |
| Uridine | **U** | 36.85 | 52.71 | 1.60E-05 |
| Guanosine | **G** | 81.15 | 105.84 | 1.19E-03 |
| Adenosine | **A** | 83.87 | 105.56 | 2.85E-04 |
| Inosine | **I** | 4.39 | 5.79 | 5.20E-05 |
| 3-Methylcytidine | **$m^3C$** | 0.03 | 0.05 | 3.00E-06 |
| 4-Methylcytidine | **$m^4C$** | 0.02 | 0.02 | 7.30E-01 |
| 5-Methylcytidine | **$m^5C$** | 0.16 | 0.24 | 2.85E-04 |
| 3-Methyluridine | **$m^3U$** | 0.06 | 0.08 | 7.02E-03 |
| 5-Methyluridine | **$m^5U$** | 0.25 | 0.34 | 1.36E-04 |
| 5,6-Dihydrouridine | **DHU** | 0.61 | 0.87 | 1.68E-03 |
| Pseudouridine | **Ψ** | 5.53 | 4.52 | 3.29E-02 |
| 1-Methylguanosine | **$m^1G$** | 1.97 | 2.78 | 7.17E-04 |
| 2-Methylguanosine | **$m^2G$** | 0.13 | 0.20 | 9.52E-03 |
| 7-Methylguanosine | **$m^7G$** | 0.35 | 0.42 | 4.73E-02 |
| 2,2-Dimethylguanosine | **$m^2{}_2G$** | 0.17 | 0.25 | 2.35E-03 |
| 2,7-Dimethylguanosine | **$m^2{}_7G$** | 0.00 | 0.01 | <0.000001 |
| 1-Methyladenosine | **$m^1A$** | 0.25 | 0.36 | 1.37E-03 |
| 2-Methyladenosine | **$m^2A$** | 0.02 | 0.03 | 2.84E-02 |
| 6-Methyladenosine | **$m^6A$** | 0.03 | 0.08 | 1.02E-04 |
| 1-Methylinosine | **$m^1I$** | 0.01 | 0.03 | 4.81E-02 |
| 2'-O-Methylcytidine | **Cm** | 0.33 | 0.49 | 3.68E-03 |
| 2'-O-Methyluridine | **Um** | 0.30 | 0.42 | 2.00E-05 |
| 2'-O-Methylguanosine | **Gm** | 0.55 | 0.78 | 2.43E-04 |
| 2'-O-Methyladenosine | **Am** | 0.90 | 1.37 | 6.18E-04 |

**Table S8. AlkB-sensitive host sRNAs on HDL**

| Class | Parent Transcript | Comparison | Fold Change | p-value | padj |
|---|---|---|---|---|---|
| tDR | tRNA-Trp-CCA-1-1 | CAC_HDL_AlkB_vs_CAC_HDL | 45.43 | 5.17E-57 | 2.64E-55 |
| tDR | tRNA-Arg-TCG-1-1 | CAC_HDL_AlkB_vs_CAC_HDL | 43.99 | 1.65E-50 | 4.21E-49 |
| tDR | tRNA-Pro-CGG-1-1 | CAC_HDL_AlkB_vs_CAC_HDL | 20.61 | 6.54E-47 | 1.11E-45 |
| tDR | tRNA-Pro-AGG-1-1 | CAC_HDL_AlkB_vs_CAC_HDL | 15.84 | 2.04E-45 | 2.59E-44 |
| tDR | tRNA-Arg-ACG-1-1 | CAC_HDL_AlkB_vs_CAC_HDL | 55.02 | 8.89E-44 | 9.07E-43 |

| | | | | | |
|---|---|---|---|---|---|
| tDR | tRNA-Arg-CCG-1-1 | CAC_HDL_AlkB_vs_CAC_HDL | 53.79 | 5.83E-35 | 4.66E-34 |
| tDR | tRNA-Lys-TTT-10-1 | CAC_HDL_AlkB_vs_CAC_HDL | 16.53 | 6.40E-35 | 4.66E-34 |
| tDR | tRNA-Asn-GTT-10-1 | CAC_HDL_AlkB_vs_CAC_HDL | 33.92 | 1.42E-34 | 8.06E-34 |
| tDR | tRNA-Leu-CAG-1-1 | CAC_HDL_AlkB_vs_CAC_HDL | 23.92 | 3.89E-34 | 1.98E-33 |
| tDR | tRNA-Pro-TGG-1-1 | CAC_HDL_AlkB_vs_CAC_HDL | 9.61 | 1.58E-33 | 7.34E-33 |
| tDR | tRNA-Leu-CAA-1-1 | CAC_HDL_AlkB_vs_CAC_HDL | 24.92 | 1.18E-31 | 5.02E-31 |
| tDR | tRNA-Thr-TGT-1-1 | CAC_HDL_AlkB_vs_CAC_HDL | 21.28 | 2.60E-30 | 9.48E-30 |
| tDR | tRNA-Arg-CCT-1-1 | CAC_HDL_AlkB_vs_CAC_HDL | 34.76 | 4.26E-30 | 1.45E-29 |
| tDR | tRNA-Met-CAT-1-1 | CAC_HDL_AlkB_vs_CAC_HDL | 41.52 | 1.59E-27 | 5.07E-27 |
| tDR | tRNA-Lys-CTT-10-1 | CAC_HDL_AlkB_vs_CAC_HDL | 9.34 | 2.23E-27 | 6.69E-27 |
| tDR | tRNA-Gln-CTG-10-1 | CAC_HDL_AlkB_vs_CAC_HDL | 15.35 | 3.10E-26 | 8.77E-26 |
| tDR | tRNA-Ser-GCT-1-1 | CAC_HDL_AlkB_vs_CAC_HDL | 16.84 | 3.22E-25 | 8.64E-25 |
| tDR | tRNA-iMet-CAT-1-1 | CAC_HDL_AlkB_vs_CAC_HDL | 27.53 | 1.90E-24 | 4.84E-24 |
| tDR | tRNA-Ile-AAT-10-1 | CAC_HDL_AlkB_vs_CAC_HDL | 19.08 | 2.52E-23 | 5.83E-23 |
| tDR | tRNA-Phe-GAA-10-1 | CAC_HDL_AlkB_vs_CAC_HDL | 35.67 | 1.25E-22 | 2.76E-22 |
| tDR | tRNA-Leu-AAG-1-1 | CAC_HDL_AlkB_vs_CAC_HDL | 9.64 | 4.12E-22 | 8.75E-22 |
| tDR | tRNA-Thr-AGT-1-1 | CAC_HDL_AlkB_vs_CAC_HDL | 19.62 | 4.98E-22 | 1.01E-21 |
| tDR | tRNA-Arg-TCT-1-1 | CAC_HDL_AlkB_vs_CAC_HDL | 21.60 | 1.13E-21 | 2.23E-21 |
| tDR | tRNA-Ser-AGA-1-1 | CAC_HDL_AlkB_vs_CAC_HDL | 14.72 | 3.79E-18 | 6.91E-18 |
| tDR | tRNA-His-GTG-1-1 | CAC_HDL_AlkB_vs_CAC_HDL | 18.91 | 3.07E-17 | 5.41E-17 |
| tDR | tRNA-Cys-GCA-10-1 | CAC_HDL_AlkB_vs_CAC_HDL | 8.13 | 4.49E-17 | 7.63E-17 |
| tDR | tRNA-Val-CAC-10-1 | CAC_HDL_AlkB_vs_CAC_HDL | 8.02 | 3.33E-16 | 5.48E-16 |
| tDR | tRNA-Thr-CGT-1-1 | CAC_HDL_AlkB_vs_CAC_HDL | 14.25 | 8.07E-16 | 1.29E-15 |
| tDR | tRNA-Leu-TAG-1-1 | CAC_HDL_AlkB_vs_CAC_HDL | 6.92 | 2.38E-15 | 3.68E-15 |
| tDR | tRNA-Val-AAC-1-1 | CAC_HDL_AlkB_vs_CAC_HDL | 8.14 | 9.89E-13 | 1.48E-12 |
| tDR | tRNA-Ser-CGA-1-1 | CAC_HDL_AlkB_vs_CAC_HDL | 33.00 | 3.85E-12 | 5.61E-12 |
| tDR | tRNA-Ile-TAT-1-1 | CAC_HDL_AlkB_vs_CAC_HDL | 23.03 | 2.06E-08 | 2.91E-08 |
| tDR | tRNA-Ala-AGC-10-1 | CAC_HDL_AlkB_vs_CAC_HDL | 2.53 | 6.04E-05 | 8.32E-05 |

| | | | | | |
|---|---|---|---|---|---|
| tDR | tRNA-Ala-CGC-1-1 | CAC_HDL_AlkB_vs_CAC_HDL | 1.86 | 1.13E-03 | 1.51E-03 |
| tDR | tRNA-Gly-CCC-1-1 | CAC_HDL_AlkB_vs_CAC_HDL | 1.79 | 2.02E-03 | 2.64E-03 |
| tDR | tRNA-Ala-TGC-1-1 | CAC_HDL_AlkB_vs_CAC_HDL | 1.64 | 4.98E-03 | 6.19E-03 |
| tDR | tRNA-SeC-TCA-1-1 | CAC_HDL_AlkB_vs_CAC_HDL | 2.76 | 8.92E-03 | 1.08E-02 |
| tDR | tRNA-Cys-ACA-1-1 | CAC_HDL_AlkB_vs_CAC_HDL | 0.35 | 2.20E-02 | 2.60E-02 |
| tDR | tRNA-Glu-TTC-10-1 | CAC_HDL_AlkB_vs_CAC_HDL | 1.53 | 4.28E-02 | 4.96E-02 |
| miRNA | hsa-miR-1321 | CAC_HDL_AlkB_vs_CAC_HDL | 6.28 | 1.69E-03 | 1.09E-01 |
| miRNA | hsa-miR-320d | CAC_HDL_AlkB_vs_CAC_HDL | 0.13 | 2.09E-03 | 1.09E-01 |
| miRNA | hsa-miR-6869-5p | CAC_HDL_AlkB_vs_CAC_HDL | 0.04 | 6.61E-03 | 1.91E-01 |
| miRNA | hsa-let-7i-5p | CAC_HDL_AlkB_vs_CAC_HDL | 0.38 | 7.35E-03 | 1.91E-01 |
| miRNA | hsa-let-7g-5p | CAC_HDL_AlkB_vs_CAC_HDL | 0.48 | 1.12E-02 | 1.96E-01 |
| miRNA | hsa-miR-185-5p | CAC_HDL_AlkB_vs_CAC_HDL | 0.08 | 1.13E-02 | 1.96E-01 |
| miRNA | hsa-let-7d-5p | CAC_HDL_AlkB_vs_CAC_HDL | 0.43 | 1.91E-02 | 2.84E-01 |
| miRNA | hsa-miR-21-5p | CAC_HDL_AlkB_vs_CAC_HDL | 0.44 | 2.33E-02 | 3.01E-01 |
| miRNA | hsa-miR-9-5p | CAC_HDL_AlkB_vs_CAC_HDL | 5.07 | 2.82E-02 | 3.01E-01 |
| miRNA | hsa-miR-30d-5p | CAC_HDL_AlkB_vs_CAC_HDL | 0.45 | 2.99E-02 | 3.01E-01 |
| miRNA | hsa-miR-361-5p | CAC_HDL_AlkB_vs_CAC_HDL | 0.14 | 3.18E-02 | 3.01E-01 |
| miRNA | hsa-miR-425-5p | CAC_HDL_AlkB_vs_CAC_HDL | 0.20 | 4.38E-02 | 3.61E-01 |
| miRNA | hsa-miR-191-5p | CAC_HDL_AlkB_vs_CAC_HDL | 0.49 | 4.51E-02 | 3.61E-01 |
| yDR | Y_RNA:ENSG00000202536.1 | CAC_HDL_AlkB_vs_CAC_HDL | 0.21 | 3.07E-04 | 3.13E-02 |
| yDR | Y_RNA:ENSG00000201239.1 | CAC_HDL_AlkB_vs_CAC_HDL | 0.33 | 7.16E-04 | 3.13E-02 |
| yDR | Y_RNA:ENSG00000201498.1 | CAC_HDL_AlkB_vs_CAC_HDL | 0.24 | 8.38E-04 | 3.13E-02 |
| yDR | Y_RNA:ENSG00000201548.1 | CAC_HDL_AlkB_vs_CAC_HDL | 0.26 | 9.43E-04 | 3.13E-02 |
| yDR | Y_RNA:ENSG00000201228.1 | CAC_HDL_AlkB_vs_CAC_HDL | 0.34 | 1.10E-03 | 3.13E-02 |
| yDR | Y_RNA:ENSG00000199635.1 | CAC_HDL_AlkB_vs_CAC_HDL | 0.26 | 1.15E-03 | 3.13E-02 |
| yDR | Y_RNA:ENSG00000199515.1 | CAC_HDL_AlkB_vs_CAC_HDL | 0.35 | 1.94E-03 | 4.53E-02 |
| yDR | RNY4P23:ENSG00000201377.1 | CAC_HDL_AlkB_vs_CAC_HDL | 0.31 | 2.22E-03 | 4.53E-02 |
| yDR | Y_RNA:ENSG00000200141.1 | CAC_HDL_AlkB_vs_CAC_HDL | 0.43 | 3.34E-03 | 6.04E-02 |

| | | | | | |
|---|---|---|---|---|---|
| yDR | Y_RNA:ENSG00000200201.1 | CAC_HDL_AlkB_vs_CAC_HDL | 0.37 | 3.81E-03 | 6.22E-02 |
| yDR | Y_RNA:ENSG00000222493.1 | CAC_HDL_AlkB_vs_CAC_HDL | 0.44 | 9.57E-03 | 1.36E-01 |
| yDR | RNY4P31:ENSG00000253018.1 | CAC_HDL_AlkB_vs_CAC_HDL | 0.42 | 1.00E-02 | 1.36E-01 |
| yDR | Y_RNA:ENSG00000222601.1 | CAC_HDL_AlkB_vs_CAC_HDL | 0.40 | 1.16E-02 | 1.45E-01 |
| yDR | RNY4P15:ENSG00000202012.1 | CAC_HDL_AlkB_vs_CAC_HDL | 0.47 | 1.71E-02 | 1.87E-01 |
| yDR | Y_RNA:ENSG00000201644.1 | CAC_HDL_AlkB_vs_CAC_HDL | 0.35 | 1.81E-02 | 1.87E-01 |
| yDR | Y_RNA:ENSG00000199584.1 | CAC_HDL_AlkB_vs_CAC_HDL | 0.38 | 1.83E-02 | 1.87E-01 |
| yDR | Y_RNA:ENSG00000201938.1 | CAC_HDL_AlkB_vs_CAC_HDL | 0.46 | 2.07E-02 | 1.92E-01 |
| yDR | Y_RNA:ENSG00000252759.1 | CAC_HDL_AlkB_vs_CAC_HDL | 0.45 | 2.12E-02 | 1.92E-01 |
| yDR | Y_RNA:ENSG00000199357.1 | CAC_HDL_AlkB_vs_CAC_HDL | 0.39 | 2.61E-02 | 2.07E-01 |
| yDR | Y_RNA:ENSG00000202357.1 | CAC_HDL_AlkB_vs_CAC_HDL | 0.36 | 2.64E-02 | 2.07E-01 |
| yDR | Y_RNA:ENSG00000239180.1 | CAC_HDL_AlkB_vs_CAC_HDL | 0.50 | 2.79E-02 | 2.07E-01 |
| yDR | Y_RNA:ENSG00000200309.1 | CAC_HDL_AlkB_vs_CAC_HDL | 0.46 | 3.09E-02 | 2.07E-01 |
| yDR | RNY4P16:ENSG00000201638.1 | CAC_HDL_AlkB_vs_CAC_HDL | 0.46 | 3.11E-02 | 2.07E-01 |
| yDR | RNY4P27:ENSG00000200211.1 | CAC_HDL_AlkB_vs_CAC_HDL | 0.51 | 3.11E-02 | 2.07E-01 |
| yDR | Y_RNA:ENSG00000252652.1 | CAC_HDL_AlkB_vs_CAC_HDL | 0.47 | 3.17E-02 | 2.07E-01 |
| yDR | Y_RNA:ENSG00000252522.1 | CAC_HDL_AlkB_vs_CAC_HDL | 0.43 | 3.39E-02 | 2.08E-01 |
| yDR | Y_RNA:ENSG00000200615.1 | CAC_HDL_AlkB_vs_CAC_HDL | 0.56 | 3.54E-02 | 2.08E-01 |
| yDR | Y_RNA:ENSG00000201451.1 | CAC_HDL_AlkB_vs_CAC_HDL | 0.45 | 3.69E-02 | 2.08E-01 |
| yDR | Y_RNA:ENSG00000200121.1 | CAC_HDL_AlkB_vs_CAC_HDL | 0.43 | 3.74E-02 | 2.08E-01 |
| yDR | Y_RNA:ENSG00000200090.1 | CAC_HDL_AlkB_vs_CAC_HDL | 0.30 | 3.83E-02 | 2.08E-01 |
| yDR | RNY4P8:ENSG00000200735.1 | CAC_HDL_AlkB_vs_CAC_HDL | 0.59 | 4.51E-02 | 2.37E-01 |
| tDR | tRNA-Arg-ACG-1-1 | Cntl_HDL_AlkB_vs_Cntl_HDL | 59.93 | 7.97E-44 | 4.23E-42 |
| tDR | tRNA-Arg-CCG-1-1 | Cntl_HDL_AlkB_vs_Cntl_HDL | 77.16 | 2.51E-43 | 6.65E-42 |
| tDR | tRNA-Arg-TCG-1-1 | Cntl_HDL_AlkB_vs_Cntl_HDL | 49.83 | 7.17E-42 | 1.27E-40 |
| tDR | tRNA-Trp-CCA-1-1 | Cntl_HDL_AlkB_vs_Cntl_HDL | 23.70 | 2.11E-39 | 2.79E-38 |
| tDR | tRNA-Ser-GCT-1-1 | Cntl_HDL_AlkB_vs_Cntl_HDL | 14.30 | 3.45E-35 | 3.22E-34 |
| tDR | tRNA-Leu-CAG-1-1 | Cntl_HDL_AlkB_vs_Cntl_HDL | 23.64 | 3.64E-35 | 3.22E-34 |

| | | | | | |
|---|---|---|---|---|---|
| tDR | tRNA-Leu-CAA-1-1 | Cntl_HDL_AlkB_vs_Cntl_HDL | 23.79 | 1.05E-34 | 7.97E-34 |
| tDR | tRNA-iMet-CAT-1-1 | Cntl_HDL_AlkB_vs_Cntl_HDL | 21.03 | 1.22E-34 | 8.09E-34 |
| tDR | tRNA-Phe-GAA-10-1 | Cntl_HDL_AlkB_vs_Cntl_HDL | 18.57 | 2.39E-33 | 1.41E-32 |
| tDR | tRNA-Lys-TTT-10-1 | Cntl_HDL_AlkB_vs_Cntl_HDL | 12.58 | 8.55E-30 | 3.78E-29 |
| tDR | tRNA-Asn-GTT-10-1 | Cntl_HDL_AlkB_vs_Cntl_HDL | 18.32 | 1.05E-28 | 4.08E-28 |
| tDR | tRNA-Met-CAT-1-1 | Cntl_HDL_AlkB_vs_Cntl_HDL | 25.01 | 1.08E-28 | 4.08E-28 |
| tDR | tRNA-Gln-CTG-10-1 | Cntl_HDL_AlkB_vs_Cntl_HDL | 9.72 | 2.20E-27 | 7.79E-27 |
| tDR | tRNA-Arg-TCT-1-1 | Cntl_HDL_AlkB_vs_Cntl_HDL | 19.19 | 2.82E-27 | 9.36E-27 |
| tDR | tRNA-Thr-TGT-1-1 | Cntl_HDL_AlkB_vs_Cntl_HDL | 25.09 | 4.69E-26 | 1.46E-25 |
| tDR | tRNA-Ile-AAT-10-1 | Cntl_HDL_AlkB_vs_Cntl_HDL | 15.77 | 1.40E-25 | 4.11E-25 |
| tDR | tRNA-Arg-CCT-1-1 | Cntl_HDL_AlkB_vs_Cntl_HDL | 30.44 | 1.71E-25 | 4.77E-25 |
| tDR | tRNA-Pro-AGG-1-1 | Cntl_HDL_AlkB_vs_Cntl_HDL | 14.12 | 3.64E-25 | 9.66E-25 |
| tDR | tRNA-Thr-CGT-1-1 | Cntl_HDL_AlkB_vs_Cntl_HDL | 15.21 | 8.97E-23 | 2.26E-22 |
| tDR | tRNA-Pro-CGG-1-1 | Cntl_HDL_AlkB_vs_Cntl_HDL | 14.76 | 4.95E-22 | 1.19E-21 |
| tDR | tRNA-His-GTG-1-1 | Cntl_HDL_AlkB_vs_Cntl_HDL | 18.08 | 3.52E-20 | 7.77E-20 |
| tDR | tRNA-Ser-AGA-1-1 | Cntl_HDL_AlkB_vs_Cntl_HDL | 8.88 | 1.52E-19 | 3.23E-19 |
| tDR | tRNA-Pro-TGG-1-1 | Cntl_HDL_AlkB_vs_Cntl_HDL | 7.36 | 6.19E-19 | 1.26E-18 |
| tDR | tRNA-Cys-GCA-10-1 | Cntl_HDL_AlkB_vs_Cntl_HDL | 4.18 | 1.88E-17 | 3.68E-17 |
| tDR | tRNA-Ile-TAT-1-1 | Cntl_HDL_AlkB_vs_Cntl_HDL | 42.28 | 8.67E-17 | 1.64E-16 |
| tDR | tRNA-Val-CAC-10-1 | Cntl_HDL_AlkB_vs_Cntl_HDL | 6.51 | 9.12E-16 | 1.67E-15 |
| tDR | tRNA-Leu-AAG-1-1 | Cntl_HDL_AlkB_vs_Cntl_HDL | 7.08 | 3.07E-15 | 5.43E-15 |
| tDR | tRNA-Thr-AGT-1-1 | Cntl_HDL_AlkB_vs_Cntl_HDL | 14.12 | 4.07E-15 | 6.95E-15 |
| tDR | tRNA-Lys-CTT-10-1 | Cntl_HDL_AlkB_vs_Cntl_HDL | 5.25 | 3.80E-13 | 6.30E-13 |
| tDR | tRNA-Ala-AGC-10-1 | Cntl_HDL_AlkB_vs_Cntl_HDL | 3.43 | 9.05E-12 | 1.45E-11 |
| tDR | tRNA-Leu-TAG-1-1 | Cntl_HDL_AlkB_vs_Cntl_HDL | 4.35 | 2.06E-08 | 3.13E-08 |
| tDR | tRNA-Ser-CGA-1-1 | Cntl_HDL_AlkB_vs_Cntl_HDL | 9.90 | 1.03E-07 | 1.52E-07 |
| tDR | tRNA-Val-AAC-1-1 | Cntl_HDL_AlkB_vs_Cntl_HDL | 3.86 | 3.20E-07 | 4.58E-07 |
| tDR | tRNA-Ala-CGC-1-1 | Cntl_HDL_AlkB_vs_Cntl_HDL | 2.39 | 8.90E-06 | 1.24E-05 |

| | | | | | |
|---|---|---|---|---|---|
| tDR | tRNA-Ala-TGC-1-1 | Cntl_HDL_AlkB_vs_Cntl_HDL | 2.16 | 5.38E-05 | 7.31E-05 |
| tDR | tRNA-SeC-TCA-1-1 | Cntl_HDL_AlkB_vs_Cntl_HDL | 3.19 | 1.12E-04 | 1.48E-04 |
| tDR | tRNA-Gly-CCC-1-1 | Cntl_HDL_AlkB_vs_Cntl_HDL | 1.82 | 2.45E-03 | 3.17E-03 |
| tDR | tRNA-Gly-TCC-1-1 | Cntl_HDL_AlkB_vs_Cntl_HDL | 1.67 | 1.62E-02 | 1.99E-02 |
| tDR | tRNA-Sup-TTA-1-1 | Cntl_HDL_AlkB_vs_Cntl_HDL | 2.14 | 1.87E-02 | 2.25E-02 |
| miRNA | hsa-miR-184 | Cntl_HDL_AlkB_vs_Cntl_HDL | 2.01 | 3.11E-02 | 9.63E-01 |
| yDR | Y_RNA:ENSG00000199635.1 | Cntl_HDL_AlkB_vs_Cntl_HDL | 4.54 | 1.99E-03 | 3.50E-01 |
| yDR | Y_RNA:ENSG00000200769.1 | Cntl_HDL_AlkB_vs_Cntl_HDL | 0.35 | 1.81E-02 | 9.45E-01 |
| yDR | Y_RNA:ENSG00000201749.1 | Cntl_HDL_AlkB_vs_Cntl_HDL | 0.34 | 2.13E-02 | 9.45E-01 |
| yDR | Y_RNA:ENSG00000199801.1 | Cntl_HDL_AlkB_vs_Cntl_HDL | 0.31 | 2.15E-02 | 9.45E-01 |
| yDR | RNY4P28:ENSG00000202151.1 | Cntl_HDL_AlkB_vs_Cntl_HDL | 0.49 | 4.14E-02 | 9.89E-01 |

**Table S9. Significant differentially abundant HDL-sRNAs in CAC+HDL at the parent level.**

| Gene | Fold Change | p-value |
|---|---|---|
| tRNA-Arg-CCG-1-1 | 1.89 | 1.90E-02 |
| tRNA-Cys-GCA-10-1 | 1.85 | 1.48E-02 |
| tRNA-Arg-TCG-1-1 | 1.81 | 2.28E-02 |
| tRNA-Arg-ACG-1-1 | 1.79 | 2.70E-02 |
| tRNA-Trp-CCA-1-1 | 1.75 | 1.86E-02 |
| tRNA-Asn-GTT-10-1 | 1.67 | 3.24E-02 |
| tRNA-Thr-TGT-1-1 | 1.67 | 4.08E-02 |
| tRNA-Met-CAT-1-1 | 1.66 | 4.48E-02 |
| tRNA-Ser-GCT-1-1 | 1.63 | 3.44E-02 |
| tRNA-Glu-TTC-10-1 | 1.58 | 2.09E-02 |
| tRNA-Asp-GTC-1-1 | 1.57 | 3.93E-02 |
| tRNA-Gln-CTG-10-1 | 1.50 | 4.46E-02 |

**Table S10. Significant differentially abundant tDRs (read level) on CAC$^{+}$HDL**

| tDR | Length | Comparison | Fold Change | p-value |
|---|---|---|---|---|
| GUUAGUACUCUGCGUUGUGGCCGCAGCAACCUCGGUUCGAAUCCGAGUCACGGCAC | 56 | CAC_HDL_AlkB_vs_Cntl_HDL_AlkB | 13.09 | 2.76E-04 |
| UUGUGGGUUCGAGUCCCAUCUUGGUCG | 27 | CAC_HDL_AlkB_vs_Cntl_HDL_AlkB | 11.37 | 1.97E-07 |

| | | | | |
|---|---|---|---|---|
| AUUCUAGGUUCGACUCCUGGCUGGCUCG | 28 | CAC_HDL_AlkB_vs_Cntl_HDL_AlkB | 9.38 | 3.52E-03 |
| AUCAGAAGAUUCUAGGUUCGACUCCUGGCUGGCUCG | 36 | CAC_HDL_AlkB_vs_Cntl_HDL_AlkB | 8.24 | 1.23E-02 |
| CCGGAGCUGGGGAUUGUGGGUUCGAGUCCCAUCUGGGUCG | 40 | CAC_HDL_AlkB_vs_Cntl_HDL_AlkB | 8.18 | 1.24E-02 |
| CAGGUUCGACUCCUGGCUGUCUCG | 24 | CAC_HDL_AlkB_vs_Cntl_HDL_AlkB | 7.92 | 4.52E-06 |
| GGUUCGACUCCUGGCUGGCUCG | 22 | CAC_HDL_AlkB_vs_Cntl_HDL_AlkB | 7.92 | 4.66E-03 |
| UCGCUGGUUCGAUUCCGGCUCGAAGGAC | 28 | CAC_HDL_AlkB_vs_Cntl_HDL_AlkB | 7.57 | 2.00E-02 |
| GGGGUUCGAUUCCCCGACUGGGAG | 24 | CAC_HDL_AlkB_vs_Cntl_HDL_AlkB | 7.02 | 1.17E-04 |
| UCUGACUGCGGAUCAGAAGAUUCUAGGUUCGACUCCUGGCUGGCUCG | 47 | CAC_HDL_AlkB_vs_Cntl_HDL_AlkB | 6.85 | 1.46E-02 |
| GAAGAUUGCAGGUUCGAGUCCUGCCGCGGUCG | 32 | CAC_HDL_AlkB_vs_Cntl_HDL_AlkB | 6.18 | 4.41E-02 |
| UCCCUGGUGGUCUAGUGGCUAGGAUUCGGCGC | 32 | CAC_HDL_AlkB_vs_Cntl_HDL_AlkB | 5.69 | 5.90E-03 |
| UGCUGGGCCCAUAACCCAGAGGUCGAUGGAUCGAAACCAUCCUCUGCUA | 49 | CAC_HDL_AlkB_vs_Cntl_HDL_AlkB | 5.55 | 2.59E-02 |
| UUUACACGCAGAAGGUCCUGGGUUCGAGCCCCAGUGGAACCA | 42 | CAC_HDL_AlkB_vs_Cntl_HDL_AlkB | 5.50 | 4.52E-02 |
| UGGUGGUUCGAGCCCACCCAGGGAC | 25 | CAC_HDL_AlkB_vs_Cntl_HDL_AlkB | 5.44 | 2.61E-02 |
| CUCCCCUGGAGGCGUGGGUUCGAAUCCCACUUCUGACA | 38 | CAC_HDL_AlkB_vs_Cntl_HDL_AlkB | 5.17 | 1.89E-02 |
| AGAGCAUGAGGCUCUUAAUCUCAGGGUCGUGGGUUCGAGCCCCACGUUGGGCG | 53 | CAC_HDL_AlkB_vs_Cntl_HDL_AlkB | 4.99 | 6.92E-05 |
| GGUUAGGAUUCGGCGCUCUCACCGCCGCGGCCCGGGUUCGAUUCCCGGUCAGGGAA | 56 | CAC_HDL_AlkB_vs_Cntl_HDL_AlkB | 4.94 | 1.46E-02 |
| CAGGUUCGACUCCUGGCUGGCU | 22 | CAC_HDL_AlkB_vs_Cntl_HDL_AlkB | 4.94 | 1.28E-02 |
| CCUCGGUUCGAAUCCGAGUCACGGCAC | 27 | CAC_HDL_AlkB_vs_Cntl_HDL_AlkB | 4.90 | 4.70E-02 |
| UUGGGGGUUCGAGUCCCUUCGUGGUCG | 27 | CAC_HDL_AlkB_vs_Cntl_HDL_AlkB | 4.88 | 6.03E-06 |
| AUCACGUCUGCUUUACACGCAGAAGGUCCUGGGUUCGAGCCCCAGUGGAACCA | 53 | CAC_HDL_AlkB_vs_Cntl_HDL_AlkB | 4.85 | 2.93E-02 |
| AGAGCAUGAGUCUCUUAAUCUCAGGGUCGUGGGUUCGAGCCCCACGUUGGGCG | 53 | CAC_HDL_AlkB_vs_Cntl_HDL_AlkB | 4.75 | 1.28E-04 |
| AGGUUCGACUCCUGGCUGGCUCGCC | 25 | CAC_HDL_AlkB_vs_Cntl_HDL_AlkB | 4.63 | 1.68E-02 |
| GUCGCUGGUUCGAUUCCGGCUCGAAUGA | 28 | CAC_HDL_AlkB_vs_Cntl_HDL_AlkB | 4.61 | 6.23E-05 |
| GGGGGUUCGAGUCCCUUCGUGGUCG | 25 | CAC_HDL_AlkB_vs_Cntl_HDL_AlkB | 4.41 | 8.24E-05 |
| AUCCAGCGAUCCGAGUUCAAAUCUCGGUGGAACCU | 35 | CAC_HDL_AlkB_vs_Cntl_HDL_AlkB | 4.38 | 3.92E-02 |

| | | | | |
|---|---|---|---|---|
| UAGCGGUUAGGAUUCCUGGUUUUCACCCAGGUGGCCCGGGUUCGACUCCCGGUAUGGGAA | 60 | CAC_HDL_AlkB_vs_Cntl_HDL_AlkB | 4.38 | 4.32E-04 |
| ACCGGGGUUCGAUUCCCCGACUGGGAG | 27 | CAC_HDL_AlkB_vs_Cntl_HDL_AlkB | 4.27 | 6.15E-05 |
| CCAGGGUUCAAGUCCCUUUUCGGGCG | 26 | CAC_HDL_AlkB_vs_Cntl_HDL_AlkB | 3.98 | 1.83E-04 |
| UAGGAUUCGGCGCUCUCACCGCCGCGGCCCGGGUUCGAUUCCCGGUCAGGGAA | 53 | CAC_HDL_AlkB_vs_Cntl_HDL_AlkB | 3.91 | 1.23E-02 |
| UCAAAUCUCGGUGGAACCUC | 20 | CAC_HDL_AlkB_vs_Cntl_HDL_AlkB | 3.73 | 2.63E-02 |
| CUAGUGGUUAGGAUUCGGCGCUCUCACCGCCGCGGCCCGGGUUCGAUUCCCGGUCAGGGA | 60 | CAC_HDL_AlkB_vs_Cntl_HDL_AlkB | 3.72 | 1.64E-03 |
| CGGGGUUCGAUUCCCCGACGGGG | 23 | CAC_HDL_AlkB_vs_Cntl_HDL_AlkB | 3.69 | 4.59E-02 |
| ACCAGGGUUCAAGUCCCUGUUCGGGCG | 27 | CAC_HDL_AlkB_vs_Cntl_HDL_AlkB | 3.66 | 2.12E-04 |
| UGUGGGUUCGAGUCCCAUCUGGGUCGCC | 28 | CAC_HDL_AlkB_vs_Cntl_HDL_AlkB | 3.66 | 1.04E-02 |
| UUCCAGGUUCGACUCCUGGCUGGCUCG | 27 | CAC_HDL_AlkB_vs_Cntl_HDL_AlkB | 3.65 | 3.13E-02 |
| ACUCAAGUUCUGGUCUCCGGAUGGAGGCGUGGGUUCGAAUCCCACUUCUGACA | 53 | CAC_HDL_AlkB_vs_Cntl_HDL_AlkB | 3.64 | 2.62E-02 |
| CAUUUGACUGCAGAUCAAGAGGUCCCUGGUUCAAAUCCGGGUGCCCCCU | 49 | CAC_HDL_AlkB_vs_Cntl_HDL_AlkB | 3.61 | 3.93E-02 |
| CGUGUUCAAAUCACGUCGGGGUCA | 24 | CAC_HDL_AlkB_vs_Cntl_HDL_AlkB | 3.54 | 2.33E-03 |
| UGGGUUCGAACCCCACUCCUGGUACC | 26 | CAC_HDL_AlkB_vs_Cntl_HDL_AlkB | 3.52 | 3.35E-02 |
| UCUAGCGGUUAGGAUUCCUGGUUUUCACCCAGGCGGCCCGGGUUCGACUCCCGGUGUGGGAA | 62 | CAC_HDL_AlkB_vs_Cntl_HDL_AlkB | 3.50 | 7.68E-04 |
| UUCCAGGUUCGACUCCUGGCUGGCUCN | 27 | CAC_HDL_AlkB_vs_Cntl_HDL_AlkB | 3.50 | 1.82E-03 |
| CUAGUGGUUAGGAUUCGGCGCUCUCACCGCCGCGGCCCGGGUUCGAUUCCCGGUCAGGGAAC | 62 | CAC_HDL_AlkB_vs_Cntl_HDL_AlkB | 3.50 | 1.48E-03 |
| GUAGAGCAUGAGACUCUUAAUCUCAGGGUCGUGGGUUCGAGCCCCACGUUGGGCG | 55 | CAC_HDL_AlkB_vs_Cntl_HDL_AlkB | 3.48 | 3.90E-02 |
| UAGGAUUCCUGGUUUUCACCCAGGCGGCCCGGGUUCGACUCCCGGUGUGGGAA | 53 | CAC_HDL_AlkB_vs_Cntl_HDL_AlkB | 3.44 | 1.89E-02 |
| GUGAGGGUUCGAGUCCCUUCGUGGUCG | 27 | CAC_HDL_AlkB_vs_Cntl_HDL_AlkB | 3.43 | 6.91E-04 |
| AGGUCGCUGGUUCGAUUCCGGCUCGAAGGA | 30 | CAC_HDL_AlkB_vs_Cntl_HDL_AlkB | 3.40 | 3.32E-02 |
| CCGGGGUUGGAUUCCCCGACGGGGAG | 26 | CAC_HDL_AlkB_vs_Cntl_HDL_AlkB | 3.34 | 6.85E-04 |
| UUGAGUGUUCGAGUCCCUUCGUGGUCG | 27 | CAC_HDL_AlkB_vs_Cntl_HDL_AlkB | 3.34 | 1.84E-03 |
| CGGGCUCGAUUCCCGGUCAGGGAA | 24 | CAC_HDL_AlkB_vs_Cntl_HDL_AlkB | 3.34 | 1.11E-04 |

| | | | | |
|---|---|---|---|---|
| GCAUUGGUGUUUCAGUGGUAGAAUUCUCGCCU | 32 | CAC_HDL_AlkB_vs_Cntl_HDL_AlkB | 3.30 | 1.94E-03 |
| CGUGUUCAAUUCACGUCGGGGUCA | 24 | CAC_HDL_AlkB_vs_Cntl_HDL_AlkB | 3.24 | 2.13E-03 |
| AGGGGUCGAGUCCCUUCGUGGUCG | 24 | CAC_HDL_AlkB_vs_Cntl_HDL_AlkB | 3.20 | 1.50E-03 |
| UAUAGUGGUUAGUACUCUGCGUUGUGGCCGCAGCAACCUCGGUUCGAAUCCGAGUCACGGCA | 62 | CAC_HDL_AlkB_vs_Cntl_HDL_AlkB | 3.17 | 2.68E-03 |
| UUCCUGGUUUUCACCCAGGCGGCCCGGGUUCGACUCCCGGUGUGGGAA | 48 | CAC_HDL_AlkB_vs_Cntl_HDL_AlkB | 3.14 | 2.38E-02 |
| CGAGUUCAAAUCACGUCGGGGUCA | 24 | CAC_HDL_AlkB_vs_Cntl_HDL_AlkB | 3.13 | 5.90E-04 |
| CAGGGGUCAAGUCCCUGUUCGGGCG | 25 | CAC_HDL_AlkB_vs_Cntl_HDL_AlkB | 3.13 | 8.31E-04 |
| AAAUCCCGGACGAGCCCC | 18 | CAC_HDL_AlkB_vs_Cntl_HDL_AlkB | 3.11 | 4.80E-02 |
| UUGGUGGUUCGAGCCCACCCAGGGACNCC | 29 | CAC_HDL_AlkB_vs_Cntl_HDL_AlkB | 3.08 | 3.56E-03 |
| GUUCGACUCCUGGCUGGCUCG | 21 | CAC_HDL_AlkB_vs_Cntl_HDL_AlkB | 3.07 | 1.35E-02 |
| UCCAGUGUUCAAGUCCCUGUUCGGGCG | 27 | CAC_HDL_AlkB_vs_Cntl_HDL_AlkB | 3.07 | 4.00E-03 |
| ACAGGGUUCAAGUCCCUGUUCGGGCG | 26 | CAC_HDL_AlkB_vs_Cntl_HDL_AlkB | 3.02 | 2.21E-03 |
| UGUGUUCGAACCCCACUCCUGGUA | 24 | CAC_HDL_AlkB_vs_Cntl_HDL_AlkB | 3.02 | 1.57E-03 |
| UUGAGGUUUCGAGUCCCUUCGUGGUCG | 27 | CAC_HDL_AlkB_vs_Cntl_HDL_AlkB | 3.01 | 4.41E-04 |
| GAUGGUUCGAGUCCCUUCGUGGUCG | 25 | CAC_HDL_AlkB_vs_Cntl_HDL_AlkB | 3.01 | 7.13E-04 |
| UCACGCGGGAGACCGGGGUUCGAUUCCCCGACGGGGAG | 38 | CAC_HDL_AlkB_vs_Cntl_HDL_AlkB | 3.00 | 4.08E-02 |
| AGAGCAUUUGACUGCAGAUCAAGAGGUCCCCGGUUCAAAUCCGGGUGCCCCCU | 53 | CAC_HDL_AlkB_vs_Cntl_HDL_AlkB | 3.00 | 4.81E-02 |
| CCAGGGUUCAAGUCCCUGUUCGGUCG | 26 | CAC_HDL_AlkB_vs_Cntl_HDL_AlkB | 2.99 | 3.27E-03 |
| UAGUGGUUAGGAUUCGGCGCUCUCACCGCCGCGGCCCGGGUUCGAUUCCCGGUCAGGGAACC | 62 | CAC_HDL_AlkB_vs_Cntl_HDL_AlkB | 2.99 | 8.66E-03 |
| CUAGUGGUUAGGAUUCGGCGCUCUCACCGCCGCGGCCCGGGUUCGAUUCCCGGUCAGGGAACC | 63 | CAC_HDL_AlkB_vs_Cntl_HDL_AlkB | 2.98 | 4.21E-03 |
| CGGGUUCGAUUCCCGGUCAGGGAACC | 26 | CAC_HDL_AlkB_vs_Cntl_HDL_AlkB | 2.96 | 3.92E-02 |
| ACCGGGGUUCGAUUCCCCGACGGGGAGCC | 29 | CAC_HDL_AlkB_vs_Cntl_HDL_AlkB | 2.94 | 3.83E-02 |
| UUGAGGGUUCGAGUCCCCUCGUGGUCG | 27 | CAC_HDL_AlkB_vs_Cntl_HDL_AlkB | 2.90 | 7.14E-04 |
| CUAGUGGUUAGGAUUCGGCGCUCUCACCGCCGCGGCCCGGGUUCGAUUCCCGGUCAGGGAA | 61 | CAC_HDL_AlkB_vs_Cntl_HDL_AlkB | 2.89 | 9.87E-03 |
| UCUAGUGGUUAGGAUUCGGCGCUCUCACCGCCGCGGCCCGGGUUCGAUUCCCGGUCAGGGAA | 62 | CAC_HDL_AlkB_vs_Cntl_HDL_AlkB | 2.86 | 6.71E-03 |

| | | | | |
|---|---|---|---|---|
| GUCGCGGGUUCGAUUCCGGCUCGAAGGA | 28 | CAC_HDL_AlkB_vs_Cntl_HDL_AlkB | 2.86 | 9.71E-03 |
| CUAGCGGUUAGGAUUCCUGGUUUUCACCCAGGCGGCCCGGGUUCGACUCCCGGUGUGGGAA | 61 | CAC_HDL_AlkB_vs_Cntl_HDL_AlkB | 2.85 | 9.61E-03 |
| CGAGUUCAAAUCUCGGUGGAACCUC | 25 | CAC_HDL_AlkB_vs_Cntl_HDL_AlkB | 2.84 | 4.61E-02 |
| GCAGGGUCGAGUCCUGCCGCGGUCG | 25 | CAC_HDL_AlkB_vs_Cntl_HDL_AlkB | 2.84 | 9.52E-03 |
| CGGGGCUCGAUUCCCCGACGGGGAG | 25 | CAC_HDL_AlkB_vs_Cntl_HDL_AlkB | 2.83 | 1.86E-03 |
| UAGUGGUUAGGAUUCGGCGCUCUCACCGCCGCGGCCCGGGUUCGAUUCCCGGUCAGGGAA | 60 | CAC_HDL_AlkB_vs_Cntl_HDL_AlkB | 2.83 | 1.53E-02 |
| AGGGUUCGAGUCCCUUCGUGGUC | 23 | CAC_HDL_AlkB_vs_Cntl_HDL_AlkB | 2.82 | 4.95E-02 |
| CUAGCGGUUAGGAUUCCUGGUUUUCACCCAGGUGGCCCGGGUUCGACUCCCGGUAUGGGAA | 61 | CAC_HDL_AlkB_vs_Cntl_HDL_AlkB | 2.80 | 7.46E-03 |
| UGGUGGUUCGAGCCCACCCAGGGACGCC | 28 | CAC_HDL_AlkB_vs_Cntl_HDL_AlkB | 2.80 | 3.03E-02 |
| UUGAGGGAUCGAGUCCCUUCGUGGUCG | 27 | CAC_HDL_AlkB_vs_Cntl_HDL_AlkB | 2.80 | 7.16E-04 |
| UGAGGGUUCGAGUCCCUUCGUGUUCG | 26 | CAC_HDL_AlkB_vs_Cntl_HDL_AlkB | 2.80 | 2.26E-03 |
| UUUAGGGUUCGAGUCCCUUCGUGGUCG | 27 | CAC_HDL_AlkB_vs_Cntl_HDL_AlkB | 2.78 | 7.54E-04 |
| AUGGUGGUUCGAGCCCACCCAGGGACGCC | 29 | CAC_HDL_AlkB_vs_Cntl_HDL_AlkB | 2.78 | 8.96E-03 |
| CGUGGGUUCGAGCCCCACGUUGGGCGNC | 28 | CAC_HDL_AlkB_vs_Cntl_HDL_AlkB | 2.78 | 3.05E-03 |
| CGGGUUCGAUGCCCGGGCGGCGCA | 24 | CAC_HDL_AlkB_vs_Cntl_HDL_AlkB | 2.76 | 2.96E-03 |
| GCGGGAGACCGGGGUUCGAUUCCCCGACGGGGAG | 34 | CAC_HDL_AlkB_vs_Cntl_HDL_AlkB | 2.76 | 4.77E-02 |
| AGAGCAUGAGACUCUUAAUCUCAGGGUCGUGGGUUCGAGCCCCACGGUGGGCG | 53 | CAC_HDL_AlkB_vs_Cntl_HDL_AlkB | 2.76 | 1.37E-02 |
| GUGGUUAGUACUCUGCGUUGUGGCCGCAGCAACCUCGGUUCGAAUCCGAGUCACGGCA | 58 | CAC_HDL_AlkB_vs_Cntl_HDL_AlkB | 2.75 | 2.77E-02 |
| AACUCUGCGUUGUGGCCGCAGCAACCUCGGUUCGAAUCCGAGUCACGGCA | 50 | CAC_HDL_AlkB_vs_Cntl_HDL_AlkB | 2.75 | 1.29E-02 |
| UAUAGUGGUGAGUAUCCCCGCCUGUCACGCGGGAGACCGGGGUUCGAUUCCCCGACGGGGAG | 62 | CAC_HDL_AlkB_vs_Cntl_HDL_AlkB | 2.74 | 3.23E-02 |
| UUGAGGGUUCGAGUCCCUUCGUUGUCG | 27 | CAC_HDL_AlkB_vs_Cntl_HDL_AlkB | 2.71 | 3.83E-03 |
| UGCGUGUUCGAAUCACGUCGGGGUCA | 26 | CAC_HDL_AlkB_vs_Cntl_HDL_AlkB | 2.71 | 1.41E-02 |
| UUAGUACUCUGCGUUGUGGCCGCAGCNACCUCGGUUCGAAUCCGAGUCACGGCA | 54 | CAC_HDL_AlkB_vs_Cntl_HDL_AlkB | 2.71 | 6.58E-03 |
| CCGGGGUUCGAUUCCCCGACGUGGAG | 26 | CAC_HDL_AlkB_vs_Cntl_HDL_AlkB | 2.71 | 2.16E-04 |

| | | | | |
|---|---|---|---|---|
| AGAGCAUGAGACUCUUAAUCUCAGGGUCGUGGGUUCGAGCCCCACGUUGGGCG | 53 | CAC_HDL_AlkB_vs_Cntl_HDL_AlkB | 2.68 | 4.60E-02 |
| AGGAUUCCUGGUUUUCACCCAGGCGGCCCGGGUUCGACUCCCGGUGUGGGAA | 52 | CAC_HDL_AlkB_vs_Cntl_HDL_AlkB | 2.68 | 4.15E-02 |
| CGGGUUCGAGUCCCGGGCGGCGCA | 24 | CAC_HDL_AlkB_vs_Cntl_HDL_AlkB | 2.67 | 6.06E-03 |
| UAGCGGUUAGGAUUCCUGGUUUUCACCCAGGCGGCCCGGGUUCGACUCCCGGUGUGGGAA | 60 | CAC_HDL_AlkB_vs_Cntl_HDL_AlkB | 2.67 | 8.58E-03 |
| AGAGCAUGGGACUCUUAAUCCCAGGGUCGUGGGUUCGAUCCCCACGUUGGGCG | 53 | CAC_HDL_AlkB_vs_Cntl_HDL_AlkB | 2.67 | 1.01E-02 |
| CGUGUUCGAUUCCCGGGCGGCGCA | 24 | CAC_HDL_AlkB_vs_Cntl_HDL_AlkB | 2.66 | 3.98E-02 |
| CCGGGGUUCGAUUCCCCNACGGGGAG | 26 | CAC_HDL_AlkB_vs_Cntl_HDL_AlkB | 2.66 | 1.61E-03 |
| CCAGGGUUCAAGUCCCUGCUCGGGCG | 26 | CAC_HDL_AlkB_vs_Cntl_HDL_AlkB | 2.66 | 5.57E-03 |
| UAGUGGUUAGUACUCUGCGUUGUGGCCGCAGCAACCUCGGUUCGAAUCCGAGUCACGGCAC | 61 | CAC_HDL_AlkB_vs_Cntl_HDL_AlkB | 2.66 | 3.36E-02 |
| UGUGGGUUCGAGUCCCAUCUGGGGUGNC | 28 | CAC_HDL_AlkB_vs_Cntl_HDL_AlkB | 2.65 | 2.78E-03 |
| UGAGGGUUCGAGUCCCUUAGUGGUCG | 26 | CAC_HDL_AlkB_vs_Cntl_HDL_AlkB | 2.65 | 2.41E-03 |
| GAGGUUCGACUCCUGGCUGGCUCG | 24 | CAC_HDL_AlkB_vs_Cntl_HDL_AlkB | 2.64 | 7.86E-03 |
| UGUGGGUUCGAGUCCCAUCUGGGGUGCC | 28 | CAC_HDL_AlkB_vs_Cntl_HDL_AlkB | 2.63 | 2.80E-02 |
| GAGGGUUCGAGUCCCUUCGUGGGCG | 25 | CAC_HDL_AlkB_vs_Cntl_HDL_AlkB | 2.63 | 4.72E-03 |
| UAGGUUCGACUNCUGGCUGGCUCG | 24 | CAC_HDL_AlkB_vs_Cntl_HDL_AlkB | 2.63 | 2.04E-03 |
| GAGGUUUCGAGUCCCUUCGUGGUCG | 25 | CAC_HDL_AlkB_vs_Cntl_HDL_AlkB | 2.62 | 2.91E-03 |
| AGCAUUUGACUGCAGAUCAAGAGGUCCCCGGUUCAAAUCCGGGUGCCCCCU | 51 | CAC_HDL_AlkB_vs_Cntl_HDL_AlkB | 2.62 | 3.16E-02 |
| CGUGAUCGAAUCACGUCGGGGUCA | 24 | CAC_HDL_AlkB_vs_Cntl_HDL_AlkB | 2.61 | 5.07E-03 |
| UGGGGUCGAAUCCCAUCCUCGUCG | 24 | CAC_HDL_AlkB_vs_Cntl_HDL_AlkB | 2.61 | 6.10E-03 |
| CAGGUGCGACUCCUGGCUGGCUCG | 24 | CAC_HDL_AlkB_vs_Cntl_HDL_AlkB | 2.59 | 2.34E-03 |
| CCGGGGUUCGAAUCCCCGACGGGGAG | 26 | CAC_HDL_AlkB_vs_Cntl_HDL_AlkB | 2.57 | 3.21E-03 |
| CAAGGUUCGACUCCUGGCUGGCUCG | 25 | CAC_HDL_AlkB_vs_Cntl_HDL_AlkB | 2.57 | 5.19E-03 |
| GACCGGGGUUCGAUUCCCCGACGGGGGG | 28 | CAC_HDL_AlkB_vs_Cntl_HDL_AlkB | 2.57 | 1.06E-03 |
| UUGAUGGUUCGAGUCCCUUCGUGGUCG | 27 | CAC_HDL_AlkB_vs_Cntl_HDL_AlkB | 2.56 | 4.88E-03 |
| AGUGGGUUCGAGUCCCAUCUGGGUCG | 26 | CAC_HDL_AlkB_vs_Cntl_HDL_AlkB | 2.56 | 1.04E-02 |

| | | | | |
|---|---|---|---|---|
| UGGGUUCGAACCCCACGCCUGGUA | 24 | CAC_HDL_AlkB_vs_Cntl_HDL_AlkB | 2.56 | 1.46E-02 |
| CAGGUUCGAGUCCUGCCGCGGNCG | 24 | CAC_HDL_AlkB_vs_Cntl_HDL_AlkB | 2.56 | 2.20E-03 |
| AGUGGUUAGGAUUCGGCGCUCUCACCGCCGCGGCCCGGGUUCGAUUCCCGGUCAGGGAA | 59 | CAC_HDL_AlkB_vs_Cntl_HDL_AlkB | 2.52 | 1.68E-02 |
| UUGAGGGUUCGAGUCCCUUCGUGGUNG | 27 | CAC_HDL_AlkB_vs_Cntl_HDL_AlkB | 2.52 | 1.71E-03 |
| CAGUUUCGACUCCUGGCUGGCUCG | 24 | CAC_HDL_AlkB_vs_Cntl_HDL_AlkB | 2.51 | 1.39E-02 |
| UGGGUUCGAUUCCCCGACGGGGAG | 24 | CAC_HDL_AlkB_vs_Cntl_HDL_AlkB | 2.51 | 1.01E-03 |
| GAGGGUUCGAGUCCCUUCGUGUUCG | 25 | CAC_HDL_AlkB_vs_Cntl_HDL_AlkB | 2.50 | 4.52E-03 |
| UUGAGGGUUCGAGUCCCUUCGGGGUCG | 27 | CAC_HDL_AlkB_vs_Cntl_HDL_AlkB | 2.49 | 8.78E-03 |
| UAGUUUCGACUCCUGGCUGGCUCG | 24 | CAC_HDL_AlkB_vs_Cntl_HDL_AlkB | 2.48 | 4.06E-03 |
| CGGGAUCGAUUCCCGGGCGGCGCA | 24 | CAC_HDL_AlkB_vs_Cntl_HDL_AlkB | 2.48 | 3.44E-03 |
| UUCCGGGUUCGAGUCCCGGCGGAGUCN | 27 | CAC_HDL_AlkB_vs_Cntl_HDL_AlkB | 2.47 | 1.13E-02 |
| AGCGUGUUCGAAUCACGUCGGGGUCA | 26 | CAC_HDL_AlkB_vs_Cntl_HDL_AlkB | 2.47 | 3.94E-03 |
| GAGGGUUCGAGUCCCUUCGGGGUCG | 25 | CAC_HDL_AlkB_vs_Cntl_HDL_AlkB | 2.47 | 5.70E-03 |
| AACGGGGUUCGAUUCCCCGACGGGGAG | 27 | CAC_HDL_AlkB_vs_Cntl_HDL_AlkB | 2.47 | 2.89E-03 |
| UGUGGGUUCGAGUCCCAUCUGGGGUGN | 27 | CAC_HDL_AlkB_vs_Cntl_HDL_AlkB | 2.46 | 2.53E-02 |
| CGGGUACGAUUCCCGGGCGGCGCA | 24 | CAC_HDL_AlkB_vs_Cntl_HDL_AlkB | 2.45 | 6.58E-03 |
| ACCGGGGUUCGAUUCCCCGACGGGGAU | 27 | CAC_HDL_AlkB_vs_Cntl_HDL_AlkB | 2.45 | 1.33E-03 |
| GAGUGUUCGAGUCCCUUCGUGGUCG | 25 | CAC_HDL_AlkB_vs_Cntl_HDL_AlkB | 2.45 | 1.16E-02 |
| UGAGGGUUCGAGUCCCUUCGGGGUCG | 26 | CAC_HDL_AlkB_vs_Cntl_HDL_AlkB | 2.45 | 6.07E-03 |
| UGGUGGUUCGAGCCCACCCAGGGACGNC | 28 | CAC_HDL_AlkB_vs_Cntl_HDL_AlkB | 2.45 | 1.50E-02 |
| UUAGUACUCUGCGUUGUGGCCGCAGCAACCUCGGUUCGAAUCCGAGUCACGGCA | 54 | CAC_HDL_AlkB_vs_Cntl_HDL_AlkB | 2.44 | 4.66E-02 |
| ACCGGGGUUCGAUUCCCCGACGGGGAGCCC | 30 | CAC_HDL_AlkB_vs_Cntl_HDL_AlkB | 2.44 | 6.92E-03 |
| GACCGGGGUUCGAUACCCCGACGGGGAG | 28 | CAC_HDL_AlkB_vs_Cntl_HDL_AlkB | 2.42 | 2.78E-03 |
| AGAGCAUGGGACUCUUAAUCCCAGGGUCGUGGGUUCGAGCCCCACGUUUGGCG | 53 | CAC_HDL_AlkB_vs_Cntl_HDL_AlkB | 2.42 | 1.67E-02 |
| UUUUGGGUUCGAGUCCCAUCUGGGUCG | 27 | CAC_HDL_AlkB_vs_Cntl_HDL_AlkB | 2.41 | 4.98E-03 |
| GAGGGUUCGAGUCCCUUCGUGGUCU | 25 | CAC_HDL_AlkB_vs_Cntl_HDL_AlkB | 2.41 | 7.71E-03 |
| CGUGGGUUCGAGCCCCACGUUGGGCGCC | 28 | CAC_HDL_AlkB_vs_Cntl_HDL_AlkB | 2.41 | 4.33E-02 |

| | | | | |
|---|---|---|---|---|
| CCGGGUUCGAUUCCCGGUCAGGGAC | 25 | CAC_HDL_AlkB_vs_Cntl_HDL_AlkB | 2.41 | 7.43E-03 |
| CCAGGGUUCAAGUCCCCGUUCGGGCG | 26 | CAC_HDL_AlkB_vs_Cntl_HDL_AlkB | 2.40 | 4.34E-03 |
| CGUGGGUUCGAAUCCCAUCCUCGUCU | 26 | CAC_HDL_AlkB_vs_Cntl_HDL_AlkB | 2.40 | 8.16E-03 |
| CGGGUUCGACUCCCGGUGUGGUAA | 24 | CAC_HDL_AlkB_vs_Cntl_HDL_AlkB | 2.40 | 2.35E-03 |
| UUGAGGGUUCGAGUCCCUUCGNGGUCG | 27 | CAC_HDL_AlkB_vs_Cntl_HDL_AlkB | 2.40 | 9.31E-03 |
| CCAGGGNUCAAGUCCCUGUUCGGGCG | 26 | CAC_HDL_AlkB_vs_Cntl_HDL_AlkB | 2.39 | 4.27E-03 |
| GAGGGUUCGAGUCCCUUAGUGGUCG | 25 | CAC_HDL_AlkB_vs_Cntl_HDL_AlkB | 2.39 | 1.19E-02 |
| AGUGGUGAGUAUCCCCGCCUGUCACGCGGGAGACCGGGGUUCGAUUCCCCGACGGGGAG | 59 | CAC_HDL_AlkB_vs_Cntl_HDL_AlkB | 2.38 | 2.87E-02 |
| UAGAGCAUGGGACUCUUAAUCCCAGGGUCGUGGGUUCGAGCCCCACGGUGGGCG | 54 | CAC_HDL_AlkB_vs_Cntl_HDL_AlkB | 2.38 | 4.18E-02 |
| UGAGGGUUCGAGUCCCAUCUGGGUCG | 26 | CAC_HDL_AlkB_vs_Cntl_HDL_AlkB | 2.38 | 6.50E-03 |
| GAGGGUUCGAGUCCCUUCGUGGUCG | 25 | CAC_HDL_AlkB_vs_Cntl_HDL_AlkB | 2.37 | 3.36E-02 |
| UUGAGGGUUCGAGUCCCUUCGUGGUAG | 27 | CAC_HDL_AlkB_vs_Cntl_HDL_AlkB | 2.36 | 7.02E-03 |
| CGGGUUCGAUUCCCGGUCGGGGAA | 24 | CAC_HDL_AlkB_vs_Cntl_HDL_AlkB | 2.36 | 2.98E-03 |
| UAGGUUCGACUCCUGGCUGGCNCG | 24 | CAC_HDL_AlkB_vs_Cntl_HDL_AlkB | 2.36 | 7.62E-03 |
| UUGAGGGUUCGAGUCCCGUCGUGGUCG | 27 | CAC_HDL_AlkB_vs_Cntl_HDL_AlkB | 2.36 | 1.21E-02 |
| UGUGGGUUCGAGUCCCAUCUGGGGUU | 26 | CAC_HDL_AlkB_vs_Cntl_HDL_AlkB | 2.35 | 7.03E-03 |
| CCGGGGUUCAAGUCCCUGUUCGGGCG | 26 | CAC_HDL_AlkB_vs_Cntl_HDL_AlkB | 2.35 | 5.29E-03 |
| CGGGUUCGAUUCCCGGGCGGCGCA | 24 | CAC_HDL_AlkB_vs_Cntl_HDL_AlkB | 2.34 | 2.40E-02 |
| UUGUGGGUUCGAGUCCCAUCUGGGUCU | 27 | CAC_HDL_AlkB_vs_Cntl_HDL_AlkB | 2.33 | 8.06E-03 |
| AGAGCAUGGGACUCUUAAUCCCAGGGNCGUGGGUUCGAGCCCCACGUUGGGCG | 53 | CAC_HDL_AlkB_vs_Cntl_HDL_AlkB | 2.33 | 3.07E-02 |
| CGUGAUCAAGUCACGUCGGGGUCA | 24 | CAC_HDL_AlkB_vs_Cntl_HDL_AlkB | 2.32 | 1.95E-02 |
| UAGGUUCGACUCCUGGCUGGCUCU | 24 | CAC_HDL_AlkB_vs_Cntl_HDL_AlkB | 2.32 | 6.33E-03 |
| CAGGUUCGNGUCCUGCCGCGGUCG | 24 | CAC_HDL_AlkB_vs_Cntl_HDL_AlkB | 2.32 | 1.73E-02 |
| GAGGGUUCGAGUCCCUUCGUGGUAG | 25 | CAC_HDL_AlkB_vs_Cntl_HDL_AlkB | 2.32 | 8.16E-03 |
| UUGAGGGUUCGAGUCCCUUCGUGGUCU | 27 | CAC_HDL_AlkB_vs_Cntl_HDL_AlkB | 2.32 | 1.56E-02 |
| CGUGGGUUCGAAUCCCACUUCUUACA | 26 | CAC_HDL_AlkB_vs_Cntl_HDL_AlkB | 2.31 | 1.09E-02 |
| UUGUGGGUUCGAGUCCCAGCUGGGUCG | 27 | CAC_HDL_AlkB_vs_Cntl_HDL_AlkB | 2.30 | 8.51E-03 |

| | | | | |
|---|---|---|---|---|
| CAGGUUCGACUCCUGGCUGGCUCU | 24 | CAC_HDL_AlkB_vs_Cntl_HDL_AlkB | 2.29 | 2.44E-02 |
| GCAUUGGUGGUUCAGUGGUAGAAUUCNCGC | 30 | CAC_HDL_AlkB_vs_Cntl_HDL_AlkB | 2.29 | 1.73E-02 |
| UAGUGGUUAGGAUUCGGCGCUCUCACCGCCGCGGCCCGGGUUCGAUUCCCGGUCAGGGAAC | 61 | CAC_HDL_AlkB_vs_Cntl_HDL_AlkB | 2.28 | 2.10E-02 |
| UUGAGGGUUCGAGUCCCUUCGUGGUCN | 27 | CAC_HDL_AlkB_vs_Cntl_HDL_AlkB | 2.28 | 2.01E-02 |
| CCGGGGUUCAAUUCCCCGACGGGGAG | 26 | CAC_HDL_AlkB_vs_Cntl_HDL_AlkB | 2.27 | 1.37E-02 |
| CCGGGGUUCGAUUCCCCGACGGGGGG | 26 | CAC_HDL_AlkB_vs_Cntl_HDL_AlkB | 2.27 | 7.38E-03 |
| AGCAUUUGACUGCAGAUCAAGAGGUCNCCGGUUCAAAUCCGGGUGCCCCCU | 51 | CAC_HDL_AlkB_vs_Cntl_HDL_AlkB | 2.27 | 2.18E-02 |
| UAGUGGUGAGUAUCCCCGCCUGUCACGCGGGAGACCGGGGUUCGAUUCCCCGACGGGGAG | 60 | CAC_HDL_AlkB_vs_Cntl_HDL_AlkB | 2.27 | 3.55E-02 |
| AGUUUCGACUCCUGGCUGGCUCG | 23 | CAC_HDL_AlkB_vs_Cntl_HDL_AlkB | 2.27 | 1.09E-02 |
| CGGGGUUCGAUUACCCGACGGGGAG | 25 | CAC_HDL_AlkB_vs_Cntl_HDL_AlkB | 2.27 | 1.33E-02 |
| AGGUACGACUCCUGGCUGGCUCG | 23 | CAC_HDL_AlkB_vs_Cntl_HDL_AlkB | 2.26 | 7.82E-03 |
| GCAUUGGUGGUUCAGUGGUAGAAUUCUCGC | 30 | CAC_HDL_AlkB_vs_Cntl_HDL_AlkB | 2.26 | 2.36E-02 |
| UUGAGGGUUCGAGUCCCUUAGUGGUCG | 27 | CAC_HDL_AlkB_vs_Cntl_HDL_AlkB | 2.25 | 1.41E-02 |
| CGUGGGUUCGAAUCCCAUCCUCGUCGCC | 28 | CAC_HDL_AlkB_vs_Cntl_HDL_AlkB | 2.24 | 3.53E-02 |
| CAGGGUUCAAGUCCCUGUNCGGGCG | 25 | CAC_HDL_AlkB_vs_Cntl_HDL_AlkB | 2.24 | 1.17E-02 |
| UUGUGGGUUCGAGUCCCAUCGGGGUCG | 27 | CAC_HDL_AlkB_vs_Cntl_HDL_AlkB | 2.24 | 2.91E-02 |
| CAGGGUUCAAGUCCCAGUUCGGGCG | 25 | CAC_HDL_AlkB_vs_Cntl_HDL_AlkB | 2.23 | 6.10E-03 |
| AGAGCAUGGGACUCUUAAUCCCAGGGUCGUGGGUUCGAGCCCCACGUGGGGCG | 53 | CAC_HDL_AlkB_vs_Cntl_HDL_AlkB | 2.22 | 4.43E-02 |
| AGGGUUCGAGUCCCCUCGUGGUCG | 24 | CAC_HDL_AlkB_vs_Cntl_HDL_AlkB | 2.20 | 1.33E-02 |
| UUGAGGGUUCGAGUCCCUUCGUGGUCG | 27 | CAC_HDL_AlkB_vs_Cntl_HDL_AlkB | 2.20 | 3.09E-02 |
| CAGGGUUCAAGUCCCUGUUGGGGCG | 25 | CAC_HDL_AlkB_vs_Cntl_HDL_AlkB | 2.20 | 7.39E-03 |
| UAGGUUCGACUCCUGGCUGGCUAG | 24 | CAC_HDL_AlkB_vs_Cntl_HDL_AlkB | 2.19 | 1.07E-02 |
| UGAGGGUUCGAGUCCCUUCGUUGUCG | 26 | CAC_HDL_AlkB_vs_Cntl_HDL_AlkB | 2.19 | 1.06E-02 |
| CCAGGGUUCAAGCCCCUGUUCGGGCG | 26 | CAC_HDL_AlkB_vs_Cntl_HDL_AlkB | 2.19 | 2.45E-02 |
| CGUGUUCGAAUCACGUCGGGGUCA | 24 | CAC_HDL_AlkB_vs_Cntl_HDL_AlkB | 2.19 | 4.82E-02 |
| UAGAGGGUUCGAGUCCCUUCGUGGUCG | 27 | CAC_HDL_AlkB_vs_Cntl_HDL_AlkB | 2.18 | 1.75E-02 |

| | | | | |
|---|---|---|---|---|
| CGUGGGUUCGAACCCCACACCUGGUA | 26 | CAC_HDL_AlkB_vs_Cntl_HDL_AlkB | 2.18 | 1.82E-02 |
| GAGGGUUCGAGUCCCGUCGUGGUCG | 25 | CAC_HDL_AlkB_vs_Cntl_HDL_AlkB | 2.18 | 1.89E-02 |
| UAGGUGCGACUCCUGGCUGGCUCG | 24 | CAC_HDL_AlkB_vs_Cntl_HDL_AlkB | 2.18 | 9.38E-03 |
| AGGGUUCGAGUCCCUUCGUGGNCG | 24 | CAC_HDL_AlkB_vs_Cntl_HDL_AlkB | 2.18 | 2.56E-02 |
| UUGCAGGUUCGAGUCCUGCCUCGGUCG | 27 | CAC_HDL_AlkB_vs_Cntl_HDL_AlkB | 2.17 | 2.16E-02 |
| CGUGGGUUCGAACCCCACUCCAGGUA | 26 | CAC_HDL_AlkB_vs_Cntl_HDL_AlkB | 2.16 | 1.65E-02 |
| ACCGGGGUUCGAAUCCCCGACGGGGAG | 27 | CAC_HDL_AlkB_vs_Cntl_HDL_AlkB | 2.16 | 7.50E-03 |
| CGUGGGUUCGAACCCCACGCCUGGUA | 26 | CAC_HDL_AlkB_vs_Cntl_HDL_AlkB | 2.16 | 3.85E-02 |
| CGGGGUCGAUUCCCGGGCGGCGCA | 24 | CAC_HDL_AlkB_vs_Cntl_HDL_AlkB | 2.15 | 2.88E-02 |
| CAGGGUCGAGUCCUGCCGCGGUCG | 24 | CAC_HDL_AlkB_vs_Cntl_HDL_AlkB | 2.15 | 1.70E-02 |
| CCGGGGUUCGAUUCCCCGACGGGNAG | 26 | CAC_HDL_AlkB_vs_Cntl_HDL_AlkB | 2.15 | 7.96E-03 |
| CACGCGGGAGACCGGGGUUCGAUUCCNCGACGGGGAG | 37 | CAC_HDL_AlkB_vs_Cntl_HDL_AlkB | 2.14 | 4.47E-02 |
| GCAGUGGUGGUUCAGUGGUAGAAUUCUCGC | 30 | CAC_HDL_AlkB_vs_Cntl_HDL_AlkB | 2.14 | 4.22E-02 |
| UCCAGGGUUCAAGUCCCUGUUAGGGCG | 27 | CAC_HDL_AlkB_vs_Cntl_HDL_AlkB | 2.14 | 2.01E-02 |
| AUUGAGGGUUCGAGUCCCGUCGUGGUCG | 28 | CAC_HDL_AlkB_vs_Cntl_HDL_AlkB | 2.14 | 2.28E-02 |
| CCGGUGUUCGAUUCCCCGACGGGGAG | 26 | CAC_HDL_AlkB_vs_Cntl_HDL_AlkB | 2.14 | 1.87E-02 |
| UAGGGUCGACUCCUGGCUGGCUCG | 24 | CAC_HDL_AlkB_vs_Cntl_HDL_AlkB | 2.14 | 1.35E-02 |
| UUGAGGGUNCGAGUCCCUUCGUGGUCG | 27 | CAC_HDL_AlkB_vs_Cntl_HDL_AlkB | 2.13 | 1.62E-02 |
| UCGAUGGAUCGAAACCAUCCUCUGCUN | 27 | CAC_HDL_AlkB_vs_Cntl_HDL_AlkB | 2.13 | 3.71E-02 |
| UUGCAGGUUCGAGUCCUGCCGCUGUCG | 27 | CAC_HDL_AlkB_vs_Cntl_HDL_AlkB | 2.12 | 3.74E-02 |
| CGUGGGUUCGAAUCCCAUCCUCGUCGNC | 28 | CAC_HDL_AlkB_vs_Cntl_HDL_AlkB | 2.12 | 2.78E-02 |
| AGAGCAUGGGACUCUUAAUCCCAGGGUCGUGGGGUCGAGCCCCACGUUGGGCG | 53 | CAC_HDL_AlkB_vs_Cntl_HDL_AlkB | 2.11 | 4.44E-02 |
| CNGGGUUCAAGUCCCUGUUCGGGCG | 25 | CAC_HDL_AlkB_vs_Cntl_HDL_AlkB | 2.11 | 1.65E-02 |
| UAGNUUCGACUCCUGGCUGGCUCG | 24 | CAC_HDL_AlkB_vs_Cntl_HDL_AlkB | 2.11 | 3.13E-02 |
| CAGGGUUCAAGUGCCUGUUCGGGCG | 25 | CAC_HDL_AlkB_vs_Cntl_HDL_AlkB | 2.11 | 1.51E-02 |
| AGGGUUCGAGACCCUUCGUGGUCG | 24 | CAC_HDL_AlkB_vs_Cntl_HDL_AlkB | 2.11 | 1.68E-02 |
| UAGGUUCGACUCGUGGCUGGCUCG | 24 | CAC_HDL_AlkB_vs_Cntl_HDL_AlkB | 2.10 | 1.95E-02 |
| CCCGGGUUCGAUUCCCGGUCAGGGAG | 26 | CAC_HDL_AlkB_vs_Cntl_HDL_AlkB | 2.09 | 1.77E-02 |

| | | | | |
|---|---|---|---|---|
| GACCGGGGUUCGAUUCCCCGACGGGGNG | 28 | CAC_HDL_AlkB_vs_Cntl_HDL_AlkB | 2.09 | 1.78E-02 |
| CGUGGGUUCGAACCCCACUCCGGGUA | 26 | CAC_HDL_AlkB_vs_Cntl_HDL_AlkB | 2.09 | 2.84E-02 |
| UAGGUUCGACUCCUGGCUGGGUCG | 24 | CAC_HDL_AlkB_vs_Cntl_HDL_AlkB | 2.09 | 1.78E-02 |
| UAGGUUCGGCUCCUGGCUGGCUCG | 24 | CAC_HDL_AlkB_vs_Cntl_HDL_AlkB | 2.08 | 1.48E-02 |
| CCGGGGUUCGAUUCCCCGACGGGGAU | 26 | CAC_HDL_AlkB_vs_Cntl_HDL_AlkB | 2.08 | 2.45E-02 |
| CGUGGGUUCGAACCCCACUCCUGGUA | 26 | CAC_HDL_AlkB_vs_Cntl_HDL_AlkB | 2.07 | 4.49E-02 |
| CGAGUUCAAGUCACGUCGGGGUCA | 24 | CAC_HDL_AlkB_vs_Cntl_HDL_AlkB | 2.07 | 3.80E-02 |
| CCGGGGUUCGAUUCCCAGACGGGGAG | 26 | CAC_HDL_AlkB_vs_Cntl_HDL_AlkB | 2.07 | 1.02E-02 |
| CGGGUUCGNUUCCCGGGCGGCGCA | 24 | CAC_HDL_AlkB_vs_Cntl_HDL_AlkB | 2.07 | 1.46E-02 |
| CAGGGUUCAAGUCCCUGAUCGGGCG | 25 | CAC_HDL_AlkB_vs_Cntl_HDL_AlkB | 2.07 | 2.05E-02 |
| UGGGAUCGAAUCCCACUCCUGACA | 24 | CAC_HDL_AlkB_vs_Cntl_HDL_AlkB | 2.07 | 2.07E-02 |
| UAGGUUCGACUCCUGGCUGGCACG | 24 | CAC_HDL_AlkB_vs_Cntl_HDL_AlkB | 2.07 | 1.39E-02 |
| CAGGUUCGACUCCUGGCGGGCUCG | 24 | CAC_HDL_AlkB_vs_Cntl_HDL_AlkB | 2.07 | 4.73E-02 |
| CGUGUACAAGUCACGUCGGGGUCA | 24 | CAC_HDL_AlkB_vs_Cntl_HDL_AlkB | 2.07 | 4.48E-02 |
| ACCUGGGUUCGAUUCCCCGACGGGGAG | 27 | CAC_HDL_AlkB_vs_Cntl_HDL_AlkB | 2.07 | 3.55E-03 |
| CCGGGGUUCGAUUCCCCGGCGGGGAG | 26 | CAC_HDL_AlkB_vs_Cntl_HDL_AlkB | 2.06 | 1.19E-02 |
| CGGGUUCGAUUCCCGGGCGGCUCA | 24 | CAC_HDL_AlkB_vs_Cntl_HDL_AlkB | 2.06 | 2.83E-02 |
| AGGGAUCGAGUCCCUUCGUGGUCG | 24 | CAC_HDL_AlkB_vs_Cntl_HDL_AlkB | 2.06 | 1.86E-02 |
| ACGAAACCGGGCGGAAACA | 19 | CAC_HDL_AlkB_vs_Cntl_HDL_AlkB | 2.06 | 4.22E-02 |
| UGGGGUCGAACCCCACUCCUGGUA | 24 | CAC_HDL_AlkB_vs_Cntl_HDL_AlkB | 2.05 | 3.45E-02 |
| CGCGGGGUCGAUCCCCGUACGGGCCA | 26 | CAC_HDL_AlkB_vs_Cntl_HDL_AlkB | 2.05 | 3.95E-02 |
| CCGNGGUUCGAUUCCCCGACGGGGAG | 26 | CAC_HDL_AlkB_vs_Cntl_HDL_AlkB | 2.05 | 1.36E-02 |
| UGGGUUCGAACCCCACUCCUGUUA | 24 | CAC_HDL_AlkB_vs_Cntl_HDL_AlkB | 2.05 | 1.81E-02 |
| UAGGUUCGACUCCUGUCUGGCUCG | 24 | CAC_HDL_AlkB_vs_Cntl_HDL_AlkB | 2.04 | 2.89E-02 |
| GACCGGGGUUCGAUUCCCCGACGGGGAG | 28 | CAC_HDL_AlkB_vs_Cntl_HDL_AlkB | 2.04 | 4.00E-02 |
| AGGGGUUCGAUUCCCCGACGGGGAG | 25 | CAC_HDL_AlkB_vs_Cntl_HDL_AlkB | 2.03 | 4.35E-02 |
| AUGUGGGUUCGAGUCCCAUCUGGGUCG | 27 | CAC_HDL_AlkB_vs_Cntl_HDL_AlkB | 2.03 | 3.89E-02 |
| CGGGUUCGAUUCNCGGUCAGGGAA | 24 | CAC_HDL_AlkB_vs_Cntl_HDL_AlkB | 2.03 | 6.34E-03 |

| | | | | |
|---|---|---|---|---|
| CGGGGUUCGAUUCCCCGACGGUGAG | 25 | CAC_HDL_AlkB_vs_Cntl_HDL_AlkB | 2.03 | 8.96E-03 |
| CCGGGGUUCGAUUCCCCGNCGGGGAG | 26 | CAC_HDL_AlkB_vs_Cntl_HDL_AlkB | 2.02 | 4.13E-03 |
| AGGGUCGACUCCUGGCUGGCUCG | 23 | CAC_HDL_AlkB_vs_Cntl_HDL_AlkB | 2.02 | 3.41E-02 |
| AGGGUUCGAGUCCCUUCGUGUUCG | 24 | CAC_HDL_AlkB_vs_Cntl_HDL_AlkB | 2.01 | 8.69E-03 |
| CGAGGGUUCGAAUCCCACUUCUGACA | 26 | CAC_HDL_AlkB_vs_Cntl_HDL_AlkB | 2.01 | 2.40E-02 |
| CGGGUUCGACUCCCGGUCUGGGAA | 24 | CAC_HDL_AlkB_vs_Cntl_HDL_AlkB | 2.01 | 1.43E-02 |
| CCGGGGUACGAUUCCCCGACGGGGAG | 26 | CAC_HDL_AlkB_vs_Cntl_HDL_AlkB | 2.01 | 8.73E-03 |
| UUGAGGGUUCGAGUCCCAUCUGGGUCG | 27 | CAC_HDL_AlkB_vs_Cntl_HDL_AlkB | 2.00 | 2.99E-02 |
| CGUGGGUACGAGCCCCACGUUGGGCG | 26 | CAC_HDL_AlkB_vs_Cntl_HDL_AlkB | 2.00 | 2.13E-02 |
| CAGGUUCGAGUCCUUCCGCGGUCG | 24 | CAC_HDL_AlkB_vs_Cntl_HDL_AlkB | 2.00 | 3.07E-02 |
| UUGUUGGUUCGAGUCCCAUCUGGGUCG | 27 | CAC_HDL_AlkB_vs_Cntl_HDL_AlkB | 2.00 | 2.24E-02 |
| CAGGGCUCAAGUCCCUGUUCGGGCG | 25 | CAC_HDL_AlkB_vs_Cntl_HDL_AlkB | 2.00 | 2.78E-02 |
| UGGGAUCGAACCCCACUCCUGGUA | 24 | CAC_HDL_AlkB_vs_Cntl_HDL_AlkB | 1.99 | 2.91E-02 |
| CAGGUUCGACUCCUGGCUGGCNCG | 24 | CAC_HDL_AlkB_vs_Cntl_HDL_AlkB | 1.99 | 3.29E-02 |
| CAGGGUUCAAGNCCCUGUUCGGGCG | 25 | CAC_HDL_AlkB_vs_Cntl_HDL_AlkB | 1.99 | 2.81E-02 |
| CGUGGGAUCGAGCCCCACGUUGGGCG | 26 | CAC_HDL_AlkB_vs_Cntl_HDL_AlkB | 1.99 | 3.85E-02 |
| GCAUUGNUGGUUCAGUGGUAGAAUUCUCGCCU | 32 | CAC_HDL_AlkB_vs_Cntl_HDL_AlkB | 1.98 | 2.43E-02 |
| CGGGGUUCGAUUGCCCGACGGGGAG | 25 | CAC_HDL_AlkB_vs_Cntl_HDL_AlkB | 1.98 | 3.36E-02 |
| GGGGUUCGAAUCCCCGACGGGGAG | 24 | CAC_HDL_AlkB_vs_Cntl_HDL_AlkB | 1.98 | 1.02E-02 |
| CGUGGGUUCGAAUCCCAUCCUCGACG | 26 | CAC_HDL_AlkB_vs_Cntl_HDL_AlkB | 1.98 | 2.22E-02 |
| CAGGUUCGAGUCCUGCCGCGGACG | 24 | CAC_HDL_AlkB_vs_Cntl_HDL_AlkB | 1.98 | 1.76E-02 |
| AGAGGGUUCGAGUCCCUUCGUGGUCG | 26 | CAC_HDL_AlkB_vs_Cntl_HDL_AlkB | 1.98 | 1.91E-02 |
| CAGGGGUUCGAUUCCCCGACGGGGAG | 26 | CAC_HDL_AlkB_vs_Cntl_HDL_AlkB | 1.97 | 1.20E-02 |
| GCGGGAGACCGGGGUUCGAUUCCCCGNCGGGGAG | 34 | CAC_HDL_AlkB_vs_Cntl_HDL_AlkB | 1.96 | 2.95E-02 |
| AGGAUCGACUCCUGGCUGGCUCG | 23 | CAC_HDL_AlkB_vs_Cntl_HDL_AlkB | 1.96 | 2.83E-02 |
| CAGGUGCGAGUCCUGCCGCGGUCG | 24 | CAC_HDL_AlkB_vs_Cntl_HDL_AlkB | 1.96 | 2.98E-02 |
| CAGGUUCUACUCCUGGCUGGCUCG | 24 | CAC_HDL_AlkB_vs_Cntl_HDL_AlkB | 1.96 | 3.01E-02 |
| CCCGGGUACGAUUCCCGGUCAGGGAA | 26 | CAC_HDL_AlkB_vs_Cntl_HDL_AlkB | 1.96 | 2.71E-02 |

| | | | | |
|---|---|---|---|---|
| UAGGUUCGACUCCUGGCUGGCUCG | 24 | CAC_HDL_AlkB_vs_Cntl_HDL_AlkB | 1.96 | 2.94E-02 |
| CGGGUUCGAGUCCCGGUCAGGGAA | 24 | CAC_HDL_AlkB_vs_Cntl_HDL_AlkB | 1.95 | 1.45E-02 |
| AGGGUUCGAGUCCCUUCGGGGUCG | 24 | CAC_HDL_AlkB_vs_Cntl_HDL_AlkB | 1.95 | 3.52E-02 |
| CCGGGGUUCGAUUACCCGACGGGGAG | 26 | CAC_HDL_AlkB_vs_Cntl_HDL_AlkB | 1.95 | 2.08E-02 |
| CCUGGGUUCGAUUCCCCGACGGGGAG | 26 | CAC_HDL_AlkB_vs_Cntl_HDL_AlkB | 1.95 | 1.31E-02 |
| AGGGUUCGAGUCCCUGCGUGGUCG | 24 | CAC_HDL_AlkB_vs_Cntl_HDL_AlkB | 1.95 | 3.72E-02 |
| CGGGUGCGAUUCCCGGUCAGGGAA | 24 | CAC_HDL_AlkB_vs_Cntl_HDL_AlkB | 1.94 | 7.32E-03 |
| CGGGGUUCGGUUCCCCGACGGGGAG | 25 | CAC_HDL_AlkB_vs_Cntl_HDL_AlkB | 1.93 | 2.29E-02 |
| CCAGGGUUCAAGUCCCUGUUAGGGCG | 26 | CAC_HDL_AlkB_vs_Cntl_HDL_AlkB | 1.93 | 2.36E-02 |
| UAGGUACGACUCCUGGCUGGCUCG | 24 | CAC_HDL_AlkB_vs_Cntl_HDL_AlkB | 1.93 | 4.27E-02 |
| CGUGGGUUCGAGCCCCACGUGGGGCG | 26 | CAC_HDL_AlkB_vs_Cntl_HDL_AlkB | 1.93 | 3.58E-02 |
| UAGGUUCGNCUCCUGGCUGGCUCG | 24 | CAC_HDL_AlkB_vs_Cntl_HDL_AlkB | 1.93 | 2.94E-02 |
| UCCAGGGAUCAAGUCCCUGUUCGGGCG | 27 | CAC_HDL_AlkB_vs_Cntl_HDL_AlkB | 1.93 | 2.73E-02 |
| CCGGGGUUCGAUACCCCGACGGGGAG | 26 | CAC_HDL_AlkB_vs_Cntl_HDL_AlkB | 1.93 | 2.58E-02 |
| CCAGGGAUCAAGUCCCUGUUCGGGCG | 26 | CAC_HDL_AlkB_vs_Cntl_HDL_AlkB | 1.93 | 1.91E-02 |
| GGGGUUCGAUUCCGCGACGGGGAG | 24 | CAC_HDL_AlkB_vs_Cntl_HDL_AlkB | 1.92 | 2.19E-02 |
| CCAGGGGUCAAGUCCCUGUUCGGGCG | 26 | CAC_HDL_AlkB_vs_Cntl_HDL_AlkB | 1.92 | 3.68E-02 |
| CCGGGGUUCGAUUCCCCGACGGUGAG | 26 | CAC_HDL_AlkB_vs_Cntl_HDL_AlkB | 1.91 | 1.86E-02 |
| CAGGGUUCAAGUCCCUGUUCGGGNG | 25 | CAC_HDL_AlkB_vs_Cntl_HDL_AlkB | 1.91 | 1.98E-02 |
| CAGGGUUCAAGUCACUGUUCGGGCG | 25 | CAC_HDL_AlkB_vs_Cntl_HDL_AlkB | 1.91 | 1.93E-02 |
| UAGGAUCGACUCCUGGCUGGCUCG | 24 | CAC_HDL_AlkB_vs_Cntl_HDL_AlkB | 1.91 | 3.41E-02 |
| AGAGUUCGAGCCUCACCUGGAGCA | 24 | CAC_HDL_AlkB_vs_Cntl_HDL_AlkB | 1.91 | 3.82E-02 |
| AGGGUUCAAGUCCCUGUUGGGGCG | 24 | CAC_HDL_AlkB_vs_Cntl_HDL_AlkB | 1.90 | 3.39E-02 |
| CGGGUUCGAUUCCCGGUCAGGGAC | 24 | CAC_HDL_AlkB_vs_Cntl_HDL_AlkB | 1.90 | 2.15E-02 |
| UUGAGGGUUCGAGUCCCUUCGAGGUCG | 27 | CAC_HDL_AlkB_vs_Cntl_HDL_AlkB | 1.90 | 4.86E-02 |
| CGGGUUCGAUUCCNGGUCAGGGAA | 24 | CAC_HDL_AlkB_vs_Cntl_HDL_AlkB | 1.90 | 1.38E-02 |
| CGAGGGUUCAAGUCCCUGUUCGGGCG | 26 | CAC_HDL_AlkB_vs_Cntl_HDL_AlkB | 1.89 | 3.45E-02 |
| CCANGGUUCAAGUCCCUGUUCGGGCG | 26 | CAC_HDL_AlkB_vs_Cntl_HDL_AlkB | 1.89 | 3.29E-02 |

| | | | | |
|---|---|---|---|---|
| GGGGAUCGAUUCCCCGACGGGGAG | 24 | CAC_HDL_AlkB_vs_Cntl_HDL_AlkB | 1.89 | 8.05E-03 |
| AGGGUUCGNGUCCCUUCGUGGUCG | 24 | CAC_HDL_AlkB_vs_Cntl_HDL_AlkB | 1.88 | 4.67E-02 |
| CGUGGGGUCGAAUCCCAUCCUCGUCG | 26 | CAC_HDL_AlkB_vs_Cntl_HDL_AlkB | 1.88 | 4.60E-02 |
| CGGGUUCGAUUCCGGGUCAGGGAA | 24 | CAC_HDL_AlkB_vs_Cntl_HDL_AlkB | 1.88 | 7.96E-03 |
| CGGGGUUCGAAUCCCCGACGGGGAG | 25 | CAC_HDL_AlkB_vs_Cntl_HDL_AlkB | 1.88 | 2.51E-02 |
| UUGGUUCGAAUCCCAUCCUCGUCG | 24 | CAC_HDL_AlkB_vs_Cntl_HDL_AlkB | 1.88 | 3.72E-02 |
| AGGUUUCGAGUCCCUUCGUGGUCG | 24 | CAC_HDL_AlkB_vs_Cntl_HDL_AlkB | 1.88 | 4.45E-02 |
| CCAGGGUCCAAGUCCCUGUUCGGGCG | 26 | CAC_HDL_AlkB_vs_Cntl_HDL_AlkB | 1.88 | 3.69E-02 |
| CCGGGGUUCGAUUCCCCGACGGGGCG | 26 | CAC_HDL_AlkB_vs_Cntl_HDL_AlkB | 1.88 | 4.91E-02 |
| CCGGGGUUCGAUUCCCCGACGGGGAA | 26 | CAC_HDL_AlkB_vs_Cntl_HDL_AlkB | 1.87 | 2.70E-02 |
| CCGGGGUUCGAUUCCCCGACGGGGUG | 26 | CAC_HDL_AlkB_vs_Cntl_HDL_AlkB | 1.87 | 2.42E-02 |
| CGGGUACGAUUCCCGGUCAGGGAA | 24 | CAC_HDL_AlkB_vs_Cntl_HDL_AlkB | 1.87 | 5.37E-03 |
| CCGGGGUUCGGUUCCCCGACGGGGAG | 26 | CAC_HDL_AlkB_vs_Cntl_HDL_AlkB | 1.87 | 2.55E-02 |
| GGGGUUCGAUUCCCCGACGGGGGG | 24 | CAC_HDL_AlkB_vs_Cntl_HDL_AlkB | 1.87 | 2.82E-02 |
| CGUUGGUUCGAAUCCCAUCCUCGUCG | 26 | CAC_HDL_AlkB_vs_Cntl_HDL_AlkB | 1.87 | 3.47E-02 |
| ACCGGGGUUCGAUACCCCGACGGGGAG | 27 | CAC_HDL_AlkB_vs_Cntl_HDL_AlkB | 1.86 | 3.99E-02 |
| CCGGGGUGCGAUUCCCCGACGGGGAG | 26 | CAC_HDL_AlkB_vs_Cntl_HDL_AlkB | 1.86 | 3.13E-02 |
| CAGNGUUCAAGUCCCUGUUCGGGCG | 25 | CAC_HDL_AlkB_vs_Cntl_HDL_AlkB | 1.86 | 4.20E-02 |
| CCAGGGUUNAAGUCCCUGUUCGGGCG | 26 | CAC_HDL_AlkB_vs_Cntl_HDL_AlkB | 1.86 | 3.11E-02 |
| CCGGGGUUCGAUUCCCCGACGGGGAN | 26 | CAC_HDL_AlkB_vs_Cntl_HDL_AlkB | 1.86 | 2.11E-02 |
| CCAGGGUUCAAGUCCAUGUUCGGGCG | 26 | CAC_HDL_AlkB_vs_Cntl_HDL_AlkB | 1.86 | 3.66E-02 |
| CCCGGGUUCAAGUCCCUGUUCGGGCG | 26 | CAC_HDL_AlkB_vs_Cntl_HDL_AlkB | 1.85 | 4.41E-02 |
| CAGGUUCGAGUCCUGCCGCGGUCG | 24 | CAC_HDL_AlkB_vs_Cntl_HDL_AlkB | 1.85 | 4.95E-02 |
| CGGGAUCGAUUCCCGGUCAGGGAA | 24 | CAC_HDL_AlkB_vs_Cntl_HDL_AlkB | 1.85 | 1.47E-02 |
| CGUGGGAUCGAAUCCCAUCCUCGUCG | 26 | CAC_HDL_AlkB_vs_Cntl_HDL_AlkB | 1.84 | 3.82E-02 |
| CAGGUUCGACUCCUGGCUGGCGCG | 24 | CAC_HDL_AlkB_vs_Cntl_HDL_AlkB | 1.84 | 4.11E-02 |
| CGGGUUCGACUCCCGGUGUGGGAC | 24 | CAC_HDL_AlkB_vs_Cntl_HDL_AlkB | 1.84 | 1.73E-02 |
| ACCGGGGAUCGAUUCCCCGACGGGGAG | 27 | CAC_HDL_AlkB_vs_Cntl_HDL_AlkB | 1.84 | 3.31E-02 |

| | | | | |
|---|---|---|---|---|
| CAGGGUUCAAGUCCCUGUUCGGGGG | 25 | CAC_HDL_AlkB_vs_Cntl_HDL_AlkB | 1.83 | 3.64E-02 |
| CCGGGGUUCGAUUCACCGACGGGGAG | 26 | CAC_HDL_AlkB_vs_Cntl_HDL_AlkB | 1.83 | 2.32E-02 |
| CCGGGGUUCGAUUCCGCGACGGGGAG | 26 | CAC_HDL_AlkB_vs_Cntl_HDL_AlkB | 1.83 | 2.15E-02 |
| CCAGGGUUCAAGUNCCUGUUCGGGCG | 26 | CAC_HDL_AlkB_vs_Cntl_HDL_AlkB | 1.83 | 3.73E-02 |
| ACCGGGGUUCGAUUCCCCGACGGGGGG | 27 | CAC_HDL_AlkB_vs_Cntl_HDL_AlkB | 1.83 | 3.00E-02 |
| CCGGGGUUCGAUUCCCCGACGGGGAG | 26 | CAC_HDL_AlkB_vs_Cntl_HDL_AlkB | 1.82 | 1.44E-02 |
| CGGGUUCGAUUCCCGGUCCGGGAA | 24 | CAC_HDL_AlkB_vs_Cntl_HDL_AlkB | 1.81 | 1.78E-02 |
| ACCGGGGUUCGAUUCCCCGACNGGGAG | 27 | CAC_HDL_AlkB_vs_Cntl_HDL_AlkB | 1.81 | 4.54E-02 |
| CCGGGGUUCGANUCCCCGACGGGGAG | 26 | CAC_HDL_AlkB_vs_Cntl_HDL_AlkB | 1.81 | 3.29E-02 |
| GGGGUGCGAUUCCCCGACGGGGAG | 24 | CAC_HDL_AlkB_vs_Cntl_HDL_AlkB | 1.81 | 1.80E-02 |
| CGGGUUCGAUUCCCGGUCAGGGAG | 24 | CAC_HDL_AlkB_vs_Cntl_HDL_AlkB | 1.81 | 2.53E-02 |
| CGGGGUGCGAUUCCCCGACGGGGAG | 25 | CAC_HDL_AlkB_vs_Cntl_HDL_AlkB | 1.81 | 4.59E-02 |
| UAGGUUCUACUCCUGGCUGGCUCG | 24 | CAC_HDL_AlkB_vs_Cntl_HDL_AlkB | 1.80 | 4.73E-02 |
| ACCGGGGUUCUAUUCCCCGACGGGGAG | 27 | CAC_HDL_AlkB_vs_Cntl_HDL_AlkB | 1.80 | 4.17E-02 |
| CCGGGGUUNGAUUCCCCGACGGGGAG | 26 | CAC_HDL_AlkB_vs_Cntl_HDL_AlkB | 1.80 | 2.56E-02 |
| ACCGUGGUUCGAUUCCCCGACGGGGAG | 27 | CAC_HDL_AlkB_vs_Cntl_HDL_AlkB | 1.80 | 2.55E-02 |
| CCGGGGUUCGAUNCCCCGACGGGGAG | 26 | CAC_HDL_AlkB_vs_Cntl_HDL_AlkB | 1.79 | 3.36E-02 |
| CCGGGGUUCGAUUGCCCGACGGGGAG | 26 | CAC_HDL_AlkB_vs_Cntl_HDL_AlkB | 1.79 | 3.65E-02 |
| CGGGUUGGAUUCCCGGUCAGGGAA | 24 | CAC_HDL_AlkB_vs_Cntl_HDL_AlkB | 1.78 | 1.07E-02 |
| CGGGGUCGACUCCCGGUGUGGGAA | 24 | CAC_HDL_AlkB_vs_Cntl_HDL_AlkB | 1.77 | 3.43E-02 |
| CCAGGGUUCAAGNCCCUGUUCGGGCG | 26 | CAC_HDL_AlkB_vs_Cntl_HDL_AlkB | 1.77 | 4.98E-02 |
| CGGGUUCGACUCCCGGUGGGGGAA | 24 | CAC_HDL_AlkB_vs_Cntl_HDL_AlkB | 1.77 | 4.39E-02 |
| CGGGUUCGAUUCCCGGUCAGGGAA | 24 | CAC_HDL_AlkB_vs_Cntl_HDL_AlkB | 1.77 | 2.72E-02 |
| CUGGGUUCGAUUCCCCGACGGGGAG | 25 | CAC_HDL_AlkB_vs_Cntl_HDL_AlkB | 1.77 | 4.56E-02 |
| CCGGGGUUCGAUUCCCCGACGNGGAG | 26 | CAC_HDL_AlkB_vs_Cntl_HDL_AlkB | 1.77 | 4.95E-02 |
| ACCGGGGUNCGAUUCCCCGACGGGGAG | 27 | CAC_HDL_AlkB_vs_Cntl_HDL_AlkB | 1.75 | 4.77E-02 |
| CCAGGGUUCAAGUCGCUGUUCGGGCG | 26 | CAC_HDL_AlkB_vs_Cntl_HDL_AlkB | 1.75 | 4.62E-02 |
| CGGGUUCGAUUCCCGGGCAGGGAA | 24 | CAC_HDL_AlkB_vs_Cntl_HDL_AlkB | 1.75 | 2.64E-02 |

| | | | | |
|---|---|---|---|---|
| CGGGGUUCNAUUCCCCGACGGGGAG | 25 | CAC_HDL_AlkB_vs_Cntl_HDL_AlkB | 1.74 | 3.21E-02 |
| CGGGGAUCGAUUCCCCGACGGGGAG | 25 | CAC_HDL_AlkB_vs_Cntl_HDL_AlkB | 1.74 | 2.83E-02 |
| CGGGUUCGAUUCCCGGUCAGGGGA | 24 | CAC_HDL_AlkB_vs_Cntl_HDL_AlkB | 1.74 | 3.56E-02 |
| GGGGUUCUAUUCCCCGACGGGGAG | 24 | CAC_HDL_AlkB_vs_Cntl_HDL_AlkB | 1.74 | 4.42E-02 |
| CCGGGGUUCGAUUCCCCGACGCGGAG | 26 | CAC_HDL_AlkB_vs_Cntl_HDL_AlkB | 1.73 | 2.92E-02 |
| CGGGUUCGNUUCCCGGUCAGGGAA | 24 | CAC_HDL_AlkB_vs_Cntl_HDL_AlkB | 1.73 | 2.30E-02 |
| CGGGUUCGAUUCCCGGUCAGGUAA | 24 | CAC_HDL_AlkB_vs_Cntl_HDL_AlkB | 1.72 | 2.37E-02 |
| CCGGGGUUCGAUUNCCCGACGGGGAG | 26 | CAC_HDL_AlkB_vs_Cntl_HDL_AlkB | 1.72 | 4.03E-02 |
| GGGUUUCGAUUCCCCGACGGGGAG | 24 | CAC_HDL_AlkB_vs_Cntl_HDL_AlkB | 1.72 | 4.17E-02 |
| CGGGUUCGAAUCCCGGUCAGGGAA | 24 | CAC_HDL_AlkB_vs_Cntl_HDL_AlkB | 1.72 | 2.99E-02 |
| CGGGUUCGACUCCCGGGAUGGGAA | 24 | CAC_HDL_AlkB_vs_Cntl_HDL_AlkB | 1.72 | 2.75E-02 |
| CGGGUUCGAUACCCGGUCAGGGAA | 24 | CAC_HDL_AlkB_vs_Cntl_HDL_AlkB | 1.71 | 3.31E-02 |
| GGGGUUCGAUACCCCGACGGGGAG | 24 | CAC_HDL_AlkB_vs_Cntl_HDL_AlkB | 1.71 | 4.26E-02 |
| CGGGGUUCGAUUCCCCGGCGGGGAG | 25 | CAC_HDL_AlkB_vs_Cntl_HDL_AlkB | 1.70 | 4.75E-02 |
| GGGGUUCGAUUCCCCGACGGGGAG | 24 | CAC_HDL_AlkB_vs_Cntl_HDL_AlkB | 1.70 | 1.48E-02 |
| CUGGUUCGAUUCCCGGUCAGGGAA | 24 | CAC_HDL_AlkB_vs_Cntl_HDL_AlkB | 1.70 | 2.48E-02 |
| CGGGUUCGAUUCCCGGUCNGGGAA | 24 | CAC_HDL_AlkB_vs_Cntl_HDL_AlkB | 1.66 | 4.97E-02 |
| CGGGUUCGAUUCCCGUUCAGGGAA | 24 | CAC_HDL_AlkB_vs_Cntl_HDL_AlkB | 1.66 | 3.96E-02 |
| CNGGUUCGAUUCCCGGUCAGGGAA | 24 | CAC_HDL_AlkB_vs_Cntl_HDL_AlkB | 1.66 | 4.21E-02 |
| UCCCUGGUGGUCUAGUGGUUAUGAUUCGGCGC | 32 | CAC_HDL_AlkB_vs_Cntl_HDL_AlkB | 0.50 | 3.11E-02 |
| CGUGUUCGACUCCCGGUGUGGGAA | 24 | CAC_HDL_AlkB_vs_Cntl_HDL_AlkB | 0.38 | 1.33E-02 |
| UCCCUGGGGGUCUAGUGGUUAGGAUUCGGCGCUC | 34 | CAC_HDL_AlkB_vs_Cntl_HDL_AlkB | 0.33 | 1.71E-03 |
| CGGGGUUCGAUUCCCCUACGGGGAG | 25 | CAC_HDL_AlkB_vs_Cntl_HDL_AlkB | 0.30 | 7.04E-04 |
| UCCCUGGUGGUCUAGUGGUUAGUAUUCGGCGCUC | 34 | CAC_HDL_AlkB_vs_Cntl_HDL_AlkB | 0.28 | 4.90E-04 |
| UCCCGGGUGGUCUAGUGGUUAGGAUUCGGCGCUC | 34 | CAC_HDL_AlkB_vs_Cntl_HDL_AlkB | 0.28 | 6.80E-04 |
| UCCCUGGUGGUCGAGUGGUUAGGAUUCGGCGCUC | 34 | CAC_HDL_AlkB_vs_Cntl_HDL_AlkB | 0.24 | 1.39E-04 |
| UCCCUGGUGGUCUAGUGGUUAGGAUUNGGCGCUC | 34 | CAC_HDL_AlkB_vs_Cntl_HDL_AlkB | 0.24 | 1.54E-04 |
| CAGUCGGUAGAGCAGGG | 17 | CAC_HDL_AlkB_vs_Cntl_HDL_AlkB | 0.24 | 2.44E-02 |

| | | | | |
|---|---|---|---|---|
| GCAUUGGUGGUUCAGUGGUAGA | 22 | CAC_HDL_AlkB_vs_Cntl_HDL_AlkB | 0.24 | 4.33E-02 |
| CCGGGGUUCGAUUCCCCGUCGGGGAG | 26 | CAC_HDL_AlkB_vs_Cntl_HDL_AlkB | 0.23 | 3.95E-04 |
| UCCCUGGUGGUCUAGUGGUUAGGAUUCUGCGCUC | 34 | CAC_HDL_AlkB_vs_Cntl_HDL_AlkB | 0.22 | 4.91E-05 |
| UCCCUGGUGGUCUAGUGGUUAGGAUUCGGCGCUC | 34 | CAC_HDL_AlkB_vs_Cntl_HDL_AlkB | 0.22 | 8.43E-05 |
| UCCCUGGUGGUCUAGUGGUUAUGAUUCGGCGCUC | 34 | CAC_HDL_AlkB_vs_Cntl_HDL_AlkB | 0.22 | 5.82E-05 |
| UCCCUGGUGGUCUAGGGGUUAGGAUUCGGCGCUC | 34 | CAC_HDL_AlkB_vs_Cntl_HDL_AlkB | 0.20 | 1.00E-04 |
| AUAGGUUCGACUCCUGGCUGGCUCG | 25 | CAC_HDL_AlkB_vs_Cntl_HDL_AlkB | 0.20 | 7.55E-03 |
| AGUCGGUAGAGCAGGG | 16 | CAC_HDL_AlkB_vs_Cntl_HDL_AlkB | 0.19 | 3.67E-02 |
| GCGGGUGUAGCUCAGUGGUAGAGC | 24 | CAC_HDL_AlkB_vs_Cntl_HDL_AlkB | 0.17 | 1.87E-02 |
| AUCCCCGGCAUCUCCAC | 17 | CAC_HDL_AlkB_vs_Cntl_HDL_AlkB | 0.13 | 1.49E-02 |
| GAUAGCUCAGUCGGUAGAG | 19 | CAC_HDL_AlkB_vs_Cntl_HDL_AlkB | 0.12 | 1.70E-02 |
| GGCGGCCCGGGUUCGACUCCCGGUGUGGGAAC | 32 | CAC_HDL_AlkB_vs_Cntl_HDL_AlkB | 0.00 | 6.75E-11 |

**Table S11. Significant differentially abundant non-tDRs on HDL in ASCVD (CAC+).**

| Class | Gene | Comparison | Fold Change | p-value |
|---|---|---|---|---|
| miRNA | hsa-let-7b-5p | CAC_HDL_AlkB_vs_Cntl_HDL_AlkB | 1.95 | 1.86E-03 |
| miRNA | hsa-miR-6869-5p | CAC_HDL_AlkB_vs_Cntl_HDL_AlkB | 0.04 | 2.70E-03 |
| miRNA | hsa-let-7e-5p | CAC_HDL_AlkB_vs_Cntl_HDL_AlkB | 2.07 | 4.34E-03 |
| miRNA | hsa-miR-10a-5p | CAC_HDL_AlkB_vs_Cntl_HDL_AlkB | 1.87 | 1.51E-02 |
| miRNA | hsa-miR-10b-5p | CAC_HDL_AlkB_vs_Cntl_HDL_AlkB | 1.81 | 2.33E-02 |
| miRNA | hsa-miR-1321 | CAC_HDL_AlkB_vs_Cntl_HDL_AlkB | 3.54 | 2.57E-02 |
| miRNA | hsa-let-7a-5p | CAC_HDL_AlkB_vs_Cntl_HDL_AlkB | 1.56 | 3.69E-02 |
| yDR | RNY4P28:ENSG00000202151.1 | CAC_HDL_AlkB_vs_Cntl_HDL_AlkB | 3.93 | 3.22E-04 |
| yDR | Y_RNA:ENSG00000201178.1 | CAC_HDL_AlkB_vs_Cntl_HDL_AlkB | 6.08 | 1.79E-03 |
| yDR | Y_RNA:ENSG00000199223.1 | CAC_HDL_AlkB_vs_Cntl_HDL_AlkB | 3.23 | 2.30E-03 |
| yDR | Y_RNA:ENSG00000199584.1 | CAC_HDL_AlkB_vs_Cntl_HDL_AlkB | 0.29 | 6.91E-03 |
| yDR | Y_RNA:ENSG00000201548.1 | CAC_HDL_AlkB_vs_Cntl_HDL_AlkB | 0.29 | 7.68E-03 |
| yDR | Y_RNA:ENSG00000201678.1 | CAC_HDL_AlkB_vs_Cntl_HDL_AlkB | 2.57 | 1.24E-02 |
| yDR | Y_RNA:ENSG00000201208.1 | CAC_HDL_AlkB_vs_Cntl_HDL_AlkB | 2.71 | 1.41E-02 |
| yDR | Y_RNA:ENSG00000207061.1 | CAC_HDL_AlkB_vs_Cntl_HDL_AlkB | 2.02 | 2.08E-02 |

| | | | | |
|---|---|---|---|---|
| yDR | Y_RNA:ENSG00000206905.1 | CAC_HDL_AlkB_vs_Cntl_HDL_AlkB | 2.32 | 2.23E-02 |
| yDR | Y_RNA:ENSG00000200506.1 | CAC_HDL_AlkB_vs_Cntl_HDL_AlkB | 1.89 | 2.30E-02 |
| yDR | RNY1:ENSG00000201098.1 | CAC_HDL_AlkB_vs_Cntl_HDL_AlkB | 1.85 | 2.61E-02 |
| yDR | Y_RNA:ENSG00000199285.1 | CAC_HDL_AlkB_vs_Cntl_HDL_AlkB | 2.28 | 2.82E-02 |
| yDR | RNY4P8:ENSG00000200735.1 | CAC_HDL_AlkB_vs_Cntl_HDL_AlkB | 1.72 | 3.06E-02 |
| yDR | Y_RNA:ENSG00000200494.1 | CAC_HDL_AlkB_vs_Cntl_HDL_AlkB | 2.15 | 3.56E-02 |
| yDR | RNY3P1:ENSG00000201955.1 | CAC_HDL_AlkB_vs_Cntl_HDL_AlkB | 2.00 | 3.67E-02 |
| yDR | RNY4P11:ENSG00000252403.1 | CAC_HDL_AlkB_vs_Cntl_HDL_AlkB | 1.71 | 3.73E-02 |
| yDR | Y_RNA:ENSG00000200615.1 | CAC_HDL_AlkB_vs_Cntl_HDL_AlkB | 1.66 | 4.42E-02 |
| yDR | Y_RNA:ENSG00000238845.1 | CAC_HDL_AlkB_vs_Cntl_HDL_AlkB | 2.04 | 4.75E-02 |
| yDR | Y_RNA:ENSG00000207142.1 | CAC_HDL_AlkB_vs_Cntl_HDL_AlkB | 1.75 | 4.81E-02 |
| yDR | Y_RNA:ENSG00000201071.1 | CAC_HDL_AlkB_vs_Cntl_HDL_AlkB | 1.95 | 4.83E-02 |
| osRNA | RN7SL674P:ENSG00000239899.2 | CAC_HDL_AlkB_vs_Cntl_HDL_AlkB | 7.25 | 2.47E-03 |
| osRNA | RN7SL396P:ENSG00000244642.2 | CAC_HDL_AlkB_vs_Cntl_HDL_AlkB | 6.56 | 3.93E-03 |
| rDR | RNA5SP358:ENSG00000199843.1 | CAC_HDL_AlkB_vs_Cntl_HDL_AlkB | 1.98 | 1.16E-03 |
| rDR | RNA5S9:ENSG00000201321.1 | CAC_HDL_AlkB_vs_Cntl_HDL_AlkB | 2.02 | 1.19E-03 |
| rDR | RNA5S1:ENSG00000199352.1 | CAC_HDL_AlkB_vs_Cntl_HDL_AlkB | 1.74 | 5.70E-03 |
| rDR | RNA5SP202:ENSG00000201185.1 | CAC_HDL_AlkB_vs_Cntl_HDL_AlkB | 1.88 | 7.63E-03 |
| rDR | RNA5SP194:ENSG00000201532.1 | CAC_HDL_AlkB_vs_Cntl_HDL_AlkB | 0.29 | 1.07E-02 |
| rDR | 5S_rRNA:ENSG00000272435.1 | CAC_HDL_AlkB_vs_Cntl_HDL_AlkB | 1.80 | 1.29E-02 |
| rDR | RNA5SP402:ENSG00000252957.1 | CAC_HDL_AlkB_vs_Cntl_HDL_AlkB | 0.54 | 1.53E-02 |
| rDR | RNA5SP248:ENSG00000202472.1 | CAC_HDL_AlkB_vs_Cntl_HDL_AlkB | 1.73 | 1.95E-02 |
| rDR | RNA5SP500:ENSG00000201356.1 | CAC_HDL_AlkB_vs_Cntl_HDL_AlkB | 1.72 | 2.05E-02 |
| rDR | RNA5SP469:ENSG00000201035.1 | CAC_HDL_AlkB_vs_Cntl_HDL_AlkB | 1.69 | 2.69E-02 |
| rDR | RNA5-8SP6:ENSG00000251705.1 | CAC_HDL_AlkB_vs_Cntl_HDL_AlkB | 0.43 | 4.91E-02 |
| snDR | RNVU1-18:ENSG00000206737.1 | CAC_HDL_AlkB_vs_Cntl_HDL_AlkB | 3.29 | 2.52E-03 |
| snDR | RNU6-35P:ENSG00000207260.1 | CAC_HDL_AlkB_vs_Cntl_HDL_AlkB | 0.28 | 3.27E-03 |

| | | | | |
|---|---|---|---|---|
| snDR | RNU6-31P:ENSG00000207116.1 | CAC_HDL_AlkB_vs_Cntl_HDL_AlkB | 0.38 | 6.24E-03 |
| snDR | RNU6-671P:ENSG00000239178.1 | CAC_HDL_AlkB_vs_Cntl_HDL_AlkB | 0.54 | 1.87E-02 |
| snDR | RNU6-761P:ENSG00000206875.1 | CAC_HDL_AlkB_vs_Cntl_HDL_AlkB | 0.55 | 1.88E-02 |
| snDR | RNU6-43P:ENSG00000207029.1 | CAC_HDL_AlkB_vs_Cntl_HDL_AlkB | 0.49 | 4.12E-02 |
| snDR | RNU6-16P:ENSG00000207113.1 | CAC_HDL_AlkB_vs_Cntl_HDL_AlkB | 0.65 | 4.74E-02 |
| snDR | RNU6-15P:ENSG00000207264.1 | CAC_HDL_AlkB_vs_Cntl_HDL_AlkB | 0.65 | 4.79E-02 |

**Table S12. Significant ARG-ACG-1-1 cleaved tDR changes in CAC$^+$HDL**

| tDRs | Length | Fold Change | p-value |
|---|---|---|---|
| GTTCGACTCCTGGCTGGCTCG | 21 | 3.07 | 1.35E-02 |
| GGTTCGACTCCTGGCTGGCTCG | 22 | 7.92 | 4.66E-03 |
| CAGGTTCGACTCCTGGCTGGCT | 28 | 4.94 | 1.28E-02 |
| AGGTACGACTCCTGGCTGGCTCG | 23 | 2.26 | 7.82E-03 |
| AGTTTCGACTCCTGGCTGGCTCG | 23 | 2.27 | 1.09E-02 |
| AGGATCGACTCCTGGCTGGCTCG | 23 | 1.96 | 2.83E-02 |
| AGGGTCGACTCCTGGCTGGCTCG | 23 | 2.02 | 3.41E-02 |
| CAGGTTCGACTCCTGGCTGTCTCG | 24 | 7.92 | 4.52E-06 |
| CAGGTGCGACTCCTGGCTGGCTCG | 24 | 2.59 | 2.34E-03 |
| GAGGTTCGACTCCTGGCTGGCTCG | 24 | 2.64 | 7.86E-03 |
| CAGTTTCGACTCCTGGCTGGCTCG | 24 | 2.51 | 1.39E-02 |
| CAGGTTCGACTCCTGGCTGGCTCT | 24 | 2.29 | 2.44E-02 |
| CAGGTTCTACTCCTGGCTGGCTCG | 24 | 1.96 | 3.01E-02 |
| CAGGTTCGACTCCTGGCTGGCNCG | 24 | 1.99 | 3.29E-02 |
| CAGGTTCGACTCCTGGCGGGCTCG | 24 | 2.07 | 4.73E-02 |
| CAAGGTTCGACTCCTGGCTGGCTCG | 25 | 2.57 | 5.19E-03 |
| TTCCAGGTTCGACTCCTGGCTGGCTCN | 27 | 3.50 | 1.82E-03 |
| TTCCAGGTTCGACTCCTGGCTGGCTCG | 27 | 3.65 | 3.13E-02 |

**Table S13. Significant differentially altered protein coding genes upon sHDL-delivery of m$^1$A-tDR-ArgACG-1-21**

| Gene Symbol | Fold Change | padj |
|---|---|---|

| | | |
|---|---|---|
| CXCL5 | 34.53 | 7.64E-06 |
| SERPINB2 | 32.98 | 3.69E-37 |
| TNFSF15 | 11.81 | 4.46E-47 |
| AKR1C3 | 10.38 | 4.72E-07 |
| SLC7A11 | 8.77 | 2.08E-113 |
| TREM1 | 8.20 | 5.12E-06 |
| VNN1 | 7.85 | 2.17E-04 |
| SEL1L | 7.56 | 6.84E-61 |
| SRXN1 | 7.20 | 2.12E-13 |
| IL6 | 7.18 | 1.97E-73 |
| GCLM | 6.99 | 3.08E-23 |
| HLA-DRB5 | 6.98 | 2.01E-05 |
| YRDC | 6.71 | 1.94E-03 |
| CLEC5A | 6.42 | 2.40E-13 |
| PLEKHM3 | 6.19 | 6.66E-27 |
| RAP1GDS1 | 5.95 | 5.89E-04 |
| CSF2 | 5.85 | 2.53E-08 |
| IL1A | 5.79 | 2.75E-26 |
| THBS1 | 5.74 | 1.22E-11 |
| MS4A14 | 5.69 | 6.25E-04 |
| SPCS3 | 5.53 | 3.52E-11 |
| TFEC | 5.52 | 5.35E-16 |
| PGK1 | 5.51 | 1.21E-13 |
| MED8 | 5.31 | 1.92E-02 |
| CYP1B1 | 5.30 | 1.18E-85 |
| TNFAIP6 | 5.27 | 3.15E-27 |
| MMADHC | 5.24 | 1.61E-03 |
| STARD7 | 5.22 | 2.00E-13 |
| MRPL51 | 5.18 | 8.17E-04 |
| MS4A7 | 5.14 | 3.55E-14 |
| NAMPT | 5.13 | 1.16E-100 |
| TXNRD1 | 5.00 | 1.36E-82 |
| AGPAT5 | 4.99 | 2.36E-02 |
| CXCL8 | 4.89 | 6.81E-94 |
| DHX32 | 4.80 | 1.04E-03 |
| ZRANB1 | 4.79 | 6.40E-04 |
| LAPTM4A | 4.78 | 1.04E-06 |
| COX8A | 4.74 | 3.77E-05 |
| PID1 | 4.66 | 4.53E-05 |
| STX16 | 4.66 | 4.52E-04 |
| GTF3C4 | 4.64 | 3.85E-04 |
| SLC2A3 | 4.64 | 1.40E-55 |
| CEACAM8 | 4.63 | 9.85E-24 |
| FAM3C | 4.61 | 3.15E-03 |

| | | |
|---|---|---|
| CD274 | 4.59 | 3.19E-08 |
| NDUFB5 | 4.57 | 2.76E-03 |
| CPVL | 4.56 | 5.62E-05 |
| LYZ | 4.51 | 6.18E-97 |
| DNAJC3 | 4.51 | 9.09E-19 |
| RAB12 | 4.47 | 1.03E-06 |
| TMEM208 | 4.45 | 2.18E-02 |
| CMAS | 4.40 | 1.44E-03 |
| SDC2 | 4.40 | 7.47E-13 |
| IL23A | 4.40 | 6.55E-12 |
| IFNGR2 | 4.39 | 2.34E-14 |
| IL1B | 4.38 | 3.45E-86 |
| MMGT1 | 4.33 | 6.08E-03 |
| ATP13A3 | 4.31 | 9.93E-45 |
| CD80 | 4.31 | 2.33E-03 |
| GFPT1 | 4.25 | 2.53E-10 |
| PLAA | 4.24 | 2.75E-08 |
| PTX3 | 4.22 | 2.44E-13 |
| DNAJB9 | 4.22 | 1.46E-12 |
| CMTM6 | 4.17 | 4.35E-12 |
| NIBAN1 | 4.15 | 1.89E-28 |
| ATXN1L | 4.14 | 5.03E-06 |
| PON2 | 4.14 | 5.00E-03 |
| TXN | 4.12 | 1.91E-29 |
| MARCKS | 4.12 | 2.80E-22 |
| SRSF1 | 4.12 | 4.17E-03 |
| PTGS2 | 4.10 | 3.05E-36 |
| MSANTD3 | 4.10 | 2.71E-07 |
| TMED7 | 4.09 | 1.14E-04 |
| DYNLT1 | 4.09 | 1.23E-02 |
| FAM114A2 | 4.08 | 7.82E-03 |
| INHBA | 4.08 | 9.37E-09 |
| HSPA13 | 4.07 | 8.76E-13 |
| ACSL1 | 4.05 | 1.41E-48 |
| IFT57 | 4.04 | 1.41E-02 |
| TDP2 | 4.04 | 1.70E-20 |
| ARFGAP3 | 4.03 | 1.27E-13 |
| FCAR | 4.02 | 1.70E-06 |
| SAMSN1 | 4.00 | 1.42E-07 |
| OGFRL1 | 4.00 | 1.03E-10 |
| ATXN1 | 4.00 | 5.20E-24 |
| HSPA5 | 3.97 | 4.33E-50 |
| CS | 3.96 | 1.71E-10 |
| SCPEP1 | 3.96 | 4.53E-13 |

| | | |
|---|---|---|
| ACSL4 | 3.93 | 1.44E-12 |
| KYNU | 3.92 | 4.43E-25 |
| RNF11 | 3.90 | 2.92E-07 |
| SLAMF7 | 3.88 | 1.53E-34 |
| USP12 | 3.86 | 1.64E-21 |
| GSR | 3.82 | 4.25E-20 |
| PTPRE | 3.80 | 1.75E-35 |
| LYN | 3.78 | 9.18E-29 |
| TMX1 | 3.78 | 2.45E-03 |
| ARRDC4 | 3.77 | 8.96E-12 |
| VCP | 3.74 | 1.59E-19 |
| PDIA3 | 3.74 | 7.22E-17 |
| DERL2 | 3.73 | 4.83E-04 |
| DLD | 3.72 | 1.24E-07 |
| HIP1 | 3.72 | 2.04E-19 |
| CPD | 3.70 | 1.96E-18 |
| RAB10 | 3.69 | 1.82E-12 |
| HLA-DRB1 | 3.68 | 1.85E-34 |
| ADAM10 | 3.68 | 4.44E-14 |
| CERS6 | 3.66 | 2.09E-03 |
| CCNH | 3.65 | 1.48E-02 |
| TLR2 | 3.65 | 3.02E-04 |
| GTF2E1 | 3.64 | 2.45E-03 |
| RFFL | 3.63 | 2.15E-05 |
| KNSTRN | 3.61 | 2.14E-02 |
| CCL24 | 3.60 | 5.52E-14 |
| CLTC | 3.60 | 1.04E-28 |
| SF3B6 | 3.58 | 4.05E-03 |
| PLAUR | 3.58 | 7.47E-47 |
| PRNP | 3.57 | 1.47E-09 |
| TNIP1 | 3.55 | 2.85E-30 |
| BACH1 | 3.53 | 5.55E-18 |
| MTMR6 | 3.53 | 1.04E-06 |
| GNB4 | 3.53 | 1.88E-04 |
| S100A8 | 3.52 | 3.20E-05 |
| CXCL16 | 3.51 | 5.47E-10 |
| TIMP1 | 3.51 | 5.30E-24 |
| ATP1B1 | 3.50 | 5.03E-10 |
| HIRA | 3.48 | 3.44E-02 |
| TMED5 | 3.48 | 5.88E-08 |
| AASDHPPT | 3.47 | 3.29E-02 |
| GLB1 | 3.46 | 5.88E-08 |
| UGDH | 3.43 | 1.29E-03 |
| CD44 | 3.43 | 8.21E-55 |

| | | |
|---|---|---|
| GRK3 | 3.43 | 4.06E-06 |
| KLHL12 | 3.42 | 3.81E-03 |
| SRGN | 3.41 | 3.44E-26 |
| GRN | 3.41 | 7.28E-34 |
| NDUFV2 | 3.41 | 5.77E-03 |
| OSGIN2 | 3.41 | 9.09E-07 |
| CYP27A1 | 3.41 | 8.53E-06 |
| ABHD4 | 3.40 | 1.82E-09 |
| PTPRJ | 3.39 | 6.34E-33 |
| KCNN4 | 3.39 | 1.28E-07 |
| ATP6V1A | 3.38 | 4.02E-12 |
| MRPL38 | 3.34 | 4.06E-02 |
| ITPRIPL2 | 3.34 | 4.06E-06 |
| MEN1 | 3.33 | 1.38E-03 |
| TRAM1 | 3.33 | 2.56E-16 |
| EDRF1 | 3.33 | 4.46E-02 |
| WDFY1 | 3.33 | 1.14E-05 |
| RDH11 | 3.31 | 4.11E-03 |
| NSF | 3.31 | 3.61E-02 |
| AKR1C1 | 3.30 | 8.99E-05 |
| NCF2 | 3.30 | 2.00E-30 |
| EXOSC6 | 3.28 | 1.71E-04 |
| TMED2 | 3.27 | 4.13E-07 |
| ME1 | 3.27 | 1.63E-10 |
| STT3A | 3.27 | 9.94E-07 |
| EVI2A | 3.26 | 5.50E-03 |
| AGO4 | 3.26 | 2.66E-05 |
| SRSF9 | 3.25 | 3.93E-03 |
| ACSL5 | 3.25 | 2.05E-09 |
| HMGCR | 3.25 | 1.14E-07 |
| TNIP3 | 3.24 | 2.17E-04 |
| HLA-DRA | 3.24 | 3.00E-49 |
| COPS2 | 3.24 | 5.12E-08 |
| ELL2 | 3.23 | 4.29E-06 |
| SOCS6 | 3.23 | 1.64E-02 |
| IL2RA | 3.22 | 2.47E-05 |
| BID | 3.22 | 1.75E-09 |
| TGFBR1 | 3.21 | 1.64E-03 |
| SMS | 3.20 | 2.78E-09 |
| COP1 | 3.20 | 1.83E-05 |
| EXOC5 | 3.20 | 2.62E-06 |
| GMPR2 | 3.19 | 1.87E-03 |
| AHCYL1 | 3.19 | 5.25E-08 |
| HMOX1 | 3.19 | 2.99E-60 |

| | | |
|---|---|---|
| CSGALNACT2 | 3.18 | 3.21E-12 |
| CDC123 | 3.18 | 1.97E-03 |
| MFSD14B | 3.17 | 7.51E-04 |
| GOLGA2 | 3.17 | 6.83E-12 |
| NSMCE2 | 3.15 | 3.81E-02 |
| ERBIN | 3.15 | 1.88E-15 |
| ELOA | 3.14 | 1.84E-08 |
| HYOU1 | 3.14 | 2.67E-16 |
| SLC25A46 | 3.14 | 9.21E-03 |
| NPLOC4 | 3.14 | 3.54E-16 |
| SEC24D | 3.13 | 2.26E-12 |
| ZC3H12C | 3.13 | 5.43E-13 |
| TNIK | 3.13 | 2.62E-13 |
| TMX2 | 3.13 | 6.92E-03 |
| CASP4 | 3.12 | 1.04E-06 |
| FBXO30 | 3.12 | 9.87E-05 |
| NPTN | 3.12 | 2.45E-07 |
| SUSD6 | 3.12 | 9.25E-19 |
| VWA5A | 3.11 | 7.22E-05 |
| TNKS2 | 3.11 | 6.53E-08 |
| UBE2J1 | 3.09 | 6.76E-10 |
| COPB1 | 3.09 | 2.15E-09 |
| SLC8A1 | 3.08 | 5.28E-05 |
| NUMB | 3.08 | 1.49E-16 |
| SMARCA5 | 3.08 | 1.34E-07 |
| LGALS3BP | 3.07 | 7.92E-07 |
| ASAP1 | 3.07 | 1.28E-36 |
| CAPZA1 | 3.07 | 7.89E-10 |
| UBE2Z | 3.07 | 1.77E-07 |
| VPS4B | 3.06 | 1.93E-06 |
| FLNA | 3.06 | 5.72E-53 |
| KMO | 3.05 | 7.39E-06 |
| CDC42SE2 | 3.05 | 4.55E-07 |
| NINJ1 | 3.05 | 2.21E-23 |
| RBAK | 3.04 | 1.69E-02 |
| CALU | 3.04 | 2.82E-06 |
| GANAB | 3.04 | 1.19E-10 |
| XRCC1 | 3.04 | 1.69E-02 |
| LRP12 | 3.03 | 5.49E-07 |
| ERP44 | 3.03 | 1.09E-05 |
| STRAP | 3.03 | 1.22E-04 |
| SEC63 | 3.02 | 2.12E-07 |
| NOL7 | 3.02 | 5.48E-04 |
| CNTNAP3 | 3.02 | 1.69E-02 |

| | | |
|---|---|---|
| CLEC12A | 3.01 | 8.13E-04 |
| NIBAN2 | 3.01 | 1.27E-13 |
| CKAP4 | 3.00 | 1.65E-04 |
| FAF2 | 3.00 | 5.03E-05 |
| CYB5R1 | 2.99 | 2.67E-05 |
| C5orf15 | 2.99 | 5.20E-03 |
| PSMD13 | 2.98 | 9.24E-04 |
| CRIM1 | 2.98 | 2.53E-10 |
| NCOA3 | 2.98 | 1.36E-09 |
| GHITM | 2.97 | 1.97E-09 |
| ZKSCAN8 | 2.97 | 2.78E-04 |
| SURF4 | 2.97 | 3.52E-13 |
| PDHA1 | 2.96 | 8.90E-04 |
| CPSF7 | 2.96 | 9.37E-09 |
| ZFYVE16 | 2.95 | 8.93E-18 |
| PIKFYVE | 2.95 | 5.33E-08 |
| ADAM17 | 2.95 | 9.02E-12 |
| SLC39A6 | 2.95 | 3.22E-03 |
| NAA50 | 2.95 | 2.69E-06 |
| POMP | 2.94 | 5.16E-04 |
| PANK3 | 2.94 | 2.67E-05 |
| RAB5B | 2.94 | 6.96E-05 |
| DOCK4 | 2.93 | 1.28E-09 |
| PPP3R1 | 2.93 | 4.33E-04 |
| UFC1 | 2.93 | 1.21E-02 |
| FEZ2 | 2.93 | 5.19E-03 |
| FBXL5 | 2.92 | 1.92E-06 |
| PPME1 | 2.92 | 1.64E-03 |
| PPT1 | 2.92 | 2.34E-21 |
| MTDH | 2.91 | 8.10E-09 |
| APP | 2.91 | 2.56E-10 |
| XRN2 | 2.91 | 1.13E-06 |
| BTF3 | 2.91 | 2.68E-10 |
| MDFIC | 2.90 | 8.08E-04 |
| PCGF5 | 2.90 | 1.52E-03 |
| SOD2 | 2.90 | 5.54E-66 |
| C1orf122 | 2.90 | 7.64E-04 |
| NCF1 | 2.90 | 5.05E-09 |
| ASPH | 2.90 | 4.02E-11 |
| IL7R | 2.90 | 1.72E-14 |
| SDE2 | 2.89 | 3.27E-03 |
| PPP1CB | 2.89 | 2.91E-09 |
| OSTC | 2.89 | 6.48E-03 |
| BNIP3L | 2.88 | 1.37E-04 |

| | | |
|---|---|---|
| EIF1AX | 2.88 | 1.27E-04 |
| PSAP | 2.88 | 1.46E-37 |
| SLK | 2.87 | 4.04E-06 |
| EIF4ENIF1 | 2.87 | 1.54E-03 |
| HLA-DPB1 | 2.87 | 3.83E-20 |
| KCTD20 | 2.86 | 1.60E-08 |
| PKM | 2.86 | 2.14E-59 |
| SOS1 | 2.86 | 1.05E-04 |
| ANXA5 | 2.86 | 3.16E-26 |
| NBN | 2.85 | 5.96E-16 |
| EMB | 2.85 | 2.46E-03 |
| NSFL1C | 2.85 | 2.70E-04 |
| MRC1 | 2.85 | 1.03E-42 |
| OSBP | 2.84 | 1.37E-04 |
| ILRUN | 2.83 | 5.04E-07 |
| IL12B | 2.83 | 8.05E-05 |
| ITGAX | 2.83 | 3.33E-24 |
| LPGAT1 | 2.83 | 4.77E-06 |
| DAD1 | 2.83 | 2.97E-03 |
| ZNF124 | 2.83 | 2.09E-02 |
| PSD3 | 2.82 | 6.19E-07 |
| COPA | 2.82 | 5.18E-09 |
| NAB1 | 2.82 | 4.77E-02 |
| AQP9 | 2.82 | 1.13E-09 |
| NCEH1 | 2.81 | 2.51E-09 |
| XPOT | 2.81 | 2.22E-09 |
| EPRS1 | 2.81 | 4.59E-06 |
| SEC22B | 2.80 | 2.35E-07 |
| ZNRF2 | 2.80 | 4.97E-03 |
| EDEM1 | 2.80 | 1.71E-08 |
| RAB1A | 2.80 | 1.66E-06 |
| FCN1 | 2.80 | 8.04E-11 |
| MGST1 | 2.80 | 3.05E-03 |
| TLK1 | 2.79 | 6.61E-07 |
| NFKB1 | 2.79 | 5.06E-27 |
| PEX19 | 2.79 | 1.63E-02 |
| FAM126B | 2.79 | 7.89E-03 |
| ARHGAP31 | 2.78 | 1.76E-13 |
| SERINC1 | 2.78 | 4.98E-08 |
| GNL2 | 2.78 | 3.77E-03 |
| BTG3 | 2.78 | 6.83E-03 |
| KRAS | 2.78 | 4.00E-03 |
| PSMC4 | 2.78 | 8.56E-05 |
| CBL | 2.77 | 1.04E-05 |

| | | |
|---|---|---|
| COPB2 | 2.77 | 1.74E-08 |
| PTPN12 | 2.77 | 2.05E-16 |
| WSB2 | 2.77 | 1.01E-05 |
| AP1G1 | 2.76 | 1.66E-06 |
| PGD | 2.76 | 6.88E-20 |
| CYBB | 2.76 | 1.68E-33 |
| SRP68 | 2.76 | 2.45E-07 |
| SHKBP1 | 2.76 | 3.46E-08 |
| HSPA9 | 2.76 | 1.38E-19 |
| PSMD2 | 2.76 | 3.46E-08 |
| PGM3 | 2.75 | 1.69E-02 |
| FRG1 | 2.75 | 7.24E-04 |
| ALCAM | 2.75 | 1.58E-11 |
| ENTPD7 | 2.75 | 2.77E-02 |
| TBCA | 2.75 | 3.55E-04 |
| U2SURP | 2.75 | 1.03E-03 |
| CYTIP | 2.75 | 2.70E-05 |
| MORF4L2 | 2.75 | 3.98E-07 |
| E2F6 | 2.75 | 1.38E-02 |
| MSC | 2.74 | 7.69E-08 |
| HLA-DQB1 | 2.74 | 8.10E-14 |
| YIPF5 | 2.74 | 1.14E-02 |
| PEA15 | 2.74 | 2.45E-07 |
| ZBTB5 | 2.73 | 3.86E-02 |
| OCRL | 2.73 | 1.40E-02 |
| PELI3 | 2.72 | 5.28E-03 |
| WARS1 | 2.72 | 6.38E-12 |
| TRPC4AP | 2.71 | 6.96E-06 |
| SRSF10 | 2.71 | 9.40E-05 |
| FMR1 | 2.71 | 5.57E-03 |
| RHOQ | 2.70 | 2.38E-07 |
| ARF1 | 2.70 | 1.73E-12 |
| SEC23B | 2.70 | 2.29E-05 |
| CHUK | 2.70 | 1.27E-02 |
| ITGB1 | 2.70 | 2.87E-13 |
| ATP2B1 | 2.70 | 1.82E-15 |
| DEK | 2.69 | 1.80E-04 |
| HNRNPK | 2.69 | 4.60E-19 |
| MAFG | 2.69 | 1.22E-07 |
| SLC12A6 | 2.69 | 1.86E-09 |
| TMBIM6 | 2.69 | 6.78E-19 |
| RING1 | 2.69 | 1.65E-02 |
| ACTR3 | 2.69 | 2.41E-15 |
| SEC61A1 | 2.68 | 1.41E-12 |

| | | |
|---|---|---|
| SLC43A2 | 2.68 | 2.33E-19 |
| TMEM39A | 2.68 | 1.18E-03 |
| BROX | 2.67 | 2.92E-05 |
| IGBP1 | 2.67 | 2.37E-02 |
| TAX1BP1 | 2.67 | 1.64E-08 |
| RER1 | 2.67 | 4.48E-04 |
| FERMT3 | 2.66 | 2.29E-13 |
| GK | 2.66 | 1.90E-05 |
| SAMHD1 | 2.66 | 1.66E-07 |
| TOX4 | 2.66 | 2.87E-04 |
| RDX | 2.66 | 9.41E-06 |
| SRPRA | 2.66 | 9.40E-06 |
| SLC35B1 | 2.65 | 4.95E-03 |
| RNF144B | 2.65 | 4.00E-11 |
| COPG1 | 2.65 | 2.04E-09 |
| CIR1 | 2.65 | 9.31E-03 |
| TNC | 2.65 | 1.01E-04 |
| SLC16A6 | 2.64 | 5.07E-08 |
| PTPN11 | 2.64 | 3.99E-04 |
| CLINT1 | 2.64 | 3.60E-10 |
| SNN | 2.64 | 2.92E-05 |
| NBR1 | 2.64 | 1.06E-06 |
| GMPPB | 2.64 | 4.16E-03 |
| SLC44A1 | 2.64 | 9.22E-04 |
| BTG2 | 2.64 | 2.56E-16 |
| MAN2A1 | 2.63 | 5.82E-06 |
| PAPSS1 | 2.63 | 6.12E-04 |
| GORASP2 | 2.63 | 7.69E-06 |
| CD74 | 2.62 | 6.64E-53 |
| RTN3 | 2.62 | 9.24E-09 |
| ORAI1 | 2.62 | 5.77E-03 |
| DENND5A | 2.61 | 7.71E-24 |
| UHRF1BP1L | 2.61 | 5.93E-04 |
| SLC25A24 | 2.61 | 2.20E-03 |
| EPHX1 | 2.61 | 4.85E-04 |
| ATP6AP2 | 2.61 | 4.62E-04 |
| MAPRE1 | 2.61 | 1.79E-06 |
| RAP2C | 2.61 | 1.67E-05 |
| CDV3 | 2.60 | 5.48E-11 |
| PLA2G7 | 2.60 | 7.77E-13 |
| SRC | 2.60 | 6.65E-11 |
| STAC | 2.60 | 3.81E-04 |
| STRN3 | 2.60 | 8.99E-04 |
| PARP4 | 2.60 | 9.27E-08 |

| | | |
|---|---|---|
| NSUN3 | 2.60 | 3.68E-03 |
| SCAND1 | 2.60 | 2.93E-02 |
| DUSP6 | 2.60 | 1.49E-02 |
| TMEM123 | 2.60 | 1.90E-10 |
| POU2F2 | 2.59 | 3.97E-06 |
| RABGGTB | 2.59 | 4.30E-03 |
| HADHB | 2.59 | 1.83E-05 |
| PGAM1 | 2.59 | 3.39E-04 |
| LMAN1 | 2.59 | 1.03E-05 |
| FUOM | 2.59 | 4.74E-02 |
| KDELR2 | 2.59 | 3.06E-04 |
| LRP10 | 2.59 | 2.18E-07 |
| ARCN1 | 2.59 | 2.33E-06 |
| KCNE3 | 2.58 | 5.77E-03 |
| RAP1B | 2.58 | 3.51E-10 |
| USP15 | 2.58 | 6.73E-04 |
| DESI1 | 2.58 | 3.51E-04 |
| VSIG4 | 2.58 | 8.90E-04 |
| ACSL3 | 2.58 | 1.76E-07 |
| NUS1 | 2.58 | 2.83E-04 |
| RICTOR | 2.57 | 9.80E-04 |
| APLP2 | 2.57 | 1.88E-25 |
| ENPP2 | 2.57 | 3.81E-03 |
| FAM168B | 2.57 | 2.52E-04 |
| P2RX7 | 2.57 | 5.41E-16 |
| TMEM30A | 2.57 | 3.57E-06 |
| SDCBP | 2.56 | 4.68E-12 |
| ACO1 | 2.56 | 2.08E-04 |
| MIA3 | 2.56 | 7.43E-05 |
| TFG | 2.56 | 8.10E-05 |
| HSP90B1 | 2.56 | 5.78E-15 |
| ST13 | 2.56 | 2.78E-15 |
| NEU3 | 2.56 | 9.91E-03 |
| PTBP3 | 2.55 | 3.62E-07 |
| CLIC1 | 2.55 | 2.04E-08 |
| CST3 | 2.55 | 1.28E-14 |
| GBP1 | 2.55 | 2.30E-05 |
| RABGAP1 | 2.55 | 3.27E-03 |
| SEC24C | 2.55 | 4.85E-04 |
| AGFG1 | 2.55 | 5.38E-06 |
| HILPDA | 2.55 | 1.07E-02 |
| RNF24 | 2.54 | 2.21E-05 |
| UQCRH | 2.54 | 5.79E-04 |
| DNAJB11 | 2.54 | 1.10E-03 |

| | | |
|---|---|---|
| DOCK7 | 2.54 | 4.38E-04 |
| MAF1 | 2.54 | 6.33E-04 |
| LAMP2 | 2.54 | 1.46E-07 |
| ITPRID2 | 2.54 | 2.46E-06 |
| WASF2 | 2.54 | 1.12E-09 |
| SLC38A2 | 2.53 | 4.35E-18 |
| PABPC1 | 2.53 | 4.74E-20 |
| CCNI | 2.53 | 2.81E-15 |
| SPART | 2.53 | 9.35E-04 |
| SELENOT | 2.53 | 1.28E-03 |
| GTPBP2 | 2.53 | 1.60E-07 |
| RASSF5 | 2.53 | 1.44E-12 |
| LIPA | 2.52 | 9.04E-15 |
| RHOU | 2.52 | 3.56E-04 |
| MEGF9 | 2.52 | 4.54E-04 |
| MTPN | 2.52 | 2.17E-06 |
| MFAP3 | 2.51 | 3.52E-02 |
| KLHL5 | 2.51 | 1.40E-02 |
| TOMM70 | 2.51 | 6.01E-03 |
| GSTO1 | 2.51 | 2.26E-05 |
| AK4 | 2.51 | 2.66E-02 |
| DUSP5 | 2.51 | 1.14E-09 |
| OTUD4 | 2.51 | 4.12E-05 |
| LAMP1 | 2.51 | 8.17E-23 |
| HGS | 2.50 | 1.27E-10 |
| KPNB1 | 2.50 | 1.46E-09 |
| TNFRSF4 | 2.50 | 8.39E-04 |
| AMFR | 2.50 | 4.89E-05 |
| MYOF | 2.50 | 3.01E-13 |
| SECISBP2L | 2.50 | 7.40E-04 |
| RBBP8 | 2.50 | 3.59E-03 |
| TRAK2 | 2.50 | 1.94E-05 |
| PSEN1 | 2.49 | 1.32E-10 |
| TMED10 | 2.49 | 6.20E-07 |
| PSMC5 | 2.49 | 6.03E-03 |
| ARSB | 2.49 | 2.34E-03 |
| CCDC47 | 2.49 | 4.26E-05 |
| USO1 | 2.49 | 1.57E-05 |
| MBNL2 | 2.49 | 8.65E-04 |
| SGPP2 | 2.48 | 5.15E-04 |
| ARAP1 | 2.48 | 2.42E-24 |
| MT2A | 2.48 | 2.07E-02 |
| SQSTM1 | 2.48 | 1.06E-28 |
| TFPI | 2.48 | 3.15E-02 |

| | | |
|---|---|---|
| A2M | 2.48 | 1.81E-11 |
| ZC3H13 | 2.48 | 6.88E-05 |
| NOTCH2 | 2.48 | 2.12E-07 |
| PJA2 | 2.47 | 1.46E-05 |
| EIF3A | 2.47 | 2.62E-13 |
| MAGT1 | 2.47 | 3.74E-04 |
| SNX19 | 2.47 | 6.14E-04 |
| ADIPOR1 | 2.47 | 1.18E-07 |
| PSMD11 | 2.47 | 4.12E-05 |
| EGR2 | 2.46 | 5.24E-14 |
| AZIN1 | 2.46 | 2.77E-15 |
| HLA-DPA1 | 2.46 | 1.87E-25 |
| RAB8B | 2.46 | 1.44E-06 |
| UBE2K | 2.46 | 2.30E-03 |
| ANPEP | 2.46 | 3.54E-16 |
| PLEK | 2.46 | 7.00E-18 |
| SLC31A1 | 2.46 | 1.75E-03 |
| FXR2 | 2.46 | 1.34E-02 |
| BRPF3 | 2.46 | 6.45E-08 |
| MYDGF | 2.45 | 1.29E-02 |
| CTSS | 2.45 | 1.66E-16 |
| ZBTB7B | 2.45 | 2.81E-04 |
| RAB2A | 2.45 | 5.41E-03 |
| ELOVL7 | 2.45 | 4.04E-02 |
| NUP58 | 2.45 | 4.79E-07 |
| RNF185 | 2.45 | 1.72E-02 |
| EXOC1 | 2.45 | 3.41E-02 |
| UCHL1 | 2.45 | 1.82E-02 |
| ADA2 | 2.45 | 4.51E-10 |
| FLT1 | 2.44 | 3.08E-10 |
| SGTB | 2.44 | 1.41E-03 |
| EVI2B | 2.44 | 1.25E-03 |
| BPNT2 | 2.44 | 1.31E-02 |
| UFM1 | 2.43 | 5.36E-03 |
| ZNF438 | 2.43 | 3.51E-02 |
| MTHFD2 | 2.43 | 2.23E-14 |
| POLD3 | 2.43 | 4.43E-02 |
| USP10 | 2.43 | 4.30E-03 |
| GNS | 2.42 | 4.23E-07 |
| IQGAP1 | 2.42 | 1.67E-22 |
| CANX | 2.42 | 1.88E-24 |
| SPTLC2 | 2.42 | 2.59E-02 |
| VDAC3 | 2.42 | 4.23E-03 |
| SERINC3 | 2.42 | 6.94E-07 |

| | | |
|---|---|---|
| NOTCH3 | 2.41 | 1.58E-06 |
| PCMT1 | 2.41 | 1.60E-02 |
| RPS6KA3 | 2.41 | 2.79E-04 |
| ATP6V1B2 | 2.41 | 7.19E-13 |
| LPCAT2 | 2.41 | 1.07E-03 |
| CCNK | 2.41 | 4.14E-02 |
| FAR1 | 2.40 | 1.57E-03 |
| TM9SF2 | 2.40 | 1.11E-03 |
| MREG | 2.40 | 1.35E-02 |
| PDE7A | 2.40 | 1.41E-03 |
| AKIRIN1 | 2.40 | 1.78E-04 |
| PRELID3B | 2.40 | 1.43E-02 |
| SLC4A1AP | 2.39 | 2.75E-02 |
| DNTTIP2 | 2.39 | 7.59E-06 |
| OS9 | 2.39 | 9.55E-08 |
| TALDO1 | 2.39 | 2.43E-14 |
| TCEA1 | 2.39 | 3.51E-04 |
| ANKRD13C | 2.39 | 3.34E-02 |
| FGFRL1 | 2.39 | 5.20E-03 |
| OSER1 | 2.39 | 9.63E-05 |
| ANXA7 | 2.39 | 5.31E-03 |
| CHIC2 | 2.39 | 2.81E-02 |
| JAKMIP2 | 2.38 | 3.16E-06 |
| SFT2D2 | 2.38 | 9.69E-03 |
| DMXL2 | 2.38 | 1.32E-10 |
| MAMLD1 | 2.38 | 4.97E-03 |
| EMILIN2 | 2.38 | 3.82E-13 |
| SH3GLB1 | 2.38 | 4.10E-03 |
| GLA | 2.38 | 2.05E-04 |
| PPM1A | 2.38 | 1.66E-03 |
| PIK3C3 | 2.38 | 8.33E-03 |
| RPL6 | 2.38 | 2.13E-09 |
| VKORC1L1 | 2.37 | 4.02E-02 |
| VASP | 2.37 | 8.83E-07 |
| M6PR | 2.37 | 4.59E-06 |
| CCSER2 | 2.37 | 5.26E-05 |
| EIF5 | 2.37 | 1.45E-19 |
| TP53INP2 | 2.37 | 4.35E-05 |
| AHNAK | 2.37 | 2.10E-29 |
| PSMD14 | 2.36 | 4.15E-03 |
| PIM1 | 2.36 | 2.02E-15 |
| SHOC2 | 2.36 | 1.20E-03 |
| LASP1 | 2.36 | 6.78E-11 |
| INSIG1 | 2.36 | 1.46E-08 |

| | | |
|---|---|---|
| MMP19 | 2.36 | 1.66E-03 |
| ADRM1 | 2.35 | 9.06E-06 |
| OSTM1 | 2.35 | 8.48E-03 |
| RIT1 | 2.35 | 4.12E-05 |
| NUCKS1 | 2.35 | 1.52E-03 |
| TPM4 | 2.35 | 2.29E-09 |
| IPO7 | 2.35 | 1.09E-07 |
| CD55 | 2.35 | 1.63E-09 |
| FBXL3 | 2.35 | 1.06E-02 |
| SLC25A36 | 2.35 | 1.71E-03 |
| VDAC1 | 2.35 | 3.73E-07 |
| RELA | 2.35 | 4.26E-05 |
| RAC1 | 2.34 | 1.67E-05 |
| RPN2 | 2.34 | 1.81E-05 |
| ATP6V0C | 2.34 | 2.06E-02 |
| NR1D2 | 2.34 | 8.34E-04 |
| SSR1 | 2.34 | 2.07E-06 |
| VIM | 2.34 | 1.04E-42 |
| ERLEC1 | 2.34 | 2.21E-02 |
| IRAK3 | 2.34 | 2.85E-04 |
| GALC | 2.34 | 9.25E-03 |
| GNB1 | 2.34 | 2.61E-10 |
| CLIP1 | 2.33 | 4.65E-07 |
| MALT1 | 2.33 | 3.99E-03 |
| B4GALT5 | 2.33 | 1.52E-07 |
| BHLHE41 | 2.33 | 2.69E-09 |
| EIF4B | 2.33 | 3.31E-11 |
| ZNF746 | 2.33 | 6.15E-06 |
| SLC2A6 | 2.33 | 1.06E-09 |
| ARL6IP1 | 2.33 | 2.33E-05 |
| MARCHF6 | 2.32 | 8.63E-05 |
| CERT1 | 2.32 | 3.16E-06 |
| UBQLN1 | 2.32 | 5.75E-09 |
| ITGB2 | 2.32 | 2.31E-15 |
| S100A9 | 2.32 | 1.56E-07 |
| SAT2 | 2.32 | 4.26E-02 |
| UBA6 | 2.32 | 7.56E-05 |
| STX12 | 2.32 | 2.37E-03 |
| FTH1 | 2.31 | 3.20E-23 |
| PPP1CC | 2.31 | 4.17E-03 |
| DOT1L | 2.31 | 4.38E-12 |
| RAB21 | 2.31 | 4.83E-05 |
| SGK3 | 2.31 | 2.31E-02 |
| TMED4 | 2.31 | 2.77E-02 |

| | | |
|---|---|---|
| ZSWIM8 | 2.31 | 1.39E-06 |
| TFRC | 2.31 | 1.95E-19 |
| PLEKHB2 | 2.31 | 8.71E-10 |
| RNF139 | 2.31 | 2.21E-02 |
| SPP1 | 2.30 | 9.54E-09 |
| ADPGK | 2.30 | 1.15E-03 |
| ACTR2 | 2.30 | 5.00E-10 |
| SUCO | 2.30 | 1.21E-02 |
| RPL22 | 2.30 | 8.36E-03 |
| NUP98 | 2.30 | 3.02E-08 |
| FGR | 2.30 | 1.97E-15 |
| YME1L1 | 2.30 | 1.01E-05 |
| FCGRT | 2.30 | 5.63E-05 |
| SEC24A | 2.30 | 3.14E-03 |
| GDI2 | 2.29 | 5.98E-10 |
| MCFD2 | 2.29 | 2.05E-03 |
| NFE2L1 | 2.29 | 6.65E-09 |
| PSMD1 | 2.29 | 7.45E-06 |
| CAPNS1 | 2.29 | 1.84E-07 |
| RASA4B | 2.29 | 6.79E-03 |
| RPL9 | 2.29 | 5.18E-09 |
| SEL1L3 | 2.29 | 6.04E-03 |
| SPIRE1 | 2.29 | 5.33E-03 |
| SYVN1 | 2.28 | 5.21E-04 |
| ADH5 | 2.28 | 4.09E-02 |
| YTHDF3 | 2.28 | 2.00E-04 |
| NRDC | 2.28 | 8.53E-05 |
| ASAH1 | 2.28 | 1.44E-12 |
| BRWD3 | 2.28 | 4.59E-03 |
| HK2 | 2.28 | 3.12E-12 |
| CCDC93 | 2.28 | 2.00E-04 |
| TAF1D | 2.28 | 6.33E-04 |
| ITGAM | 2.28 | 5.52E-14 |
| ARL6IP5 | 2.27 | 1.11E-03 |
| ADIPOR2 | 2.27 | 1.22E-02 |
| MAPK1IP1L | 2.27 | 9.40E-06 |
| PICALM | 2.27 | 1.85E-09 |
| RAB14 | 2.27 | 2.58E-04 |
| MAP7D1 | 2.27 | 3.75E-12 |
| FMNL3 | 2.27 | 2.79E-13 |
| FXR1 | 2.26 | 7.98E-06 |
| AP2B1 | 2.26 | 2.70E-05 |
| CXCL1 | 2.26 | 7.61E-06 |
| NCOA1 | 2.26 | 2.52E-04 |

| | | |
|---|---|---|
| PAIP2 | 2.26 | 1.49E-02 |
| NECTIN2 | 2.26 | 7.93E-03 |
| GAA | 2.25 | 1.02E-11 |
| CAPN2 | 2.25 | 1.36E-08 |
| PSMD7 | 2.25 | 2.17E-02 |
| PPP2CA | 2.25 | 4.00E-03 |
| TWF1 | 2.25 | 1.22E-02 |
| RPS13 | 2.25 | 1.92E-07 |
| ENPP4 | 2.25 | 1.61E-02 |
| EEF1A1 | 2.25 | 7.32E-36 |
| MFSD12 | 2.24 | 7.24E-08 |
| H2AZ2 | 2.24 | 1.07E-02 |
| ATP5F1A | 2.24 | 1.08E-04 |
| TRIP12 | 2.24 | 5.68E-09 |
| SELENOS | 2.24 | 1.58E-02 |
| NEMP1 | 2.24 | 2.21E-02 |
| YARS1 | 2.24 | 2.73E-04 |
| TKT | 2.24 | 9.47E-20 |
| USP16 | 2.24 | 1.15E-02 |
| CLIC4 | 2.24 | 2.45E-11 |
| RIOK3 | 2.24 | 4.45E-03 |
| SPCS2 | 2.23 | 3.20E-02 |
| WDR1 | 2.23 | 2.85E-14 |
| RPL37A | 2.23 | 1.37E-09 |
| RAB22A | 2.23 | 4.43E-03 |
| PDIA6 | 2.23 | 2.21E-05 |
| SEC61B | 2.23 | 6.88E-03 |
| TOP2B | 2.23 | 1.25E-04 |
| BNIP2 | 2.23 | 7.13E-04 |
| ZFP91 | 2.23 | 2.59E-02 |
| TAOK1 | 2.23 | 6.95E-08 |
| PIP5K1C | 2.23 | 4.13E-10 |
| DNAJC5 | 2.23 | 3.92E-04 |
| MYL12B | 2.22 | 8.56E-05 |
| SBDS | 2.22 | 8.18E-03 |
| TMEM127 | 2.22 | 8.69E-07 |
| DPYSL2 | 2.22 | 2.43E-06 |
| PSTPIP2 | 2.22 | 1.52E-08 |
| ANXA1 | 2.22 | 6.31E-07 |
| STT3B | 2.22 | 8.60E-08 |
| CSDE1 | 2.22 | 4.02E-16 |
| KTN1 | 2.22 | 2.92E-06 |
| STX7 | 2.22 | 8.36E-03 |
| KPNA3 | 2.22 | 7.61E-03 |

| | | |
|---|---|---|
| HACD2 | 2.22 | 4.17E-02 |
| PSMB7 | 2.21 | 5.34E-03 |
| CHPF2 | 2.21 | 9.14E-03 |
| WBP11 | 2.20 | 7.85E-04 |
| ALOX5AP | 2.20 | 2.16E-04 |
| RLIM | 2.20 | 1.32E-04 |
| VAT1 | 2.20 | 5.38E-12 |
| PLIN2 | 2.20 | 3.99E-15 |
| VAMP3 | 2.20 | 8.95E-03 |
| DDOST | 2.20 | 3.00E-05 |
| ATF2 | 2.20 | 2.72E-03 |
| MFAP1 | 2.20 | 4.15E-02 |
| SIRPA | 2.20 | 3.47E-17 |
| UBXN7 | 2.20 | 6.39E-04 |
| CTNNA1 | 2.20 | 2.45E-07 |
| SETD5 | 2.20 | 2.97E-06 |
| PITPNA | 2.20 | 1.61E-02 |
| ADAM9 | 2.20 | 4.75E-06 |
| NLRC4 | 2.20 | 1.78E-02 |
| LPP | 2.19 | 3.22E-04 |
| UBR4 | 2.19 | 3.22E-19 |
| NDFIP1 | 2.19 | 4.56E-03 |
| MAML2 | 2.19 | 1.24E-04 |
| SPAG9 | 2.19 | 3.63E-09 |
| ZNF644 | 2.19 | 1.24E-02 |
| VAV1 | 2.19 | 1.23E-06 |
| SLC35F6 | 2.19 | 2.51E-02 |
| EFR3A | 2.19 | 4.67E-04 |
| MGAT4B | 2.19 | 1.45E-04 |
| PRDX1 | 2.19 | 1.93E-08 |
| MED13L | 2.19 | 3.12E-07 |
| SQOR | 2.19 | 7.81E-03 |
| ENO1 | 2.19 | 1.63E-13 |
| MED23 | 2.19 | 3.48E-02 |
| ZNF598 | 2.19 | 1.40E-02 |
| KCNE1 | 2.18 | 3.89E-02 |
| SUMO3 | 2.18 | 1.73E-02 |
| CDC37 | 2.18 | 1.10E-10 |
| PXDC1 | 2.18 | 2.03E-02 |
| PRDX3 | 2.18 | 5.77E-03 |
| EIF3L | 2.18 | 4.11E-03 |
| BLOC1S2 | 2.18 | 3.22E-02 |
| ARID5B | 2.18 | 1.84E-04 |
| STAT5B | 2.18 | 3.12E-04 |

| | | |
|---|---|---|
| CREB1 | 2.18 | 3.51E-03 |
| RB1CC1 | 2.18 | 5.34E-04 |
| PTPRC | 2.18 | 2.87E-11 |
| CLDND1 | 2.18 | 3.97E-02 |
| CANT1 | 2.17 | 6.08E-03 |
| C6orf62 | 2.17 | 5.61E-06 |
| SNX10 | 2.17 | 5.39E-03 |
| RPL35A | 2.17 | 2.39E-04 |
| KIF5B | 2.17 | 8.94E-08 |
| GOLGB1 | 2.17 | 2.80E-04 |
| TRIB3 | 2.17 | 1.61E-04 |
| PRPF8 | 2.17 | 4.75E-09 |
| SNX17 | 2.17 | 4.19E-03 |
| CALR | 2.17 | 1.38E-19 |
| NT5C2 | 2.17 | 3.60E-02 |
| PSMD4 | 2.17 | 1.75E-03 |
| FBXO7 | 2.16 | 3.65E-03 |
| EXOC4 | 2.16 | 3.17E-03 |
| DCUN1D1 | 2.16 | 2.31E-02 |
| PLD3 | 2.16 | 1.74E-27 |
| C15orf48 | 2.16 | 2.45E-05 |
| ELK1 | 2.16 | 4.39E-03 |
| FNDC3B | 2.16 | 5.33E-17 |
| SH3BGRL | 2.16 | 3.62E-07 |
| ELAVL1 | 2.16 | 4.80E-02 |
| TNIP2 | 2.16 | 7.31E-04 |
| AP3B1 | 2.16 | 1.45E-03 |
| MORF4L1 | 2.16 | 1.33E-05 |
| UBA52 | 2.15 | 1.24E-04 |
| IRF2 | 2.15 | 1.20E-02 |
| MYO1G | 2.15 | 4.11E-05 |
| RAB11B | 2.15 | 1.81E-02 |
| CTNNB1 | 2.15 | 8.18E-03 |
| CTSA | 2.15 | 1.30E-08 |
| SLU7 | 2.15 | 1.26E-03 |
| PTTG1IP | 2.15 | 3.85E-05 |
| CMPK1 | 2.15 | 2.85E-02 |
| PUM2 | 2.14 | 7.30E-05 |
| DNAJA2 | 2.14 | 2.61E-03 |
| RPL37 | 2.14 | 5.02E-12 |
| RPS4X | 2.14 | 2.71E-10 |
| TMCO1 | 2.14 | 1.63E-02 |
| MAP3K8 | 2.14 | 1.83E-05 |
| MED13 | 2.14 | 3.75E-06 |

| | | |
|---|---|---|
| RPS7 | 2.14 | 1.51E-04 |
| VDAC2 | 2.14 | 4.43E-03 |
| SARAF | 2.14 | 1.64E-07 |
| ATP5F1C | 2.13 | 2.79E-02 |
| USP8 | 2.13 | 4.16E-04 |
| VPS37C | 2.13 | 1.53E-02 |
| ARFGEF1 | 2.13 | 4.05E-05 |
| CNOT8 | 2.13 | 3.34E-02 |
| B4GALT1 | 2.13 | 4.81E-12 |
| RARS1 | 2.13 | 2.62E-02 |
| EIF4H | 2.13 | 9.46E-06 |
| RPL36AL | 2.13 | 1.07E-04 |
| CTDSP2 | 2.13 | 4.99E-04 |
| ALDH1A2 | 2.13 | 2.96E-03 |
| BCAT1 | 2.13 | 1.11E-03 |
| SMNDC1 | 2.13 | 2.04E-02 |
| TDG | 2.13 | 3.81E-02 |
| UBXN4 | 2.13 | 1.52E-04 |
| HS3ST3B1 | 2.12 | 8.48E-03 |
| PHF3 | 2.12 | 1.02E-05 |
| SLC6A6 | 2.12 | 5.01E-10 |
| RPS25 | 2.12 | 1.63E-04 |
| FLII | 2.12 | 1.04E-07 |
| RAD21 | 2.12 | 3.05E-04 |
| CYBA | 2.12 | 7.95E-12 |
| TPP1 | 2.12 | 2.40E-11 |
| ARL8B | 2.11 | 1.03E-07 |
| IGF2R | 2.11 | 2.19E-10 |
| MGLL | 2.11 | 2.69E-05 |
| FBXO42 | 2.11 | 1.58E-03 |
| AFF4 | 2.11 | 1.62E-08 |
| PPDPF | 2.11 | 9.66E-04 |
| TM9SF3 | 2.11 | 5.00E-04 |
| LRP1 | 2.11 | 1.16E-18 |
| CREG1 | 2.11 | 1.07E-08 |
| ILF2 | 2.11 | 5.21E-04 |
| LARP1 | 2.11 | 4.78E-06 |
| MAN2B1 | 2.11 | 5.66E-06 |
| MSN | 2.10 | 7.77E-13 |
| CNOT1 | 2.10 | 7.63E-07 |
| TLN1 | 2.10 | 1.14E-10 |
| EDEM3 | 2.10 | 1.15E-02 |
| ERGIC3 | 2.10 | 2.97E-03 |
| RPS6 | 2.10 | 4.01E-09 |

| | | |
|---|---|---|
| LARP4 | 2.10 | 8.48E-03 |
| CXCL3 | 2.10 | 4.05E-35 |
| FBXO38 | 2.10 | 4.38E-02 |
| DDX39B | 2.10 | 2.25E-03 |
| HECA | 2.10 | 4.25E-04 |
| TMOD3 | 2.10 | 2.91E-03 |
| MOB1A | 2.09 | 1.17E-04 |
| VCAN | 2.09 | 5.84E-05 |
| PTPN1 | 2.09 | 7.24E-05 |
| CSTA | 2.09 | 2.35E-02 |
| CSF2RB | 2.09 | 1.10E-03 |
| SNRNP200 | 2.09 | 2.20E-06 |
| TAOK3 | 2.09 | 1.14E-06 |
| TPT1 | 2.09 | 1.15E-16 |
| ACADVL | 2.09 | 9.37E-06 |
| ETV3 | 2.09 | 4.39E-04 |
| TXNDC11 | 2.09 | 5.19E-03 |
| EMP1 | 2.09 | 7.60E-05 |
| CELF1 | 2.08 | 5.26E-07 |
| RPL23 | 2.08 | 1.89E-08 |
| RAB6A | 2.08 | 1.80E-04 |
| TERF2IP | 2.08 | 4.37E-03 |
| LHFPL2 | 2.08 | 2.40E-06 |
| LIN7C | 2.08 | 3.48E-02 |
| HMBOX1 | 2.07 | 2.46E-03 |
| PTPN9 | 2.07 | 1.85E-02 |
| UBE2B | 2.07 | 4.37E-03 |
| EIF3H | 2.07 | 1.52E-03 |
| GAPDH | 2.07 | 2.65E-13 |
| MAPKAPK2 | 2.07 | 1.19E-03 |
| UHMK1 | 2.07 | 1.61E-03 |
| RAC2 | 2.07 | 1.74E-10 |
| NUP188 | 2.07 | 6.08E-06 |
| APEH | 2.07 | 8.33E-03 |
| ST3GAL2 | 2.07 | 5.94E-05 |
| NFAT5 | 2.06 | 7.59E-06 |
| FBXW11 | 2.06 | 3.01E-02 |
| CSNK1G3 | 2.06 | 4.38E-02 |
| CD209 | 2.06 | 4.36E-02 |
| ZHX2 | 2.06 | 2.44E-04 |
| METTL9 | 2.06 | 2.17E-04 |
| WDR43 | 2.06 | 1.79E-02 |
| ZFAND6 | 2.06 | 2.63E-02 |
| GOLGA5 | 2.06 | 1.41E-02 |

| | | |
|---|---|---|
| UBE2G1 | 2.06 | 1.91E-02 |
| CHCHD2 | 2.05 | 7.69E-04 |
| C5AR1 | 2.05 | 6.15E-10 |
| NR3C1 | 2.05 | 3.75E-06 |
| RPS21 | 2.05 | 1.68E-04 |
| ZFR | 2.05 | 1.89E-04 |
| WTAP | 2.05 | 1.53E-08 |
| TMX4 | 2.05 | 4.86E-02 |
| UBE2Q1 | 2.05 | 2.14E-02 |
| ACVR1 | 2.05 | 9.35E-03 |
| EDF1 | 2.04 | 1.77E-03 |
| E2F4 | 2.04 | 3.61E-02 |
| FBXO11 | 2.04 | 2.94E-03 |
| PDK1 | 2.04 | 4.60E-02 |
| RPL21 | 2.04 | 8.36E-05 |
| TMEM33 | 2.04 | 6.64E-04 |
| YWHAB | 2.04 | 1.21E-02 |
| AKIRIN2 | 2.04 | 4.08E-03 |
| XRCC6 | 2.04 | 5.51E-06 |
| ECPAS | 2.04 | 6.07E-05 |
| CD4 | 2.04 | 4.01E-03 |
| GID8 | 2.04 | 2.43E-02 |
| YPEL5 | 2.04 | 4.26E-03 |
| CD63 | 2.03 | 2.47E-07 |
| RAP2A | 2.03 | 9.60E-03 |
| SIPA1L2 | 2.03 | 1.79E-03 |
| WDR26 | 2.03 | 2.36E-08 |
| HLA-C | 2.03 | 1.27E-22 |
| HMGA1 | 2.03 | 3.54E-03 |
| CSNK1A1 | 2.03 | 4.87E-04 |
| DERL1 | 2.03 | 3.43E-02 |
| YIPF3 | 2.03 | 7.62E-03 |
| VIRMA | 2.03 | 3.84E-02 |
| GARS1 | 2.03 | 8.11E-05 |
| LIMS1 | 2.03 | 6.41E-06 |
| GPX4 | 2.03 | 1.09E-04 |
| RNF13 | 2.03 | 2.89E-06 |
| GAPVD1 | 2.03 | 6.01E-04 |
| ATP6V1E1 | 2.03 | 6.47E-04 |
| BOD1L1 | 2.02 | 2.20E-02 |
| RHBDD2 | 2.02 | 5.22E-03 |
| DIP2B | 2.02 | 5.19E-04 |
| NARS1 | 2.02 | 9.19E-03 |
| ATP6AP1 | 2.02 | 2.62E-08 |

| | | |
|---|---|---|
| MMP14 | 2.02 | 1.95E-03 |
| TRIM25 | 2.02 | 2.36E-06 |
| GRINA | 2.02 | 2.36E-07 |
| IP6K1 | 2.02 | 1.66E-03 |
| RBM3 | 2.02 | 4.23E-03 |
| TNPO1 | 2.02 | 1.20E-02 |
| RNPEP | 2.02 | 4.79E-03 |
| PIK3AP1 | 2.02 | 9.39E-10 |
| PAFAH1B2 | 2.02 | 1.71E-02 |
| SORT1 | 2.01 | 1.99E-06 |
| PPP2R2D | 2.01 | 1.61E-02 |
| RPSA | 2.01 | 6.78E-06 |
| OPA1 | 2.01 | 1.75E-02 |
| IBTK | 2.01 | 3.95E-04 |
| ZNF385A | 2.01 | 1.81E-02 |
| TMEM214 | 2.01 | 1.30E-03 |
| B2M | 2.01 | 1.17E-20 |
| MDM2 | 2.01 | 2.23E-04 |
| PTOV1 | 2.01 | 4.52E-02 |
| ARL8A | 2.01 | 1.73E-03 |
| RPL4 | 2.01 | 8.35E-07 |
| CAT | 2.01 | 2.16E-02 |
| SSU72 | 2.01 | 2.90E-03 |
| RNF145 | 2.00 | 1.02E-02 |
| PRCP | 2.00 | 4.42E-02 |
| FTL | 2.00 | 5.08E-33 |
| TMED9 | 2.00 | 6.58E-03 |
| PRPF3 | 2.00 | 5.48E-03 |
| HLX | 2.00 | 2.20E-02 |
| WDR48 | 2.00 | 1.28E-02 |
| TSPO | 2.00 | 5.83E-03 |
| STAU1 | 2.00 | 2.19E-03 |
| ADGRF3 | -2.00 | 3.78E-02 |
| MARCKSL1 | -2.00 | 1.11E-03 |
| EPHA2 | -2.00 | 3.15E-02 |
| FSCN1 | -2.00 | 3.33E-05 |
| PHLDB3 | -2.00 | 4.86E-02 |
| CNTN2 | -2.01 | 9.55E-03 |
| DCHS1 | -2.01 | 4.06E-02 |
| PTPRN2 | -2.01 | 2.06E-02 |
| IRF2BP2 | -2.01 | 3.35E-02 |
| MCF2L | -2.01 | 1.35E-03 |
| MYO7B | -2.01 | 4.65E-03 |
| PAX5 | -2.01 | 4.79E-02 |

| | | |
|---|---|---|
| DALRD3 | -2.02 | 2.20E-02 |
| PCBP3 | -2.02 | 4.68E-02 |
| SLIT1 | -2.02 | 1.55E-02 |
| PRAM1 | -2.02 | 1.66E-02 |
| CCDC12 | -2.03 | 2.20E-02 |
| DPYSL5 | -2.03 | 4.98E-02 |
| CELSR3 | -2.03 | 1.11E-02 |
| SPTB | -2.03 | 3.10E-03 |
| HSPA1A | -2.03 | 1.60E-17 |
| TOR3A | -2.03 | 2.65E-02 |
| CHIT1 | -2.04 | 3.19E-02 |
| RGS3 | -2.04 | 1.20E-03 |
| CARMIL3 | -2.04 | 4.34E-02 |
| RHOB | -2.04 | 8.02E-07 |
| PLEKHF2 | -2.05 | 1.78E-02 |
| MAST4 | -2.05 | 1.18E-02 |
| ALS2CL | -2.05 | 1.58E-02 |
| AXL | -2.06 | 3.78E-02 |
| KIF26B | -2.06 | 2.41E-02 |
| GIPC1 | -2.06 | 3.45E-02 |
| DISP3 | -2.06 | 2.10E-02 |
| RADIL | -2.06 | 1.46E-02 |
| FCHO1 | -2.06 | 8.89E-03 |
| TMC8 | -2.06 | 6.08E-03 |
| FLT4 | -2.07 | 6.23E-03 |
| ZBTB20 | -2.07 | 2.36E-02 |
| LAMA5 | -2.08 | 5.01E-04 |
| CXCR4 | -2.08 | 5.61E-03 |
| ETV1 | -2.08 | 3.18E-02 |
| FAM43A | -2.08 | 1.75E-02 |
| DNASE1 | -2.08 | 1.72E-02 |
| PDZD7 | -2.08 | 1.30E-02 |
| FER1L5 | -2.08 | 1.55E-02 |
| PRDM1 | -2.08 | 6.23E-04 |
| DOK2 | -2.09 | 1.15E-02 |
| DNAH17 | -2.09 | 6.50E-05 |
| GFRA2 | -2.09 | 4.47E-02 |
| B3GAT1 | -2.09 | 2.73E-02 |
| CECR2 | -2.09 | 3.15E-02 |
| TNXB | -2.09 | 3.11E-03 |
| TTC39A | -2.10 | 1.84E-02 |
| LAMB1 | -2.10 | 2.75E-02 |
| IL16 | -2.10 | 4.32E-03 |
| MORN1 | -2.10 | 8.23E-03 |

| | | |
|---|---|---|
| COL1A1 | -2.11 | 6.33E-03 |
| USP6 | -2.11 | 4.22E-03 |
| IQSEC2 | -2.11 | 1.96E-03 |
| CACNA1H | -2.11 | 1.17E-02 |
| ARMC9 | -2.11 | 5.07E-03 |
| CAMK2G | -2.11 | 4.79E-03 |
| PPFIA3 | -2.11 | 3.79E-02 |
| ZNF672 | -2.11 | 9.87E-03 |
| BOD1 | -2.12 | 3.47E-02 |
| AHNAK2 | -2.12 | 3.71E-03 |
| MAN1C1 | -2.12 | 1.86E-02 |
| PFAS | -2.12 | 3.63E-03 |
| SEMA3F | -2.12 | 4.82E-02 |
| TBCD | -2.13 | 1.31E-05 |
| KRI1 | -2.14 | 3.97E-02 |
| TIE1 | -2.14 | 3.01E-02 |
| ULBP2 | -2.14 | 4.97E-02 |
| MYLIP | -2.15 | 7.62E-03 |
| SHROOM2 | -2.15 | 2.71E-02 |
| SLA | -2.15 | 3.24E-03 |
| PPL | -2.15 | 1.78E-02 |
| PYGM | -2.15 | 1.54E-02 |
| SLC4A3 | -2.15 | 3.29E-02 |
| MYBPC2 | -2.16 | 2.36E-02 |
| NFIA | -2.16 | 1.78E-02 |
| ZFR2 | -2.16 | 1.42E-02 |
| PDE6B | -2.16 | 2.50E-02 |
| PLXDC1 | -2.16 | 4.98E-03 |
| MED29 | -2.16 | 2.88E-03 |
| GALK2 | -2.17 | 4.90E-02 |
| H2BC9 | -2.17 | 3.58E-02 |
| D2HGDH | -2.17 | 2.69E-02 |
| KLF4 | -2.17 | 6.49E-04 |
| MYH7 | -2.17 | 1.45E-03 |
| SOCS1 | -2.17 | 3.42E-06 |
| CDC14A | -2.18 | 3.86E-02 |
| MYOM1 | -2.18 | 2.66E-02 |
| OTOA | -2.18 | 4.86E-02 |
| VWA3A | -2.18 | 3.45E-02 |
| ANO7 | -2.18 | 9.18E-03 |
| DUOX2 | -2.18 | 7.31E-03 |
| GRIN1 | -2.19 | 2.86E-02 |
| CFAP74 | -2.19 | 1.04E-03 |
| FAHD2A | -2.19 | 4.60E-02 |

| | | |
|---|---|---|
| RAB26 | -2.19 | 3.45E-03 |
| PITPNM2 | -2.19 | 7.81E-03 |
| FAM20A | -2.20 | 1.78E-02 |
| LDB3 | -2.21 | 3.78E-02 |
| TGFA | -2.21 | 4.86E-02 |
| TRPM1 | -2.21 | 4.43E-02 |
| TAF4 | -2.21 | 1.24E-02 |
| SEPTIN4 | -2.21 | 4.74E-02 |
| ABCB9 | -2.21 | 1.61E-02 |
| ADCY1 | -2.22 | 3.40E-03 |
| TTYH1 | -2.22 | 4.24E-02 |
| MYRF | -2.22 | 2.64E-02 |
| NOS2 | -2.22 | 9.91E-03 |
| KCNQ2 | -2.23 | 2.44E-04 |
| GGT5 | -2.23 | 3.99E-02 |
| SNAP47 | -2.23 | 6.35E-03 |
| GRIN2D | -2.23 | 4.03E-02 |
| STARD13 | -2.23 | 2.35E-02 |
| MUC6 | -2.24 | 1.30E-02 |
| EVC | -2.24 | 1.94E-03 |
| ATP2B3 | -2.24 | 1.11E-02 |
| TIAM2 | -2.24 | 2.14E-02 |
| PIK3R3 | -2.25 | 3.66E-02 |
| TLL2 | -2.25 | 1.08E-02 |
| CHCHD7 | -2.25 | 4.72E-05 |
| GFPT2 | -2.25 | 3.47E-03 |
| H2AC13 | -2.25 | 2.53E-02 |
| ID3 | -2.25 | 4.94E-04 |
| GPRIN3 | -2.25 | 3.67E-03 |
| LRP5 | -2.25 | 2.28E-02 |
| CDH4 | -2.26 | 2.47E-02 |
| MLXIPL | -2.26 | 2.45E-02 |
| SAG | -2.26 | 9.42E-03 |
| WNK2 | -2.26 | 4.38E-04 |
| OSBPL5 | -2.26 | 1.99E-03 |
| SPOCD1 | -2.27 | 1.14E-02 |
| SLC4A11 | -2.27 | 1.53E-02 |
| CFAP46 | -2.27 | 2.48E-04 |
| CCL5 | -2.27 | 3.87E-05 |
| PLCB1 | -2.28 | 2.81E-02 |
| IL32 | -2.28 | 4.47E-02 |
| DNAH2 | -2.28 | 1.16E-06 |
| H3C13 | -2.28 | 1.40E-02 |
| TTYH2 | -2.28 | 1.51E-02 |

| | | |
|---|---|---|
| XXYLT1 | -2.29 | 7.81E-03 |
| AFF1 | -2.29 | 1.46E-04 |
| PADI3 | -2.29 | 3.51E-02 |
| CTTN | -2.29 | 4.51E-03 |
| SMPD3 | -2.29 | 3.74E-02 |
| BAIAP2 | -2.29 | 5.13E-06 |
| EXT1 | -2.29 | 5.19E-03 |
| SH3D21 | -2.30 | 2.43E-02 |
| INAVA | -2.30 | 1.81E-02 |
| SARDH | -2.30 | 8.62E-04 |
| TRPM5 | -2.30 | 7.60E-03 |
| SEMA6C | -2.30 | 2.73E-02 |
| SLC29A4 | -2.31 | 3.67E-02 |
| PRX | -2.31 | 1.88E-02 |
| HES1 | -2.31 | 1.05E-02 |
| XIRP1 | -2.31 | 4.34E-02 |
| HSPA1B | -2.32 | 1.04E-29 |
| LAD1 | -2.32 | 3.12E-02 |
| RNF207 | -2.32 | 2.83E-02 |
| LRIG1 | -2.32 | 2.33E-03 |
| TBC1D24 | -2.33 | 1.81E-03 |
| PLA2G4E | -2.33 | 4.55E-02 |
| MAT1A | -2.33 | 4.50E-02 |
| WDFY2 | -2.33 | 2.51E-02 |
| ALDH1A3 | -2.33 | 4.34E-02 |
| DHX58 | -2.33 | 3.27E-02 |
| TRERF1 | -2.33 | 2.08E-02 |
| ZAP70 | -2.33 | 2.40E-02 |
| EGR1 | -2.33 | 2.00E-08 |
| H3C11 | -2.34 | 1.66E-03 |
| FUT6 | -2.34 | 2.37E-02 |
| SUOX | -2.34 | 3.08E-02 |
| BTN2A2 | -2.35 | 3.37E-02 |
| COLQ | -2.35 | 3.08E-02 |
| ADCYAP1R1 | -2.35 | 3.65E-02 |
| NMRAL1 | -2.35 | 4.40E-02 |
| PLCH2 | -2.36 | 2.85E-03 |
| PLEKHA6 | -2.36 | 5.30E-03 |
| BSN | -2.36 | 2.60E-03 |
| ST8SIA5 | -2.36 | 7.11E-03 |
| BCL11A | -2.36 | 2.26E-05 |
| STOX2 | -2.37 | 2.69E-02 |
| TAB1 | -2.38 | 1.23E-02 |
| SLC22A23 | -2.38 | 7.60E-03 |

| | | |
|---|---|---|
| SEPTIN8 | -2.38 | 1.19E-02 |
| CATSPERD | -2.38 | 1.01E-02 |
| ANKRD24 | -2.39 | 1.64E-02 |
| NR1D1 | -2.39 | 1.66E-02 |
| CDKN1C | -2.39 | 8.36E-03 |
| ZNF589 | -2.39 | 3.02E-02 |
| FBXW8 | -2.40 | 3.01E-02 |
| TRIM2 | -2.40 | 6.08E-03 |
| STUM | -2.40 | 6.50E-03 |
| SLC43A1 | -2.40 | 5.22E-03 |
| PLK2 | -2.40 | 5.35E-09 |
| COL4A4 | -2.41 | 3.23E-03 |
| ASIC1 | -2.41 | 4.03E-02 |
| XKR7 | -2.41 | 3.41E-02 |
| PLEKHD1 | -2.41 | 3.48E-02 |
| NRARP | -2.42 | 1.17E-02 |
| GNLY | -2.42 | 3.51E-03 |
| PDE9A | -2.42 | 7.14E-03 |
| CHD5 | -2.42 | 3.75E-03 |
| CHRNA4 | -2.42 | 1.43E-02 |
| SIRT3 | -2.42 | 1.85E-02 |
| HEXD | -2.43 | 1.93E-02 |
| IGSF9 | -2.44 | 5.34E-04 |
| MVK | -2.44 | 9.19E-03 |
| HSPA2 | -2.44 | 1.11E-10 |
| LAMC2 | -2.44 | 3.15E-02 |
| HSH2D | -2.44 | 3.51E-02 |
| PBX4 | -2.44 | 4.56E-02 |
| GFAP | -2.44 | 3.15E-02 |
| ARHGEF37 | -2.44 | 1.24E-02 |
| ADAMTS15 | -2.45 | 3.01E-02 |
| KIFC2 | -2.45 | 4.16E-03 |
| VIL1 | -2.45 | 3.06E-02 |
| SLC30A3 | -2.46 | 3.47E-02 |
| TBX21 | -2.46 | 2.37E-02 |
| EPHB3 | -2.46 | 8.62E-03 |
| TICRR | -2.46 | 4.39E-02 |
| ALOX12B | -2.46 | 4.80E-02 |
| ZFP36 | -2.46 | 1.51E-15 |
| PAPLN | -2.46 | 1.28E-03 |
| PSEN2 | -2.47 | 2.59E-02 |
| HR | -2.47 | 9.56E-03 |
| CSKMT | -2.47 | 2.74E-06 |
| SLC22A8 | -2.48 | 4.47E-02 |

| | | |
|---|---|---|
| CASKIN1 | -2.48 | 1.03E-02 |
| COL9A3 | -2.48 | 4.16E-03 |
| CELF3 | -2.48 | 3.41E-02 |
| ST8SIA1 | -2.48 | 1.77E-02 |
| NOX5 | -2.48 | 1.75E-02 |
| ID1 | -2.49 | 1.95E-03 |
| FXYD2 | -2.49 | 3.01E-02 |
| UMODL1 | -2.49 | 4.50E-04 |
| OGDHL | -2.50 | 3.56E-03 |
| RORC | -2.50 | 4.57E-02 |
| TOP1MT | -2.51 | 1.48E-04 |
| TNK1 | -2.51 | 3.40E-02 |
| ZDHHC24 | -2.51 | 4.20E-02 |
| COQ4 | -2.51 | 2.36E-02 |
| MICAL3 | -2.51 | 4.33E-06 |
| PPP1R13L | -2.51 | 2.94E-02 |
| KATNB1 | -2.51 | 8.21E-03 |
| CNTNAP2 | -2.52 | 1.48E-02 |
| RGS9 | -2.52 | 5.34E-03 |
| MELTF | -2.52 | 1.57E-03 |
| ADAMTS13 | -2.52 | 5.01E-04 |
| ABHD5 | -2.52 | 3.36E-03 |
| PHACTR3 | -2.52 | 4.07E-02 |
| FLVCR1 | -2.53 | 4.31E-02 |
| SCN4A | -2.53 | 2.00E-03 |
| NDUFAF6 | -2.53 | 4.18E-02 |
| NLRC3 | -2.53 | 2.83E-03 |
| ADAMTS7 | -2.53 | 8.05E-05 |
| SH3RF3 | -2.53 | 8.43E-03 |
| XPNPEP2 | -2.53 | 4.11E-02 |
| PALM | -2.53 | 2.05E-02 |
| NPAS1 | -2.54 | 1.61E-02 |
| PAX3 | -2.54 | 4.90E-02 |
| PSD | -2.54 | 1.63E-03 |
| LRRC20 | -2.55 | 4.92E-02 |
| FCRLB | -2.55 | 1.81E-02 |
| APBB1 | -2.55 | 3.73E-02 |
| KAZALD1 | -2.55 | 4.03E-02 |
| LARGE2 | -2.56 | 4.04E-02 |
| DPF1 | -2.56 | 1.19E-02 |
| NLGN3 | -2.56 | 3.25E-02 |
| RNF165 | -2.56 | 5.98E-03 |
| RBFOX3 | -2.56 | 9.76E-04 |
| GCDH | -2.57 | 7.05E-03 |

| | | |
|---|---|---|
| CA11 | -2.58 | 2.29E-02 |
| RASGEF1C | -2.58 | 4.91E-02 |
| NTPCR | -2.58 | 8.54E-03 |
| DBH | -2.58 | 2.77E-02 |
| GAD1 | -2.58 | 4.04E-02 |
| SLC14A2 | -2.59 | 2.46E-02 |
| TMEM19 | -2.59 | 2.97E-02 |
| GAL3ST1 | -2.59 | 1.55E-02 |
| CDK18 | -2.59 | 4.15E-03 |
| KLHL35 | -2.60 | 9.45E-03 |
| CNIH3 | -2.60 | 1.82E-02 |
| EVPL | -2.60 | 1.83E-03 |
| ACADS | -2.60 | 3.15E-02 |
| EPS8L2 | -2.60 | 4.81E-03 |
| ZDHHC11 | -2.60 | 6.99E-03 |
| CILP | -2.61 | 3.51E-02 |
| KLHL30 | -2.62 | 4.18E-02 |
| PRMT9 | -2.62 | 3.90E-02 |
| ADCY6 | -2.63 | 2.28E-02 |
| SLC13A5 | -2.63 | 1.27E-02 |
| EPHX2 | -2.64 | 3.76E-02 |
| STX1B | -2.64 | 2.44E-02 |
| PKMYT1 | -2.64 | 3.60E-02 |
| ADAM8 | -2.65 | 7.58E-03 |
| CTH | -2.65 | 4.20E-02 |
| COMP | -2.65 | 2.93E-02 |
| TNNT2 | -2.65 | 1.91E-02 |
| FOSB | -2.65 | 1.44E-14 |
| TESC | -2.65 | 2.32E-02 |
| RIMS4 | -2.66 | 2.99E-02 |
| SCUBE1 | -2.66 | 2.29E-04 |
| HHIPL1 | -2.66 | 4.51E-03 |
| LRRC15 | -2.67 | 2.35E-02 |
| GATA4 | -2.68 | 1.95E-02 |
| A4GALT | -2.68 | 2.02E-02 |
| OASL | -2.68 | 6.60E-03 |
| HES4 | -2.68 | 4.15E-03 |
| KIFAP3 | -2.69 | 3.31E-02 |
| CAMSAP3 | -2.69 | 2.16E-02 |
| RGMA | -2.69 | 1.13E-05 |
| LBX2 | -2.70 | 2.36E-02 |
| TGFB1I1 | -2.70 | 3.58E-02 |
| ABHD11 | -2.70 | 4.85E-02 |
| SLC1A6 | -2.70 | 4.80E-02 |

| | | |
|---|---|---|
| RCL1 | -2.70 | 3.83E-02 |
| CD72 | -2.71 | 4.38E-02 |
| SOX4 | -2.71 | 6.58E-04 |
| DNAAF1 | -2.71 | 2.06E-02 |
| ARL4D | -2.72 | 2.50E-02 |
| TSTD3 | -2.72 | 1.13E-02 |
| OTOF | -2.73 | 3.86E-05 |
| ARMC2 | -2.73 | 1.86E-02 |
| TSGA10IP | -2.74 | 3.26E-02 |
| RDH10 | -2.74 | 7.80E-03 |
| TPO | -2.74 | 6.91E-04 |
| CATSPER1 | -2.74 | 6.41E-03 |
| PKP3 | -2.74 | 1.75E-02 |
| LSM10 | -2.74 | 2.54E-02 |
| OSM | -2.74 | 1.66E-02 |
| ELP6 | -2.74 | 2.94E-02 |
| NTHL1 | -2.75 | 4.73E-02 |
| ESPNL | -2.75 | 5.22E-03 |
| ICA1 | -2.76 | 3.75E-02 |
| C6orf226 | -2.77 | 3.66E-02 |
| GOLGA8A | -2.77 | 2.60E-02 |
| TENT5A | -2.77 | 8.98E-04 |
| GOLGA8B | -2.78 | 3.30E-02 |
| INSRR | -2.78 | 7.60E-03 |
| CHRNA2 | -2.78 | 1.38E-02 |
| MATN1 | -2.78 | 2.63E-02 |
| ZNF549 | -2.79 | 3.08E-02 |
| UHRF1 | -2.79 | 4.26E-03 |
| ASNS | -2.80 | 1.78E-02 |
| CHST10 | -2.80 | 4.86E-02 |
| DZIP1L | -2.81 | 2.47E-02 |
| LHX6 | -2.81 | 8.80E-03 |
| MAPK12 | -2.82 | 5.12E-04 |
| PAX2 | -2.82 | 2.14E-03 |
| SOBP | -2.83 | 1.28E-02 |
| SHQ1 | -2.83 | 4.80E-02 |
| TUBB3 | -2.84 | 6.38E-04 |
| SLC22A6 | -2.84 | 4.86E-02 |
| TRPM3 | -2.84 | 1.52E-02 |
| LIN9 | -2.84 | 1.29E-02 |
| POU6F1 | -2.84 | 9.61E-03 |
| RTEL1-TNFRSF6B | -2.84 | 2.54E-02 |
| STAP2 | -2.85 | 1.30E-02 |
| RPL3L | -2.85 | 3.08E-02 |

| | | |
|---|---|---|
| FSTL3 | -2.85 | 4.92E-02 |
| TBX1 | -2.86 | 1.25E-02 |
| ABCG8 | -2.86 | 6.52E-04 |
| MYCL | -2.86 | 2.40E-03 |
| NRG2 | -2.86 | 1.97E-02 |
| WDR54 | -2.86 | 3.36E-02 |
| MFSD4A | -2.87 | 2.13E-02 |
| ZSCAN2 | -2.87 | 3.94E-02 |
| SPNS3 | -2.87 | 1.29E-02 |
| HSPA6 | -2.87 | 8.75E-09 |
| KLHDC7B | -2.87 | 4.18E-02 |
| SHOX2 | -2.88 | 3.37E-02 |
| SLC22A14 | -2.88 | 2.53E-02 |
| IFNLR1 | -2.89 | 6.44E-04 |
| USP43 | -2.89 | 2.00E-02 |
| PARP16 | -2.90 | 9.24E-03 |
| ADGRD2 | -2.92 | 1.94E-03 |
| STOML1 | -2.92 | 1.10E-03 |
| TP53TG5 | -2.93 | 1.74E-02 |
| ANTXRL | -2.93 | 1.56E-02 |
| OGG1 | -2.93 | 8.95E-03 |
| NLGN2 | -2.93 | 8.68E-03 |
| H3C1 | -2.94 | 4.95E-03 |
| C11orf95 | -2.95 | 4.01E-02 |
| KAZN | -2.97 | 3.75E-04 |
| SYNGR1 | -2.97 | 5.50E-03 |
| KRBA1 | -2.97 | 8.59E-04 |
| SEZ6 | -2.98 | 1.01E-02 |
| ZNF844 | -2.98 | 1.08E-02 |
| PRR12 | -2.98 | 4.35E-05 |
| ODAD1 | -2.98 | 4.91E-03 |
| PRDM8 | -2.98 | 2.82E-04 |
| APOL4 | -2.99 | 2.78E-03 |
| CLIP3 | -3.00 | 5.62E-03 |
| PDE1C | -3.00 | 4.04E-02 |
| ALOX15 | -3.00 | 1.38E-02 |
| PRSS27 | -3.01 | 9.39E-03 |
| KHDC1 | -3.01 | 1.78E-02 |
| USP45 | -3.02 | 3.94E-02 |
| KLF7 | -3.03 | 1.20E-05 |
| SLC9A3R1 | -3.03 | 2.83E-04 |
| XKR5 | -3.03 | 2.95E-02 |
| ZNF710 | -3.03 | 8.28E-04 |
| PSTPIP1 | -3.04 | 5.31E-06 |

| | | |
|---|---|---|
| IL17REL | -3.05 | 2.03E-02 |
| P2RY2 | -3.05 | 8.51E-03 |
| NRSN2 | -3.05 | 1.86E-02 |
| ACP7 | -3.06 | 4.21E-02 |
| WFIKKN1 | -3.06 | 2.35E-02 |
| MAPK15 | -3.07 | 2.67E-02 |
| GAB4 | -3.08 | 4.37E-03 |
| TRIM47 | -3.09 | 1.73E-02 |
| ZNF568 | -3.09 | 4.52E-03 |
| PRDM13 | -3.09 | 3.41E-02 |
| DLX2 | -3.10 | 3.00E-05 |
| RBM43 | -3.10 | 1.41E-02 |
| RTKN | -3.10 | 2.07E-03 |
| SLC9A3 | -3.11 | 3.14E-02 |
| TRAIP | -3.11 | 1.29E-02 |
| DEGS2 | -3.13 | 3.30E-03 |
| POU2AF1 | -3.13 | 3.43E-02 |
| C16orf89 | -3.13 | 4.28E-02 |
| ATP1A3 | -3.14 | 4.62E-02 |
| PI16 | -3.14 | 1.29E-02 |
| KAT2A | -3.15 | 4.17E-03 |
| ARHGEF40 | -3.15 | 1.51E-02 |
| HCRTR1 | -3.18 | 2.17E-02 |
| TFAP2A | -3.18 | 1.41E-02 |
| DCLK2 | -3.18 | 9.92E-03 |
| TAF4B | -3.19 | 8.36E-05 |
| BICDL2 | -3.19 | 2.77E-02 |
| NTRK1 | -3.19 | 6.56E-04 |
| PRR15 | -3.20 | 1.93E-02 |
| PPP2R2C | -3.21 | 5.36E-05 |
| ENPP7 | -3.21 | 1.08E-02 |
| GIMAP6 | -3.22 | 8.30E-03 |
| ZNF569 | -3.23 | 2.48E-02 |
| H2BC5 | -3.24 | 3.41E-07 |
| LINGO1 | -3.24 | 2.55E-02 |
| AVPR2 | -3.27 | 3.74E-02 |
| AJUBA | -3.27 | 1.24E-02 |
| ADGRA1 | -3.30 | 5.61E-03 |
| ATOH8 | -3.31 | 9.19E-05 |
| GPC2 | -3.32 | 2.48E-02 |
| CATIP | -3.33 | 3.16E-02 |
| GPR183 | -3.33 | 1.03E-11 |
| TMEM132C | -3.35 | 1.89E-02 |
| SPNS2 | -3.35 | 3.34E-02 |

| | | |
|---|---|---|
| KRT33B | -3.36 | 4.47E-02 |
| PIP5KL1 | -3.37 | 6.38E-03 |
| PKN3 | -3.38 | 9.34E-05 |
| GNAI1 | -3.38 | 8.32E-03 |
| C17orf97 | -3.40 | 7.79E-03 |
| C1orf158 | -3.40 | 1.72E-02 |
| ORM1 | -3.41 | 9.17E-04 |
| SOHLH1 | -3.43 | 1.46E-02 |
| ADRA1A | -3.45 | 4.79E-02 |
| GPHN | -3.46 | 2.16E-03 |
| DPY19L4 | -3.47 | 3.47E-02 |
| PRR33 | -3.47 | 7.29E-03 |
| PYCR1 | -3.47 | 1.80E-02 |
| KBTBD13 | -3.48 | 3.96E-02 |
| ZNF470 | -3.48 | 1.89E-02 |
| MTFMT | -3.48 | 4.52E-02 |
| RALGDS | -3.49 | 6.00E-05 |
| PNMA8A | -3.50 | 4.79E-02 |
| USH1G | -3.51 | 1.81E-02 |
| SIK1B | -3.53 | 2.51E-02 |
| LRRN2 | -3.53 | 5.15E-03 |
| CBX2 | -3.54 | 1.82E-02 |
| PLAAT5 | -3.57 | 1.24E-02 |
| CLDN7 | -3.57 | 3.98E-02 |
| WNT3A | -3.58 | 1.48E-02 |
| HSD11B2 | -3.58 | 2.26E-03 |
| NPC1L1 | -3.59 | 6.13E-05 |
| SPATC1 | -3.59 | 4.28E-02 |
| WDCP | -3.59 | 2.26E-02 |
| C5orf63 | -3.64 | 3.22E-02 |
| GPRC5B | -3.64 | 3.75E-04 |
| TOMM34 | -3.66 | 9.67E-03 |
| GPR162 | -3.68 | 1.80E-02 |
| NUTM2D | -3.68 | 1.71E-02 |
| KCNQ5 | -3.69 | 2.21E-02 |
| TEX45 | -3.69 | 1.51E-02 |
| TNFRSF8 | -3.70 | 3.26E-03 |
| RGR | -3.72 | 1.81E-02 |
| GPER1 | -3.72 | 2.44E-03 |
| BRINP2 | -3.72 | 7.78E-03 |
| RAB42 | -3.72 | 1.20E-02 |
| PLD4 | -3.73 | 9.68E-03 |
| UNC5C | -3.73 | 3.99E-02 |
| RFLNA | -3.74 | 1.63E-02 |

| | | |
|---|---|---|
| CYP2A6 | -3.75 | 4.04E-02 |
| ONECUT2 | -3.77 | 4.16E-02 |
| TBX2 | -3.78 | 9.54E-04 |
| RORB | -3.78 | 2.63E-02 |
| DUOXA1 | -3.79 | 1.39E-02 |
| HHIPL2 | -3.82 | 1.20E-02 |
| SOAT2 | -3.83 | 1.78E-02 |
| MKS1 | -3.84 | 1.55E-03 |
| CELA2A | -3.87 | 1.05E-02 |
| GPRIN2 | -3.88 | 2.43E-04 |
| ARL9 | -3.91 | 4.97E-03 |
| BEAN1 | -3.91 | 1.91E-05 |
| CCNF | -3.92 | 4.26E-03 |
| LRRC45 | -3.92 | 1.23E-03 |
| CLBA1 | -3.94 | 1.04E-03 |
| P2RX6 | -4.00 | 6.33E-03 |
| MPP6 | -4.03 | 4.28E-02 |
| H2AW | -4.04 | 1.57E-03 |
| TUBA4B | -4.05 | 6.66E-03 |
| HS3ST3A1 | -4.06 | 2.43E-02 |
| DACH1 | -4.10 | 3.81E-02 |
| PTGDR2 | -4.13 | 2.17E-02 |
| CASKIN2 | -4.13 | 3.61E-04 |
| CABP1 | -4.14 | 1.30E-02 |
| PNMA6F | -4.15 | 1.39E-02 |
| PLA2G4F | -4.18 | 2.48E-04 |
| CRABP2 | -4.19 | 5.28E-03 |
| ACR | -4.21 | 2.22E-02 |
| CDX2 | -4.23 | 1.13E-02 |
| UNC5CL | -4.26 | 9.53E-03 |
| GATA5 | -4.26 | 4.58E-03 |
| PLPP7 | -4.27 | 5.20E-04 |
| LPAR5 | -4.28 | 2.19E-02 |
| H1-5 | -4.28 | 4.50E-05 |
| THEM5 | -4.29 | 2.31E-02 |
| NYAP1 | -4.30 | 1.12E-02 |
| KPNA5 | -4.31 | 6.08E-03 |
| GAREM2 | -4.32 | 1.78E-03 |
| TGFB2 | -4.33 | 4.02E-04 |
| DLX1 | -4.36 | 7.95E-04 |
| CA5A | -4.37 | 1.24E-02 |
| RBPJL | -4.39 | 5.22E-05 |
| ANO4 | -4.44 | 9.86E-03 |
| NTN3 | -4.48 | 2.71E-03 |

| | | |
|---|---|---|
| SIGIRR | -4.49 | 1.33E-04 |
| UNC5A | -4.51 | 7.89E-03 |
| NUTM2F | -4.53 | 7.81E-03 |
| SIM1 | -4.54 | 3.61E-03 |
| TBX18 | -4.54 | 1.21E-02 |
| CTRB1 | -4.56 | 1.78E-02 |
| GPR150 | -4.59 | 4.65E-03 |
| NEUROD2 | -4.74 | 1.80E-02 |
| VAX1 | -4.78 | 1.63E-02 |
| KDF1 | -4.81 | 8.48E-03 |
| SPATA31E1 | -4.84 | 1.68E-03 |
| SHC4 | -4.85 | 1.98E-03 |
| ANKRD33 | -4.85 | 1.40E-02 |
| RRAD | -4.93 | 1.81E-02 |
| FAM90A1 | -5.02 | 5.04E-03 |
| ELAC1 | -5.18 | 1.42E-02 |
| SYDE1 | -5.47 | 5.72E-03 |
| SAMD10 | -5.61 | 5.14E-03 |
| SP8 | -5.70 | 2.94E-02 |
| MRGPRE | -6.00 | 1.96E-03 |
| SYNC | -6.01 | 4.08E-03 |
| CTSW | -6.35 | 2.53E-04 |
| TAFA5 | -6.38 | 9.66E-04 |
| SCHIP1 | -6.74 | 3.02E-03 |
| POU3F1 | -7.15 | 2.71E-03 |
| C3orf22 | -7.17 | 1.17E-04 |
| CCDC102A | -7.43 | 2.43E-06 |
| CHRNG | -7.45 | 4.13E-04 |
| UCP1 | -8.10 | 7.41E-04 |
| APOA1 | -8.10 | 6.77E-45 |
| CLDND2 | -13.50 | 4.14E-06 |

**Table S14. KEGG pathway analyses of significantly altered genes upon HDL-m1A-tDR-ArgACG-1-21 delivery.**

Table S14. KEGG pathway analyses of significantly altered genes upon HDL-m1A-tDR-ArgACG-1-21 delivery.

| KEGG Pathway | Description | In Pathway | In Data | Expect | FC | FDR |
|---|---|---|---|---|---|---|
| hsa04141 | Protein processing in endoplasmic reticulum | 165 | 52 | 16.66 | 3.12 | 4.89E-12 |
| hsa04145 | Phagosome | 152 | 44 | 15.34 | 2.87 | 6.89E-09 |
| hsa05140 | Leishmaniasis | 74 | 27 | 7.47 | 3.61 | 1.03E-07 |
| hsa05134 | Legionellosis | 55 | 21 | 5.55 | 3.78 | 2.27E-06 |
| hsa04216 | Ferroptosis | 40 | 17 | 4.04 | 4.21 | 6.05E-06 |

| | | | | | | |
|---|---|---|---|---|---|---|
| hsa05323 | Rheumatoid arthritis | 90 | 25 | 9.09 | 2.75 | 9.04E-05 |
| hsa05152 | Tuberculosis | 179 | 38 | 18.07 | 2.10 | 2.80E-04 |
| hsa04142 | Lysosome | 123 | 28 | 12.42 | 2.25 | 1.08E-03 |
| hsa05145 | Toxoplasmosis | 113 | 26 | 11.41 | 2.28 | 1.43E-03 |
| hsa04940 | Type I diabetes mellitus | 43 | 14 | 4.34 | 3.23 | 1.43E-03 |
| hsa05321 | Inflammatory bowel disease (IBD) | 65 | 18 | 6.56 | 2.74 | 1.43E-03 |
| hsa04612 | Antigen processing and presentation | 77 | 20 | 7.77 | 2.57 | 1.43E-03 |
| hsa04658 | Th1 and Th2 cell differentiation | 92 | 22 | 9.29 | 2.37 | 2.20E-03 |
| hsa05120 | Epithelial cell signaling in Helicobacter pylori infection | 68 | 18 | 6.86 | 2.62 | 2.20E-03 |
| hsa05146 | Amoebiasis | 96 | 22 | 9.69 | 2.27 | 3.73E-03 |
| hsa05169 | Epstein-Barr virus infection | 201 | 37 | 20.29 | 1.82 | 4.06E-03 |
| hsa04979 | Cholesterol metabolism | 50 | 14 | 5.05 | 2.77 | 5.67E-03 |
| hsa05142 | Chagas disease (American trypanosomiasis) | 102 | 22 | 10.30 | 2.14 | 7.75E-03 |
| hsa04670 | Leukocyte transendothelial migration | 112 | 23 | 11.31 | 2.03 | 1.15E-02 |
| hsa04659 | Th17 cell differentiation | 107 | 22 | 10.80 | 2.04 | 1.39E-02 |
| hsa03050 | Proteasome | 45 | 12 | 4.54 | 2.64 | 1.89E-02 |
| hsa03060 | Protein export | 23 | 8 | 2.32 | 3.45 | 1.89E-02 |
| hsa05416 | Viral myocarditis | 59 | 14 | 5.96 | 2.35 | 2.49E-02 |
| hsa05332 | Graft-versus-host disease | 41 | 11 | 4.14 | 2.66 | 2.49E-02 |
| hsa00600 | Sphingolipid metabolism | 47 | 12 | 4.74 | 2.53 | 2.49E-02 |
| hsa04657 | IL-17 signaling pathway | 93 | 19 | 9.39 | 2.02 | 2.54E-02 |
| hsa04144 | Endocytosis | 244 | 39 | 24.63 | 1.58 | 2.84E-02 |
| hsa04933 | AGE-RAGE signaling pathway in diabetic complications | 99 | 19 | 9.99 | 1.90 | 4.81E-02 |
| hsa04062 | Chemokine signaling pathway | 189 | 31 | 19.08 | 1.62 | 4.81E-02 |
| hsa04066 | HIF-1 signaling pathway | 100 | 19 | 10.10 | 1.88 | 4.93E-02 |
| hsa05132 | Salmonella infection | 86 | 17 | 8.68 | 1.96 | 4.93E-02 |
| hsa04218 | Cellular senescence | 160 | 27 | 16.15 | 1.67 | 4.93E-02 |
| hsa04510 | Focal adhesion | 199 | 32 | 20.09 | 1.59 | 4.93E-02 |
| hsa05166 | Human T-cell leukemia virus 1 infection | 255 | 39 | 25.74 | 1.52 | 4.93E-02 |

**Table S15. Oligonucleotide sequences**

| **Name** | **Sequence** | **Type** | **Source** | **Corresponding Figure** | **Notes** |
|---|---|---|---|---|---|
| mmu Il6 primer - forward | GCTACCAAACTGGATATAATCAGGA | DNA | IDT | Fig. 1D | |
| mmu Il6 primer - reverse | CCAGGTAGCTATGGTACTCCAGAA | DNA | IDT | Fig. 1D | |
| mmu Il1b primer - forward | AGTTGACGGACCCCAAAAG | DNA | IDT | Fig. 1D | |
| mmu Il1b primer - reverse | AGCTGGATGCTCTCATCAGG | DNA | IDT | Fig. 1D | |

| | | | | | |
|---|---|---|---|---|---|
| mmu Tnfa primer - forward | CAGGGTACCGTTTTCCGAGGGTTGAATGAG | DNA | IDT | Fig. 1D | |
| mmu Tnfa primer - reverse | CAGCTCGAGGTCTTTTCTGGAGGGAGATGTG | DNA | IDT | Fig. 1D | |
| mmu Nr4a3 primer - forward | CAAAGAGAGTGGAGGAGCTATG | DNA | IDT | Fig. 1D | |
| mmu Nr4a3 primer - reverse | CAGTGAGTGTGGAAGGTGTT | DNA | IDT | Fig. 1D | |
| mmu Tmem123 primer - forward | GGGCCAGGTTTAACAAAGAAAG | DNA | IDT | Fig. 1D | |
| mmu Tmem123 primer - forward | TCAGTTGGACCCGATGTAAAG | DNA | IDT | Fig. 1D | |
| m1A_PCR primer-forward | CTAATACGACTCACTATAGGGCGGAGACGGTCGGG | DNA | IDT | Fig.4F | ArgACG-1 tRNA forward primer |
| m1A_PCR primer-reverse | GCGGAGCCCACACTCTACTCGACAGATACGAATAT | DNA | IDT | Fig.4F | ArgACG-1 tRNA reverse primer |
| tDR-ArgACG-1-21 | GUUCGACUCCUGGCUGGCUCG | RNA | Gene link | Fig.5A-F | Synthetic 21 nt ArgACG tRNA oligo |
| m1A-tDR-ArgACG-1-21 | GUUCG(m1A)CUCCUGGCUGGCUCG | RNA | Gene link | Fig.5A-F | Synthetic 21 nt m1A-ArgACG tRNA oligo |
| hsa IL1B primer - forward | TACCTGTCCTGCGTGTTGAA | DNA | IDT | Fig.5C,6A-C | |
| hsa IL1B primer - reverse | TCTTTGGGTAATTTTTGGGATCT | DNA | IDT | Fig.5C,6A-C | |
| hsa TNFA primer - forward | AGCCCATGTTGTAGCAAACC | DNA | IDT | Fig.5C,6A-C | |
| hsa TNFA primer - reverse | TCTCAGCTGGACGCCATT | DNA | IDT | Fig.5C,6A-C | |
| hsa TMEM123 primer - forward | GAAACCTACAGCGGCATCTAA | DNA | IDT | Fig.5C,6A-C | |
| hsa TMEM123 primer - reverse | ACCAACAAAGCTCCCAGTATC | DNA | IDT | Fig.5C,6A-C | |
| hsa IL6 primer - forward | ACCTAGAGTACCTCCAGAACAG | DNA | IDT | Fig.5C | |
| hsa IL6 primer - reverse | ACTGCATAGCCACTTTCCATTA | DNA | IDT | Fig.5C | |
| hsa IL12B primer - forward | TAAGATGCGAGGCCAAGAATTA | DNA | IDT | Fig.5C | |
| hsa IL12B primer - reverse | GCAGAATGTCAGGGAGAAGTAG | DNA | IDT | Fig.5C | |
| tDR-ArgACG-1-24 | GUUCGACUCCUGGCUGGCUCGCCA | RNA | Gene link | Fig.6A-C | Synthetic 24 nt ArgACG |

| | | | | | |
|---|---|---|---|---|---|
| | | | | | tRNA oligo |
| m1A-tDR-ArgACG-1-24 | GUUCG(m1A)CUCCUGGCUGGCUCGCCA | RNA | Gene link | Fig.6A-C | Synthetic 24 nt m1A-ArgACG tRNA oligo |
| m1G RNA (neg cntl) | GGAGACGGUCGGGUCCAGAUAUUCGUAUCUGUC | RNA | IDT | Fig.S4A-B | Synthetic unmodified (m1G) broccoli RNA |
| m1G RNA | GGAGACGGUCGGGUCCA(m1G)AUAUUCGUAUCUGUC | RNA | IDT | Fig.S4A-B | Synthetic m1G broccoli RNA |
| m1A RNA (neg cntl) | GGAGACGGUCGGGACCAGAUAUUCGUAUCUGUC | RNA | Gene link | Fig.S4A-B | Synthetic unmodified (m1A) broccoli RNA |
| m1A RNA | GGAGACGGUCGGG(m1A)CCAGAUAUUCGUAUCUGUC | RNA | Gene link | Fig.S4A-B | Synthetic m1A broccoli RNA |
| RT primer | GCGGAGCCCACACTCTACTCGACAGATACGAATAT | DNA | IDT | Fig.S4A-B | RT primer |
| RT-PCR-forward | CTAATACGACTCACTATAGGGCGGAGACGGTCGGG | DNA | IDT | Fig.S4A-B | RT-PCR-forward |
| RT-PCR-reverse | GCGGAGCCCACACTCTACTCGACAGATACGAATAT | DNA | IDT | Fig.S4A-B | RT-PCR-reverse |
